# Network analysis of mouse brain proteomes resolves molecular and cellular endophenotypes of aging and Aβ pathology

**DOI:** 10.64898/2026.11.05.764669

**Authors:** Sruti Rayaprolu, Sneha Malepati, Eric B. Dammer, Kyla Langdon, Rashmi Kumari, Amanda D. Brandelli, Claudia Espinosa-Garcia, Duc D. Duong, Upasna Srivastava, Prateek Kumar, Ali Tfaily, Jiahui Chen, Xin Zhou, Yinlong Li, Ahmad F. Chaudhary, Steven Liang, Nicholas T. Seyfried, Srikant Rangaraju

## Abstract

Aging and Alzheimer’s disease (AD) pathology exert both shared and distinct effects on the brain, yet their relative contributions to molecular alterations remain poorly defined. Here we performed tandem mass tag proteomics across the lifespan in wild-type and 5xFAD mice and integrated these data with human brain proteomic networks to resolve aging- and amyloid-β (Aβ)-associated molecular programs. We quantified 8,535 proteins and identified 51 co-expression modules representing neuronal, mitochondrial, glial, vascular and proteostatic pathways. Aging emerged as the predominant driver of proteomic remodeling, with widespread alterations in RNA metabolism, protein translation, and neuronal pathways occurring independently of Aβ pathology. In contrast, Aβ selectively modified a subset of age-dependent processes, including suppression of mitochondrial ATP biosynthetic machinery and amplification of astrocytic and microglial activation states. These key aging and disease-related molecular signatures were highly conserved in human AD brain proteomes. Together, our findings establish aging as the primary determinant of brain proteome organization while revealing specific mechanisms through which Aβ pathology reshapes age-associated biological processes. Our results also highlight molecular mechanisms of human aging and AD pathology that can be modeled in mouse models.

## INTRODUCTION

Alzheimer’s disease (AD) is among the most common neurodegenerative disorders worldwide, affecting 6.9 million individuals over the age of 65 in the United States alone, and represents a growing public health challenge as the population ages (Alzheimer’s Association 2026 Facts and Figures)^1,2^. Although extracellular amyloid-beta (Aβ) plaques and intracellular neurofibrillary tangles composed of hyperphosphorylated tau are defining neuropathological features of AD^3–5^, the molecular mechanisms that drive disease onset and progression remain poorly understood. Aging is the strongest risk factor for late-onset AD^2,6^, increasing susceptibility to neurodegeneration through widespread changes to synaptic, glial, vascular, metabolic, and proteostatic mechanisms^7–9^. Many of these processes are also implicated in the development of AD, suggesting that the disease progression may arise through complex interactions between normal aging and pathological processes^10^. Consequently, distinguishing molecular changes associated with aging from those specifically driven by AD pathology remains a major challenge in the field.

Large-scale proteomic studies of post-mortem human brain tissue have consistently identified coordinated changes in synaptic, mitochondrial, immune, vascular and proteostasis-related pathways and have revealed highly reproducible co-expression networks associated with disease progression (Dammer et al., 2015; Ping et al., 2018; Johnson et al., 2020; Wingo et al., 2019; Higginbotham et al., 2020). These studies have provided important insights into the molecular architecture of AD; however, delineating aging-associated mechanisms from AD-associated mechanisms remains difficult because most analyses rely on post-mortem brain tissue obtained at end stage of disease when neurodegeneration and neuronal loss are already widespread^11^. Furthermore, interpretation of human post-mortem studies is confounded by the prolonged and heterogenous nature of disease progression, as well as the presence of mixed pathologies and age-related comorbidities. While longitudinal studies integrating ante-mortem cognitive, biofluid and imaging measures with post-mortem brain and biofluid-derived “omics” data^12,13^ address these limitations, disentangling the relative contributions of aging and AD pathology to molecular alterations in the human brain remains a major challenge.

Mouse models of AD pathology provide an opportunity to address these limitations by enabling longitudinal assessment of molecular changes during again and disease progression. Among the available amyloidosis models, the 5xFAD transgenic mouse model is one of the most widely used and extensively characterized models of Aβ pathology. 5xFAD mice express five familial early-onset AD mutations in *APP* and *PSEN1*^14,15^, resulting in dysfunction of APP protein processing, accelerated Aβ42 production, progressive Aβ plaque accumulation, robust glial activation, and progressive neuritic pathology^14,16^. While 5xFAD mice do not fully recapitulate human AD pathology, particularly tau aggregation and extensive neurodegeneration, they provide a powerful system for investigating the how aging and Aβ pathology independently and jointly reshape the brain proteome over time.

In this study, we performed tandem-mass tag mass spectrometry (TMT-MS) across the lifespan of 5xFAD and wild-type (WT) mice to disentangle the proteomic changes associated with aging from those driven by Aβ pathology. Using weighted protein co-expression network analysis, we identified protein modules representing distinct cellular and biological processes and characterized their relations with age, genotype, and Aβ levels. We further integrated mouse and human AD brain proteomic networks to identify conserved molecular signatures associated with aging, Aβ pathology, and their interaction. Together, these analyses provide a comprehensive, time-resolved view of brain proteomic molecular alterations and highlight age-dependent signatures that are either preserved or uniquely altered in the context of Aβ-associated neurodegeneration.

## RESULTS

### Network analysis of brain proteomes resolves modules associated with aging, amyloidosis, cell type and disease-related mechanisms

We performed TMT-MS on 86 cortical brain homogenates (16-plex, 6 batches including global internal standards per batch) from WT and 5xFAD mice at various ages (1.8 to 14 months) and quantified 8,535 proteins after data normalization and outlier removal (**Fig. 1A, Supplementary Table 1**). We also performed bulk RNA sequencing from all 86 mice and identified 21,829 mRNA transcripts (**Supplementary Table 5**). Compared to WT mice, 5xFAD mice showed age-dependent progressive increase in protein levels of human Aβ42, along with increased protein levels of key markers implicated in AD pathogenesis: Apoe, Gfap, Clu, Trem2, C1qa, Itm2b, and Mdk (**Fig. 1B**). The number of differentially expressed proteins (DEPs) between 5xFAD and WT mice increased with age, starting at 6 months of age (**Fig. 1C, Supplementary Table 3**). Of the DEPs, 16 proteins including App, Psen1, Ncstn, and Apbb1 were consistently differentially expressed at all ages (**Fig. 1D**). Variance partitioning analysis demonstrated that age accounted for the largest proportion of variance in the brain proteome, followed by genotype, while the effect of TMT-plex batch was successfully removed by batch correction (**Fig. 1E**). GSVA of proteomic data also showed how AD pathology impacted the brain proteome in an age dependent manner, including activation of immune pathways, antigen presentation and lysosomal mechanisms, and suppression of mitochondrial respiratory function, translation and synaptic pathways (**Supplementary Fig 1**). The 8,535 proteins were then used to build a protein co-expression network using the weighted gene co-expression network analysis (WGCNA) algorithm^17^. This resolved the proteome into 51 modules (annotated as M1-M51), which were further grouped into six distinct clusters (**Fig. 1F**). Gene ontology (GO) analysis was used to determine the biology over-represented in each module as well as in each of the six clusters (**Fig. 1F, Supplemental Data 3**). Cluster 1 modules were enriched in proteins involved in RNA metabolism, splicing, translation, and the post-synaptic compartment. Cluster 2 modules included neuronal/synaptic, ion channel, signal transduction, endosomal, and DNA transcription proteins. Cluster 3 modules included axonal, nuclear and lipid transport, mitochondrial, and gene transcription proteins. Cluster 4 modules included neuronal and mitochondrial proteins. Cluster 5 modules primarily included mitochondria, glycolysis, metabolism, and myelin sheath proteins. Lastly, cluster 6 modules included immune/defense response, extracellular matrix, lysosome, ER/Golgi, and cholesterol transport related proteins.

**Figure 1.**
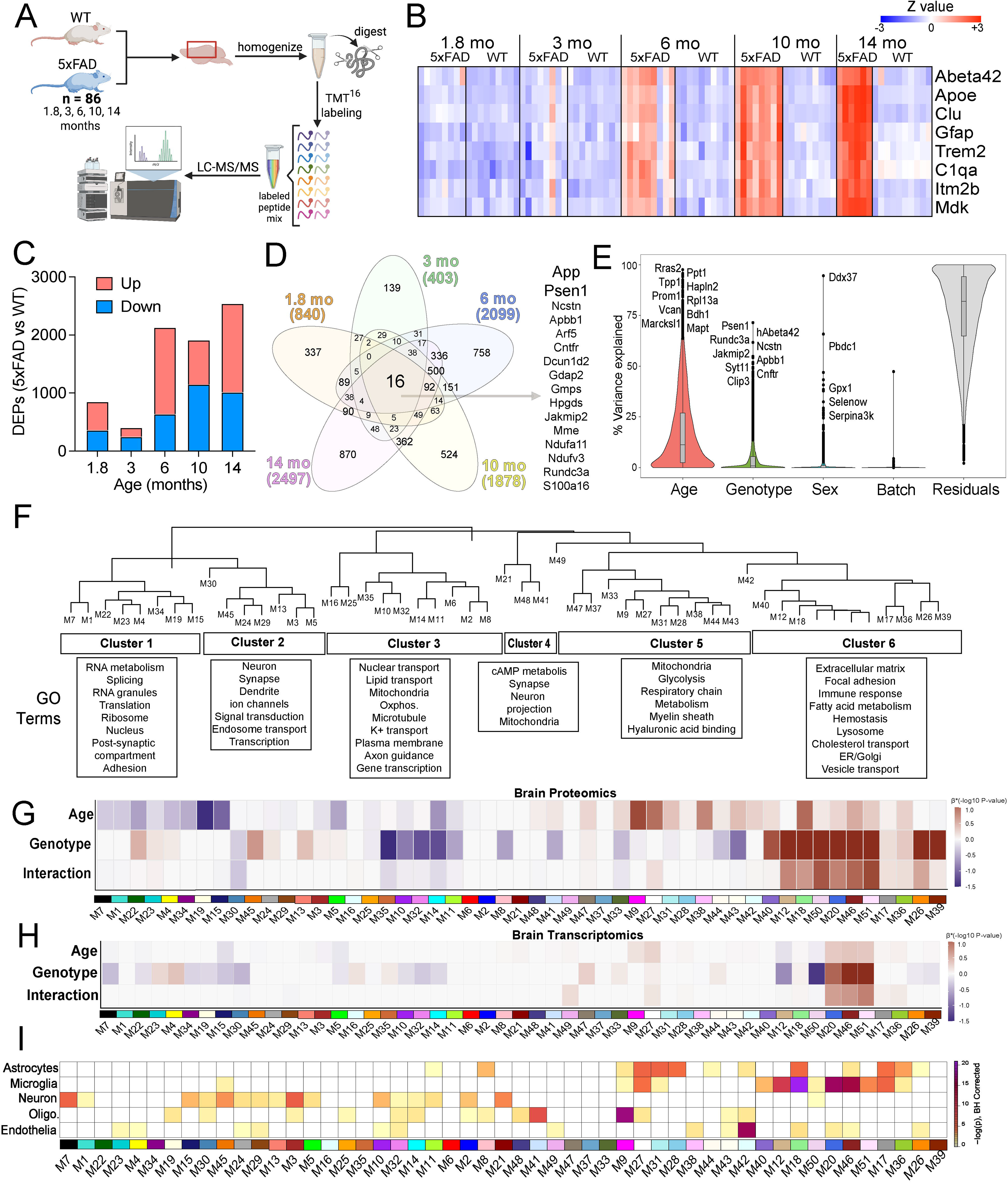
TMT-MS profiling of the aging 5xFAD mouse brain resolves 51 protein co-expression modules. (A) Study design. Brain tissue from 86 5xFAD and wild-type (WT) mice aged 1.8, 3, 6, 10 and 14 months was homogenized, digested, labelled with TMT-16plex reagents and analyzed by LC-MS/MS. (B) Z-scored abundance of amyloid- and glia-associated proteins in individual mice, grouped by age and genotype. (C) Number of differentially expressed proteins (5xFAD versus WT) at each age, increased (red) and decreased (blue). (D) Overlap of differentially expressed proteins across the five ages; the 16 proteins altered at every age are listed. (E) Percentage of variance in protein abundance explained by age, genotype, sex, batch and residual terms; proteins with the largest contribution to each term are labelled. (F) Dendrogram of the 51 co-expression modules, grouped into six clusters. Boxes list representative enriched gene-ontology terms for each cluster. (G) Module-trait relationships in the brain proteome. Color denotes the signed association strength (beta x -log_10_ P) of each module eigenprotein with age, genotype and the age x genotype interaction. (H) Module-trait relationships in the brain bulk RNA-seq profiles from the same mice. A synthetic eigengene was constructed for each proteomic module, using proteomic module members that were present in the bulk RNAseq data. (I) Cell-type marker enrichment of each module (one-sided Fisher exact test; color, -log_10_ BH-adjusted P).

Given that age and genotype showed the largest effect on variance in the dataset (**Fig. 1E**), we next examined module-trait relationships for the proteome (**Fig. 1G**) and transcriptome (**Fig. 1H**). In general, we found that Cluster 5 and 6 modules increased with age while Cluster 1 and 2 modules decreased with age (**Fig. 1G**). Several age-related modules in Cluster 6 (extracellular matrix, immune response) also increased with genotype (**Fig. 1G**; e.g., M12, M18, M20) while modules M26 and M39 showed a strong association with age only (**Fig. 1G**). At the transcriptome level, only three of Cluster 6 modules show strong age and genotype associations – M20, M46, M51 (**Fig. 1H**), while two modules showed negative trait association – M12, M20 (**Fig. 1H**). Discordant associations with age and amyloidosis were observed in Cluster 1 (RNA metabolism/splicing) and few Cluster 5 (mitochondria, glycolysis) modules (**Fig. 1H**). In general, the transcriptome showed weaker associations with module traits (**Fig. 1H**) compared to the proteome, suggesting that aging- and amyloidosis-associated biological processes are more robustly reflected at the proteome level. Lastly, we performed cell type enrichment analysis using reference mRNA and protein marker lists^18,19^ to evaluate whether the modules capture molecular features of specific brain cell types (**Fig. 1I**). We found that majority of modules in Cluster 2 and 3 were enriched in neuronal markers while Cluster 5 and 6 were enriched in astrocyte markers. Microglial and myeloid markers were enriched in Cluster 6 modules, particularly M18, M20 and M46. These microglia-enriched modules also showed enrichment of astrocytic markers, suggesting shared glial mechanisms or co-expression patterns that cannot be further resolved without cell type-specific approaches. Endothelial proteins were specifically enriched in module M42. Lastly, oligodendrocyte markers were enriched in modules M9 and M41. Among amyloidosis associated modules, Cluster 6 modules showed age-amyloidosis interactions, where aging effects were magnified by amyloidosis, a feature apparent at both proteomic and transcriptomic levels. Together, these analyses reveal that aging and amyloidosis exert both shared and distinct effects on the brain proteome.

### Age-dependent impairment of RNA metabolism and protein translation occurs independently of amyloidosis

Modules within Cluster 1 (**Fig. 2A**) showed enrichment in proteins involved in RNA and protein metabolism, including gene transcription (M7), RNA splicing (M1), initiation of protein translation (M22), translation elongation (M23, M4) and proteasome function (M23, including Psmb2) (**Fig. 2B**). In general, all Cluster 1 modules showed decreased levels with aging, suggestive of age-related dysfunction in these pathways (**Fig. 2B**). With the exception of module M7, none of the Cluster 1 modules showed cell type enrichment patterns (**Fig. 1G**). M7 showed enrichment of neuronal/synaptic proteins along with RNA biosynthesis/transcription proteins. This suggests that age-related dysfunction in RNA splicing and all aspects of protein synthesis and proteasomal protein degradation occur across all brain cell types, while age-related dysfunction in gene transcription may be preferentially occurring in neurons. In contrast to strong age-related effects on Cluster 1 modules, amyloidosis had no effect on RNA biosynthesis, RNA splicing, or translation elongation modules. Interestingly, amyloidosis increased levels of translation initiation and proteasomal modules, which were more apparent at ages above 6 months, when amyloid plaque deposition is diffusely present. This finding suggests a preferential vulnerability of protein translation and degradation mechanisms to beta amyloid pathology.

**Figure 2.**
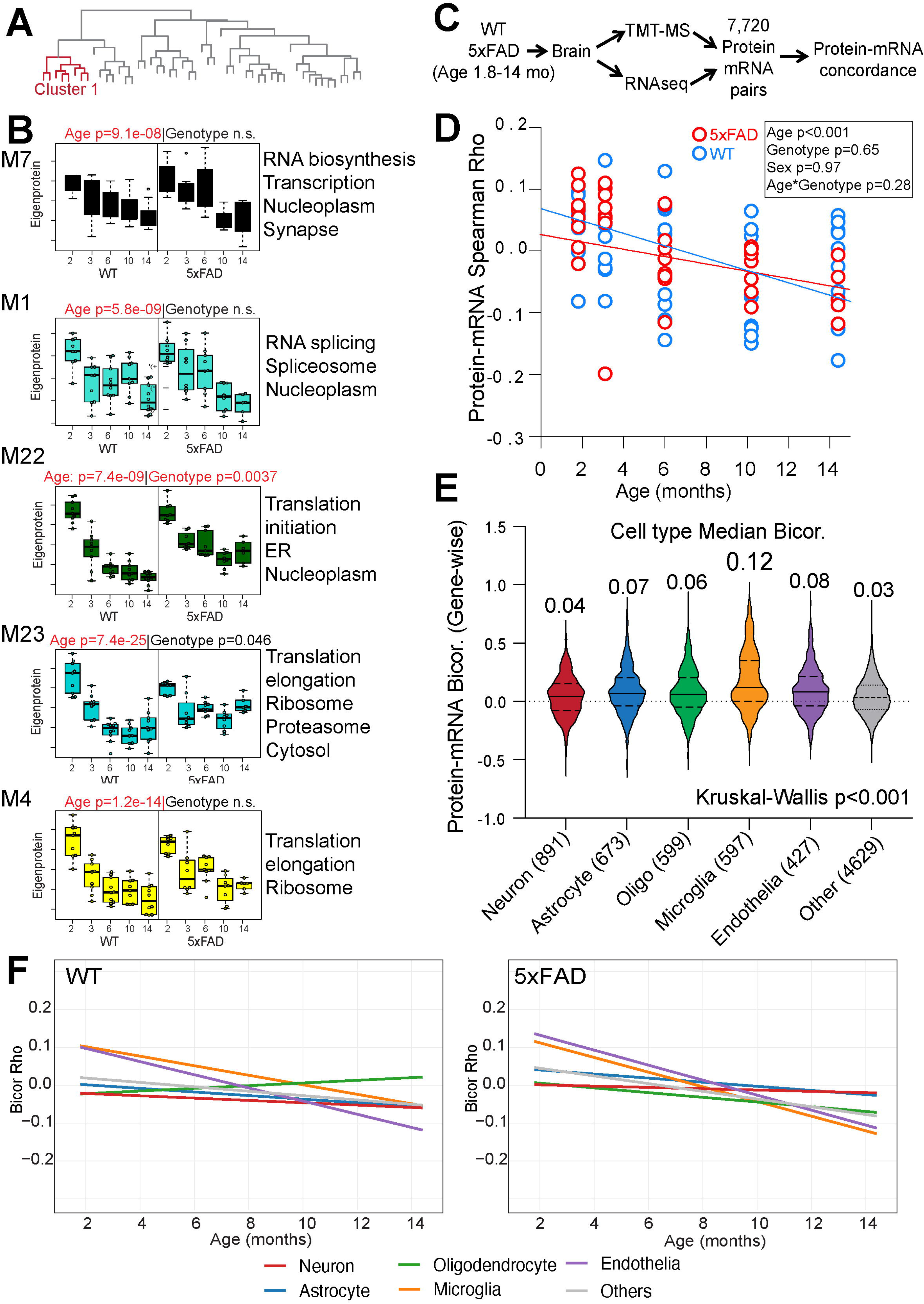
Cluster 1 modules track chronological age independently of genotype, in parallel with an age-related loss of protein-mRNA concordance. (A) Module dendrogram with Cluster 1 highlighted in red. (B) Eigenprotein values of Cluster 1 modules (M7, M1, M22, M23, M4) across age in WT and 5xFAD mice. P values for the age and genotype terms are given above each module (n.s. = not significant); representative GO terms on the right. (C) Paired proteomic and transcriptomic profiling of the same brains yielded 7,720 protein-mRNA pairs for concordance analysis. (D) Per-sample protein-mRNA concordance (Spearman rho) versus age in WT (blue) and 5xFAD (red) mice. Concordance declines with age (P < 0.001) and is unaffected by genotype, sex or their interaction with age. (E) Gene-wise protein-mRNA correlation (biweight midcorrelation) by cell-type class; the number of cell type-specific genes per class is in parentheses and the median rho is printed above each violin (Kruskal-Wallis P < 0.001). (F) Age trajectories of gene-wise protein-mRNA correlation for each cell-type class, shown separately for WT and 5xFAD mice.

Since our brain proteomic analysis suggests that aging is associated with progressive dysfunction in RNA metabolism and proteostasis, we hypothesized that this age-dependent dysfunction should also impact protein and mRNA concordance in an age-dependent manner. Protein-mRNA concordance was assessed across 7,720 matched protein-mRNA pairs quantified in both proteomic and RNA sequencing datasets was assessed (**Fig, 2C**). Across all 86 samples, the overall protein-mRNA concordance was low **(Supplementary Fig. 2A, R^2^=0.003**) consistent with known low-to-modest correlations between protein and mRNA, particularly in the brain^20,21^. We observed a significant age-dependent decline in protein–mRNA concordance (**Fig 2D**, p < 0.01). In contrast. neither genotype (p = 0.65), sex (p = 0.97), nor age-by-genotype interaction (p = 0.28) significantly influenced concordance (**Fig 2D**), indicating that transcript–protein discordance arises primarily as a consequence of aging rather than amyloidosis. Proteins that had highest positive protein-mRNA concordance (e.g., C1qa, C1qb, Ctss) were enriched in GO terms related to complement component C1q complex and microglial cell activation (**Supplementary Fig. 2B**). Conversely, proteins that had negative protein-mRNA concordance (e.g., Thy 1, Ncan, Rsph4a) were enriched in GO terms such as GDP-dissociation inhibitor binding, growth cone membrane, and regulation of action potential firing (**Supplementary Fig. 2B**). Next, we evaluated protein-mRNA concordance among brain cell types and found that it varied significantly across cell types (Kruskal–Wallis, p < 0.001), with microglia exhibiting the highest median protein–mRNA bicorrelation (bicor = 0.12), followed by endothelial (0.08), astrocytic (0.07), and oligodendrocytic (0.06) markers, whereas neuronal proteins displayed the weakest transcript–protein correspondence (bicor = 0.04) (**Fig. 2E**). Furthermore, the age-dependent decrease in protein-mRNA discordance was consistently observed across all cell types (**Supplementary Fig. 3**). Amyloidosis did not affect protein-mRNA discordance in neurons, astrocytes, oligodendrocytes or endothelial cells but exhibited a distinct effect in microglia (**Fig. 2F**).

To summarize, our analyses reveal proteomic signatures of age-dependent dysfunction in RNA transcription, RNA splicing, protein translation, and protein degradation that are associated with a progressive increase in protein-mRNA discordance in the brain. This aging induced discordance is observed across all major brain cell types and, importantly, is not affected by amyloidosis.

### Identifying differential effects of aging and beta amyloidosis on neurons, neuronal sub-populations and excitatory-inhibitory balance

Clusters 2 and 3 are enriched for neuronal markers (**Fig. 1E**); however, showed divergent association with amyloidosis (**Fig. 1G**). Thus, we asked whether this divergence reflects differential responses of two major neuronal subclasses – excitatory (glutamatergic) or inhibitory (GABAergic) neurons – to amyloidosis. Thus, Clusters 2 and 3 were assessed for pan-excitatory and pan-inhibitory cell-type enrichment (Fisher’s exact test (FET)) modules using reference single-nucleus transcriptomic datasets of mouse neurons^22,23^.

Cluster 2 modules were predominantly enriched in excitatory neuronal markers, while Cluster 3 modules showed preferential enrichment in inhibitory neuronal markers (**Fig. 3A**). M3 (glutamatergic transmission, post-synaptic density), M13 (synaptic vesicle ATPase, synapse/dendrite), M24 (calcium transport, neuronal spine), M45 (synaptic vesicle, including Sv2b, syntaxin binding) and M29 (post-synaptic density, protein Ser/Thr phosphatase complex) were the excitatory neuron-enriched modules in Cluster 2 (**Fig. 3B**). Specifically, M3 and M24 showed age-dependent decreased levels without effects of amyloidosis (**Fig. 3B right panel**, ANOVA p = 7.9e-05 M3; p = 0.055 M24). Excitatory modules M45 and M13 showed an elevated abundance in 5xFAD mice as compared to WT mice (**Fig. 3B**, p = 0.0042 M13; p = 0.00026 M45), independent of age. M8, M10 and M11 were three Cluster 3 modules enriched in inhibitory neuronal markers (**Fig. 3C**). Inhibitory module M8 (neuron projection, neutral amino acid transport) exhibited an age-dependent increase that was blunted by amyloidosis (**Fig. 3C right panel,** ANOVA p = 0.031 M8). M10 (neuron projection, microtubule complex) also showed age-dependent increase but displayed a significant decrease in the presence of amyloidosis (**Fig. 3C right panel,** ANOVA p = 9.5e-08 M10). In contrast, M11 showed age-dependent decline that was further accelerated by amyloidosis (**Fig. 3C right panel,** ANOVA p = 5.0e-07 M11). Within Cluster 3, we were unable to further resolve modules to specific GABAergic interneuron subtypes (e.g., Pvalb, Sst, Vip interneurons) using available transcriptomic atlases ^22,23^. Nevertheless, some modules contained canonical markers of interneuron classes. For example, M41 contained hub proteins of PV interneurons, including Pvalb, while M32 contained Kcnc1 and Ank1, established markers of fast-spiking GABAergic interneurons^23^. Heatmap representation of module eigenprotein levels across all ages showed excitatory neuron modules (M7, M15, M24, M45, M3, M13) with early and sustained upregulation in 5xFAD relative to WT, whereas inhibitory neuron modules (M10, M11, M8) exhibited a pattern of progressive downregulation in 5xFAD mice that deepened with age, most prominently from 6 months onward (**Fig. 3D**).

**Figure 3.**
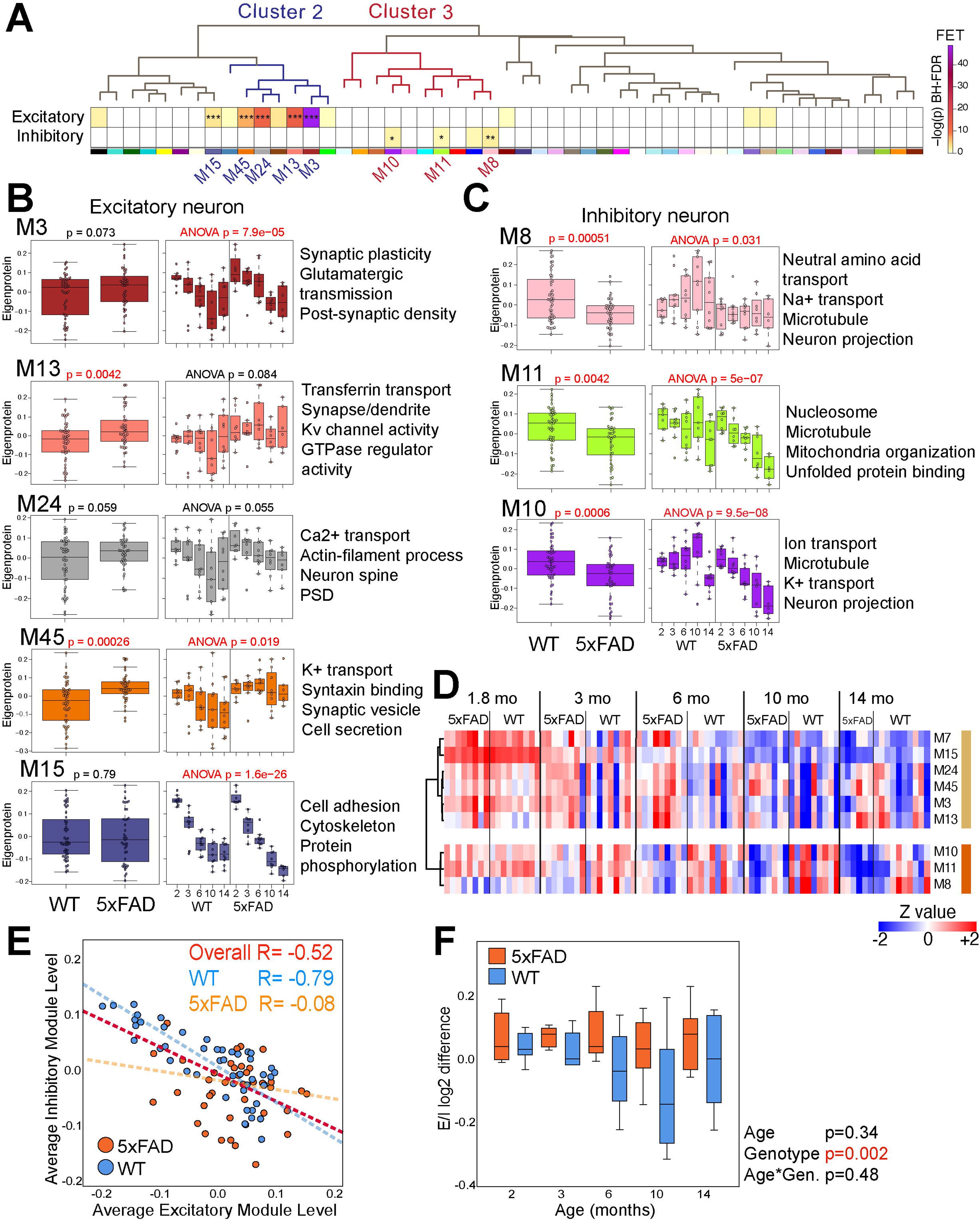
Excitatory and inhibitory neuronal modules follow distinct age trajectories, and their reciprocal coupling is lost in 5xFAD mice. (A) Module dendrogram with Cluster 2 (blue) and Cluster 3 (red). Heatmaps below show enrichment of excitatory and inhibitory neuron markers in each module (Fisher exact test; color, -log_10_ BH-adjusted P; asterisks indicate significant modules). (B) Excitatory neuron modules (M3, M13, M24, M45, M15). For each module: eigenprotein by genotype (left) and by age within WT and 5xFAD (right, ANOVA P). Representative GO terms listed on the right. (C) Inhibitory neuron modules M8, M11 and M10. For each module: eigenprotein by genotype (left) and by age within WT and 5xFAD (right, ANOVA P). Representative GO terms listed on the right. (D) Z-scored eigenprotein values of the excitatory and inhibitory modules in individual mice, grouped by age and genotype. (E) Mean inhibitory versus mean excitatory module eigenprotein for each mouse. The inverse relationship seen in WT mice (R = −0.79) is absent in 5xFAD mice (R = −0.08); overall R = −0.52. (F) Proteomic estimation of excitatory/inhibitory balance (log_2_ difference between mean excitatory and mean inhibitory module levels) by age and genotype. Genotype P = 0.002; age P = 0.34; age x genotype P = 0.48.

We further assessed relationships between excitatory and inhibitory modules across genotype (WT and 5xFAD mice). Across all animals, excitatory and inhibitory module levels were negatively correlated (**Fig. 3E**; Spearman R = −0.52). This negative correlation was substantially stronger in WT mice (**Fig. 3E**; R= −0.79) although this was absent in 5xFAD mice (**Fig. 3E**; R = −0.08). This implies an inverse relationship between excitatory and inhibitory neurons under homeostatic conditions, which is disrupted by amyloidosis. To quantify this imbalance directly, we estimated excitatory/inhibitory (E/I) ratio at the proteomic level for each animal across the ages and genotypes (**Fig. 3F**). There was a significant effect of genotype (p = 0.002), with 5xFAD mice displaying consistently elevated E/I ratios relative to WT controls, whereas age alone was not a significant predictor (p = 0.34) and there was no significant age-by-genotype interaction (p = 0.48). In an independent cohort of 9-10 month old WT and 5xFAD mice, we performed synaptic vesicle 2A (SV2A) positron emission tomography (PET) imaging to assess synaptic density (**Supplementary Fig. 4**). At the level of whole brain, we found no significant differences between 5xFAD and WT mice without accounting for sex differences. However, lower SV2A tracer binding was observed in female 5xFAD compared to WT females across all brain regions with the exception of the cerebellum, a region known to be devoid of Aβ plaques in 5xFAD mice (**Supplementary Fig. 4C**). We also performed behavioral studies in an independent cohort of 9-10 month old 5xFAD and WT mice, using both spatial (Morris water maze) and associate learning (contextual and cued fear conditioning). While 5xFAD and WT mice performed similarly on the Morris Water maze platform, 5xFAD mice showed impairments in contextual fear conditioning paradigms (**Supplementary Fig. 5).** Taken together, we conclude that the proteomic phenotype of E/I imbalance in 5xFAD mice, is associated with impaired synaptic density as well as behavioral deficits, at stages where Aβ pathology is well-established in 5xFAD animals.

### Differential effects of aging and amyloidosis on brain mitochondria

We identified 13 modules enriched in mitochondrial proteins (**Fig. 4A**). Few mitochondria-enriched modules belonged to Clusters 3 and 4 (which also showed neuronal-marker enrichment, e.g., M35, M14), while the remaining mitochondrial modules mapped to Cluster 5, which showed predominantly glial marker enrichment (**Fig. 1E**). M35 in Cluster 3 showed strong enrichment for proteins involved in oxidative phosphorylation (Oxphos), respiratory chain complex, and inner mitochondrial membrane (**Fig. 4B**). M14, also in Cluster 3, was highly enriched in mitochondrial translational machinery (e.g., Mrpl, Mrps), reflective of protein translation in the mitochondria (**Fig. 4B**). In contrast, mitochondrial modules in Cluster 5 (e.g., M27, M31, M28) showed enrichment in proteins involved in metabolism located in either the mitochondrial matrix or inner mitochondrial membrane (**Fig. 4B**).

**Figure 4.**
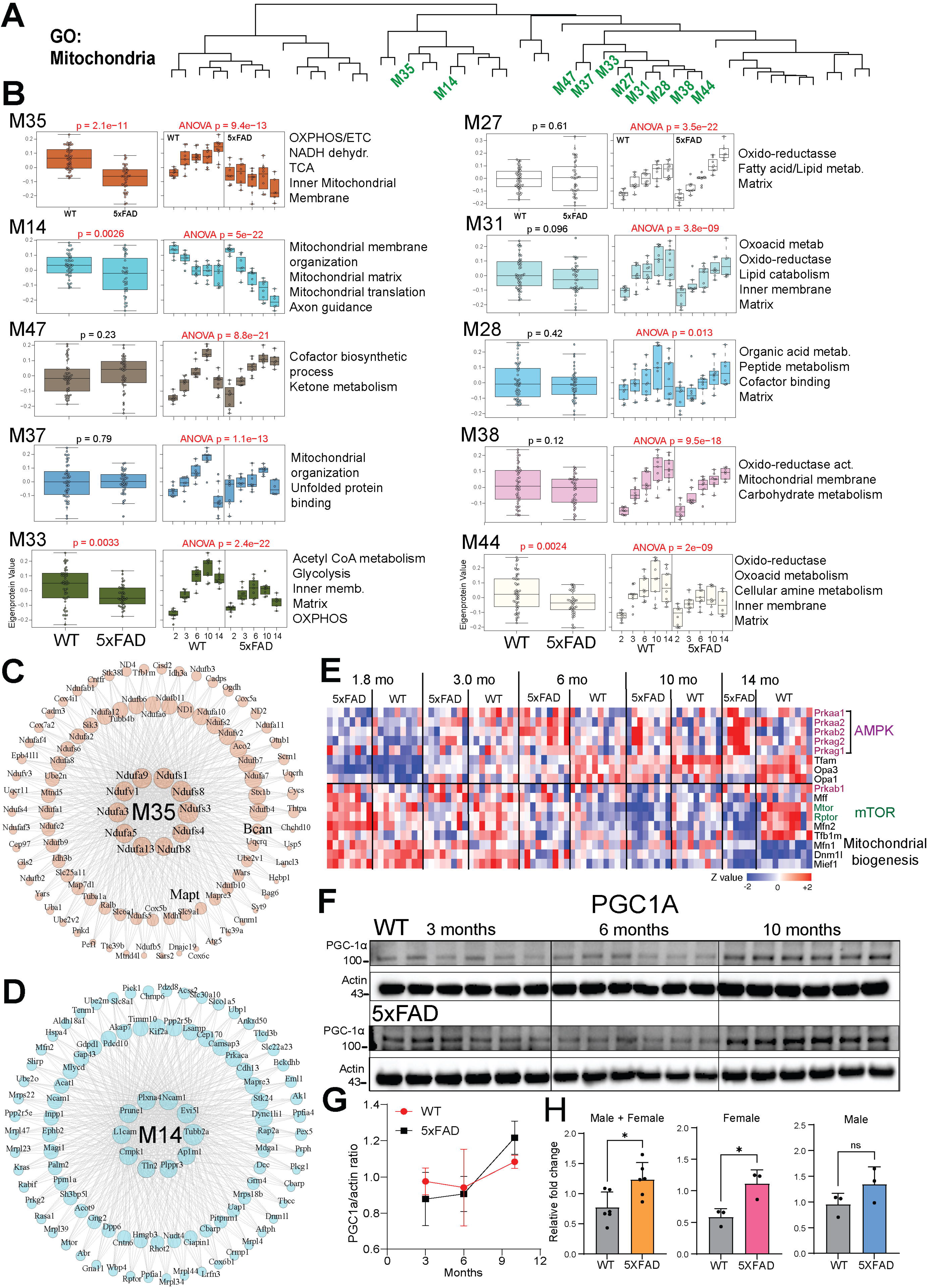
Mitochondrial modules decline with age, and the complex I module M35 is selectively reduced in 5xFAD mice. (A) Module dendrogram with the ten mitochondria-enriched modules highlighted in green. (B) Eigenprotein of each mitochondrial module by genotype with ages pooled (left) and by age within WT and 5xFAD (right, ANOVA P). Representative GO terms at right. (C) Co-expression network of M35, the oxidative phosphorylation / complex I module. Node size scales with intramodular connectivity (kME); the most connected hub proteins are labelled and are dominated by NADH dehydrogenase (Nduf) subunits. (D) As in (C), for M14. (E) Z-scored abundance of AMPK subunits, mTOR pathway components and mitochondrial biogenesis and dynamics proteins in individual mice, grouped by age and genotype. (F) Representative immunoblots of PGC-1alpha and beta-actin in WT and 5xFAD brain at 3, 6 and 10 months. (G) PGC-1alpha normalized to actin across age in WT (red) and 5xFAD (black) mice; mean +/- s.e.m. (H) PGC-1alpha levels in 5xFAD relative to WT for all mice and for females and males separately. *P < 0.05; ns, not significant.

In WT brains, aging generally increased the levels of all mitochondrial modules except for M14, which showed an age-dependent decrease (**Fig. 4B**; ANOVA p = 5e-22) which was accelerated by amyloidosis. These aging patterns were only impacted by 5xFAD genotype (amyloidosis) in M35 (p = 2.1e-11), M14(p = 0.0026), M33 (p = 0.0033), and M44 (p = 0.0024).

All other mitochondrial modules (related to metabolism of fatty acids, lipids, peptides, carbohydrates, ketones) seemed not to be impacted by amyloidosis. M35 had the strongest effect of amyloidosis and contained hub proteins that are key members of the respiratory chain complex, such as Ndufa, Ndufas, Ndufb proteins (**Fig. 4C**). Atg5, a critical regulator of autophagy, was also a member of M35 suggesting that amyloidosis may negatively impact the autophagic process. Interestingly, Mapt (encoding mouse Tau protein) and peri-neuronal net protein (Bcan) were also members of M35. M14 (mitochondrial translation) was enriched in axon guidance proteins such as Plxna4, Ncam1, L1cam (**Fig. 4D**)^24,25^. Other amyloidosis-dependent mitochondrial modules (M44, M33) showed no cell type-enrichment patterns suggestive of general effects of amyloidosis on mitochondrial across all brain cell types. We interrogated module protein membership with the MitoCarta3.0 database across three annotations: broad functional group, subcellular localization, and detailed functional group (**Supplementary Fig. 7**). M32 showed strong enrichment for OXPHOS components and inner mitochondrial membrane localization, while M27, M31, M44, and m43 enriched across metabolism (amino acid, fatty acid, and lipid) and mitochondrial matrix (**Supplementary Fig. 7**). These findings provide further granularity into the functional identity of the mitochondria enriched modules, suggesting that mitochondrial dysfunction is organized into separate domains rather than a global response.

To understand whether aging and amyloidosis effects on mitochondrial modules are due to altered regulation of mitochondrial biogenesis, mitophagy or mitochondrial fusion/fission, we assessed levels of these mitochondrial regulatory proteins in our dataset (**Fig. 4E**). Positive regulators of mitochondrial biogenesis (e.g., TFAM) as well as positive regulators of mitochondrial maintenance (e.g., Opa1/3) increased with age. Protein subunits of the AMPK complex (e.g., Prkaa1/2, Prkaab1/b2, Prkag1), another major driver of energy sensing and mitochondrial biogenesis, showed age-dependent increases except for one subunit (Prkab1) which showed no significant aging effects. Conversely, negative regulators of mitochondrial biogenesis (e.g., mTOR, Rptor) were slightly decreased with age, particularly after 6 months of age (**Fi. 4E**). Mitochondrial fusion (Mfn1/2) as well as mitochondrial fission (Dnm1l, Mief1) proteins were also decreased with age, suggesting that aging in mouse brain may not impact fusion/fission balance (**Fig. 4E**). Since PGC1a (a critical regulator of mitochondrial biogenesis) was not detected by TMT-MS, we performed western blot analysis of brain lysates from 3, 6, and 10 month old WT and 5xFAD mice and observed higher levels of PGC1A protein in older versus younger mice, as well as higher levels in 5xFAD versus WT brain (**Fig. 4F**). Quantification of PGC-1α normalized to actin confirmed a genotype-dependent increase, with 5xFAD mice displaying elevated PGC-1α/actin ratios relative to WT, peaking around 6 months before partially declining at later ages (**Fig. 4G**). 5xFAD mice showed significant increase in PGC1A relative fold change compared to WT; however, this effect was driven primarily by females (**Fig. 4H**, p < 0.05) and not by males (**Fig. 4H**, not significant).

Taken together, we infer that brain aging in mice promotes overall mitochondrial maintenance and biogenesis without impacting fission/fusion balance, while amyloidosis has a selective inhibitory effect of inner mitochondrial membrane machinery involved in oxidative phosphorylation and matrix proteins involved in mitochondrial translation; and these effects of amyloidosis may be preferential to mitochondria in neurons.

### Unique Change in the proteins representing glial vascular and immune cells of the mouse brain

Modules within Clusters 5 and 6 (**Fig. 5A**) showed enrichment of microglial (**Fig. 5B**; M40, M12, M20, M51, M46), astrocyte (**Fig. 5C**; M27, M31, M28, M36), both astrocyte and microglial (**Fig. 5D**; M18, M17), oligodendrocyte (**Fig. 5E**; M9, M41) and endothelial (**Fig. 5F**; M42) cell type-specific markers. Microglial modules showed the strongest amyloidosis-dependent increase in levels compared to any cell type examined (**Fig. 5B**). Several disease-associated microglia (DAM) proteins (e.g., Trem2, Apoe, Lpl), complement-related proteins (C1qa, C1qb, C1qc), lysosomal proteins (Ctsb, Ctsd, Lamp2) and interferon-related proteins (Ifit3b, Stat1) were increased with pathological progression. M40, enriched for defense response, fatty acid metabolism, Ca2+ binding, and kinase activity, showed significant effects of both age and amyloidosis (**Fig. 5B**; p = 00028 genotype, p = 4.5e-08 age). M12 (myeloid differentiation, cell-cell adhesion, late endosome, heparin binding) similarly showed significant age and amyloidosis effects, consistent with sustained microglial activation (**Fig. 5B**; p = 2.6e-13 genotype, p = 5e-23 age). M20 (vacuole, late endosome, cholesterol transport) and M51 (response to stimulus and ECM-related genes including Trem2, Mdk, and Apoe) displayed amyloidosis-driven increases without a significant independent age effect (**Fig. 5B**; p = 9.1e08 M20, p = 3.4e-08, M51), suggesting an early, sustained pathological shift in microglial state rather than a progressive trajectory. M46, enriched for complement pathway, lysosome, and cell surface functions, displayed significant effects of both age and genotype, with the most pronounced effect observed at late disease stages of amyloidosis (**Fig. 5B**; p = 3.2e-07).

**Figure 5.**
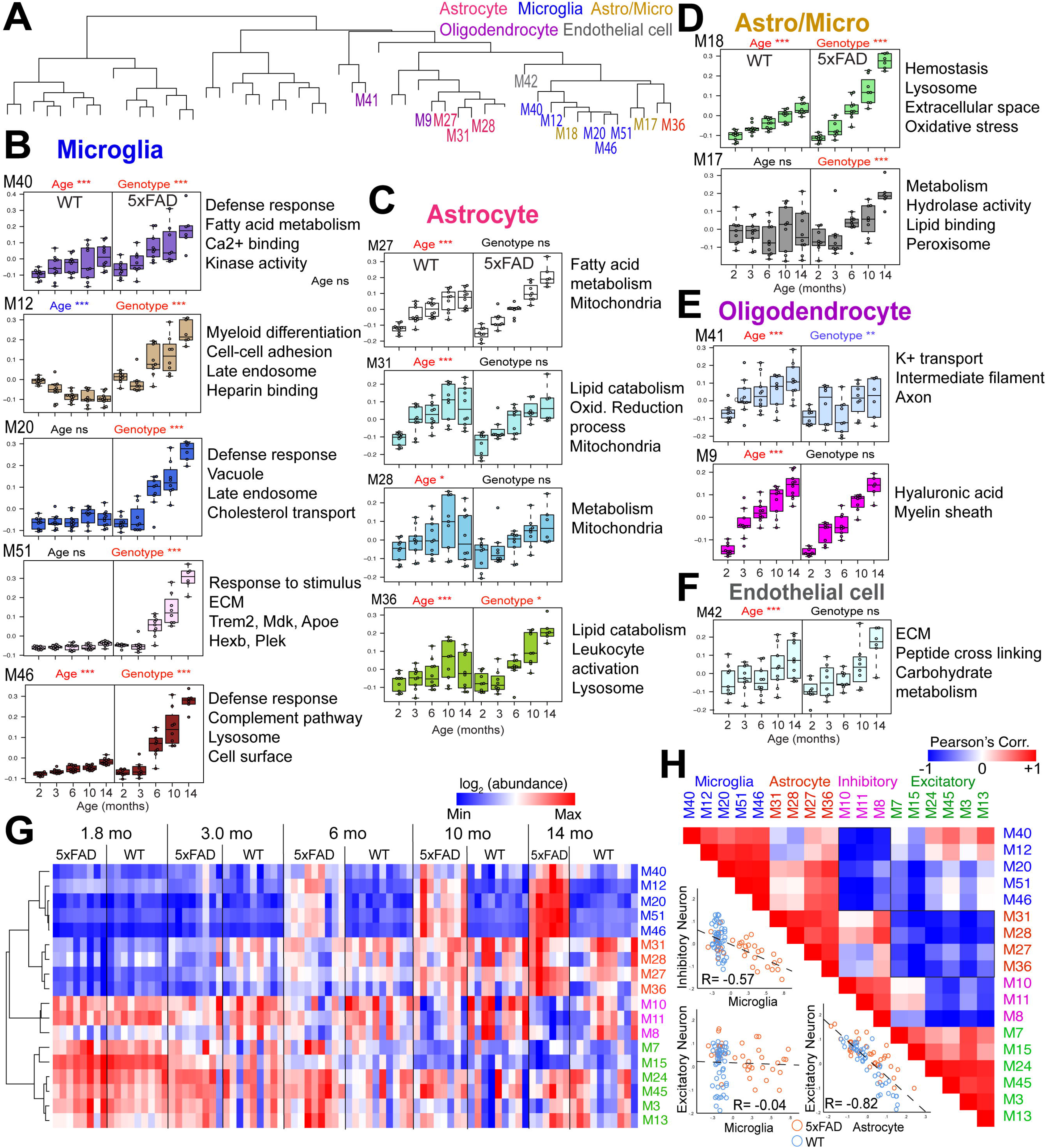
Glial modules are induced by amyloid pathology and are inversely coupled to neuronal modules. (A) Module dendrogram with modules enriched for astrocyte, microglia, mixed astrocyte/microglia, oligodendrocyte and endothelial markers labelled by cell class. (B-F) Module eigenprotein across age in WT and 5xFAD mice for microglial (B), astrocyte (C), astrocyte/microglia (D), oligodendrocyte (E) and endothelial (F) modules. Significance of the age and genotype terms is indicated above each module (ns, not significant; *P < 0.05; **P < 0.01; ***P < 0.001); representative GO terms at right. (G) Min-max scaled log2 abundance of the 18 glial and neuronal module eigenproteins in individual mice, ordered by age and genotype; rows are hierarchically clustered by similarity. (H) Pairwise Pearson correlation among the same 18 module eigenproteins. Insets show the underlying relationships for inhibitory neuron versus microglial (R = −0.57), excitatory neuron versus microglial (R = −0.04) and excitatory neuron versus astrocyte (R = −0.82) modules; WT blue, 5xFAD orange.

In contrast, astrocyte-enriched modules showed robust age-dependent increase and amyloidosis-independent effect in both WT and 5xFAD mice, with the exception of M36 (**Fig. 5C**). Modules M27, M31, and M28, all enriched for fatty acid metabolism, lipid catabolism, oxidative reduction, and mitochondrial processes, increased with age in both WT and 5xFAD animals without reaching significance for genotype (**Fig. 5C**; p = 1e-24 M27, p = 3.8e-08, M31, p = 013 M28). Module M36, enriched for lipid catabolism, leukocyte activation, and lysosomal biology, was the exception, displaying a significant genotype effect (p = 0.03) in addition to a strong age effect (p = 6.1e-10), suggesting a disease-specific astrocyte reactive component within this module.

Two modules displayed mixed astrocyte/microglia enrichment and showed divergent amyloidosis responses (**Fig. 5D**). M18, associated with hemostasis, lysosome, extracellular space, and oxidative stress, was significantly elevated in 5xFAD mice with strong effects of both age and amyloidosis (**Fig. 5D**; p = 002 genotype, p = 1.3e-31 age). M17, enriched for metabolism, hydrolase activity, lipid binding, and peroxisomal biology, showed a significant amyloidosis effect without a significant age interaction (**Fig. 5D**; p = 0.00014), indicating a stable disease-associated shift in this shared glial module. Furthermore, several reactive astrocyte markers, particularly pro-inflammatory A1 markers including Gfap, Gbp2, Vim, and Psmb8, were present in microglia-enriched modules (M46, M20, M18).

Oligodendrocyte modules displayed significant age-dependent increases in both WT and 5xFAD mice with variable amyloidosis effects (**Fig. 5E**). Beyond microglial and astrocyte modules, other glial and vascular modules were also identified. Oligodendrocyte-enriched modules M9 and M41 contained canonical myelin proteins such as Mbp, Plp1, Mag, and Mog and showed an age-dependent increase in WT mice (**Fig. 5E**). M41, enriched intermediate filaments such as Nefm, Nefl, Nefh, axonal and K+ transport proteins, showed a modest reduction with amyloidosis (**Fig. 5E**, p = 0.0013 genotype). M41 also contained some GABA-ergic neuronal proteins such as Pvalb (parvalbumin), suggesting that this module may capture effects of aging and amyloidosis on PV+ interneurons and their unique myelination patterns^26^. Conversely, amyloidosis had no effect on module M9 levels, which was enriched for myelin sheath (Plp1, Mbp) and hyaluronic acid (**Fig. 5E**, p = 1e-24). Module M42 was the only endothelial-enriched module which contained proteins involved in extracellular matrix and carbohydrate metabolism as well as canonical endothelial markers (e.g., Tpm1, Tpm2, Vwf) and vascular smooth muscle proteins (e.g., Myh11). M42 levels increased with age but no effect of amyloidosis was observed (**Fig. 5F**; p = 1.3e-05).

Together, these findings reveal that age and amyloidosis engage glial proteomic changes in a highly cell-type-specific manner. Microglial modules were the most strongly and consistently affected by amyloidosis while astrocyte modules were driven predominantly by aging. Mixed astrocyte/microglia modules suggest partial mechanistic convergence between the two cell types under amyloid pathology. Amyloidosis selectively disrupts axonal and cytoskeletal integrity within oligodendrocyte-associated modules while leaving the core myelin proteome relatively intact. Lastly, the limited representation of vascular modules reflects the inherent under-sampling of the cerebrovascular compartment.

### Insights into glia-neuronal interactions in aging and amyloidosis

To further leverage the cell type resolution of our proteomic network analysis, we examined the data for cross-cell type correlation patterns between microglial, astrocyte, and excitatory and inhibitory neuronal modules (**Fig. 5G-H**). Microglial modules increased with both age and amyloidosis (**Fig. 5G**; M40, M12, M20, M51, M46), astrocyte modules increased predominantly with age (**Fig. 5G**; M31, M28, M27, 36), excitatory neuronal modules declined with age largely independently of amyloidosis (**Fig. 5G**; M7, M15, M25, M45, M3, M13), and inhibitory neuronal modules showed amyloidosis-induced reductions specifically in older animals (6–14 months; **Fig. 5G**; M10, M11, M8). Pearson correlation analysis across all module pairs revealed selectivity in glial-neuron relationships (**Fig. 5H**). Microglial modules were inversely correlated with inhibitory neuronal modules (average R= −0.57) but not with excitatory neuronal modules (average R= −0.04) (**Fig. 5H**). In contrast, astrocyte modules had strong inverse correlations with excitatory neuronal modules (average R= −0.82) but not with inhibitory neuronal modules (average R= −0.2). We also observed that the relationships between astrocyte, microglia and both neuronal modules varied by genotype (5xFAD or WT) (**Supplementary Table 11-12**). Microglial modules had a modest inverse correlation with inhibitory neuronal modules in WT mice (R= −0.33) but had strong negative correlation with inhibitory neuronal modules in 5xFAD mice (R= −0.67). Microglia modules had no association with excitatory modules in WT mice (R= 0.02) but were negatively associated with excitatory modules in 5xFAD mice (R= −0.41). Astrocyte modules had strong inverse relationships with excitatory modules in both WT (R= −0.91) and 5xFAD (R= −0.8) mice.

While correlational in nature, these patterns support a model in which distinct cellular mechanisms for glial cells in age-related and amyloidosis-induced pathology, where microglia may negatively impact inhibitory neurons specifically under amyloidosis conditions, while astrocytes exert detrimental effects on excitatory neurons with aging independent of amyloidosis.

### Conserved pathological mechanisms across mouse models of amyloidosis and human Alzheimer’s disease

To contextualize the proteomic alterations observed in 5xFAD mice with human disease, we integrated our mouse brain proteomic dataset with a post-mortem human brain proteomics dataset comprising cognitively normal controls, asymptomatic AD pathology cases (AsymAD) and AD with dementia cases (AD)^21^. Our goal was to identify mechanisms shared between the preclinical amyloidosis modeled by 5xFAD mice and human AD pathology. We assessed the conservation of human-derived proteomic network within in the mouse proteomic network and identified proteins that are differentially enriched in human AD or in AsymAD relative to controls and show concordant changes in 5xFAD mice compared to and WT mice.

The human proteomic network was derived from over 10,000 proteins and contained 44 modules)^21^. Module preservation analysis using established Zsummary thresholds^27^ identified these six highly preserved (Zsummary >10: M13, M18, M28, M20, M3, M1) and thirteen moderately preserved (10> Zsummary >5: M5, M21, M2, M44, M26, M27, M6, M11, M22, M30, M38, M8, M39) human modules within the mouse proteomic network (**Fig. 6A**, **Fig. 6B**). To further evaluate cross-species conservation, we compared module eigengene expression across AD, AsymAD, and control human brain samples for all significantly preserved modules **(Fig. 6C)**.

**Figure 6.**
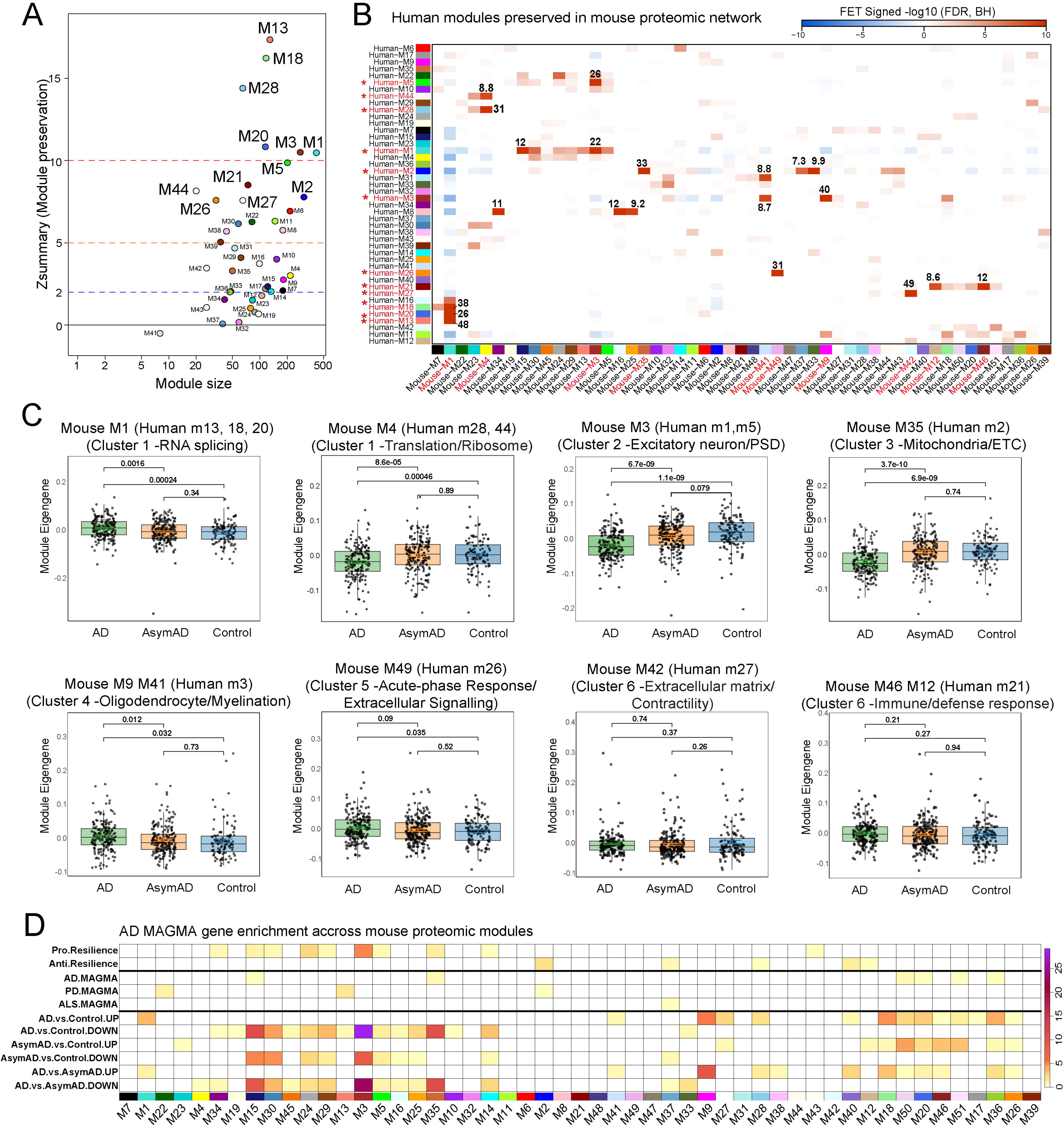
Mouse brain modules are preserved in the human brain proteome and are enriched for AD risk, resilience and disease-associated signatures. (A) Zsummary preservation statistic plotted against module size. Dashed lines mark Zsummary = 2, 5 and 10, the conventional thresholds for weak, moderate and strong preservation respectively; the best-preserved modules are labelled. (B) Overlap between human and mouse modules (one-sided Fisher exact test). Color denotes the signed -log_10_ BH-adjusted P values are printed for the strongest overlaps. (C) Human module eigengene by diagnosis (AD, asymptomatic AD and control) for eight mouse-human module pairs spanning the six clusters. Post-hoc pairwise P values are shown above each panel. (D) Enrichment across the 51 mouse modules of AD, PD and ALS GWAS risk genes (MAGMA), cognitive resilience-associated genes, and up- and down-regulated proteins from the human AD, asymptomatic AD and AD-versus-asymptomatic AD contrasts. Color, -log_10_ BH-FDR.

Human modules M13, M18 and M20 all mapping to mouse module M1, were enriched in RNA splicing factors including Eftud2, Srrm2, Prpf8, showed significantly elevated expression in symptomatic AD relative to controls and AsymAD (**Fig. 6C**), with no difference between the latter two groups. A similar pattern was observed for human module M3, which mapped to mouse modules M9 and M41 and was enriched in oligodendrocyte and myelination proteins (e.g., Lgi3, Mobp, Myo1d, Plp1), as well as some parvalbumin interneuron-associated markers (e.g., Acan, Bin1).

In contrast, several modules showed significantly reduced eigenprotein expression in AD brain samples. Human modules M28 and M44 mapped to mouse module M4, with decreased expression of translational and ribosomal proteins (e.g. Rpl30, Rps7, Farsb) in AD (**Fig. 6C**). Human modules M1 and M5 mapped to mouse module M3, enriched in excitatory neuronal markers (Strn4, Syngap) and glutamatergic synaptic proteins (Homer1, Dlg3, Psd), demonstrated decreased eigengene expression in symptomatic AD, but not AsymAD cases (**Fig. 6C**), suggesting that excitatory neuronal dysfunction could be associated with the more aggressive disease stage. Similarly, the mitochondrial marker-enriched human module M2 mapped to mouse module M35, enriched in several Complex I subunit proteins (e.g., Ndufs1, Ndufb8, Ndufa5), and showed lower eigengene expression in AD but not AsymAD (**Fig. 6C**), implicating suppression of oxidative phosphorylation as a late-stage disease phenomenon.

While the aforementioned cross-species trends identified cellular and metabolic functions that may become altered in the AD with dementia, our analyses also nominated cross-species preserved modules that did not seem to have significant differences in expression across controls and AD groups. Mouse module M42, enriched in cytoskeleton and contractility proteins (e.g., Tpm2, Ehd2, Mylk), mapped to human module M27, and eigengene expression demonstrated no significant changes between Control, AsymAD, or AD. Mouse modules M46 and M12 mapped to human module M21 and demonstrated no difference in eigengene expression among the three groups. These two mouse modules were enriched with select immune and defense proteins, which seemingly may demonstrate no or limited change in expression during the progression of amyloidosis. M46 contained complement proteins (C1q a-c), microglial markers (Msn, Aif1/Iba1), MHC proteins, lysosomal proteins, interferon-related proteins (Ifit3b), reactive astrocyte proteins (Gfap) and AD risk genes (e.g., Tmem106b). M12 contained microglial markers mainly involved in myeloid differentiation (e.g., Ctsa), cell-cell adhesion (e.g., Clip3, Clstn1), and vesicular trafficking (e.g., Arsb, Rab33a). Lastly, human module M26 mapped to mouse module M49 (coagulation, acute-phase response, extracellular region), and contained blood-derived proteins (fibrinogens, apoproteins, heme-related proteins, and serpins). While this module had no change in eigengene expression between an AD and AysymAD comparison, as well as AsymAD and control comparison, it appears that this module had slightly elevated eigengene expression in the symptomatic AD group as compared to controls.

To further evaluate the translational relevance of mouse cortical proteomic modules to human neurodegenerative disease and resilience-associated molecular signatures, we performed enrichment analyses across three categories of human datasets: human cognitive resilience-associated gene sets^28^, disease-risk genes identified using MAGMA (Multi-marker Analysis of GenoMic Annotation) for AD, Parkinson’s Disease (PD), and Amyotrophic Lateral Sclerosis (ALS), and lastly, differential expression signatures comparing data from AD, AsymAD, and control brains (**Fig. 6D**)^21,29^. Resilience-associated signatures showed selective enrichment, with some modules enriched for either pro-resilience (M3, M24, M35, M4, M15, M30, M29, M5, M14, M43) or anti-resilience (significantly enriched in modules M2, M40, M37, M28, M12) gene sets. Among MAGMA-derived neurodegenerative disease risk signatures, AD associated genes demonstrated the most widespread enrichment across mouse proteomic modules (significantly enriched in M15, M35, M50, M20, M51, M36), followed by PD associated genes (significantly enriched in M22, M13, M2), and then ALS-associated genes (enriched in M37). Differential expression-based enrichment analysis comparing AD, AsymAD, and control human cases revealed widespread overlap between human AD-associated changes and mouse cortical proteomic modules. Both upregulated and downregulated gene signatures showed substantial enrichment across multiple modules, indicating that the mouse proteomic networks capture diverse molecular features of human disease progression. The strongest enrichment was observed for genes downregulated in AD relative to controls and AsymAD cases, particularly within M15, cluster 2 modules M3, M30, M29, M24, cluster 3 modules M35, M14, M25. Several of these modules also showed enrichment for genes altered in AsymAD versus control comparisons, although these signals were generally less pronounced than those observed in symptomatic AD. Not surprisingly, most of these mouse modules were also those that were enriched in pro-resilience genes. In contrast, select modules were enriched for genes upregulated in AD, namely M1 (RNA splicing), cluster 5 modules M9, M27 M28, and cluster 6 modules M18, M50, M20, M36, suggesting that functions like immune response, glial and metabolic activity are boosted during amyloidosis progression. Together, these findings demonstrate that specific mouse cortical proteomic modules recapitulate multiple dimensions of human neurodegenerative disease biology, including genetic risk, disease-associated expression changes, and resilience-related molecular programs.

Lastly, we performed a cross-species comparative analysis of aging and amyloidosis-related effects on the brain proteome (**Fig. 7A**). To model aging effects, we estimated an age effect (beta co-efficient) in WT mice. To model an aging effect in human brain, using brain LFQ MS data from 190 non-disease control post-mortem DLPFC samples. We then assessed the overall correlation of aging effects across species using proteins quantified in both datasets. Aging effects across species showed moderate correlation (Spearman R=0.18, p<0.001, **Fig. 7B**) with 57% of proteins shown concordant changes. GO analysis of concordant changes showed that extracellular matrix and small molecular metabolism pathways are increased with aging in both species while gene transcription, protein translation and synapse/axon pathways are decreased with aging in both species (**Fig. 7D**). Proteins that showed highest level of conserved aging effects included ACAN, HAPLN2 and HTRA1. To compare amyloidosis-induced effects across species, we estimated amyloidosis-related effects in mice by estimating an Aβ effect coefficient (5xFAD vs WT) after adjusting for aging effect using data from 86 animals. In humans, we used TMT-MS data from 488 ROSMAP cases in which age effects were removed and estimated AD effect coefficient comparing AD to non-disease controls (excluding cases with mild cognitive impairment). Similar to our findings with aging, we observed a moderate overall correlation of disease effects across species (Spearman R=0.16, 56% concordance) that were independent of aging effects (**Fig. 7C**). GO analysis of concordant changes showed that immune effector, antigen presentation, T cell-related, extracellular vesicle and phagosome pathways (e.g., APP, MDK, APOE, CD44, GFAP, NTN1, OLFML3, SMOC1) are increased with amyloidosis/AD pathology in both species while oxidative phosphorylation and mitochondrial pathways are decreased with disease pathology in both species (**Fig. 7E**). Since human AD has both amyloidosis and tauopathy neuropathological features, we also estimated a pure amyloidosis effect in humans (using the CERAD score of amyloid plaque burden) and found similar patterns of conserved effects across species. This cross-species analysis highlights brain proteomic signatures of aging and of AD pathology that are conserved across humans and mouse models, providing a framework that can help guide pre-clinical modeling of aging and AD using mouse models. The results also reveal the pathways that are not conserved across species and may therefore not faithfully recapitulate human biology.

**Figure 7.**
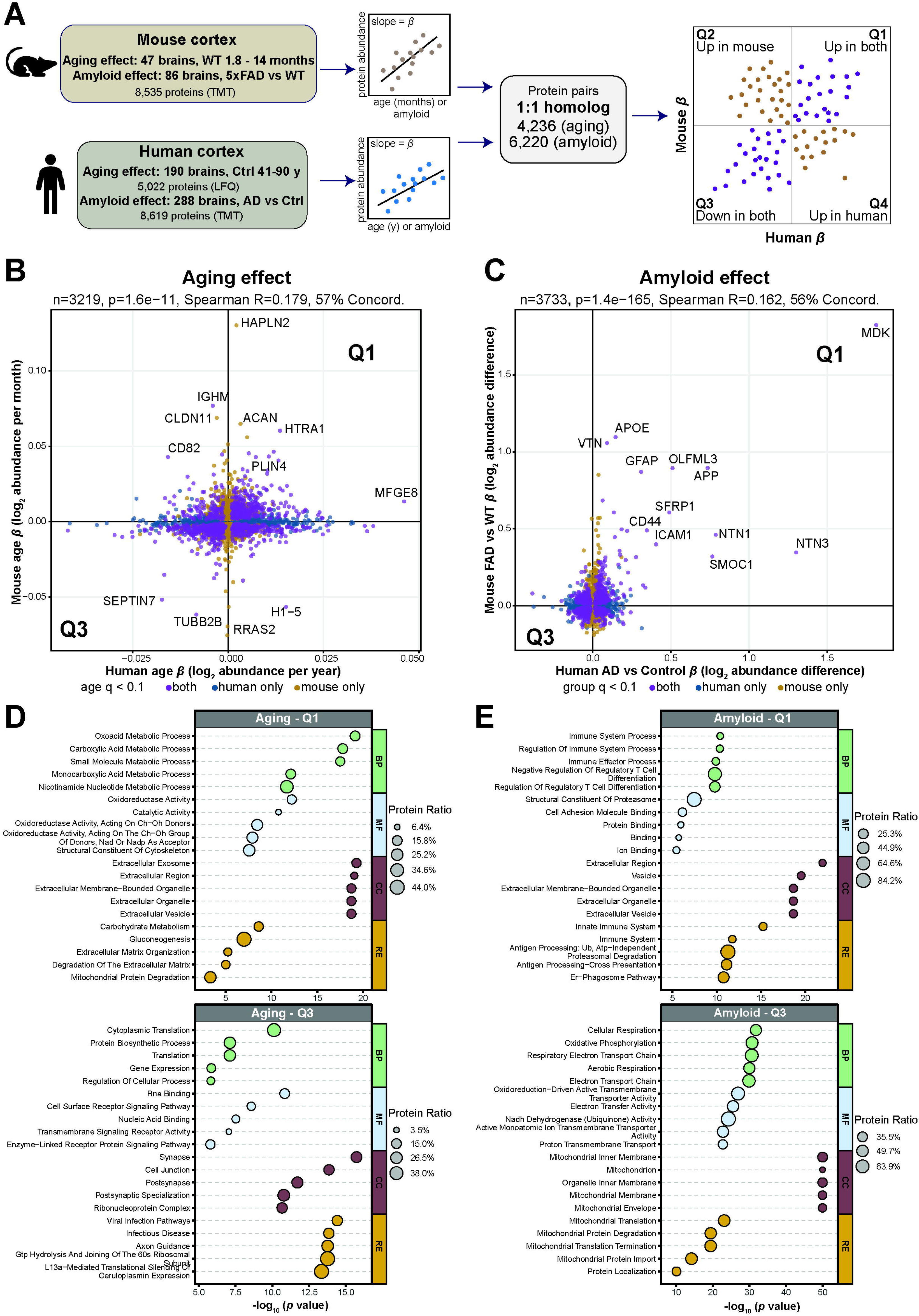
Aging and amyloid effect sizes are weakly but significantly concordant between mouse and human cortex. (A) Analysis design. Per-protein effect sizes (beta) for age and for amyloid pathology were estimated separately in the mouse cortical proteome and in human cortical proteomes, then matched through 1:1 orthologues, yielding 4,236 aging and 6,220 amyloid protein pairs. Quadrants Q1-Q4 define concordant increases, discordant pairs and concordant decreases. (B) Mouse age beta versus human age beta. Points are colored by which species reached significance (q < 0.1); Spearman rho and the number of orthologue pairs plotted are shown. (C) Mouse 5xFAD-versus-WT beta versus human AD-versus-control beta, displayed as in (B). (D) Gene-ontology and pathway enrichment of the concordantly increased (Q1, top) and concordantly decreased (Q3, bottom) proteins in the aging comparison. Dot size, fraction of the term’s proteins recovered; x axis, -log10 P. (E) As in (D), for the amyloid comparison.

## DISCUSSION

Disentangling the molecular effects of aging from those of disease pathology remains a central challenge in understanding neurodegeneration. In the context of AD where aging and pathology are closely intertwined, resolving their respective contributions may be critical to understanding disease-specific mechanisms and informing therapeutic strategies. In this study, we sought to define how aging and amyloidosis independently and jointly shape the brain proteome of 5xFAD mice. Integrating TMT-MS proteomics and transcriptomics with network-based analyses (WGCNA), we resolved the brain proteome into multiple co-expression modules spanning distinct functional and cellular domains, including pathways related to RNA and protein metabolism, neuronal and synaptic systems, mitochondrial function, and glial and immune responses. This allowed us to identify coordinated molecular programs associated with aging, amyloidosis, and their intersection. In contrast to the transcriptome, suppressive effects of aging on RNA splicing and translation, and of amyloidosis on neurons and mitochondria, were more apparent at the proteomic level. In contrast, conserved patterns were also observed, particularly those involving activation of immune and glial responses. Using an integrative proteomics approach across mouse and humans, we also identify molecular pathways that show concordant age-related and amyloidosis-related changes in mice and human brain, providing an important framework that can guide studies of human aging and AD pathology in mouse models.

A prominent observation from our dataset is that age accounts for a substantial proportion of proteomic variance, exceeding that of genotype effects even in the aggressive over-expression 5xFAD model of cerebral amyloidosis^30,31^. This observation is consistent with prior transcriptomic and proteomic studies demonstrating that aging is a major determinant of molecular heterogeneity in the brain^32,33^. Within this framework, we identify age-associated changes in pathways related to RNA processing, protein synthesis, and protein degradation. These findings align with established hallmarks of aging, including impaired proteostasis, altered RNA metabolism, and reduced translational fidelity^10,34–36^. The observed increase in protein–mRNA discordance with age further extends prior reports of modest transcript–protein correlation in the brain and is consistent with the possibility that aging alters the relationship between transcriptional output and protein abundance^35,37^. While the underlying mechanisms cannot be directly inferred from our data, these patterns are compatible with age-related changes in post-transcriptional regulation, ribosomal function, and protein turnover^34,38^. Notably, these effects were largely independent of amyloidosis, supporting the interpretation that they reflect intrinsic features of aging rather than secondary consequences of Aβ pathology. While age-related changes in RNA processing and proteostasis modules did not indicate cell type-specific enrichment patterns, our analyses protein-RNA discordance did indicate differential levels of age-dependent protein-mRNA discordance in different cell types. Future studies using cell type-specific profiling of proteomes and transcriptomes using Simultaneous protein and RNA -omics (SPARO) could address how the proteome and transcriptome alterations are coordinated in a cell type-specific manner^39^.

In contrast with aging effects, cerebral amyloidosis was associated with more selective and context-dependent alterations, particularly within neuronal and synaptic domains. Patterns observed across neuronal modules suggest differential effects of Aβ pathology on excitatory and inhibitory neurons, consistent with heightened vulnerability of inhibitory interneurons, consistent with prior work demonstrating subtype-specific neuronal vulnerability in AD^11,23^. The observed shifts in synaptic protein organization and module relationships, together with functional readouts, align with evidence that early Aβ pathology is associated with alterations in synaptic function and network activity prior to overt neurodegeneration^23,40^. Within this context, our findings support the interpretation that amyloidosis perturbs the balance and coordination of neuronal circuits rather than inducing uniform neuronal decline. Mirroring the documented E/I imbalance in 5xFAD brain, we derived a brain proteomic surrogate of E/I balance using a composite of excitatory and inhibitory neuronal protein modules with a similar pattern if increased E/I imbalance with progressive amyloidosis. Such circuit-level alterations have been proposed to contribute to early cognitive dysfunction and network instability in AD^41–44^.

Mitochondrial pathways exhibited both age- and amyloidosis-associated signatures, but with distinct patterns across modules. Aging was associated with coordinated changes across multiple mitochondrial modules and regulators of mitochondrial function, consistent with prior studies implicating mitochondrial remodeling and altered energy metabolism as features of the aging brain^34,45,46^. In support of increased age-related mitochondrial biogenesis and maintenance, we observed higher levels of PGC1A, AMPK, TFAM proteins with aging^47^. In contrast, amyloidosis was associated with more selective alterations in subsets of mitochondrial proteins, particularly those linked to specific compartments such as the inner mitochondrial membrane, where proteins involved in oxidative phosphorylation/ATP production reside. This pattern suggests that mitochondrial changes in aging may reflect broader adaptations in mitochondrial biogenesis and maintenance, whereas amyloid pathology may preferentially suppress oxidative phosphorylation, indicative of brain energy failure. Prior work has demonstrated that Aβ can interfere with mitochondrial processes including oxidative phosphorylation, protein import, and mitochondrial dynamics and our findings are consistent with these reports at a proteome-wide level^46,48–50^. More broadly, these results support the interpretation that mitochondrial alterations in aging and disease are not uniform, instead they reflect differential vulnerability across distinct mitochondrial processes. Based on our observations, future studies should be focused on assessing mitochondrial dysfunction in a cell type-specific manner, particularly in neuronal subtypes and astrocytes, especially since they comprise the majority of the cell population in the mouse brain.

Glial and vascular modules highlight a striking divergence in how aging- and amyloidosis-associated responses reshape non-neuronal cell biology in the cortex. Amyloidosis elicited robust and coordinated increases in microglial modules enriched for canonical disease-associated microglia (DAM) proteins, complement components, and lysosomal machinery. This is consistent with evidence that Aβ drives a transition towards phagocytic, lipid-processing, and inflammatory microglial states^51,52^. The enrichment of complement and shared astrocyte–microglia modules further suggests that these responses are not restricted to microglia but instead reflect coordinated neuroimmune programs that have been implicated in synaptic remodeling and neurodegeneration^53^. In contrast, astrocyte-enriched modules were predominantly shaped by aging rather than amyloidosis, capturing pathways related to lipid metabolism and mitochondrial function, in line with studies showing that astrocytes undergo substantial metabolic and bioenergetic remodeling during brain aging^54,55^. The age-dependent increases in oligodendrocyte and endothelial modules are similarly consistent with emerging evidence for myelin remodeling and cerebrovascular alterations as key features of the aging brain^56–58^. Notably, the selective reduction of an oligodendrocyte-associated module containing neurofilament and GABAergic markers in 5xFAD mice suggests a vulnerability of myelinated inhibitory circuits, aligning with recent work highlighting specialized interactions between parvalbumin interneurons and oligodendrocytes^59^. Together, these findings support a model in which amyloidosis preferentially amplifies microglial and shared neuroimmune pathways, whereas aging exerts broader effects on astrocytic, myelin-related and vascular programs, underscoring the complementary but distinct contributions of these processes to brain aging and AD-related pathology.

The relationships observed between glial and neuronal modules provide additional context for interpreting these findings. Prior studies have demonstrated that glial activation can influence synaptic function, neuronal excitability, and circuit stability^53,60,61^. The differential associations we observed between glial modules and neuronal subtypes raise the possibility that distinct glial responses may differentially impact specific neuronal populations. For example, an inverse relationship between microglial and inhibitory neuronal modules was observed, particularly in 5xFAD brains, while no relationship between microglial and excitatory neuronal modules were observed. This is consistent with recent literature where AD pathology-related microglia preferentially degrade or engulf peri-neuronal nets around inhibitory neurons prior to interneuronal loss^62^. Interestingly, astrocyte modules were negatively associated with excitatory (glutamatergic) module abundance in both WT and 5xFAD brain, suggesting disease-independent inverse relationships between these two cell types, that could be further characterized by structural and functional studies. The correlative associations between glial and neuronal systems observed in our bulk proteomic study can guide future hypothesis-based targeting and modulation to assess glial-neuronal interactions in aging and disease.

Integrative analysis of 5xFAD mouse and human AD brain proteomic networks revealed a substantial degree of conservation, supporting the relevance of pre-clinical amyloidosis modules for captures key aspects of human disease biology. Notably, a subset of human brain proteomic modules showed preservation in the mouse brain proteomic network, indicating that core mechanisms, such as RNA splicing, are conserved across species. Importantly, modules that were preserved across both proteomes exhibited coordinated alterations across control, asymptomatic AD, and AD cases, highlighting their potential relevance to disease progression. The mitochondrial module M35, conserved across species and containing oxidative phosphorylation, was notably lower in 5xFAD brain as well as in AD human brain, representing a conserved AD-vulnerable mitochondrial function. Whether correction of this mitochondrial defect at early stages of AD pathology impacts disease progression, neuroinflammation or synaptic dysfunction, needs to be clarified. Using a complimentary approach to model the effects of aging and amyloidosis across species by leveraging existing human brain proteomics data from non-disease cases across the aging spectrum, and from AD cases, we also derived proteins and molecular pathways that show conserved aging effects, or conserved AD-related effects in human AD and in an amyloidosis mouse model of AD pathology (concordant: lysosomal/endolysosomal, axon growth, astrocytic glutamate transfer; aging only: myelination, ribosome, synaptic/presynaptic; amyloid only: autophagy, immune/microglial activation). These findings suggest that specific proteomic modules may represent robust and translatable molecular pathways underlying AD pathology and provides a framework for prioritizing targets that are both mechanistically informative and warrant further study in models of amyloid pathology.

While these findings provide a systems-level framework for understanding the intersection of aging and amyloidosis, strengths and limitations of this study should be considered. A key strength of this study is the integration of large-scale, age-dependent longitudinal proteomics with network-based analyses across both aging and amyloidosis, enabling the identification of coordinated molecular programs that would be difficult to resolve using single-modality approaches. The inclusion of multiple age points spanning early to later stages further provides a framework to examine how molecular trajectories evolve over time, rather than capturing static endpoints. Such temporal resolution is difficult to achieve in human studies, which largely rely on cross-sectional or end-stage post-mortem tissue sampling and are often confounded by inter-individual variability and co-existing pathologies. At the same time, our analyses are based on bulk cortical tissue, which constrains resolution of cell type–specific and spatially localized processes. Although network-based and enrichment approaches provide insight into cell type associations, they cannot fully capture cellular heterogeneity or state-specific changes. In addition, while the 5xFAD mouse model recapitulates key aspects of Aβ pathology, including early and robust amyloid deposition, and thereby enables the study of temporally defined amyloid-associated molecular changes^31^, it does not encompass other defining features of human AD, such as tau pathology and extensive neurodegeneration^63^, which may limit direct translational interpretation^15,64^.

In summary, our study presents a comprehensive, multi-layered proteomic resource that captures the dynamic interplay between aging and amyloidosis across molecular, cellular, and functional domains of the brain. By distinguishing broad, coordinated changes associated with aging from more selective alterations associated with amyloid pathology, our findings support a model in which aging establishes a molecular context within which disease-associated processes emerge. This work provides a framework for dissecting how age-related molecular remodeling shapes vulnerability to neurodegeneration and highlights the importance of targeting both foundational aging processes and disease-specific mechanisms in the development of therapeutic strategies. Our findings also highlight targets and pathways involve in human aging and AD pathology that can be modeled in mouse aging and amyloidosis models.

## METHODS

### Animal studies

All experiments involving animal procedures were approved by the Emory University Institutional Animal Care and Use Committee (IACUC, PROTO201700821) and were in accordance with the ARRIVE guidelines.

C57BL/6 J (wildtype [WT]) mice (JAX #000664), and 5xFAD (B6.Cg-Tg(APPSwFlLon,PSEN1*M146L*L286V)6799Vas/Mmjax, JAX #034848) mouse lines were used for experiments in this study. Genotyping was performed using primers and polymerase chain reaction (PCR) conditions as listed on vendor website (Jackson Labs). WT and 5xFAD mice (equal males and females, total n=86) were used for the purpose of this study with data collected from ≥ 3 mice per age group and per sex for all experiments. Mice were closely monitored under standard conditions until aged to 1.8 (n=8 5xFAD: 4 males, 4 females, n=10 WT 5 males, 5 females), 3.1 (n=8 5xFAD: 4 males, 4 females, n=10 WT 5 males, 5 females), 6 (n=9 5xFAD: 5 males, 4 females, n=10 WT 5 males, 5 females), 10 (n=8 5xFAD: 4 males, 4 females, n=9 WT 5 males, 4 females), or 14 (n=6 5xFAD: 4 males, 2 females, n=10 WT 5 males, 5 females) months respectively for each cohort. Two additional cohorts were also used in this study: a separate western blot cohort (3, 6, 10 months; n=6/age/genotype, 3 males, 3 females) and an independent behavioral and SV2A PET cohort (9-10 months (n=9 5xFAD: 5 males, 4 females and n=8 WT: 5 males, 3 females.). Mice were housed in the Department of Animal Resources at Emory University under a 12 h light/12 h dark cycle and kept at controlled temperature (72°F) and humidity (40-50%rh) conditions in a pathogen-free environment with ad libitum access to food and water.

### Euthanasia and tissue collection

Mice from each age cohort were fully anesthetized with isoflurane and euthanized by decapitation, followed by transcardial perfusion with ice-cold 1X PBS. The brain was removed from the skull and hemi-dissected along the mid-sagittal line. The left hemisphere was fixed in 4% paraformaldehyde (PFA) for 24 h and then transferred to 30% sucrose after through washes in PBS. The right hemisphere was further dissected into forebrain and hindbrain (containing cerebellum and brainstem). A small portion of the forebrain was snap frozen on dry ice for RNA sequencing and the remaining forebrain tissue was snap frozen on dry ice for proteomic analyses.

### Brain tissue homogenization and protein digestion

Briefly, frozen forebrain tissue was homogenized in 8 M urea buffer (1:5 by the weight of the tissue; 8 M urea, 10 mM Tris, 100 mM NaH₂PO₄, pH 8.5) containing 1x HALT protease and phosphatase inhibitors (Thermo Fisher). Tissue and buffers were homogenized in RINO tubes (Next Advance) containing ∼100 µl stainless steel beads (0.9–2.0 mm blend) and in a Bullet Blender at 4 °C for two 5-min cycles. Lysates were transferred to new Eppendorf LoBind tubes, sonicated in three cycles (5 s at 30% amplitude, 15 s on ice), centrifuged (15,000 x g, 5 min), and supernatants collected in new tubes. Protein concentration was quantified by BCA assay (Pierce). For protein digestion, 100 µg of each sample was reduced with 1 mM dithiothreitol (30 min, RT), alkylated with 5 mM iodoacetamide (30 min, dark), then digested with Lysyl endopeptidase (1:100 w/w, overnight). Samples were diluted 7-fold in 50 mM ammonium bicarbonate, followed by trypsin (1:50 w/w, 16 h). Peptides solutions were acidified to a final concentration of 1% (vol/vol) formic acid (FA) and 0.1% (vol/vol) trifluoroacetic acid (TFA) and desalted on 30-mg HLB columns (Oasis). Columns were first conditioned by rinsing with 1 ml methanol, washing with 1 ml 50% acetonitrile (ACN), and then 2 × 1 ml 0.1% TFA, washed with 2 × 1 ml 0.1% TFA, and finally eluted with 2 × 0.5 ml 50% ACN. Equal peptide aliquots from each sample were pooled as the global internal standard (GIS), which was split and labeled for each TMT batch as described below.

### Isobaric tandem mass tag (TMT) peptide labeling

Prior to labeling, mouse brain samples (wild-type and 5xFAD at different ages) were randomized across batches according to covariates including genotype, age, and sex, to minimize confounding effects. For each batch, peptides derived from individual samples, along with a pooled GIS were labeled using Thermo Fisher Scientific TMT 16-plex (Thermo Fisher Scientific, A44520R) as per kit instructions and desalted by established protocols (REF). Batches 1-4 contained two GIS channels (TMTpro-126 and −134N) while batches 5-6 contained one GIS channel (TMTpro-126). The remaining channels were reserved for randomized individual samples.

### High-pH off-line fractionation

Dried samples (30%) were re-suspended in high pH loading buffer (0.07% vol/vol NH4OH, 0.045% vol/vol FA, 2% vol/vol ACN) and loaded onto a Water’s BEH 1.7 um 2.1mm by 150mm. An Thermo Vanquish was used to carry out the fractionation. Solvent A consisted of 0.0175% (vol/vol) NH4OH, 0.01125% (vol/vol) FA, and 2% (vol/vol) ACN; solvent B consisted of 0.0175% (vol/vol) NH4OH, 0.01125% (vol/vol) FA, and 90% (vol/vol) ACN. The sample elution was performed over a 25 min gradient with a flow rate of 0.6 mL/min. A total of 96 individual equal volume fractions were collected across the gradient and dried to completeness using a SpeedVac.

### Mass spectrometry analysis and data acquisition

All samples (∼1ug for each fraction) were loaded and eluted by Ultimate 3000 RSLCnano (Thermofisher Scientific) with a 150 μm i.d. capillary column with 1.7 μm C18 CSH (Water’s) over a 21 min gradient. Mass spectrometry was performed with a high-field asymmetric waveform ion mobility spectrometry (FAIMS) Pro frontend equipped Orbitrap Eclipse (Thermo) in positive ion mode using data-dependent acquisition with 2 second top speed cycles for each FAIMS compensative voltage (CV). Each cycle consisted of one full MS scan followed by as many MS/MS events that could fit within the given 1.5 second cycle time limit. MS scans were collected at a resolution of 120,000 (410-1600 m/z range, 4×10^5 AGC, 50 ms maximum ion injection time, FAIMS CV of −45). Only precursors with charge states between 2+ and 6+ were selected for MS/MS. All higher energy collision-induced dissociation (HCD) MS/MS spectra were acquired at a resolution of 30,000 (0.7 m/z isolation width, 35% collision energy, 125k AGC target, 86 ms maximum ion time, TurboTMT on). Dynamic exclusion was set to exclude previously sequenced peaks for 20 seconds within a 10-ppm isolation window.

### Database searches for protein quantification

TMT reporter-ion data were searched and quantified in Proteome Discoverer (PD; Thermo Fisher Scientific) as a single multiconsensus across all batches. We exported the Proteins.txt table and kept master proteins identified at medium or high protein-level FDR confidence. Raw (unnormalized) reporter abundances were taken for all channels. Each UniProt accession was mapped to a mouse gene symbol, and proteins were indexed as Symbol|UniProtID. 8,535 proteins were kept with less than 50% missing values across 86 samples.

### Batch correction and normalization

Within-batch effects were removed with a modified TAMPOR (Tunable Approach for Median Polish of Ratio) algorithm. Within each TMT batch, every protein’s abundance in each channel was divided by its median across that batch’s global internal standard (GIS) channel(s). The resulting ratios were then centered row-wise across batches: each batch’s GIS-derived ratios were scaled so that their within-batch median matched the median across batches. Sample-wise (column) median centering was deliberately omitted. As TAMPOR was run without two-way median polish, the algorithm converged in 1 iteration.

### Covariate regression

For each protein, we fit a linear model of log2 abundance with age, sex, genotype and batch as covariates. Coefficients were estimated by ordinary nonparametric bootstrap (1,000 resamples; R package boot), and each coefficient’s median across resamples was used. Batch effects were then subtracted using the batch coefficients only. Genotype and age were kept in the model so that variance attributable to them was protected from removal.

### Outlier detection

Outlier samples were assessed by sample-network connectivity. Connectivity came from a biweight midcorrelation (bicor)–based sample adjacency matrix (WGCNA fundamentalNetworkConcepts). Samples whose connectivity was more than ±3 SD from the mean were removed, and this was repeated until none remained. No outliers were identified, leaving 8,535 proteins × 86 samples (5 ages × 2 genotypes; n = 6–10 per group).

### Differential expression analysis

Within each age group, FAD and wild-type mice were compared by one-way ANOVA with Tukey post hoc testing (equivalent to a two-sample t-test for two groups), with BH FDR correction. Proteins that showed statistically significant differences after correction, along with their adjusted P values, are reported in **Supplementary Table 3**. Visualization of differential expression as volcano plots was performed using the ggplot2 package in R (version 3.5.2) or with the matplotlib package in Python (versions 3.3.2 and 3.3.4) running in Python version 3.8.5.

### Variance partitioning

Using the variance Partition R package (v1.28.9), variance partitioning analysis (VPA) was used to determine the relative contributions of age, genotype, sex, and experimental batch to variation in brain protein abundance. A linear mixed-effects model was used to estimate the variance components for each protein, with genotype, sex, and batch modeled as random effects and age included as fixed variables. The fitExtractVarPartModel function was used to estimate the residual unexplained variance and the percentage of variance explained by each factor. Using ggplot2 (v4.0.3), variance contributions across proteins were compiled and displayed as violin plots. The proteins that were most significantly correlated with age, genotype, sex, or batch were identified and annotated to show the largest variance contributions for each of the individual covariates.

### Weighted gene co-expression network analysis (WGCNA)

A signed, weighted protein co-expression network was constructed from the batch-corrected log₂ abundance matrix of 8,535 proteins across 86 samples using the WGCNA package (version [1.69 — confirm]) in R [version]. Pairwise biweight midcorrelations (bicor) between all protein pairs were converted to a signed adjacency matrix using a soft-thresholding power of β = 11, selected as the lowest power achieving a scale-free topology fit of R² > 0.8 across a tested range of β = 4–12 (**Supplemental Data 1**). Modules were detected with WGCNA::blockwiseModules() with all proteins assigned to a single block, using deepSplit = 2, a minimum module size of 10 proteins, a merge cut height of 0.07, the mean topological overlap matrix (TOM) denominator, and a partitioning-around-medoids stage that respected the dendrogram (pamRespectsDendro = TRUE, reassignThresh = 0.05). Module eigenproteins were defined as the first principal component of each module, and module membership (kME) as the correlation between each protein and its module eigenprotein (WGCNA::signedKME).

Module assignments were then refined iteratively: proteins with intramodular kME < 0.30 were moved to the unassigned (grey) set, unassigned proteins with a maximum kME > 0.30 were assigned to the corresponding module, and proteins whose kME to another module exceeded their intramodular kME by ≥ 0.10 were reassigned to the higher-kME module. Eigenproteins and kME values were recalculated after each pass, and the procedure was repeated until no further reassignments occurred, requiring 5 of a permitted 30 iterations. The final network comprised 51 modules (M1–M51) ranging in size from 18 to 789 proteins, with [N] proteins ([N]%) remaining unassigned. Module eigenproteins were tested for differences by genotype and by genotype × age group using omnibus F-tests from linear models, and were correlated (bicor) with sample traits, including age within each genotype.

### Gene ontology and cell type marker enrichment analyses

Functional characterization of both differentially expressed proteins and co-expressed protein modules was performed using Gene Ontology (GO) annotation (**Supplemental Data 3**). Analyses were carried out with GO Elite (version 1.2.5), following previously described approaches, and results were refined and visualized using a custom R script.

Module overlap with mouse brain cell-type marker lists, a union of Sharma et al. and Zhang et al. markers for five cell types (see Supplementary Table 4; endothelial cells, microglia, astrocytes, neurons, and oligodendrocytes) was tested with one-tailed Fisher’s exact tests. The background was all network proteins, and p-values were Benjamini–Hochberg (BH) FDR-corrected.

### Network preservation

The WGCNA::modulePreservation() function was applied to evaluate the preservation of modules across different datasets. This function was also used to test how varying levels of missing data affected stability of the consensus network. Composite preservation statistics (Zsummary scores) were calculated by comparing the consensus network as the reference against each target dataset or missing-value threshold, with 500 random permutations performed for each test. To ensure reproducibility, the random seed was fixed at 1, and the quickCor option was disabled. An additional approach to preservation was carried out using synthetic eigenproteins. Specifically, module members from the consensus network whose intramodular membership scores (kME) ranked within the top 20th percentile were grouped together to construct synthetic modules in each target dataset. For synthetic modules containing at least four proteins, synthetic weighted eigengenes were calculated using the WGCNA::moduleEigengenes() function, representing the variance across all member proteins in the target network. Preservation statistics, along with correlation scatter plots of module eigengenes versus sample traits in each target dataset, were then computed and visualized.

### Effect-size models

Human and mouse brain proteomes were compared at the level of per-protein regression coefficients (beta) rather than ranks or overlap counts, so that the magnitude and direction of each protein’s association could be read on one axis per species. All coefficients were estimated by ordinary least squares, one model per protein, with log2 relative abundance as the response; features complete across all samples were fitted through a single shared design matrix and features with missing values were fitted individually on their own complete cases, so residual degrees of freedom vary across features. Two-sided p values were obtained from the t statistic of the coefficient of interest and converted to q values by the Benjamini-Hochberg procedure within each species’ table, independently of the other species.

For the age contrast, mouse log2 abundance was regressed on age with sex as a covariate (y ∼ age + sex) in the 47 wild-type animals only (8,535 protein features; median n = 47 per protein; 88.5% of features fitted at the full n), so that beta is the change in log2 abundance per month of age in the absence of the 5xFAD transgene. The matching human coefficient is the age term of the control-only analysis of the frontal cortex aging proteome (y ∼ age + sex; n = 190 individuals; 5,022 protein features), expressed per year of age. For the disease-group contrast, mouse log2 abundance was regressed on genotype with age and sex as covariates (y ∼ genotype + age + sex) across all 86 animals (39 5xFAD, 47 wild-type; wild-type as reference; 8,534 protein features, the amyloid feature itself excluded), and the human coefficient is the AD-versus-control difference described in 5.1. Because mouse age is expressed in months and human age in years, the two age axes are not unit-for-unit comparable, and the fitted spline describes the shape of the cross-species relationship rather than an equivalence of slopes.

### Scatter construction and significance coloring

Each paired protein is one point, the human coefficient on the abscissa and the mouse coefficient on the ordinate. Points were colored by where the protein reached an FDR threshold of q < 0.10: in both species, in human only, in mouse only, or in neither. The threshold is applied to each species’ own adjusted p value, so “significant in both” requires q < 0.10 twice and is not a joint test. A penalised cubic regression spline with shrinkage was fitted to every plotted point by REML (mgcv::gam with s(x, bs = “cs”)), the basis dimension set from the number of distinct abscissa values, and the shaded band is the pointwise 95% confidence interval of the fitted mean rather than a prediction interval. Pearson and Spearman correlations, the fraction of sign-concordant proteins, and the spline’s effective degrees of freedom and deviance explained are reported for all shared proteins and, as a sensitivity check, for the subsets significant in either and in both species. The spline is always fitted to all plotted points; on cropped panels the view alone is restricted to the 0.5-99.5% quantiles of each axis.

### Definition of the concordant protein lists

Enrichment was tested on proteins that were sign-concordant across the two species and reached q < 0.10 in human, in mouse, or in both. Requiring significance in only one species rather than both admits proteins whose cross-species agreement is supported by one well-powered measurement and a same-signed but under-powered one and roughly triples the input relative to a both-species requirement. Because concordance confines the selection to quadrants I and III of the scatter, the two quadrants were carried forward as separate lists so that enrichment is never averaged over proteins moving in opposite directions: for the age contrast, 725 proteins up in both species and 1,102 down in both; for the disease-group contrast, 963 up in both and 1,122 down in both.

### Gene-set enrichment of the concordant proteins

The background for every test was that contrast’s complete set of paired human symbols — 4,236 for the age contrast and 6,220 for the disease-group contrast — that is, the proteins actually measurable and testable in both species rather than the whole human proteome, so that the universe matches what the scatter could have selected from. Over-representation was assessed with GOparallel (v1.4), which wraps an adaptation of piano::runGSAhyper extended to return the depletion as well as the enrichment tail of Fisher’s exact test. For each list by gene-set pair the one-sided enrichment p, the one-sided depletion p and a signed Z score were computed, the Z being the normal deviate corresponding to half the smaller of the two tails and signed negative where depletion was the smaller tail. Enrichment p values were adjusted by Benjamini-Hochberg across all gene sets tested within each list, and gene sets with fewer than five members present in the universe were not tested. Because the adjusted table covers the enrichment tail only, depleted gene sets carry a negative Z and no adjusted p value and are read from the signed Z table rather than from the figures.

Gene sets were taken from the Bader Lab (Human_GO_AllPathways_noPFOCR_with_GO_iea release of 1 October 2026). The distributed file was repaired before use by removing blank lines, empty gene fields and memberless sets, collapsing duplicate members within a set, reducing repeated name fragments to the canonical name/type/identifier form and transliterating names to ASCII, leaving 27,739 gene sets over 20,712 unique symbols (GO biological process 13,384; molecular function 6,483; cellular component 2,092; Reactome 1,376 plus 1,451 under Reactome database identifiers). After the five-member floor, 9,408 sets were testable against the age background and 11,124 against the disease-group background. The same lists were tested against five brain cell-type marker panels — astrocyte (1,117 symbols), microglia (1,360), neuron (1,492), oligodendrocyte (989) and endothelial (1,029) — assembled in the same format; the panels are close to mutually exclusive, with only 19 of 5,967 unique symbols appearing in more than one. For the ontology runs, closely related terms were pruned within each ontology type by graph-based clustering on Cohen’s kappa of gene-symbol overlap (kappa < 0.30); no pruning was applied to the cell-type runs. Bubble plots retain terms with Z > 1.96 and at least five member proteins in the list and show the five highest-Z terms per ontology type, with bubble area proportional to the gene ratio.

### Bulk brain RNA-seq sample preparation

Snap-frozen brain tissue was homogenized in 600 μL Buffer RLT (Qiagen), and total RNA was extracted using the RNeasy Mini Kit (Qiagen) with on-column DNase digestion according to the manufacturer’s instructions. RNA concentration was measured using the Qubit 2.0 Fluorometer (Life Technologies), and RNA integrity was evaluated using an Agilent TapeStation 4200. A total of 10 ng of RNA from each sample was used for cDNA synthesis using the SMART-Seq v4 Ultra Low Input RNA Kit (Takara Bio) following the manufacturer’s protocol.Amplified cDNA was fragmented, and dual-indexed sequencing adapters were added using the Nextera XT DNA Library Preparation Kit (Illumina). Library quality and fragment size distribution were assessed on an Agilent TapeStation 4200, after which libraries were pooled at equimolar concentrations. Sequencing was performed on an Illumina NovaSeq 6000 platform using 100-bp single-end (SE100) reads, generating approximately 25 million reads per sample. RNA quality metrics were assessed for all samples before library preparation. No predefined RNA Integrity Number (RIN) cutoff was applied, and all samples were processed for library construction.

### Bulk RNA-seq data processing

Raw sequencing reads (FASTQ files) were subjected to quality control using FastQC, and sequencing quality summaries were generated with MultiQC. Adapter sequences and low-quality bases were removed where necessary. High-quality reads were aligned to the appropriate reference genome using a splice-aware aligner STAR. Gene-level read counts were quantified using featureCounts using the Subread package based on the corresponding gene annotation (GTF). The resulting raw gene count matrix was used for downstream analyses in R.

### Protein-mRNA concordance analyses

Concordance between protein and transcript expression was evaluated at the sample level across wild-type (WT) and 5xFAD cohorts. All analyses were performed in RStudio (version 2024.12.1). Protein quantification values were taken as log₂-transformed normalized intensities, and transcript expression values were log₂-transformed normalized RNA counts. For each sample, vectors of normalized protein abundances and matched mRNA expression values were generated. Only genes with both nonzero protein and RNA measurements were included. Spearman’s rank correlation coefficient (Spearman Rho, ρ) was calculated between protein and transcript expression vectors using the cor.test() function with method = “spearman”, yielding a single concordance score per sample. This approach captured monotonic relationships between modalities and is robust to non-linear effects.

In addition, biweight midcorrelation (bicor), implemented in the WGCNA package, was calculated for comparison. P values from correlation tests were obtained and adjusted for multiple testing using the Benjamini-Hochberg procedure where appropriate.

To investigate cell-type-specific trends, concordance analyses were repeated using marker gene sets corresponding to astrocytes, microglia, oligodendrocytes, neurons, and endothelial cells. Marker lists were constructed by merging previously published gene sets (Sharma et al. and Zhang et al.) and harmonized such that overlapping assignments were uniquely assigned to a single cell type. For each cell type, protein and RNA expression vectors restricted to the respective cell-type-specific gene list were correlated on a per-sample basis, producing cell-type-specific concordance scores. In addition, a global concordance measure for all genes was computed across the full set of measured protein-mRNA pairs. Visualization was performed in R using the ggplot2 package. Scatterplots were used to illustrate the distribution of concordance values across genotypes and ages. For each cell type, protein-mRNA correlation analyses as described above were also conducted on a genewise manner and filtered based on set correlation coefficient values to generate lists of most concordant, most discordant, and noncorrelated protein-mRNA gene pairs across cell types.

### Western blotting

Samples for western blots were prepared using 15μg of brain tissue homogenate boiled in 4× Laemmli buffer supplemented with β-mercaptoethanol (1:10). Samples were resolved on a BOLT 4%–12% Bis–Tris gel along with a molecular weight indicating ladder. Protein was transferred to polyvinylidene difluoride membrane using the iBLOT2 stack system. The membrane was blocked for 1h at room temperature using the StartingBlock blocking buffer. Blots were incubated with primary antibodies (PGC1α Invitrogen, PA5-72948 & β -actin-Cell signalling-4967) overnight at 4°C followed by three washes with TBS-T (0.1%),10 minutes each. Secondary antibody incubation was performed for 1h at room temperature followed again by three 10 minutes washes with TBS-T (0.1%). The membrane was imaged using the Odyssey Li-COR system or Bio-Rad ChemiDoc MP Imaging system. Densitometry analyses for band density were done using ImageJ software and statistical significance was measured using t-test on Prism 10 software.

### SV2A positron emission tomography (PET) imaging in 5xFAD and WT mice

The general procedure for microPET studies was based on previously with minor modification in this work. Dynamic microPET imaging study was performed on a Genisys 8 PET scanner (Sofie Biosciences, Culver, CA). Mice(independent of those used for TMT-MS studies) were anesthetized with 1–2% (v/v) isoflurane throughout the imaging session. The radioligand, [^18^F]SDM-8 (ca. 40 μCi/100 μL) was administered via a pre-installed tail vein catheter. The dynamic scan was acquired for 60 min. The PET dynamic images were reconstructed using G8 software (Analysis Tools and System Setup/Diagnostics Tool, Sofie Biosciences) and analyzed with PMOD 4.3 (PMOD technology, Switzerland, License ID: 1340). Standardized uptake values (SUV) were calculated and used to generate time-activity curves for the brain regions (cortex, thalamus, hippocampus, striatum, and cerebellum). SUV = (radioactivity per mL tissue/injected radioactivity) × body weight.

### Morris water maze behavioral testing

Behavior was run in an independent cohort of 9-10 month-old 5xFAD and WT mice from the same colony: 5xFAD N = 9 mice: 5 males, 4 females and WT N = 8 mice: 5 males, 3 females. Morris Water Maze training took place in a round, water-filled tub (52-inch diameter) in an environment rich with extra maze cues and a small platform 1 cm below the surface. White tempera paint (non-toxic) was added to the water to make it opaque so that the mice could not see the platform. Mice were placed in the water maze with their paws touching the wall from 4 different starting positions (N, S, E, W) in water that starts at 25°C and typically declines to 22°C by the time a whole group of mice is tested. An invisible escape platform was located in the same spatial location 1 cm below the water surface independent of a subject’s start position on a particular trial. In this manner, subjects were able to utilize extra maze cues to determine the platform’s location. Each subject was given 4 trials/day for 5 consecutive days with a 15-min intertrial interval. The maximum trial length was 60 s and if subjects did not reach the platform in the allotted time, they were manually guided to it. Upon reaching the invisible escape platform, subjects were left on it for an additional 5 s to allow for survey of the spatial cues in the environment to guide future navigation to the platform. After each trial, subjects were dried and kept in a dry plastic holding cage filled with paper towels to allow them time to dry off. The holding cage was placed half-on, half-off a heating pad and mice were monitored closely. Following the 5 days of task acquisition, each mouse was subjected to a probe trial (60 s) to assess their spatial memory, in which the platform was removed, and the amount of time spent in the target quadrant which previously contained the escape platform during task acquisition was measured. All trials were videotaped and performance analyzed by means of MazeScan (Clever Sys, Inc.). For each trial, the escape latency (s), the traveled distance (mm), and swimming speed (mm/s) were recorded. Data of four trials for each animal in each daily session were averaged and analyzed with a two-way repeated measures ANOVA, followed by Sidak’s post hoc test. In the probe trial, the time spent at the target quadrant was analyzed using an unpaired-t test (5xFAD vs WT) or a one-way ANOVA followed by Sidak’s post hoc test (when separating data by sex).

### Fear conditioning

Cued and contextual conditioning is a fear conditioning task which measures the ability of a rodent to form and retain an association between aversive experience and environmental cues. On 9-10 month old 5xFAD and WT mice, we used a standard fear conditioning paradigm that occurs over a period of three consecutive days. Mice were placed in the fear conditioning apparatus (7” W, 7” D X 12” H, Coulbourn) and allowed to explore the enclosure for 3 min. Following this habituation period, 3 conditioned stimulus (CS)-unconditioned stimulus (US) pairings were presented with a 1 min intertrial interval. The CS consisted of a 20 second 85 db tone and US consisted of 2 seconds of a 0.5 mA footshock, which co-terminated with each CS presentation. This level of foot shock is relatively modest; by comparison, stress-induced reinstatement of drug seeking uses 15 min of intermittent 0.8 mA footshock. Shock was delivered via a Precision Animal Shocker (Colbourn) connected to each fear-conditioning chamber. The shock level was set and verified prior to each training session. One minute following the last CS-US presentation, animals were returned to their home cage. On day 2, the animals were presented with a context test, during which subjects were placed in the same chamber used during conditioning on Day 1, and the amount of freezing time was recorded via a camera and the software provided by Colbourn. No shocks were given during the context test. On day 3, a tone test was presented, during which time subjects were exposed to the CS in a novel compartment. Initially, animals were allowed to explore the novel context for 2 min. Following this habituation period, the 85 db tone was presented for 6-min, and the amount of freezing time recorded. No shocks were given during the tone test. The amount of freezing time was analyzed with a two-way repeated measures ANOVA, followed by Sidak’s (5xFAD vs WT) or Tukey’s post hoc tests (when separating data by sex).

## DATA AVAILABILITY

### PRIDE

The mass spectrometry proteomics data generated in this study have been deposited in the ProteomeXchange Consortium via the PRIDE partner repository under the dataset identifier PXD073498.

### GEO

GEO submission id for mouse brain bulk RNAseq data is **GSE343781**

Processed proteomic and transcriptomic data are also provided in supplemental datasheets.

## ACKNOWLEDGEMENTS

We acknowledge the following individuals for technical support related to this project: Hailian Xiao, Jason P Schroeder, Asiah McWhorter, Aditya Natu, Ananthramanan Shantaraman.

Next generation sequencing services were provided by the Emory National Biomedical Research Center (NBRC) Genomics Core, which is supported in part by NIH P51 OD011132. Sequencing data were acquired on an Illumina NovaSeq 6000 funded by NIH S10 OD026799.

## CONTRIBUTIONS

Sruti Rayaprolu - Acquisition of funding, study conceptualization and design, data analysis and interpretation, manuscript writing and editing

Sneha Malepati – Analysis of mass spectrometry data, manuscript writing and interpretation

Eric B. Dammer - Analysis of mass spectrometry data, manuscript writing and interpretation

Kyla Langdon - Manuscript writing and interpretation of results

Rashmi Kumari – Experiments (Western blot analysis) and manuscript editing

Amanda D Brandelli - Experiments (Western blot analysis) and manuscript editing

Claudia Espinosa-Garcia - Experiments (Western blot analysis) and manuscript editing

Duc D Duong – Acquisition of mass spectrometry data and first-tier data cleanup

Upasna Srivastava – Data analysis and manuscript writing

Prateek Kumar – Experiments and manuscript editing

Ali Tfaily - Experiments (Western blot analysis) and manuscript writing and editing

Jiahui Chen – imaging design, data analysis

Xin Zhou and Yinlong - radiochemistry and tracer production

Ahmad – data analysis

Steven Liang – imaging experiment design, data analysis

Nicholas T. Seyfried: Acquisition of funding, study conceptualization and design, manuscript editing

Srikant Rangaraju – Acquisition of funding, study conceptualization and design, data analysis and interpretation, manuscript writing and editing

## FUNDING

The work was partly by the following grants:

S. Liang: AG074218, AG078058, and S10OD034326.

S. Rangaraju and N.T.Seyfried: R01AG075820, R01NS114130

S. Rayaprolu: F32AG064862, K01AG103502

## ABBREVIATIONS

Aβ: Amyloid-beta
AD: Alzheimer’s disease
ALS: Amyotrophic lateral sclerosis
DAM: Disease-associated microglia
DEPs: Differentially expressed proteins
GO: Gene ontology
MAGMA: Multi-marker Analysis of GenoMic Annotation
PD: Parkinson’s disease
PET: Positron emission tomography
SPARO: Simultaneous protein and RNA-omics
SV2A: Synaptic vesicle glycoprotein 2A
TMT-MS: Tandem-mass tag mass spectrometry
WGCNA: Weighted gene co-expression network analysis
WT: Wild-type

---

**Supplementary Figure 1. Gene-set variation analysis of the 5xFAD brain proteome across age.** Heatmap of the mean difference in GSVA enrichment score between 5xFAD and WT mice at 2, 3, 6, 10 and 14 months, computed over gene ontology gene sets. Red, higher in 5xFAD; blue, lower. Rows are hierarchically clustered into six groups, numbered and color-coded at left; representative gene sets for each group are listed at right. Immune, complement, antigen-presentation and lysosomal sets (groups 1 and 4) rise progressively from 6 months, while mitochondrial, respiratory-chain and synaptic sets (groups 2 and 6) fall at the oldest ages.

**Supplementary Figure 2. Protein and mRNA abundance are largely uncorrelated at the gene level.** (A) Mean protein abundance versus mean mRNA abundance across the 7,720 genes with paired proteomic and transcriptomic measurements. Biweight midcorrelation rho = −0.054, R2 = 0.003; dashed line, linear fit. (B) Genes ranked by gene-wise protein-mRNA biweight midcorrelation. Genes at the positive and negative extremes of the distribution are labelled. The most concordant genes are dominated by complement, MHC class I and microglial transcripts; the most discordant include myelin and neuronal surface proteins.

**Supplementary Figure 3. Protein-mRNA concordance declines with age in a cell type-specific manner.** Gene-wise protein-mRNA correlation (biweight midcorrelation rho) in each mouse, computed separately over neuronal, astrocytic, oligodendrocytic, microglial and endothelial marker genes, plotted against age. WT blue, 5xFAD red; dashed lines are per-genotype linear fits. Panels expand the cell-type trajectories summarized in Fig. 2F; endothelial and microglial concordance declines most steeply with age.

**Supplementary Figure 4. Synaptic density measured by SV2A PET is unchanged in 5xFAD mice when sexes are combined.** (A) Representative [18F]SDM-8 PET images in female and male WT and 5xFAD mice, shown in three orthogonal planes on a common SUV scale. (B) Regional [18F]SDM-8 standardized uptake value in whole brain, cortex, thalamus, hippocampus, striatum and cerebellum with sexes combined. No region differs by genotype. (C) Regional time-activity curves across the 60-min scan for female and male WT and 5xFAD mice. (D) Regional SUV analyzed separately in females (top) and males (bottom). Significant genotype differences are confined to female mice. ns, not significant; *P < 0.05; **P < 0.01. Data are mean +/- SEM with individual mice shown.

**Supplementary Figure 5. 5xFAD mice show impaired contextual fear memory with preserved spatial learning.** (A) Fear-conditioning paradigm: tone-shock training followed by contextual and cued fear testing. (B) Contextual and (C) cued fear conditioning. Freezing across the test session in WT and 5xFAD mice (left) and split by sex (right). (D) Morris water maze acquisition across five training days: escape latency, path length and swim speed, by genotype (left) and by sex (right). (E) Morris water maze probe trial: time spent in the target quadrant, by genotype. (F) Probe trial performance split by sex. Data are mean +/- SEM with individual mice shown. *P < 0.05.

**Supplementary Figure 6. MitoCarta3.0 annotation enrichment across the 51 protein co-expression modules.** Enrichment of each module for MitoCarta3.0 (A) functional groups, (B) sub-mitochondrial localizations and (C) detailed functional categories. One-sided Fisher exact test; color denotes -log10 BH-adjusted P and the adjusted P value is printed in significant cells. Modules are ordered as in Fig. 1F with the module color bar above each heatmap. M35 is by far the most strongly enriched module (OXPHOS, adjusted P = 9.8e-57; inner mitochondrial membrane, adjusted P = 2.9e-41), supporting its assignment as the complex I / respiratory chain module in Fig. 4.

**Supplemental Data 1. Intramodular protein co-expression networks for all 51 modules of the mouse cortical proteome.** Each page shows the network neighborhood of one module, with modules presented in the order of the module eigenprotein dendrogram. Nodes are the 100 proteins with the highest intramodular membership (kME) for that module; modules containing fewer than 100 proteins are shown in full.

**Supplemental Data 2. Global network architecture, module–trait relationships, and eigenprotein trajectories for all 51 modules.** WGCNA cluster dendrogram of the 8,535 proteins quantified across 86 cortices (WT and 5xFAD) followed by eigenprotein (eigengene) network: dendrogram of the 51 module eigenproteins and the corresponding heatmap of pairwise eigenprotein relatedness. Additional data include module–trait relationship heatmap and panels: (left) module eigenprotein by genotype with all ages pooled, (center left) module eigenprotein by age-by-genotype group (1.8, 3.1, 6, 10.2 and 14.4 months in wild-type and 5xFAD), and (center right, right) module eigenprotein against age within wild-type and within 5xFAD animals.

**Supplemental Data 3. Gene ontology enrichment of all 51 protein co-expression modules.** Modules are shown in numerical order, M1 (turquoise) through M51 (thistle1). For each module the five most strongly enriched terms are plotted for each of the three ontologies: biological process (green), molecular function (blue) and cellular component (maroon).

