## Supplemental Data 1 for "Network analysis of mouse brain proteomes resolves molecular and cellular endophenotypes of aging and Aβ pathology"

**M7 black module**

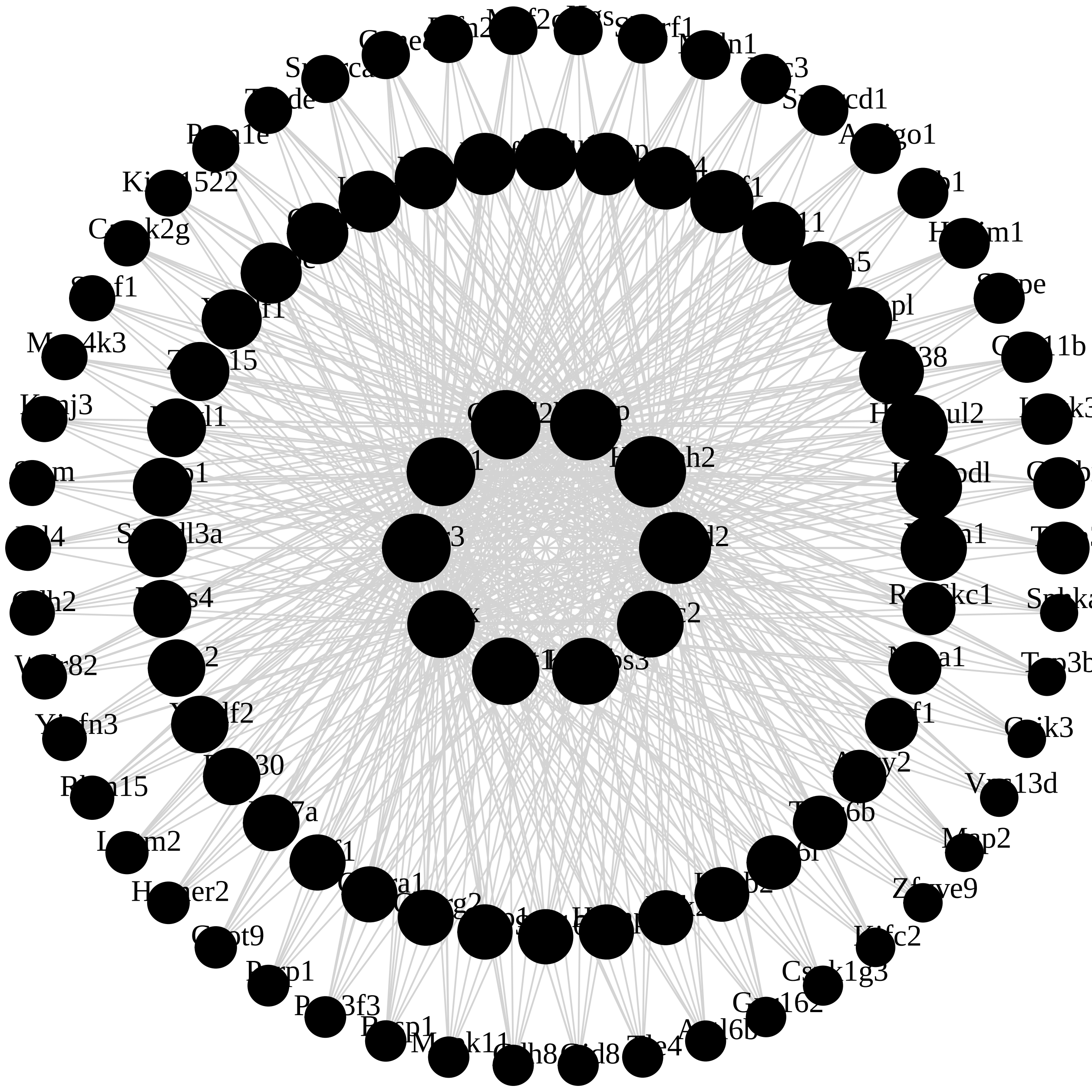

**M1 turquoise module**

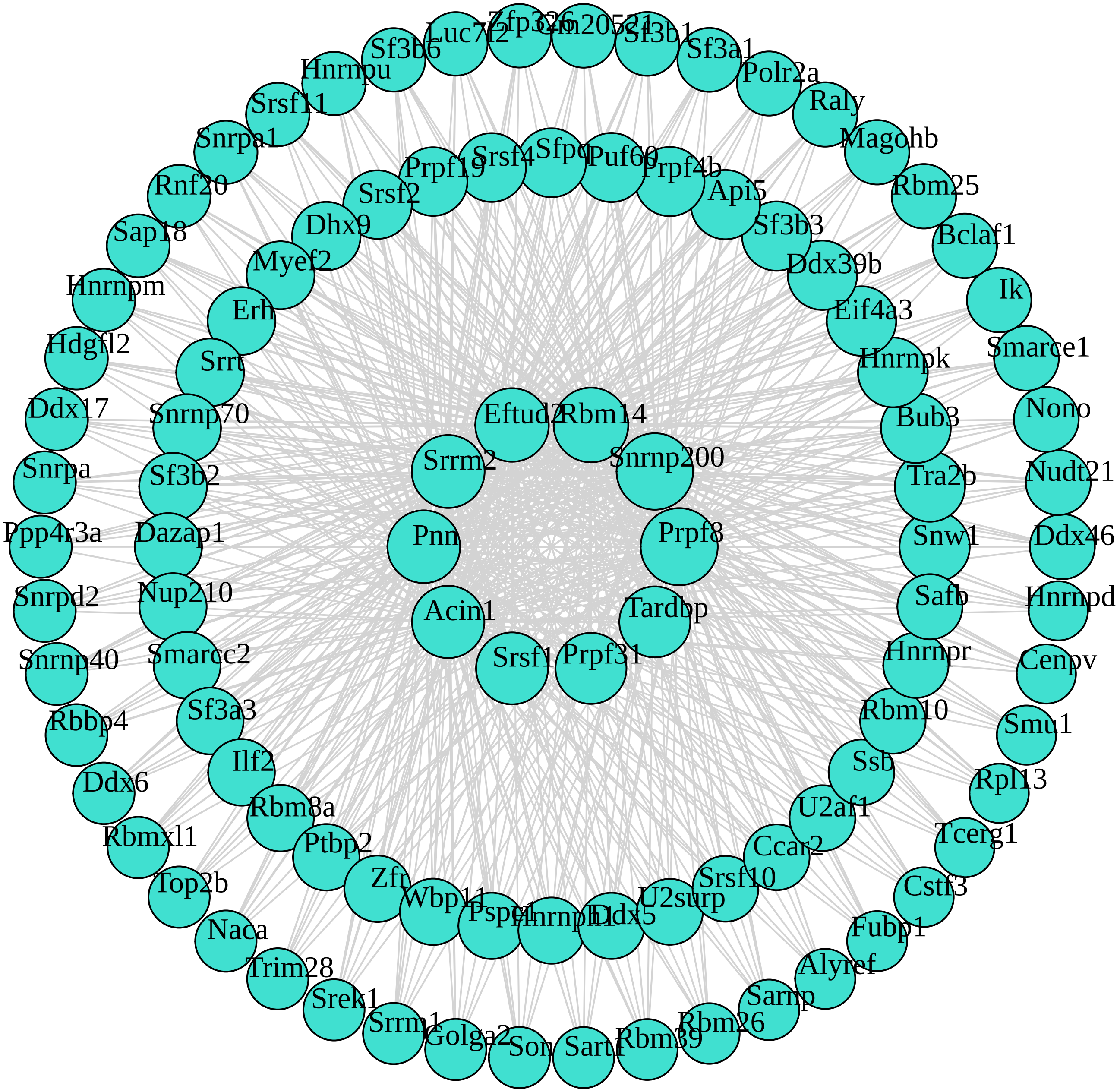

**M22 darkgreen module**

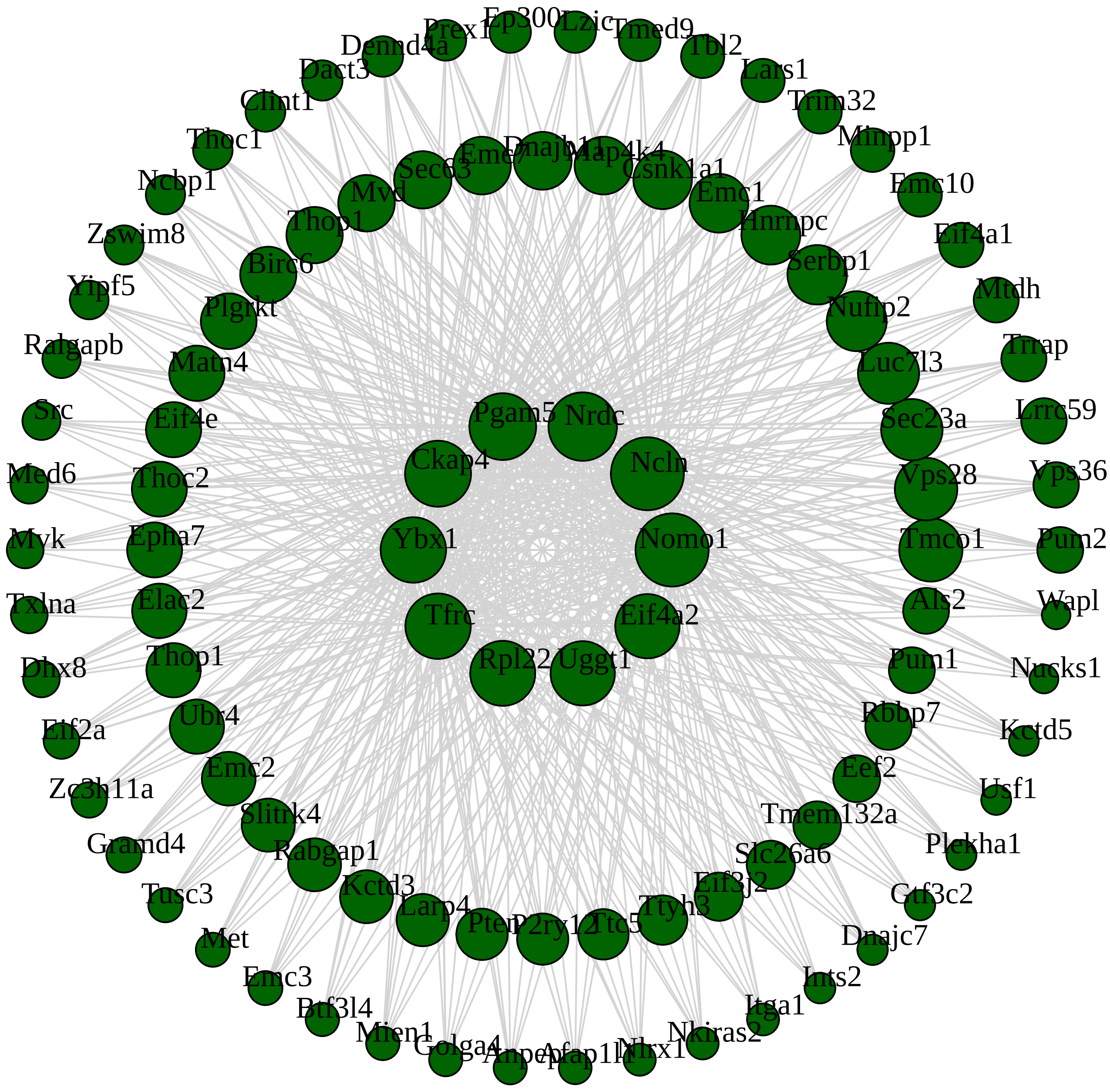

**M23 darkturquoise module**

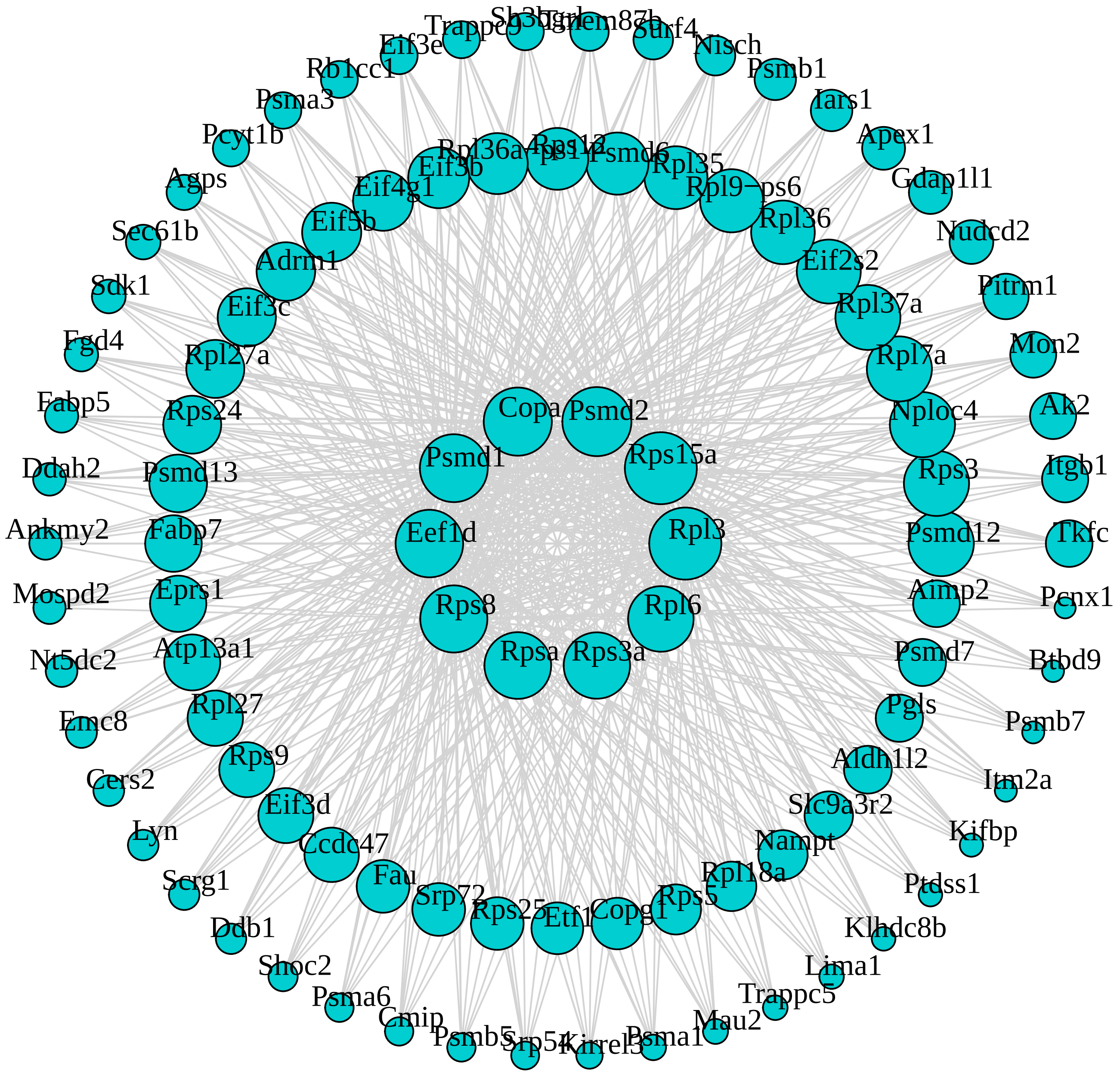

**M4 yellow module**

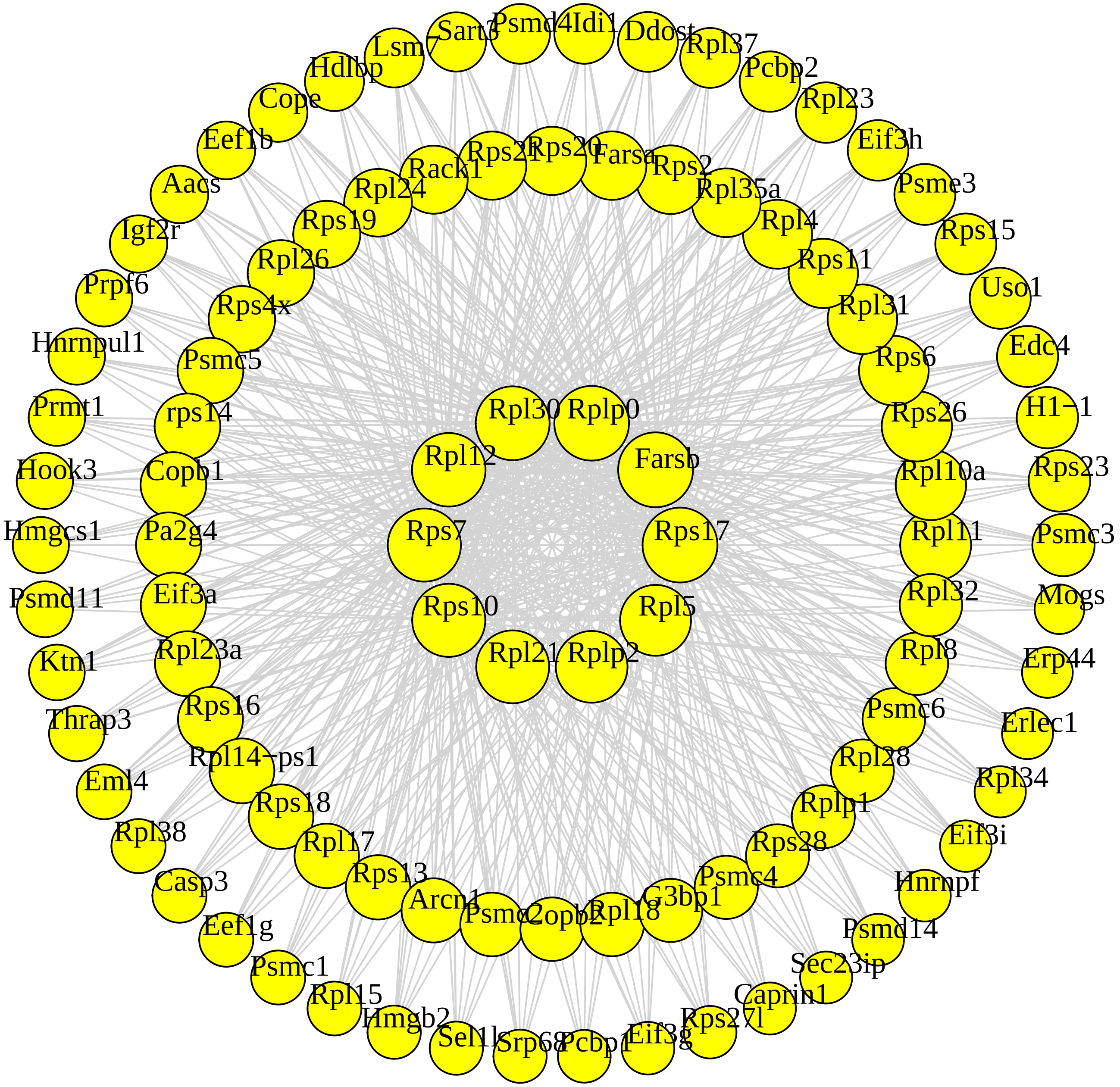

**M34 darkmagenta module**

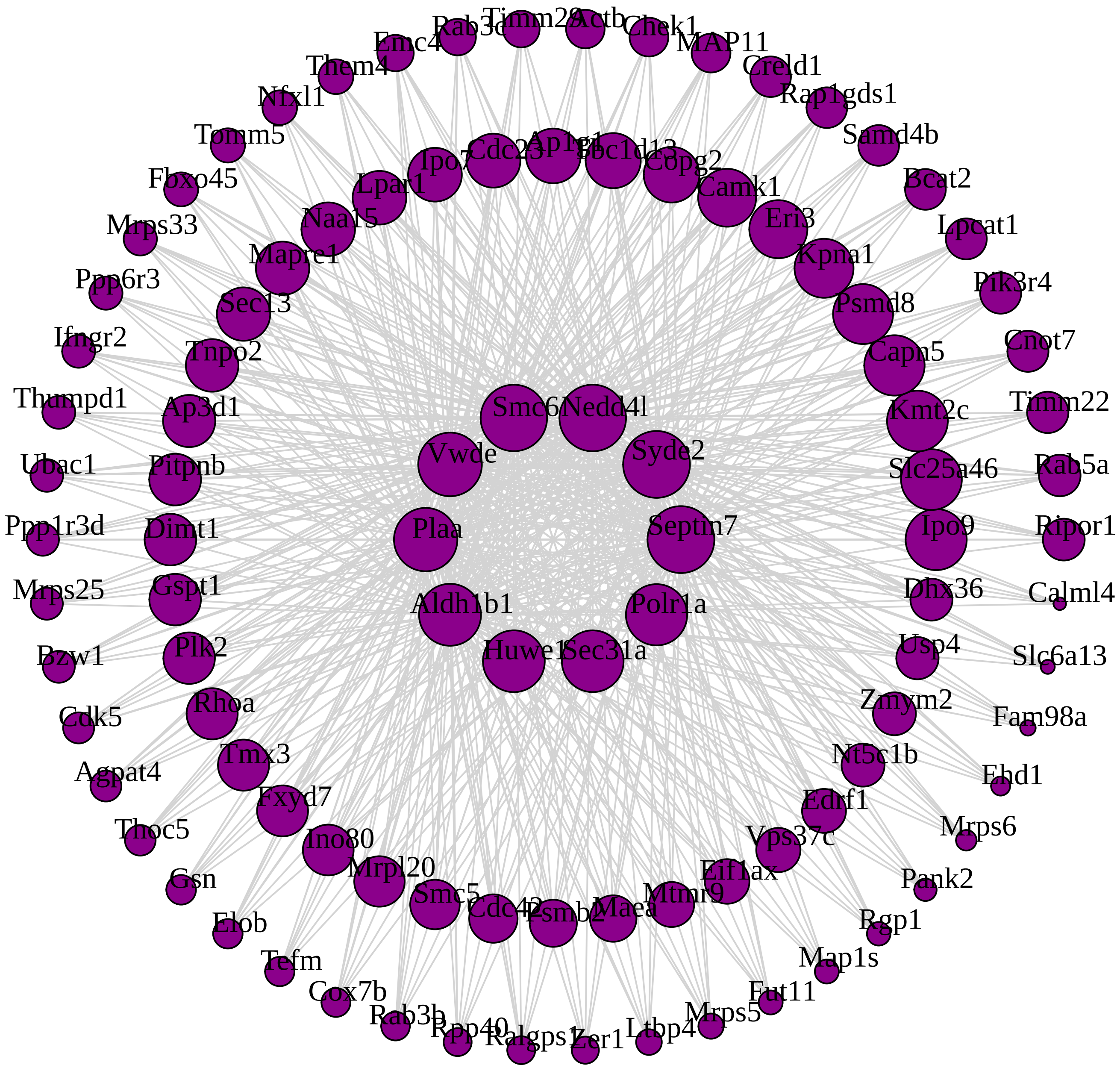

**M19 lightyellow module**

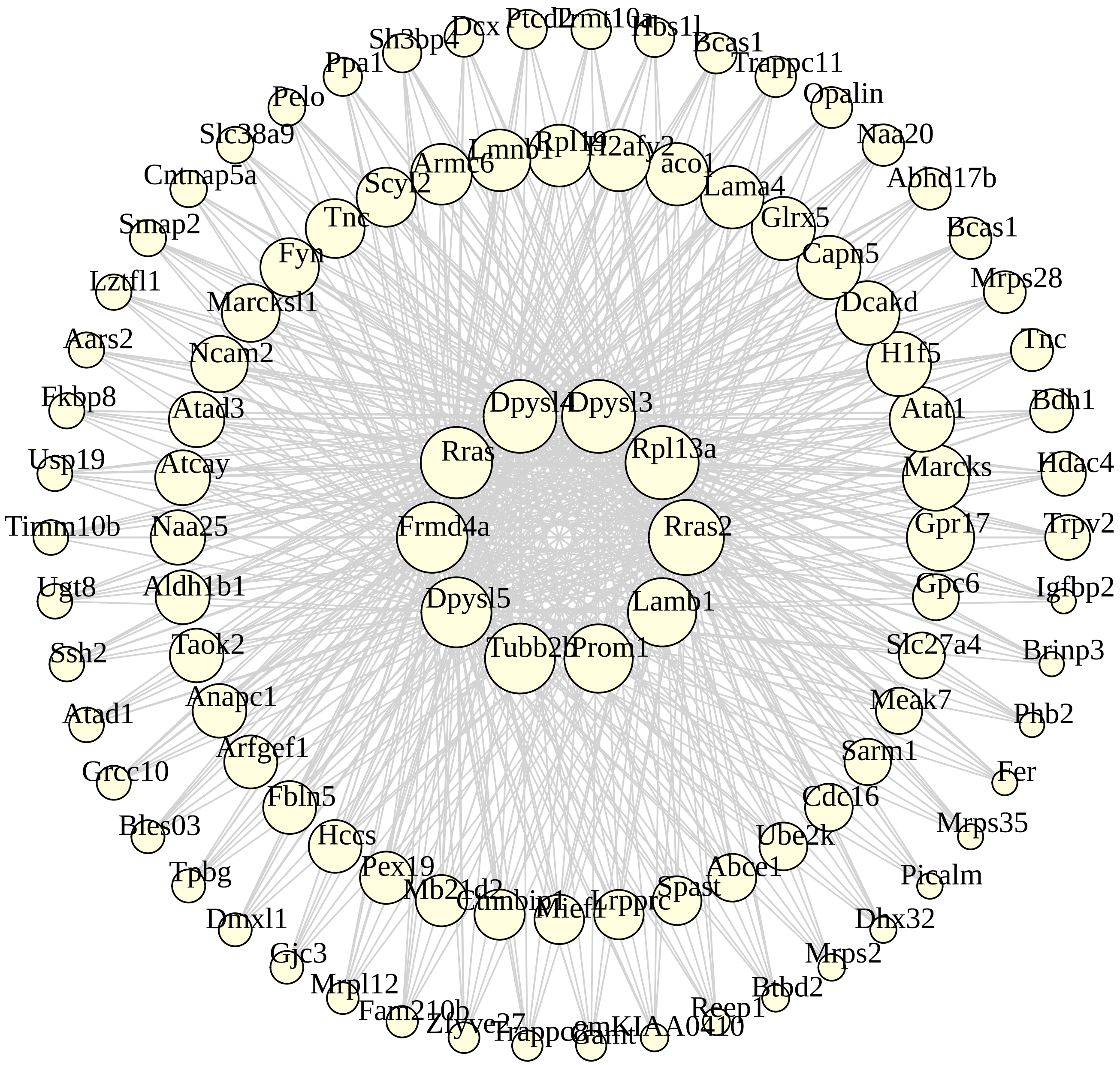

**M15 midnightblue module**

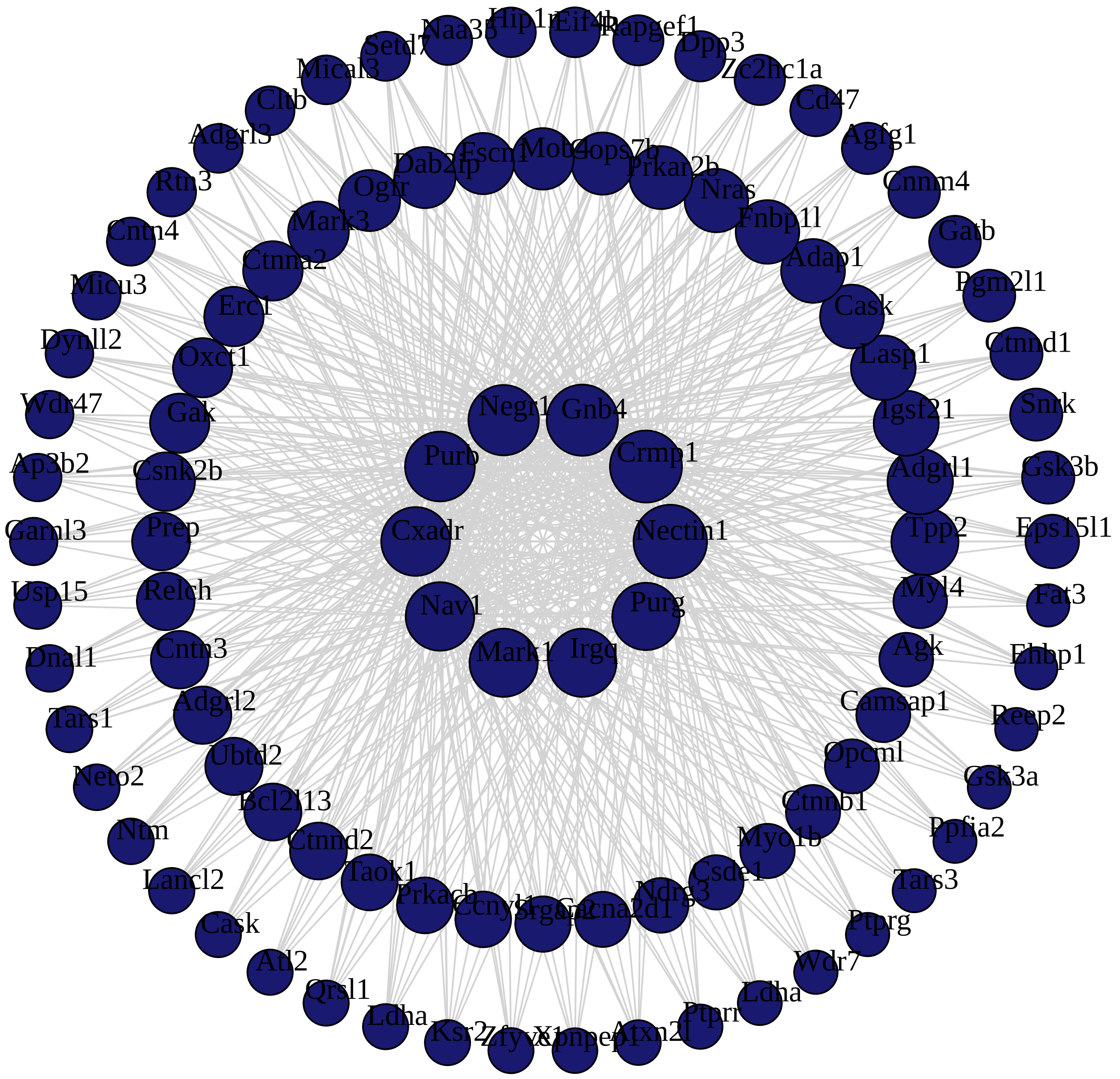

#### M30 steelblue module

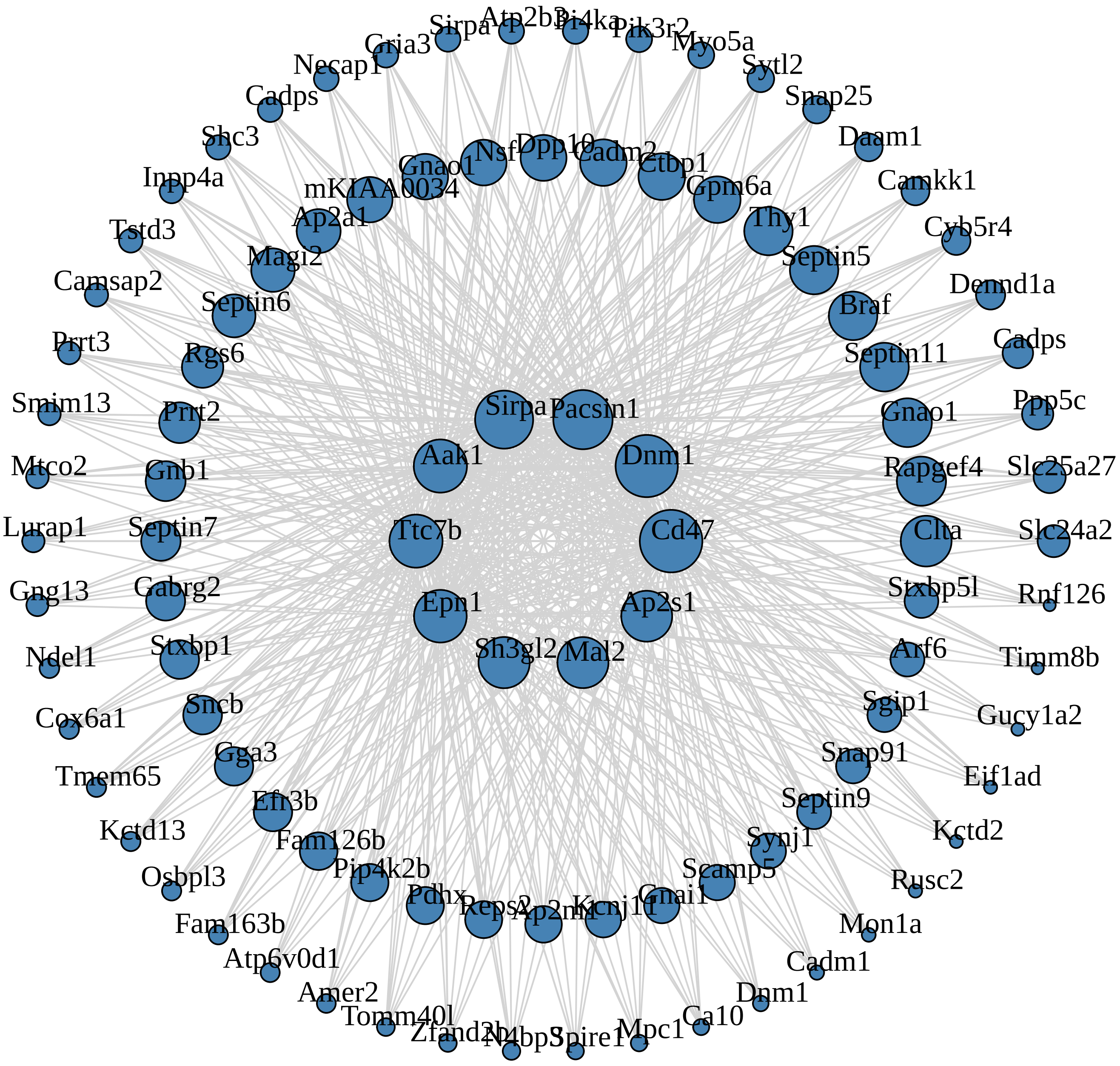

### M45 darkorange2 module

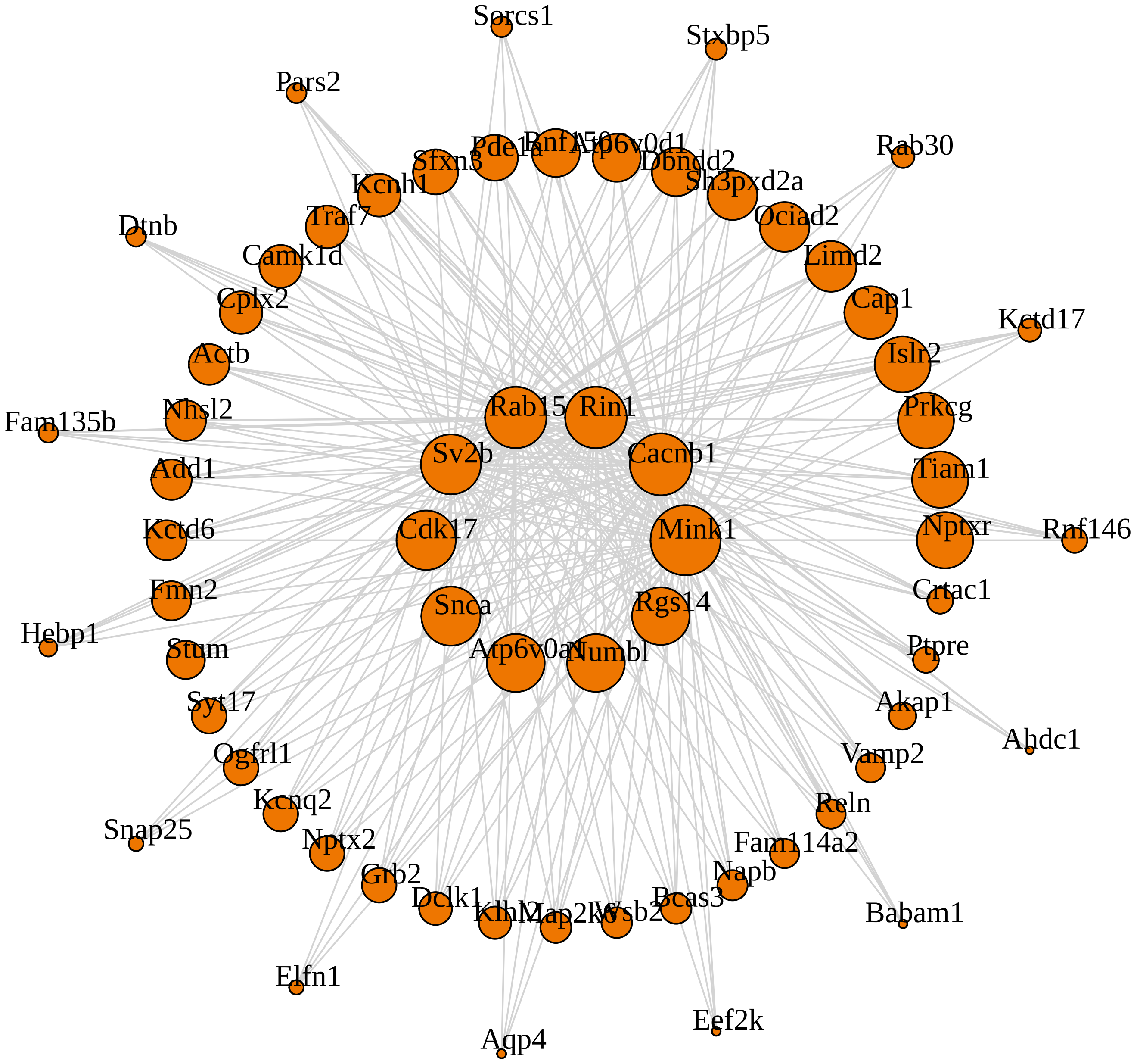

**M24 darkgrey module**

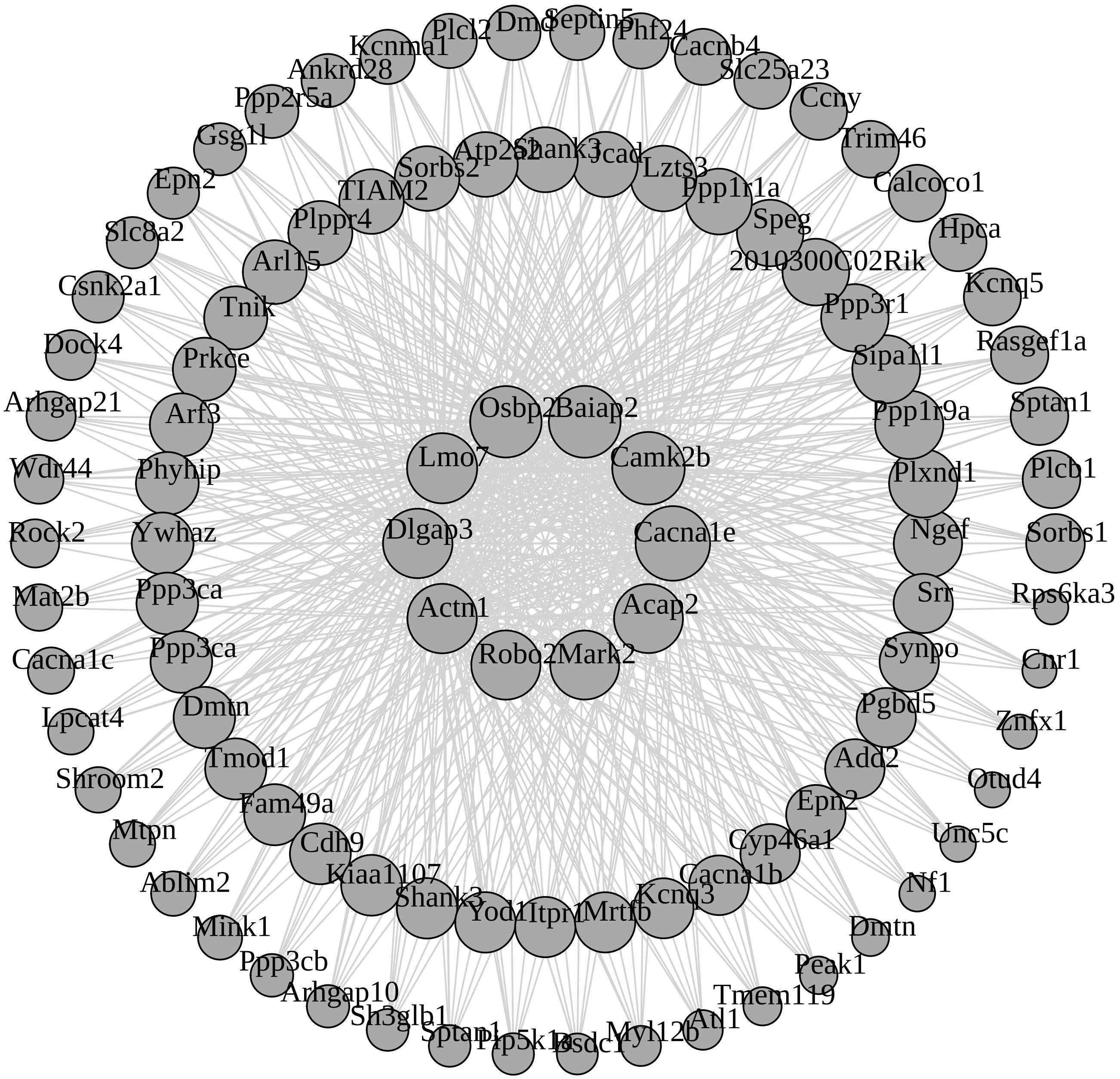

**M29 saddlebrown module**

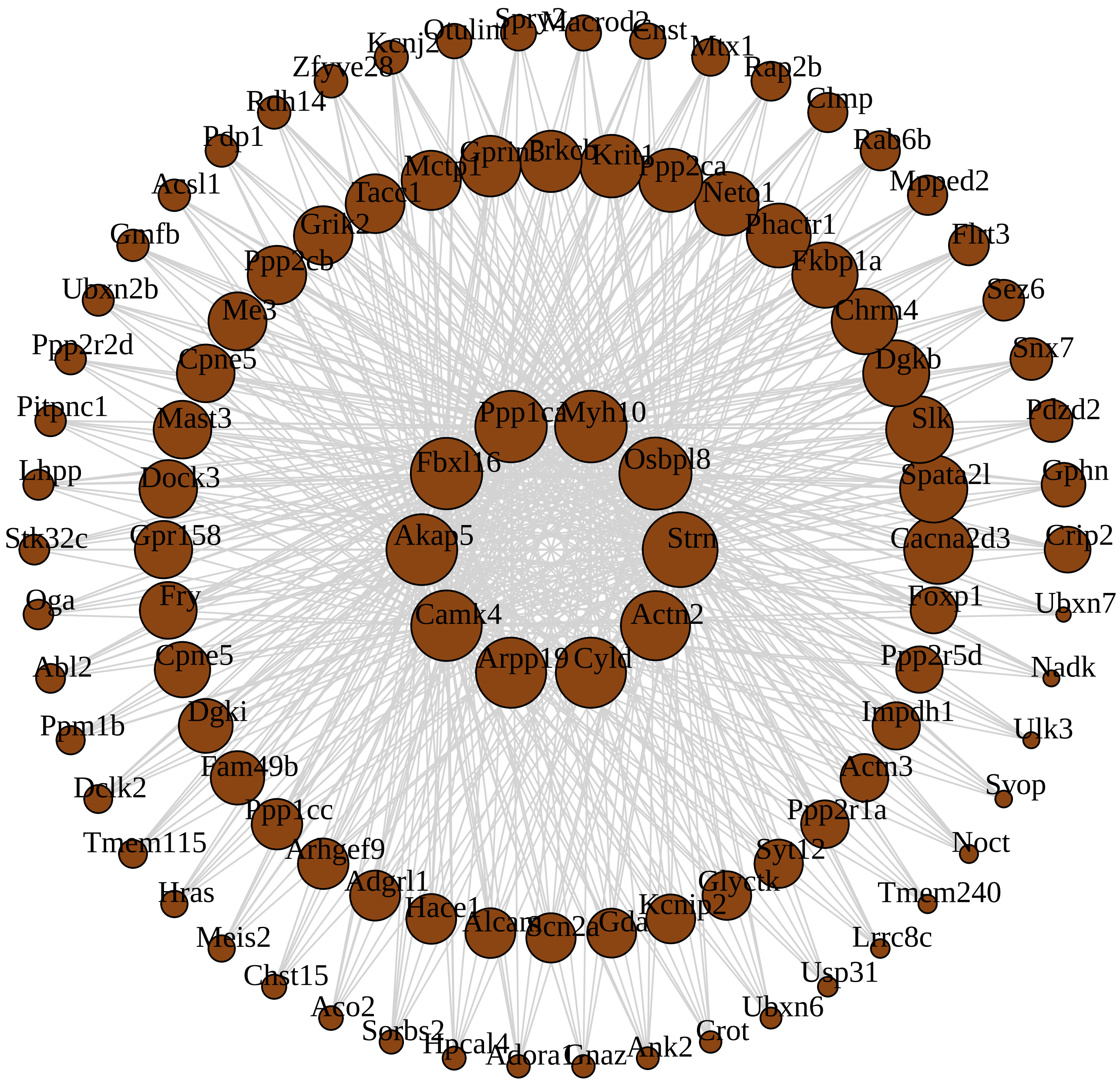

**M13 salmon module**

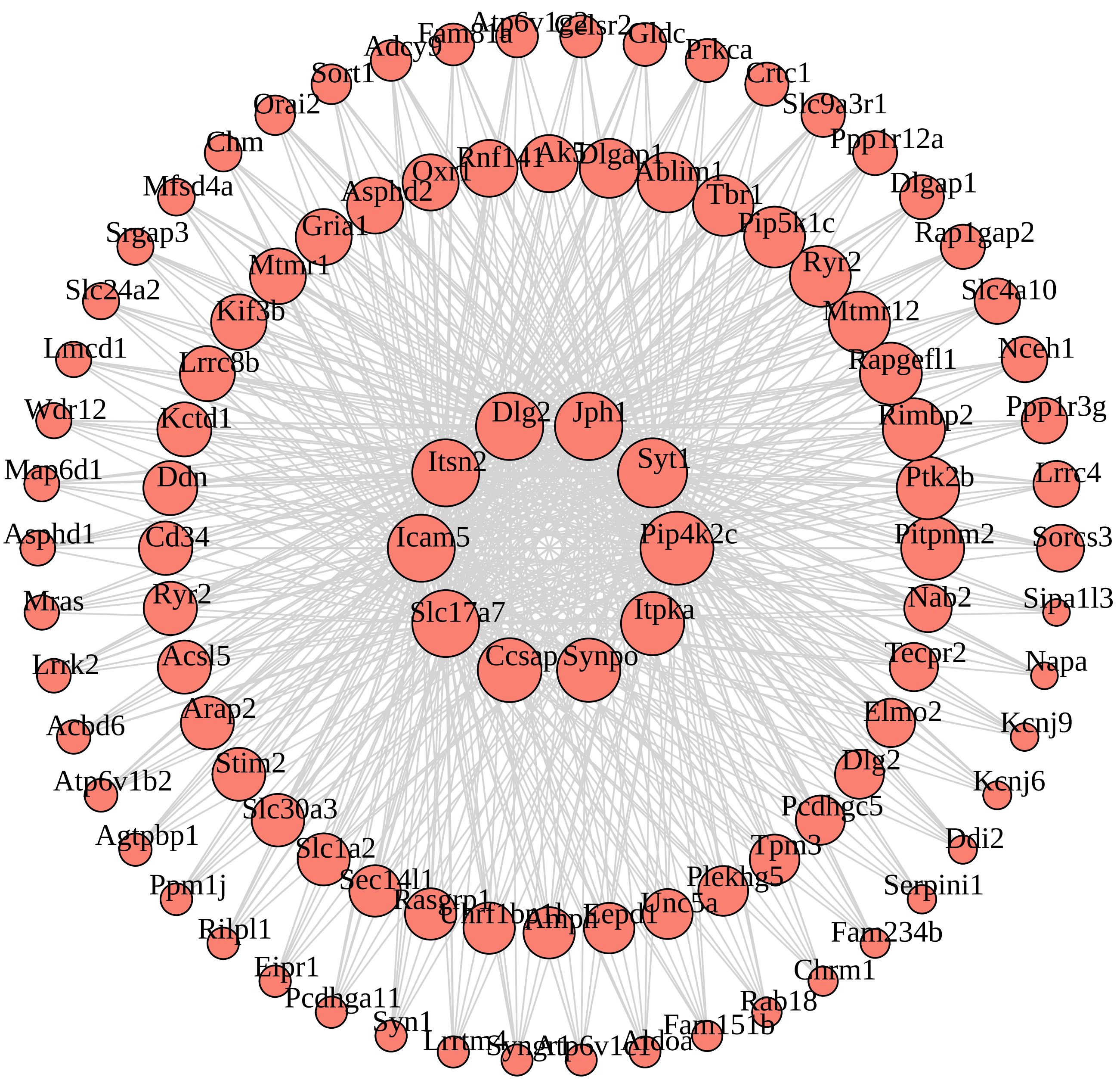

**M3 brown module**

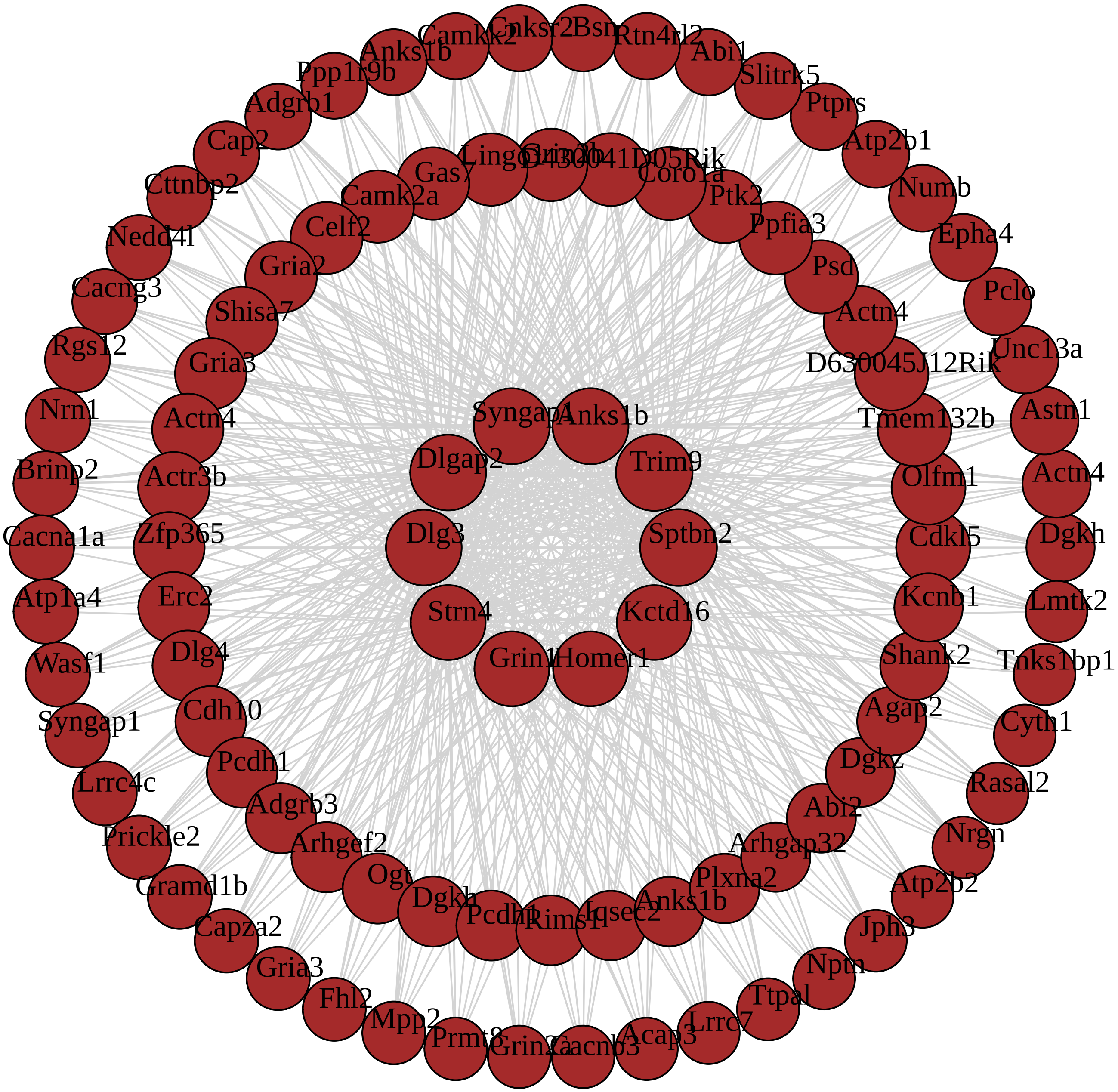

#### M5 green module

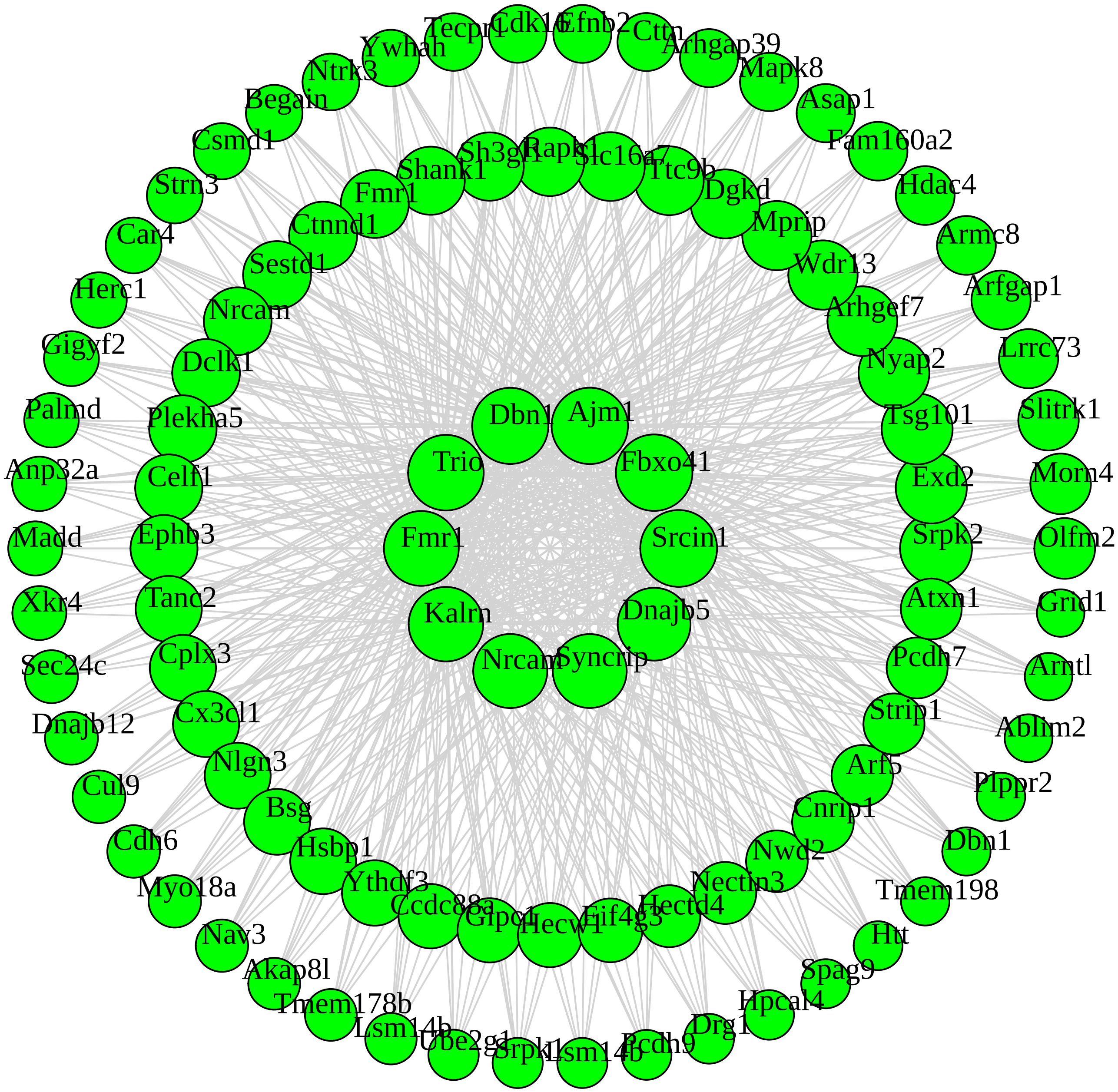

**M16 lightcyan module**

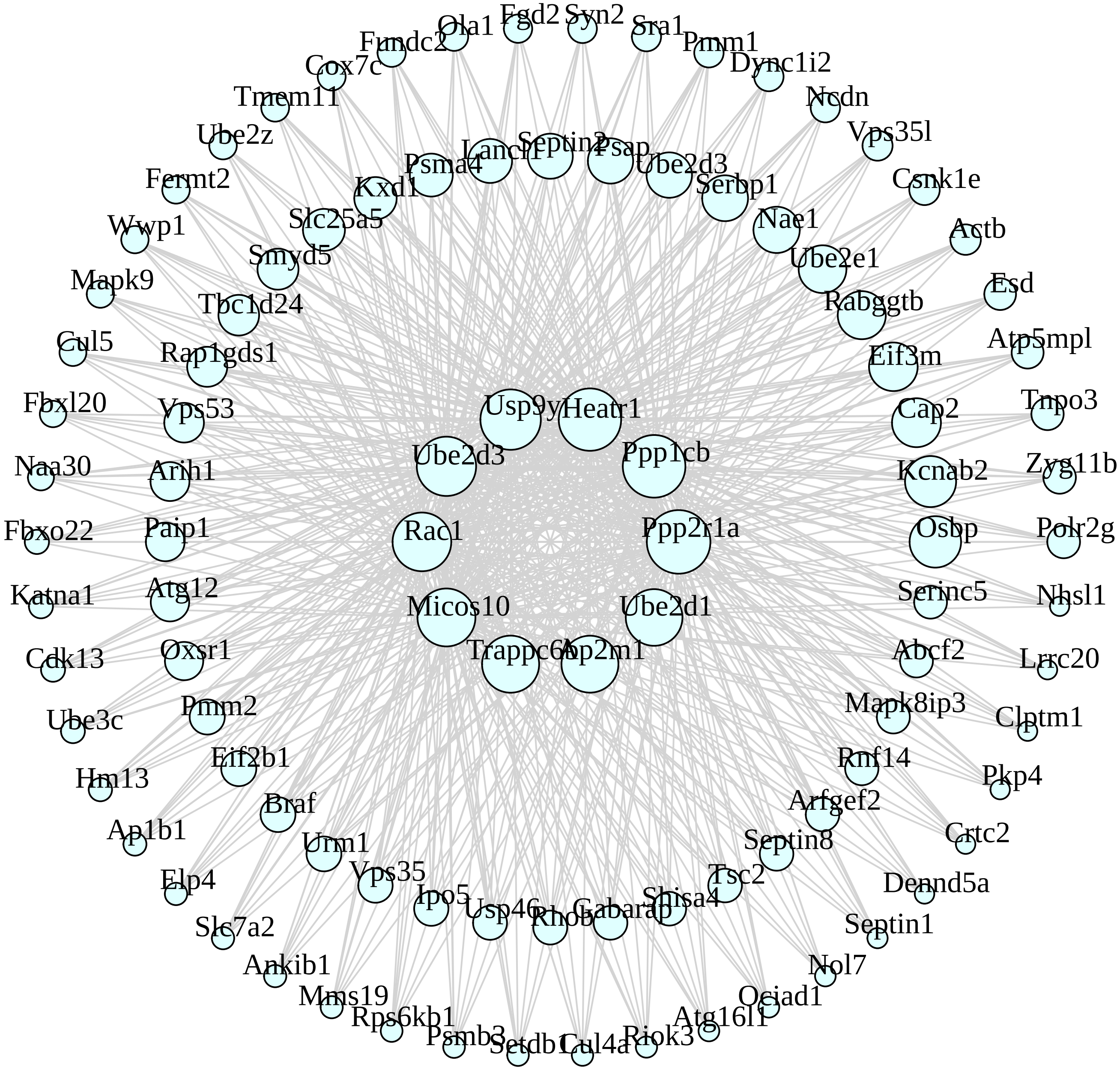

**M25 orange module**

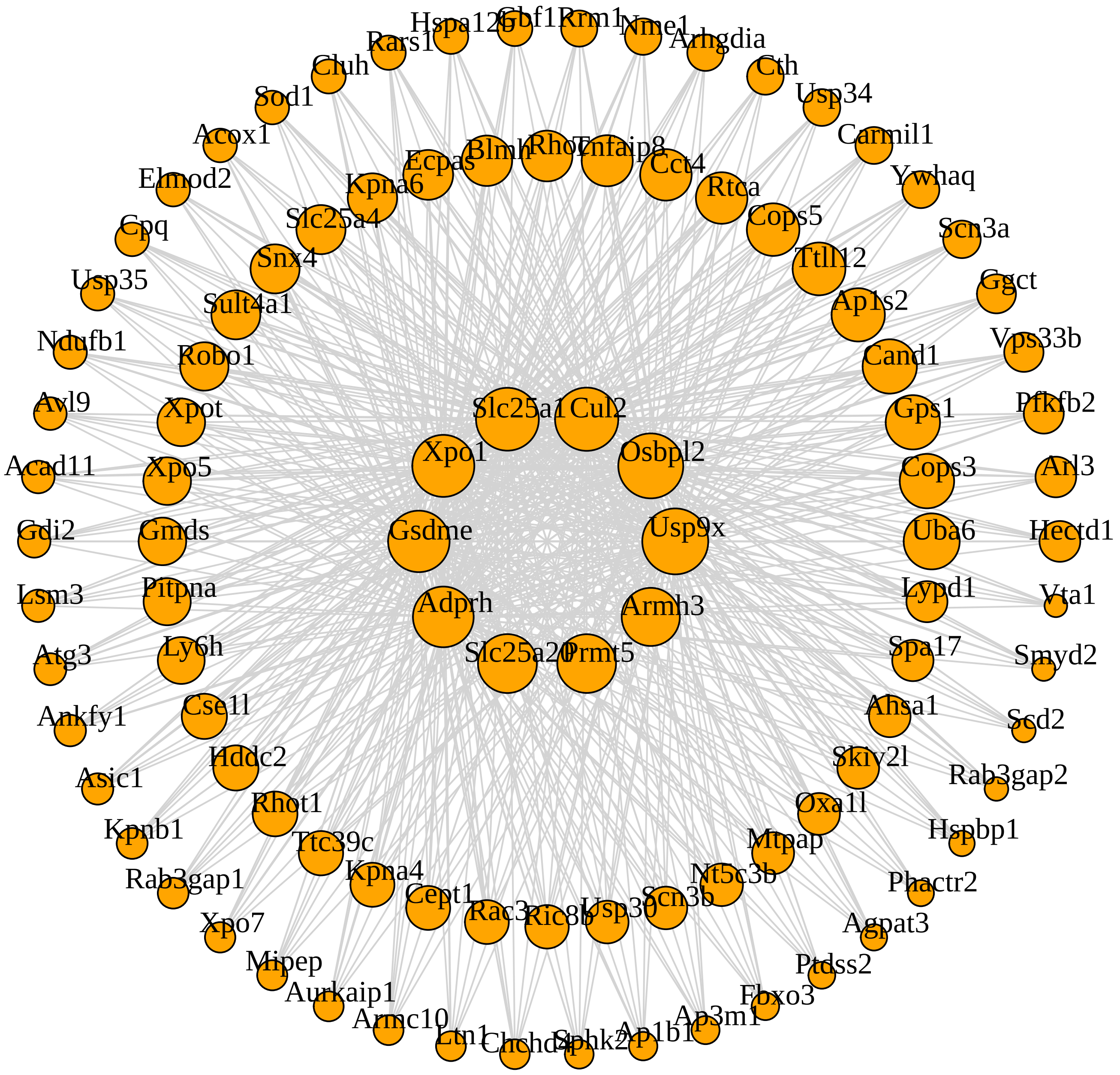

M35 sienna3 module

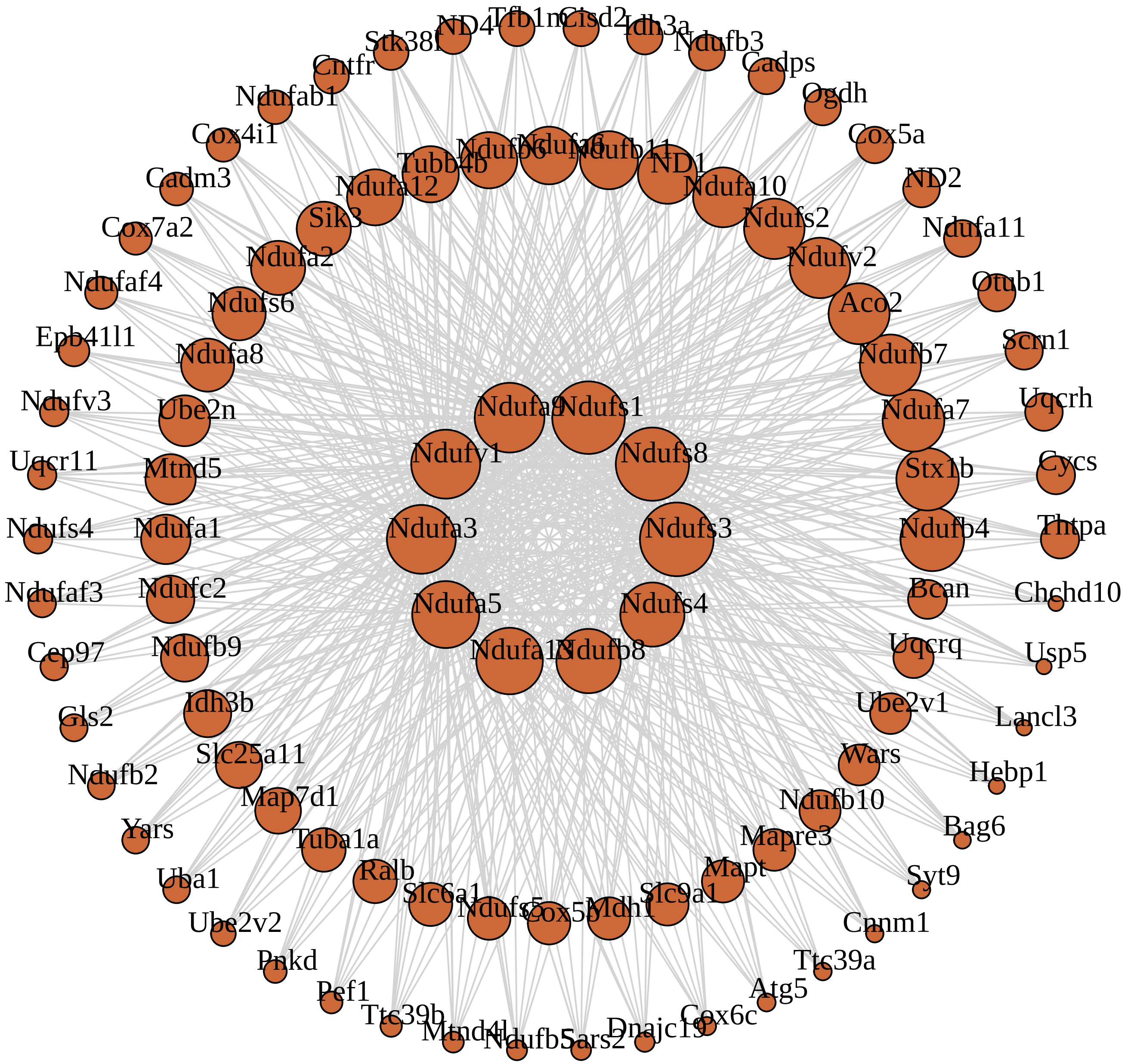

**M10 purple module**

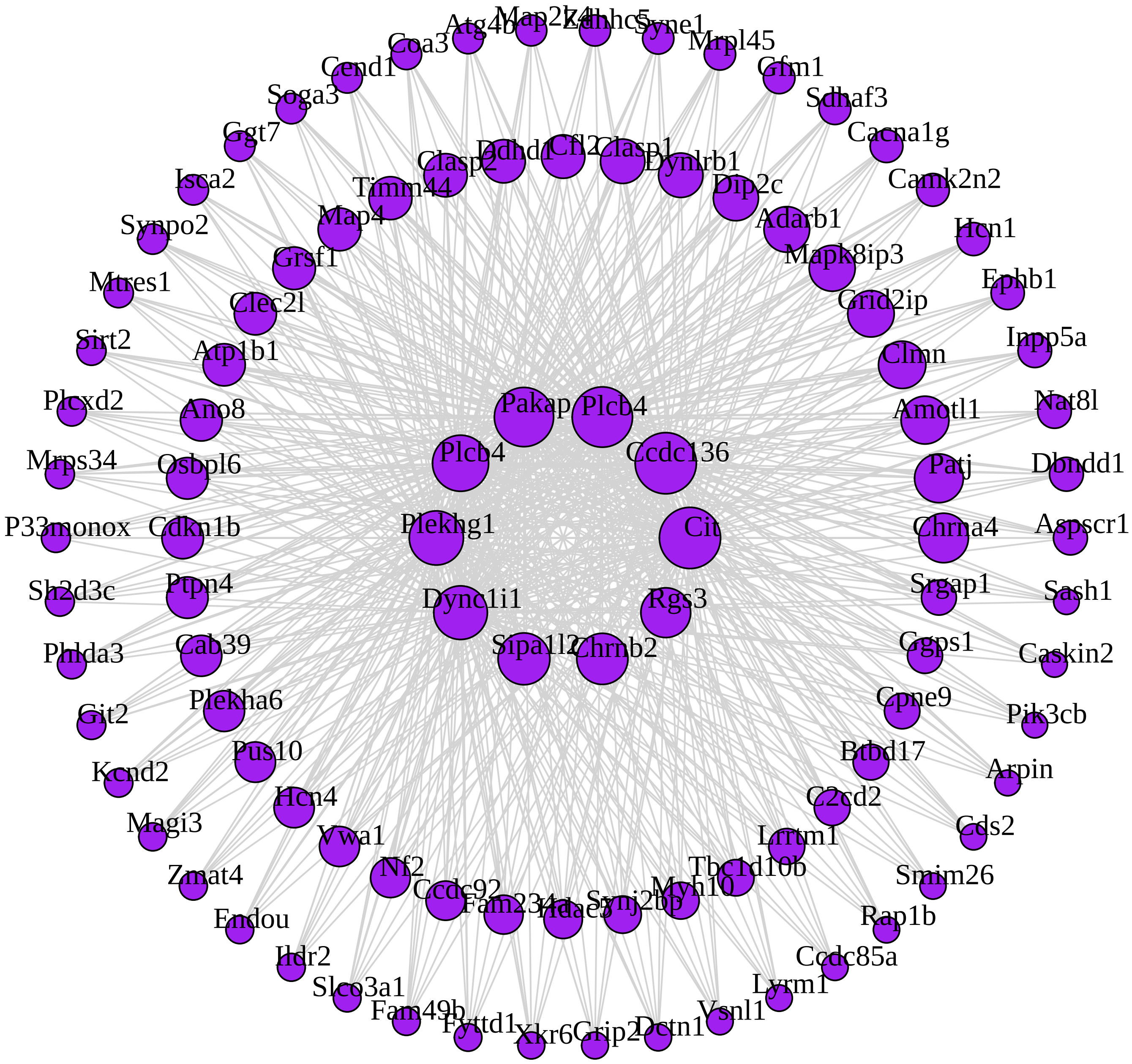

**M32 violet module**

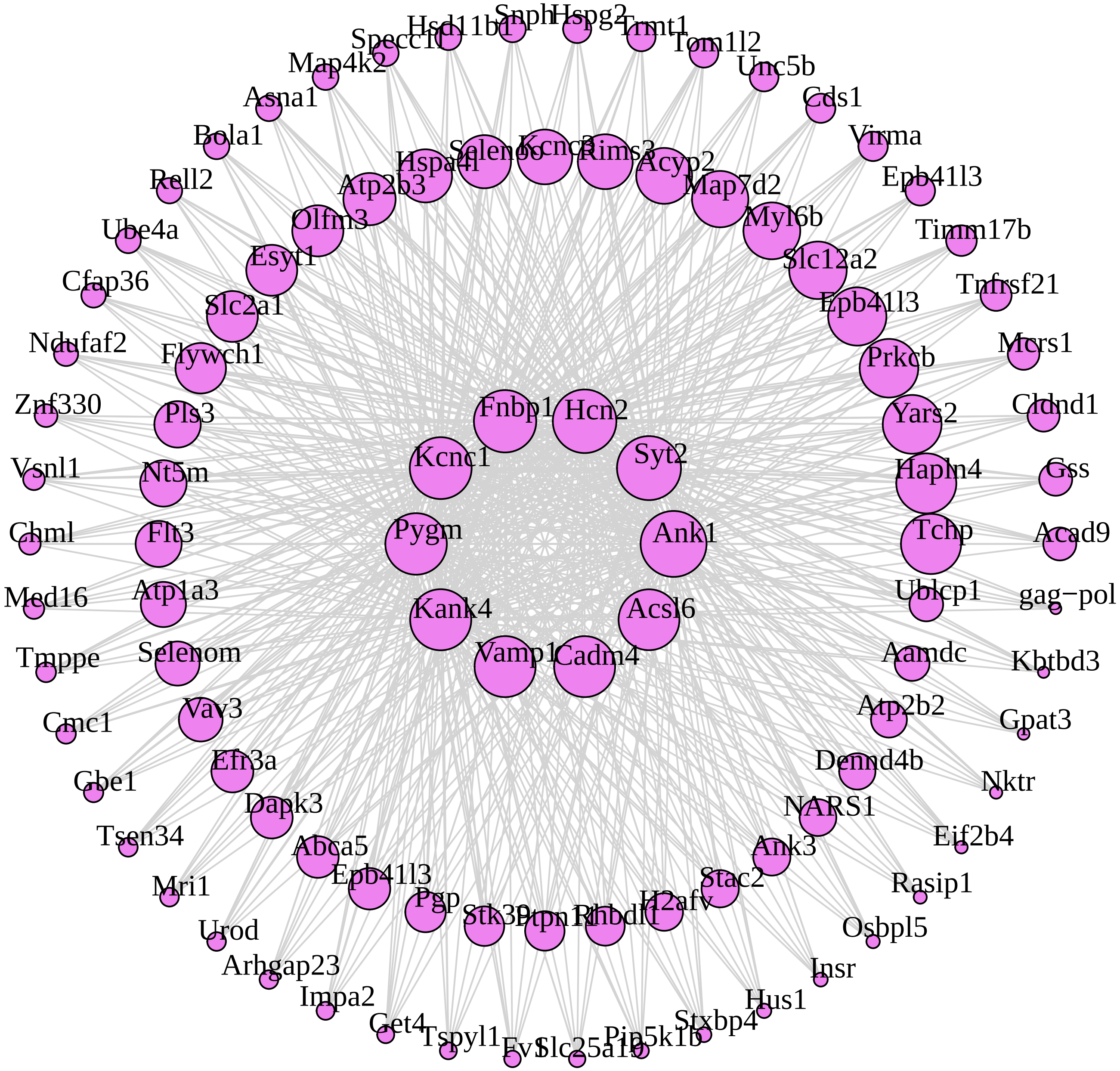

**M14 cyan module**

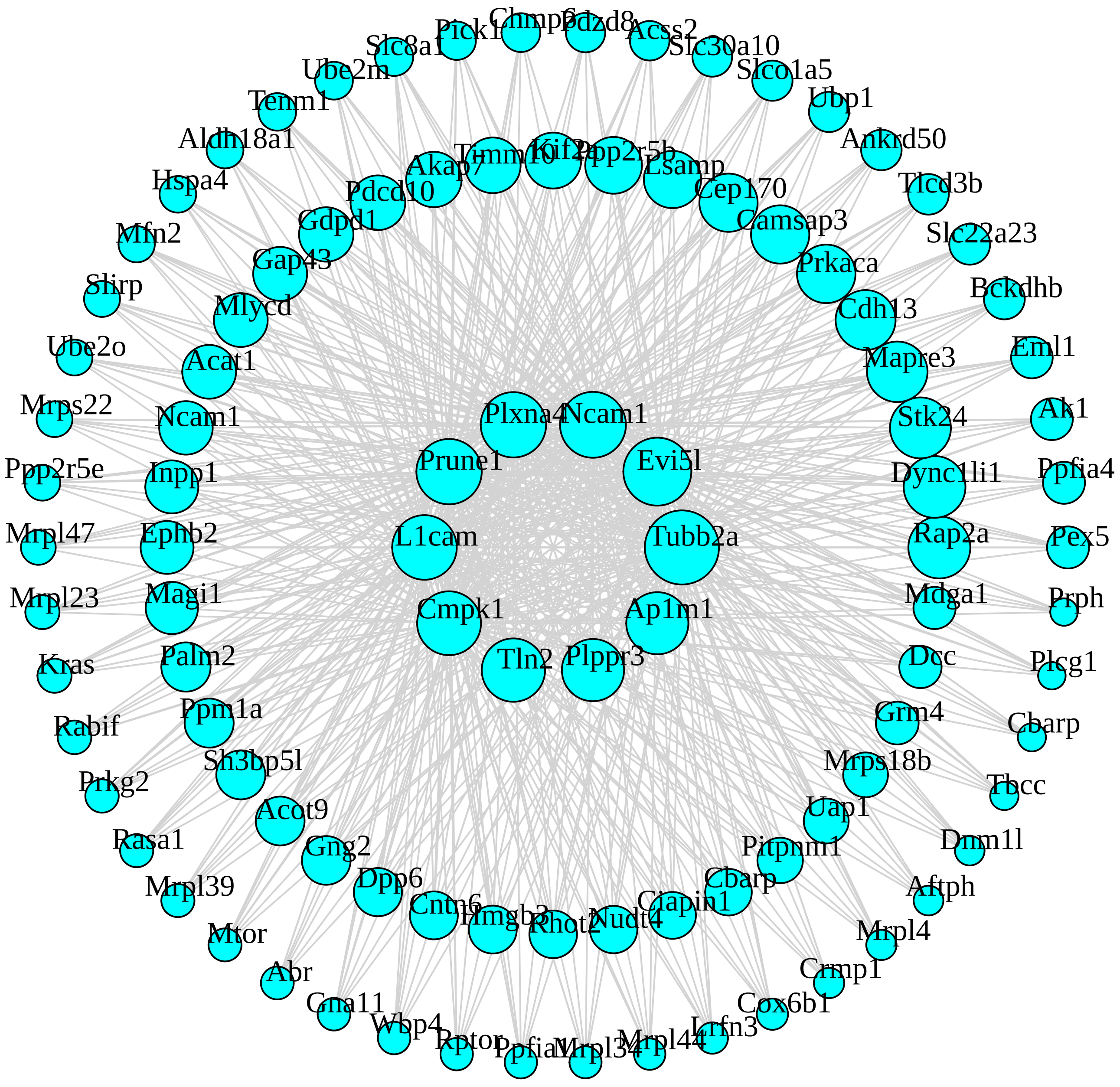

**M11 greenyellow module**

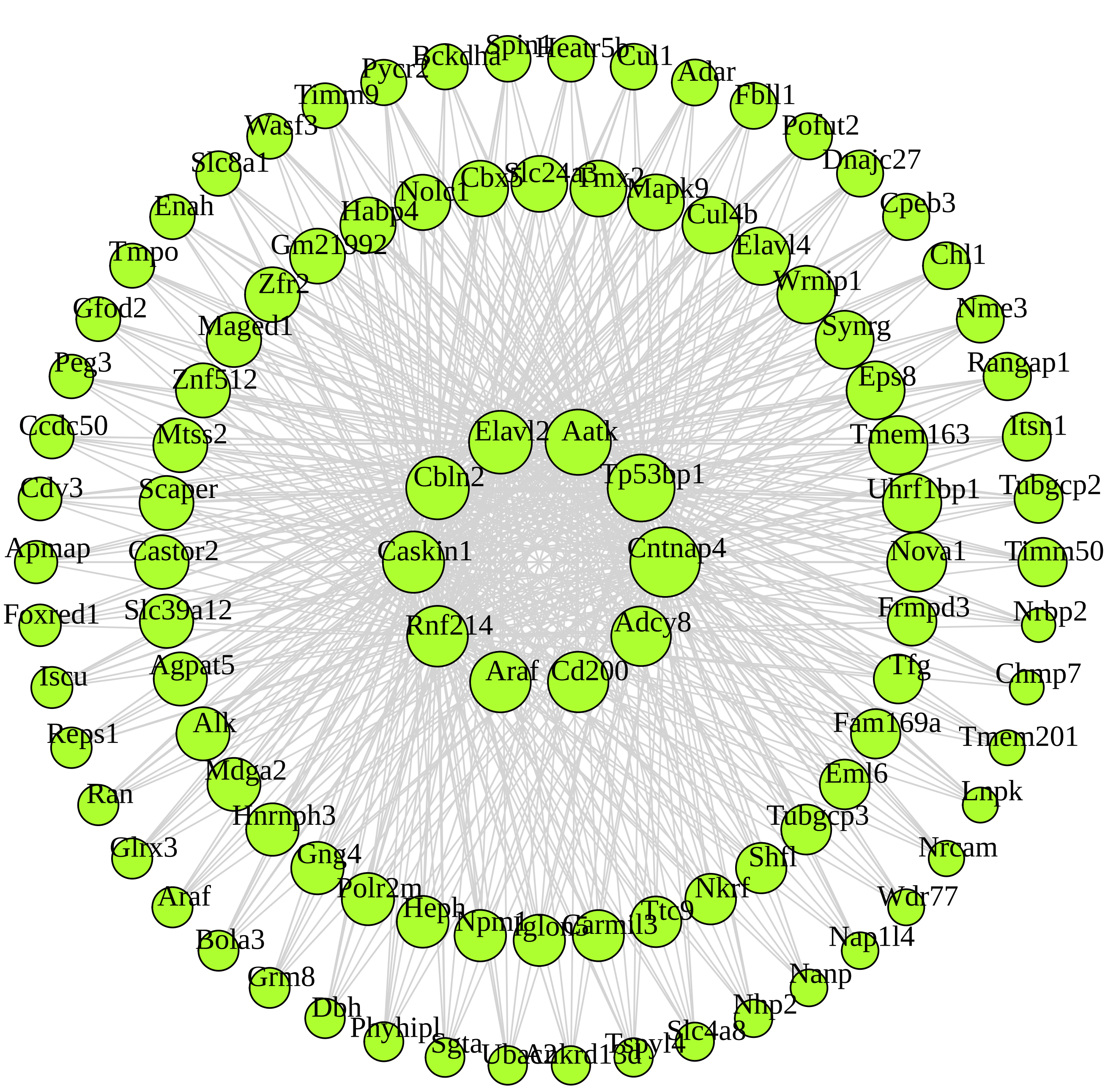

**M6 red module**

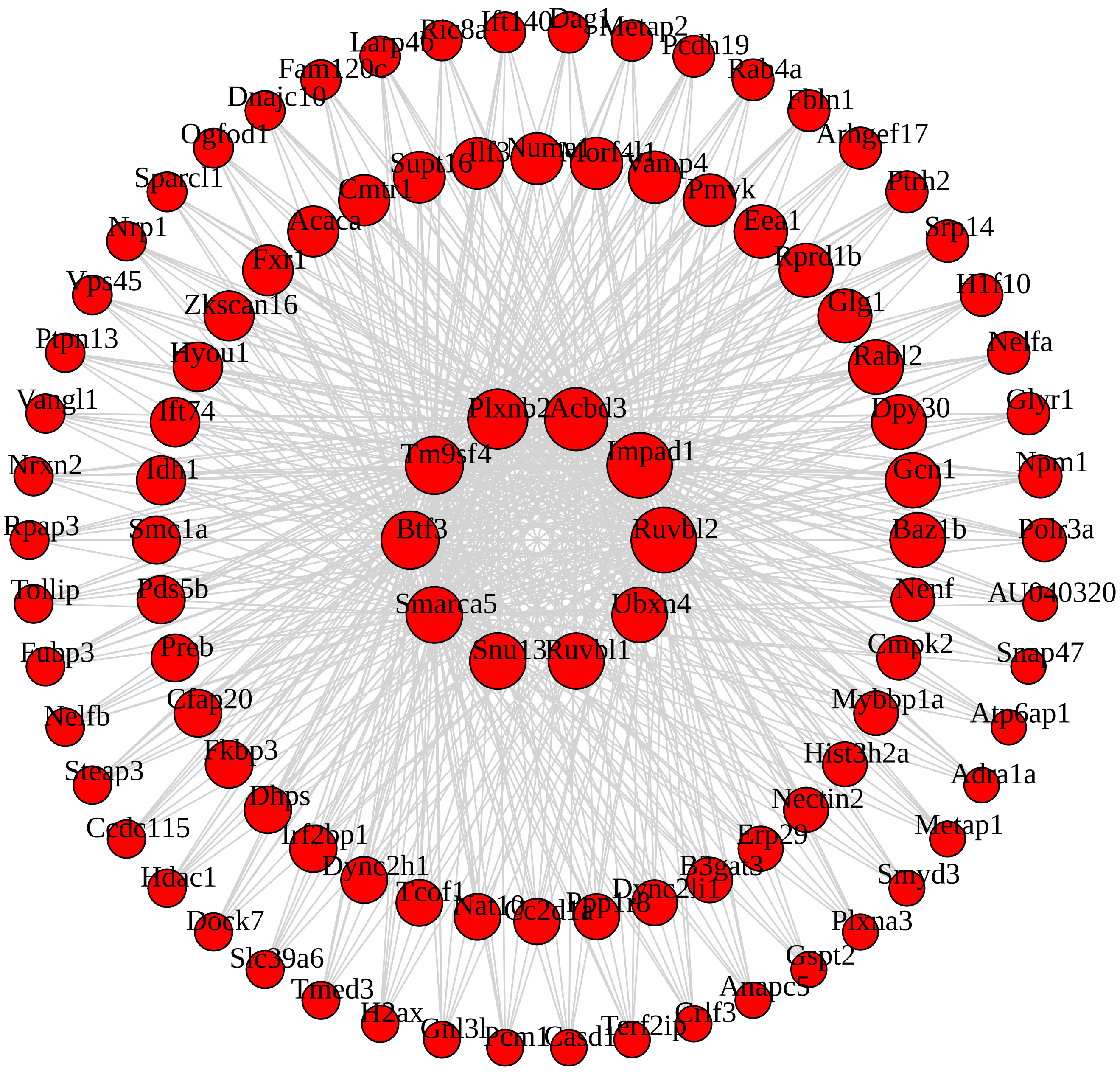

**M2 blue module**

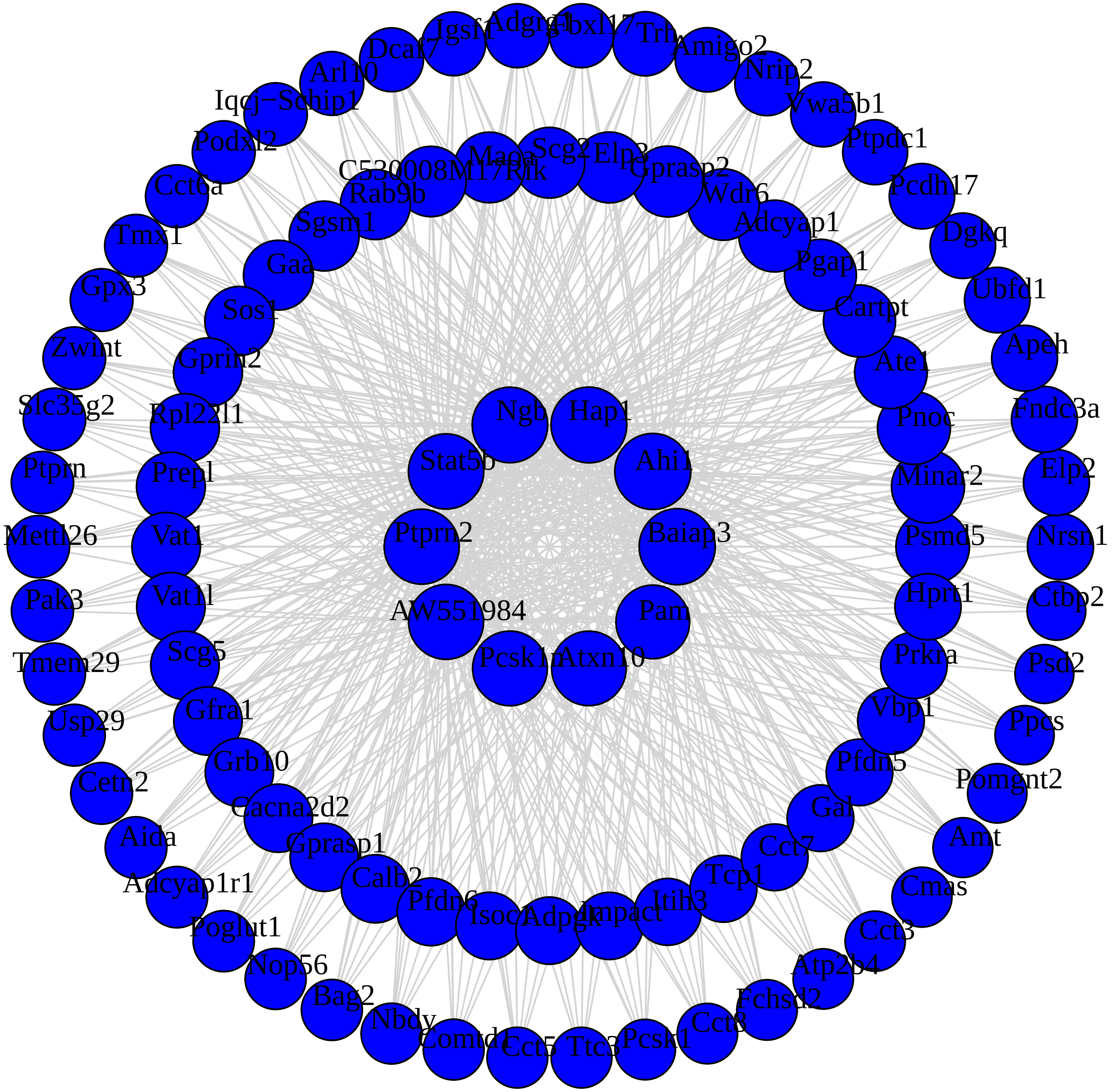

#### M8 pink module

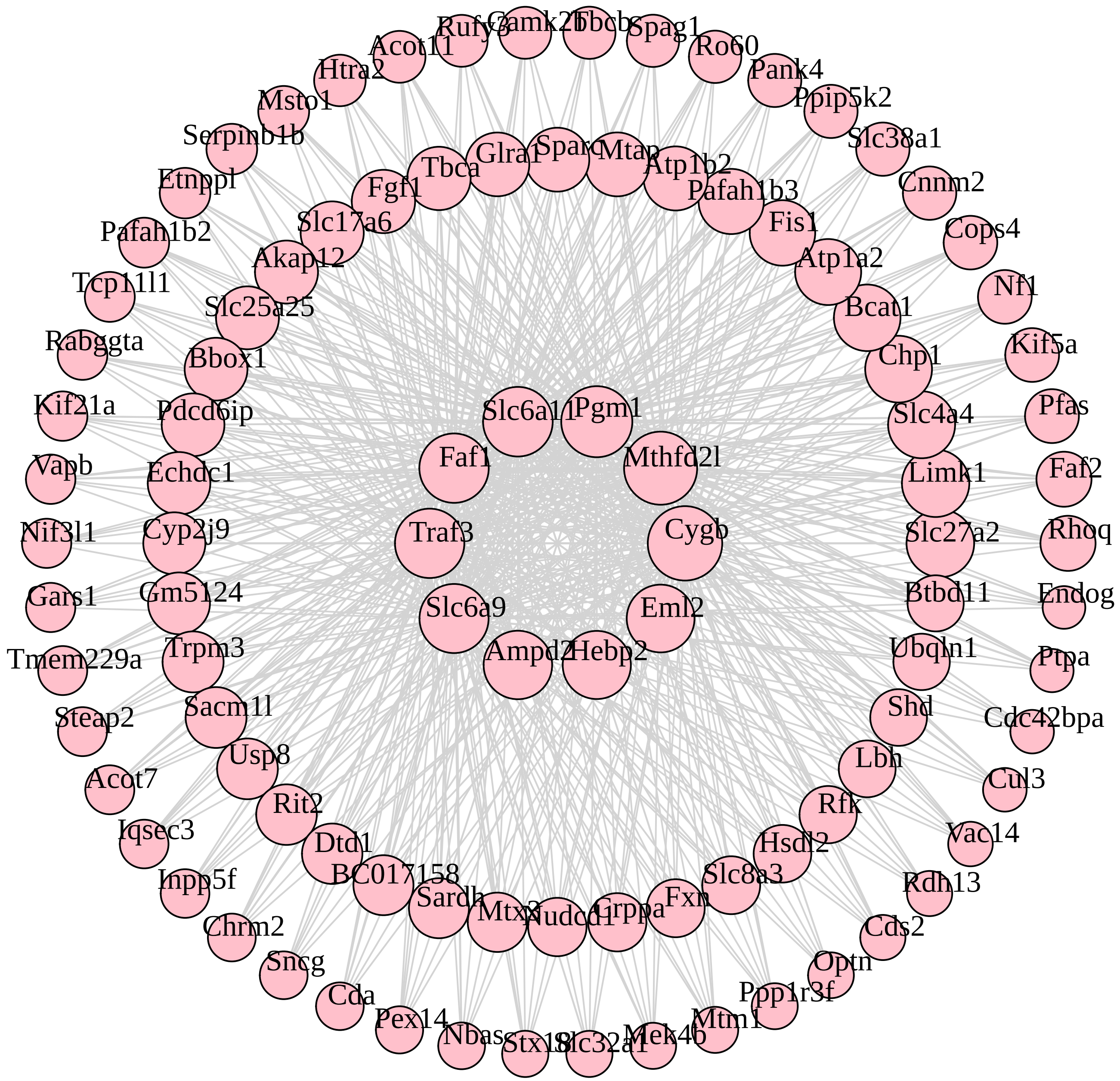

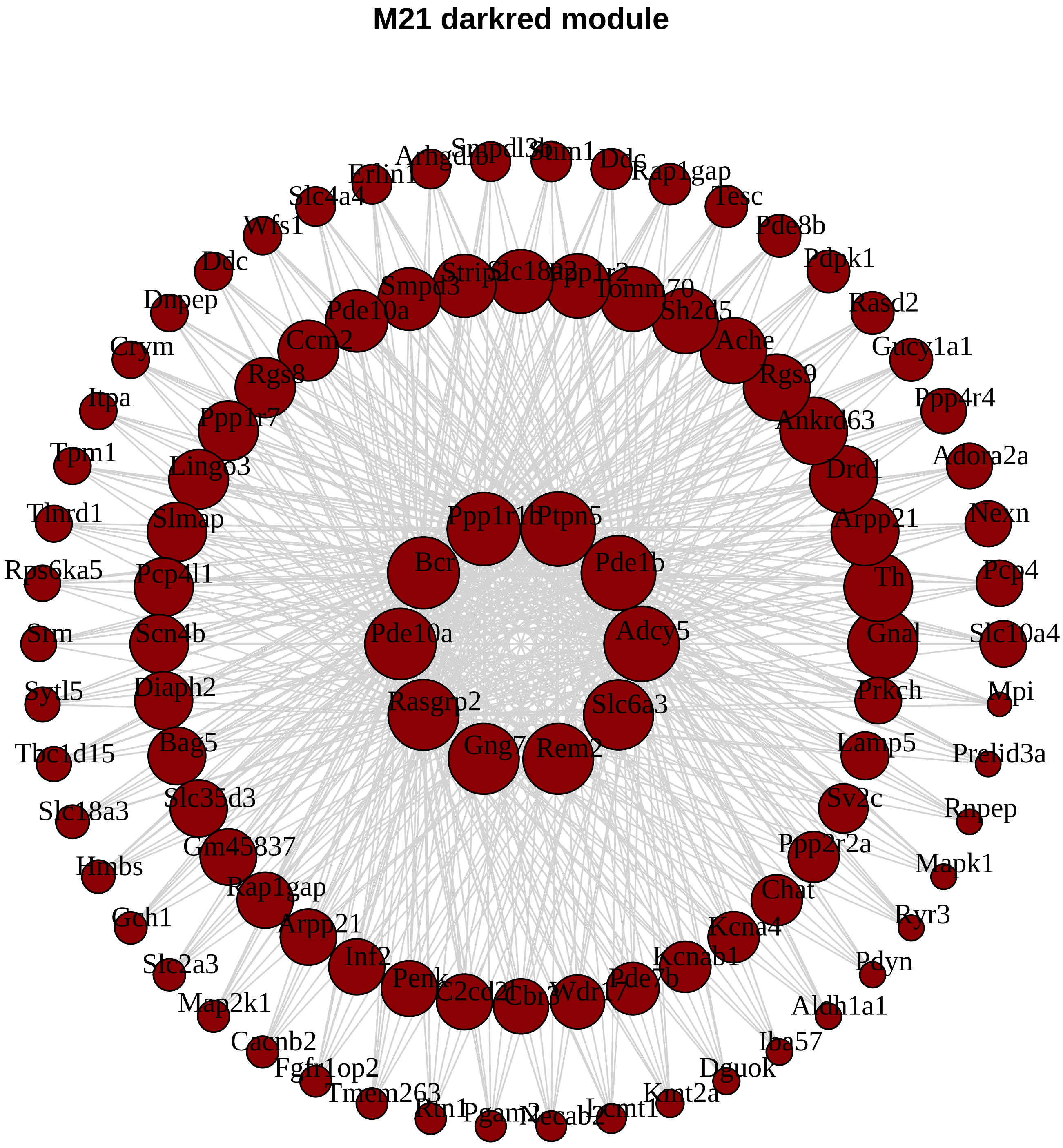
M21 darkred module
Erlin1
Slc4a4
Wfs1
Ddc
Dnpep
Crym
Itpa
Tpm1
Tlnrd1
Rps6ka5
Srm
Sytl5
Tbc1d15
Slc18a3
Hmbs
Gch1
Slc2a3
Map2k1
Cacnb2
Fgfr1op2
Tmem263
Rap1gap
Tesc
Pde8b
Pdpk1
Rasd2
Gucy1a1
Ppp4r4
Adora2a
Nexn
Pcp4
Slc10a4
Mpi
Prelid3a
Rnpep
Mapk1
Ryr3
Pdyn
Aldh1a1
Iba57
Dguok
Kmt2a
Smpd3
Tomm70
Pde10a
Sh2d5
Ccm2
Ache
Rgs8
Rgs9
Ppp1r7
Ankrd63
Lingo3
Drd1
Slmap
Arpp21
Pcp4l1
Th
Scn4b
Gnal
Diaph2
Prkch
Bag5
Lamp5
Slc35d3
Sv2c
Gm45837
Ppp2r2a
Rap1gap
Chat
Arpp21
Kcna4
Inf2
Kcnab1
Penk
Ppp1r1b
Ptpn5
Bcr
Pde1b
Pde10a
Adcy5
Rasgrp2
Slc6a3
Gng7
Rem2

**M48 darkslateblue module**

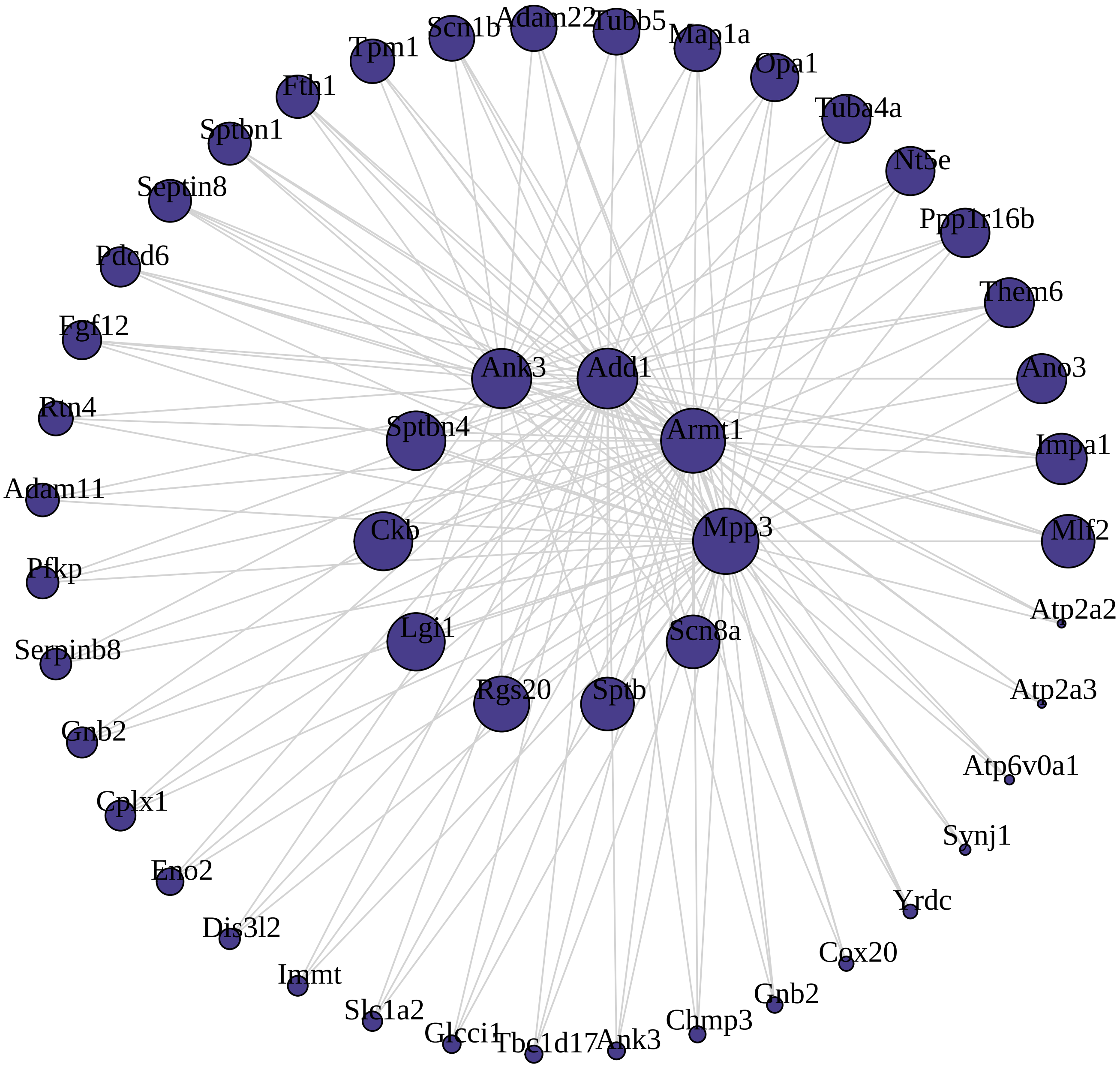

### M41 lightsteelblue1 module

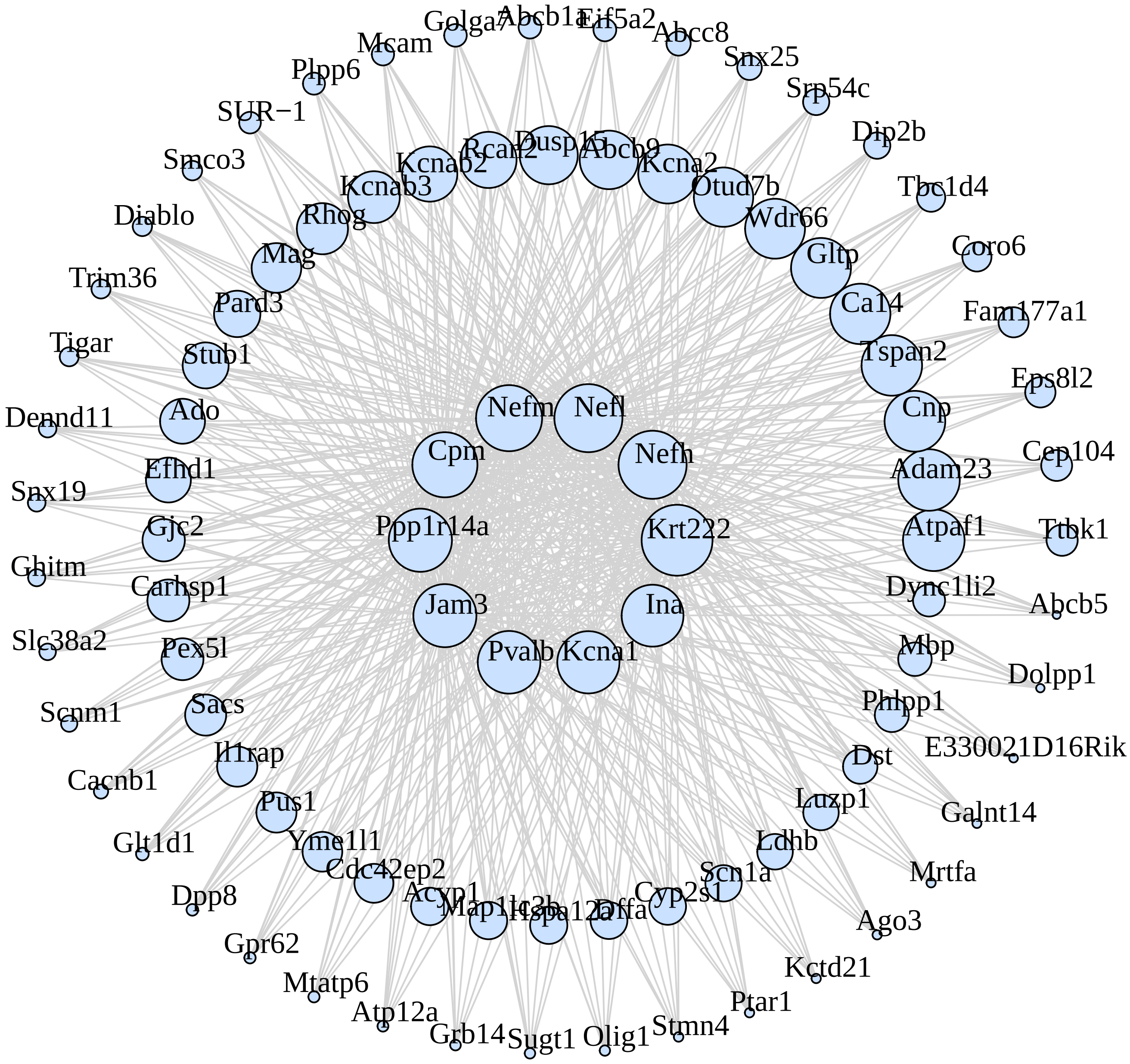

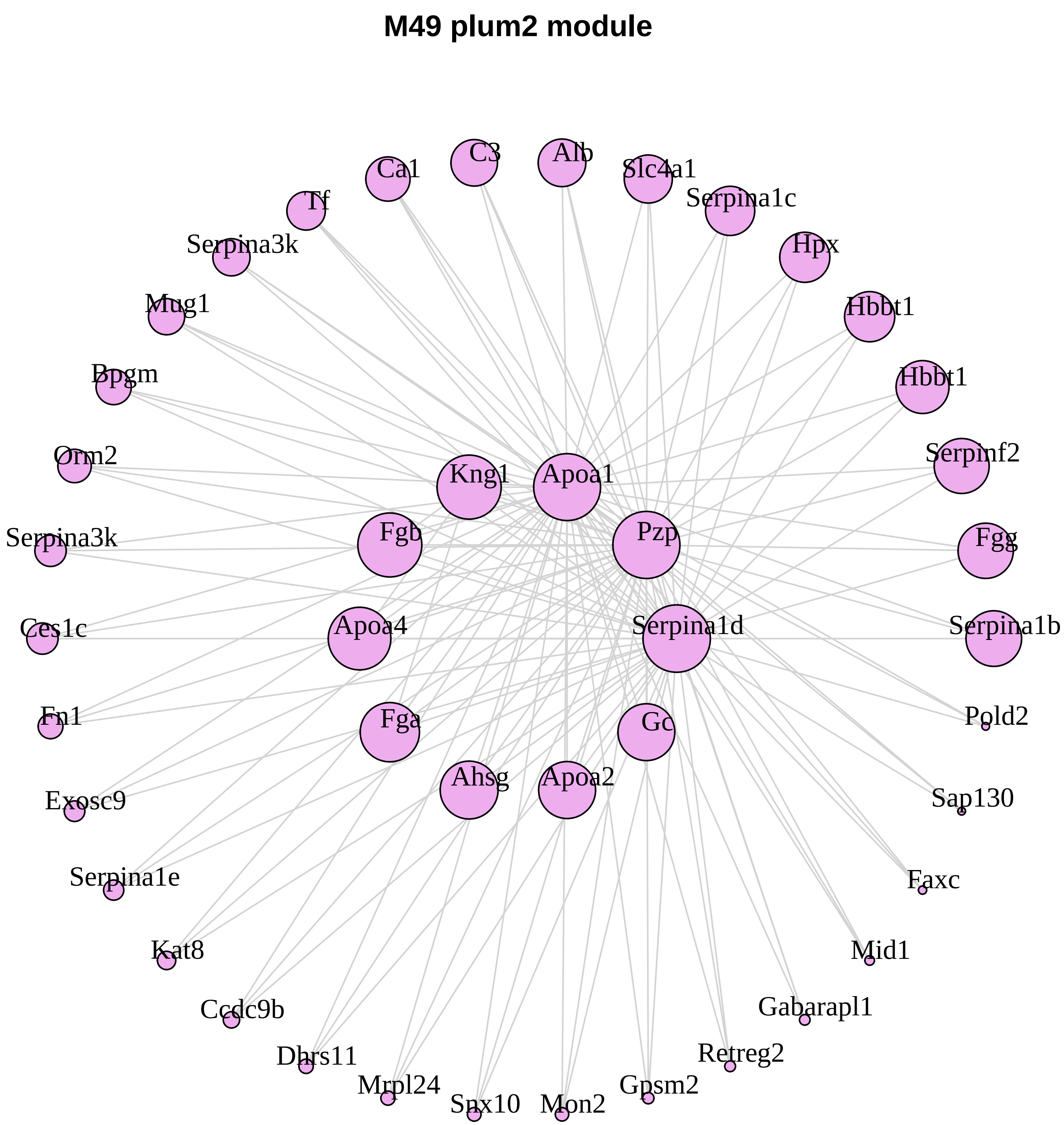
M49 plum2 module
C3
Alb
Ca1
Slc4a1
Tf
Serpina1c
Serpina3k
Hpx
Mug1
Hbbt1
Bpgm
Hbbt1
Orm2
Kng1
Apoa1
Serpinf2
Serpina3k
Fgb
Pzp
Fgg
Ces1c
Apoa4
Serpina1d
Serpina1b
Fn1
Fga
Gc
Pold2
Exosc9
Ahsg
Apoa2
Sap130
Serpina1e
Faxc
Kat8
Mid1
Ccdc9b
Gabarapl1
Dhrs11
Retreg2
Mrpl24
Gpsm2
Snx10
Mon2

M47 bisque4 module

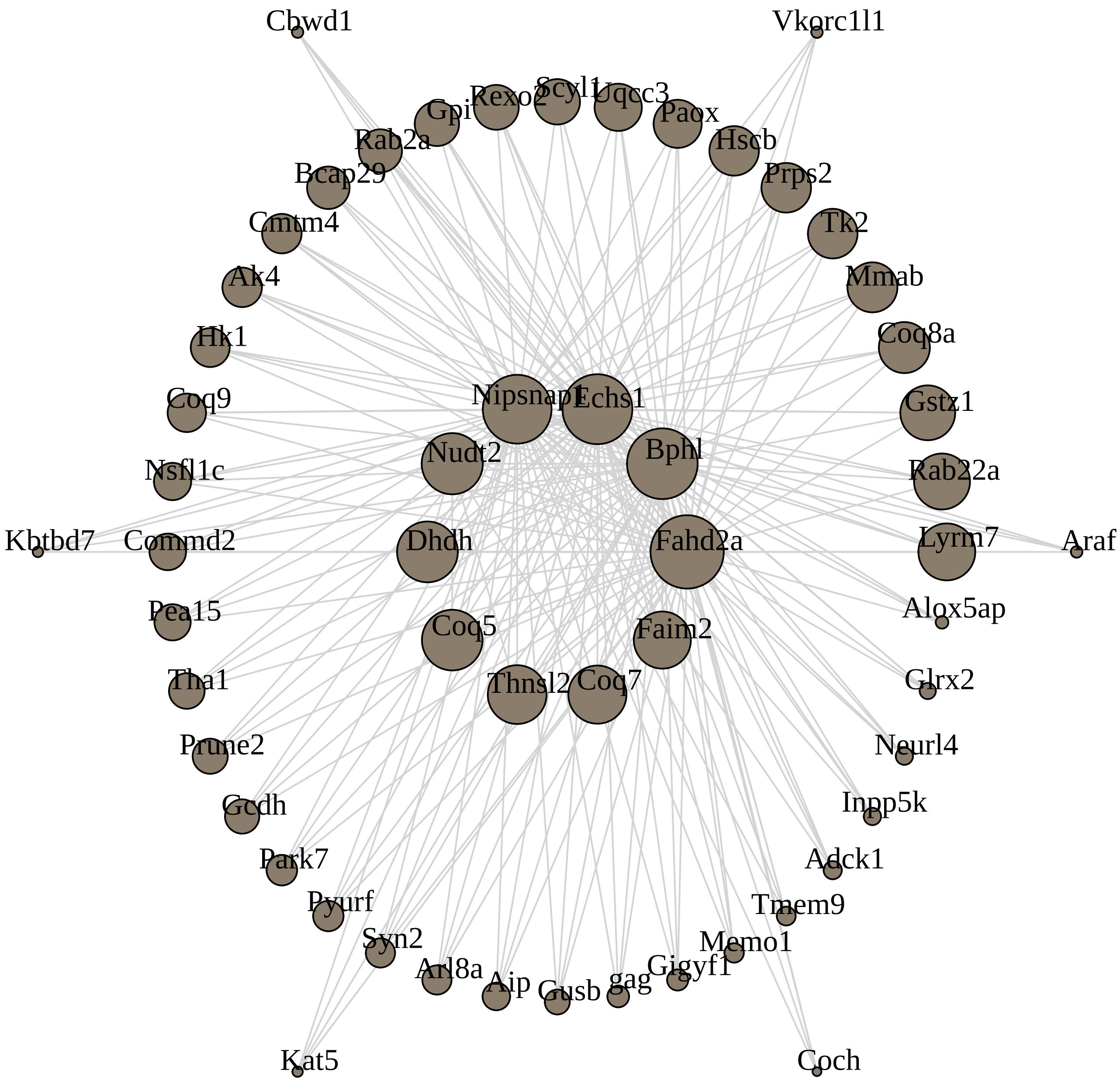

**M37 skyblue3 module**

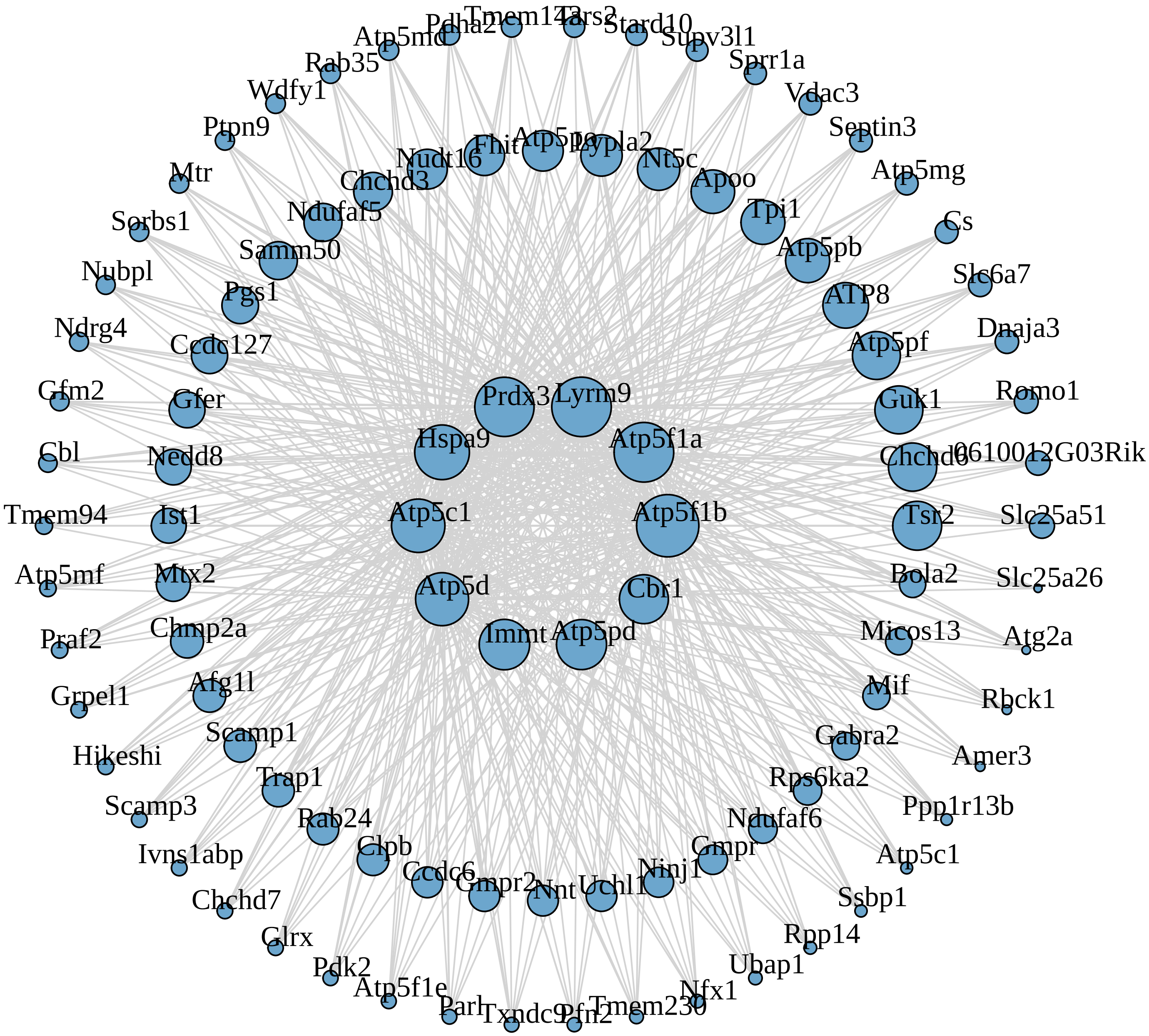

**M33 darkolivegreen module**

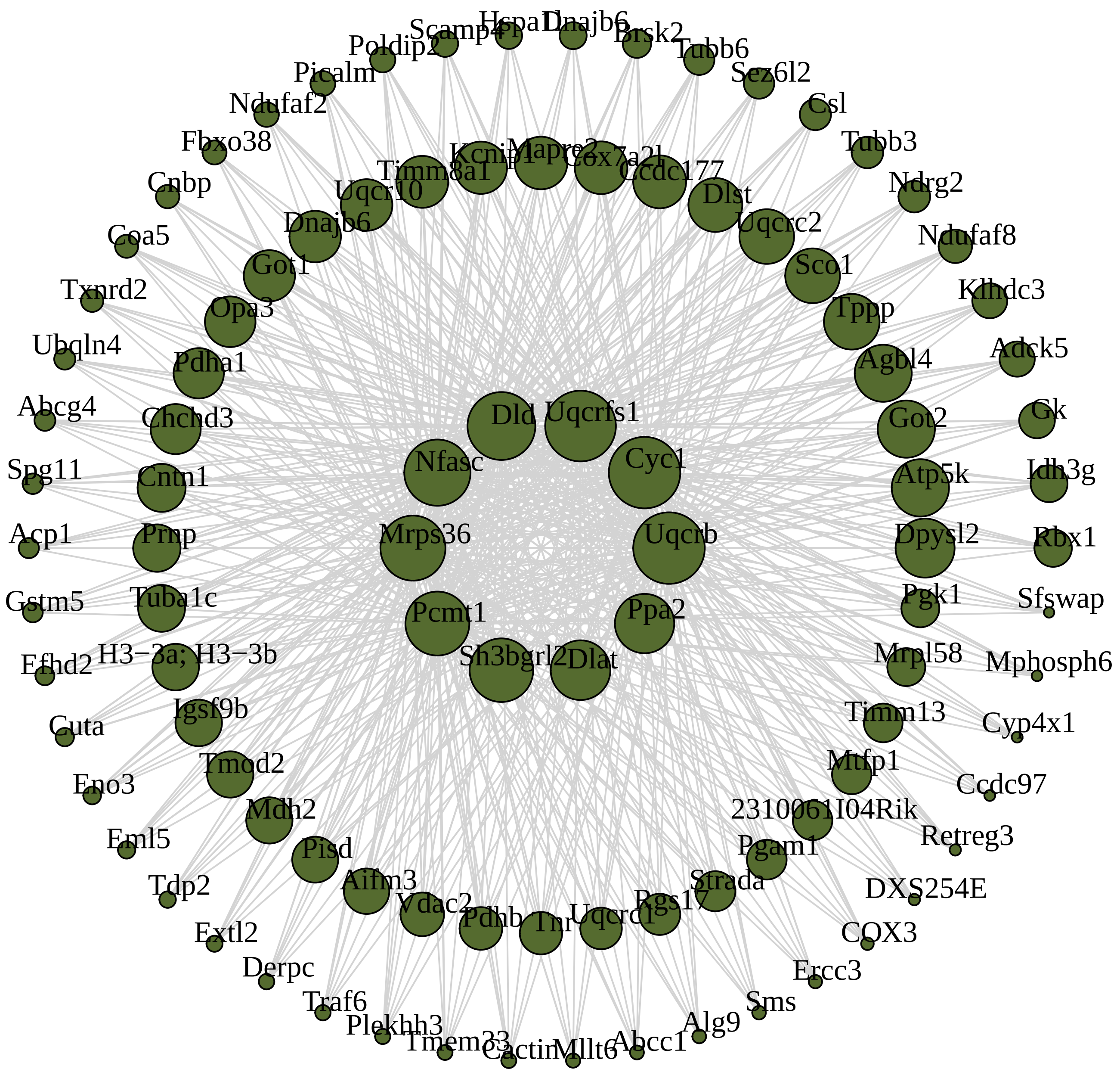

**M9 magenta module**

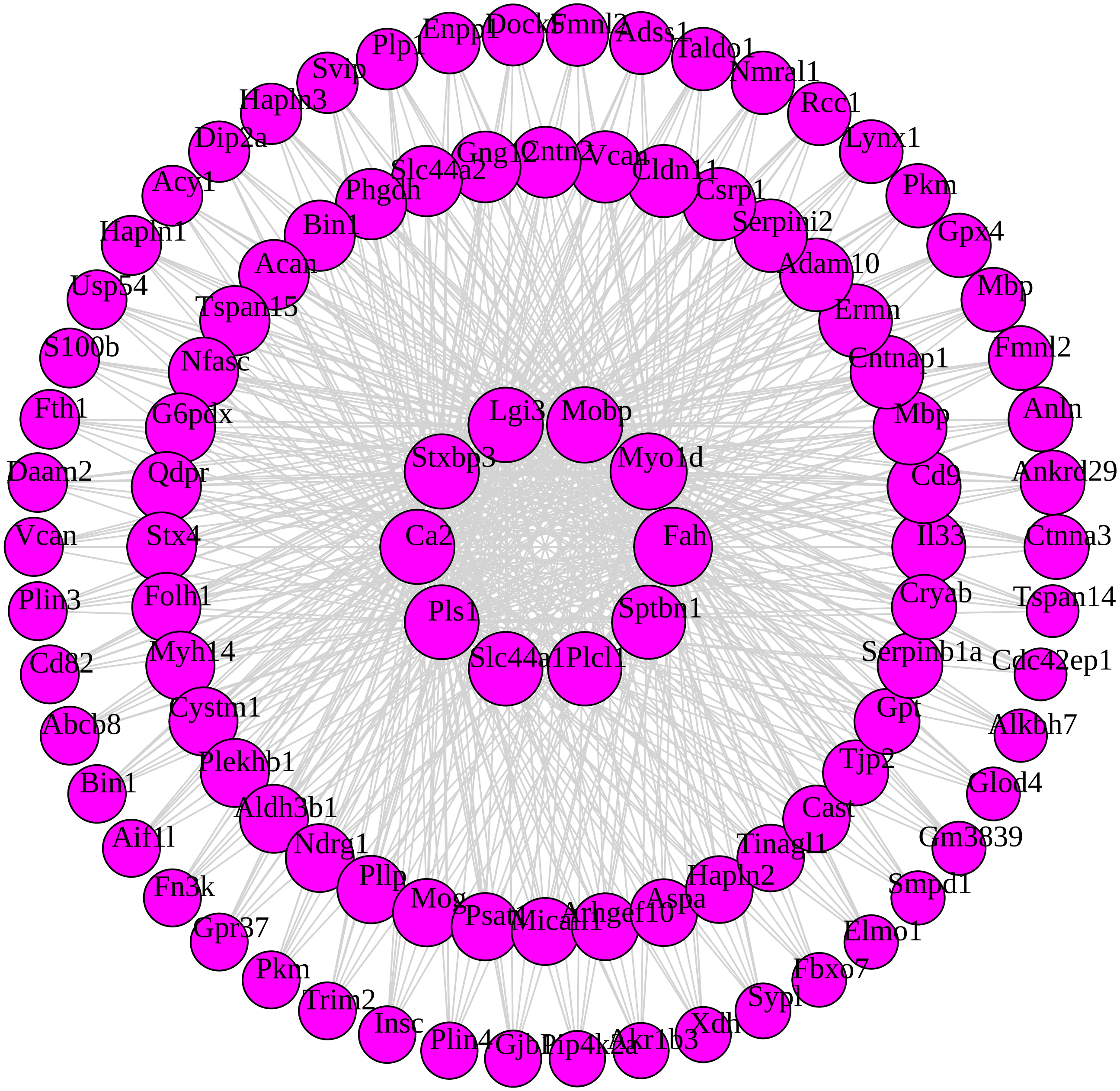

**M27 white module**

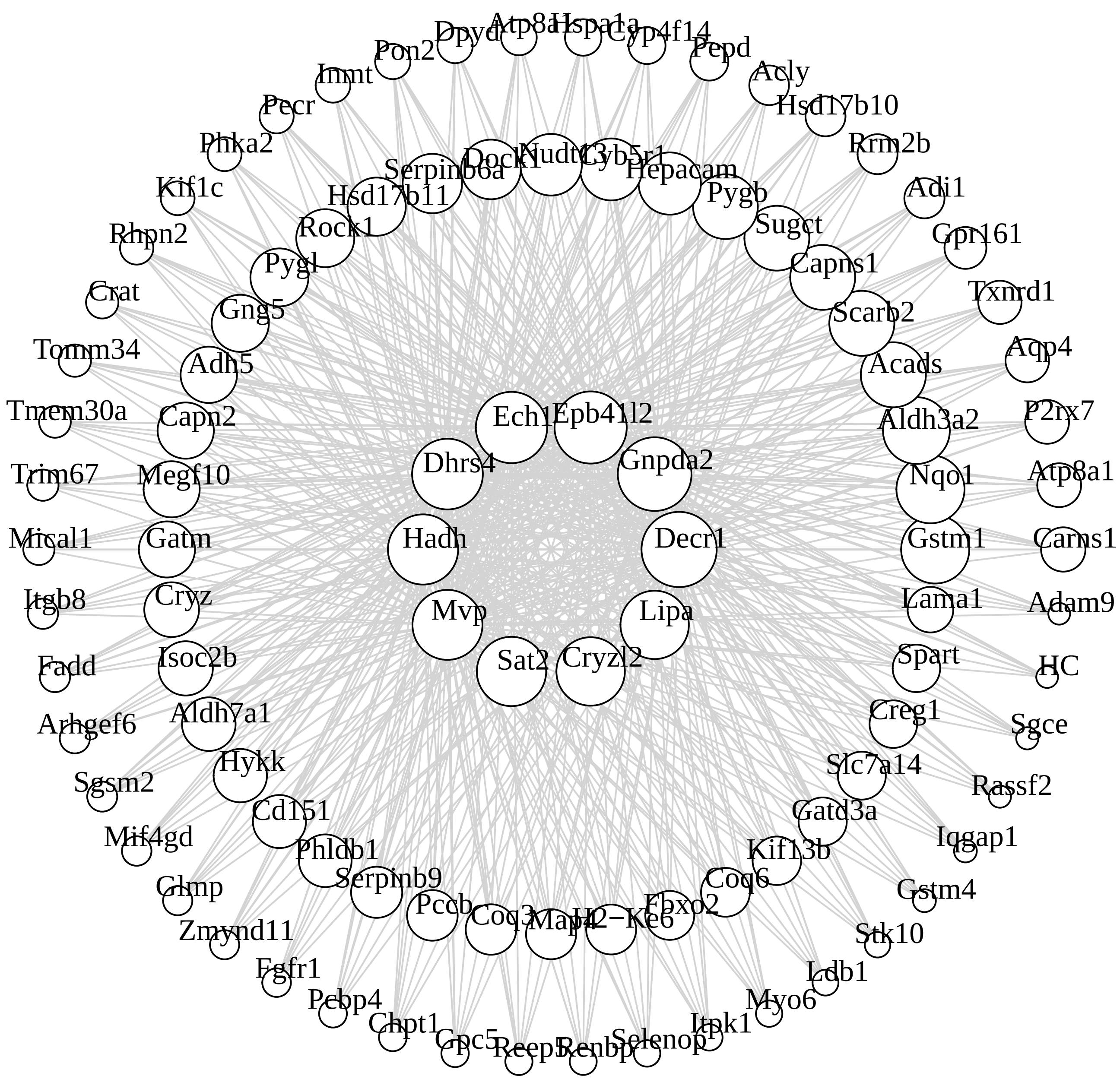

**M31 paleturquoise module**

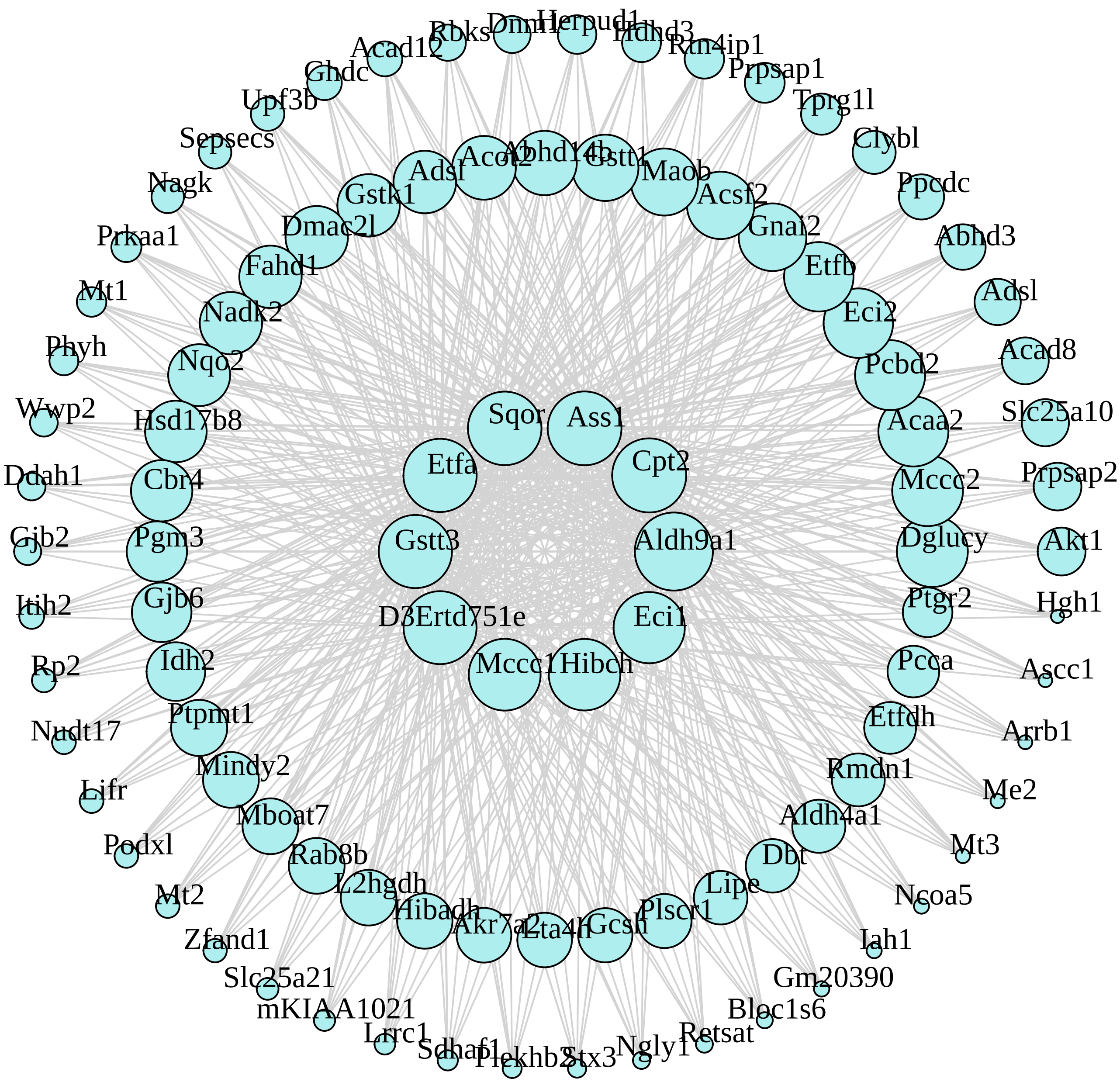

**M28 skyblue module**

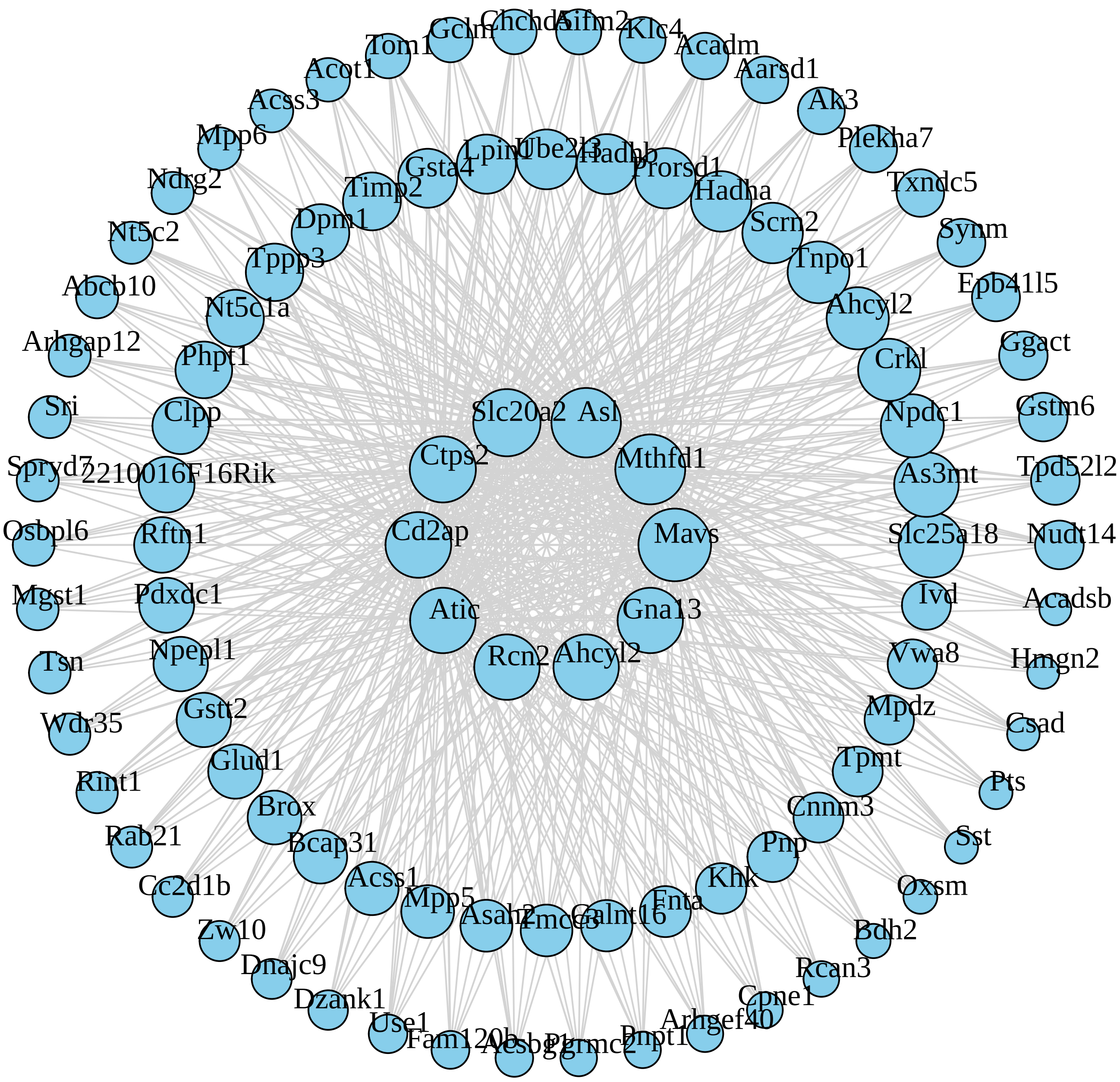

**M38 plum1 module**

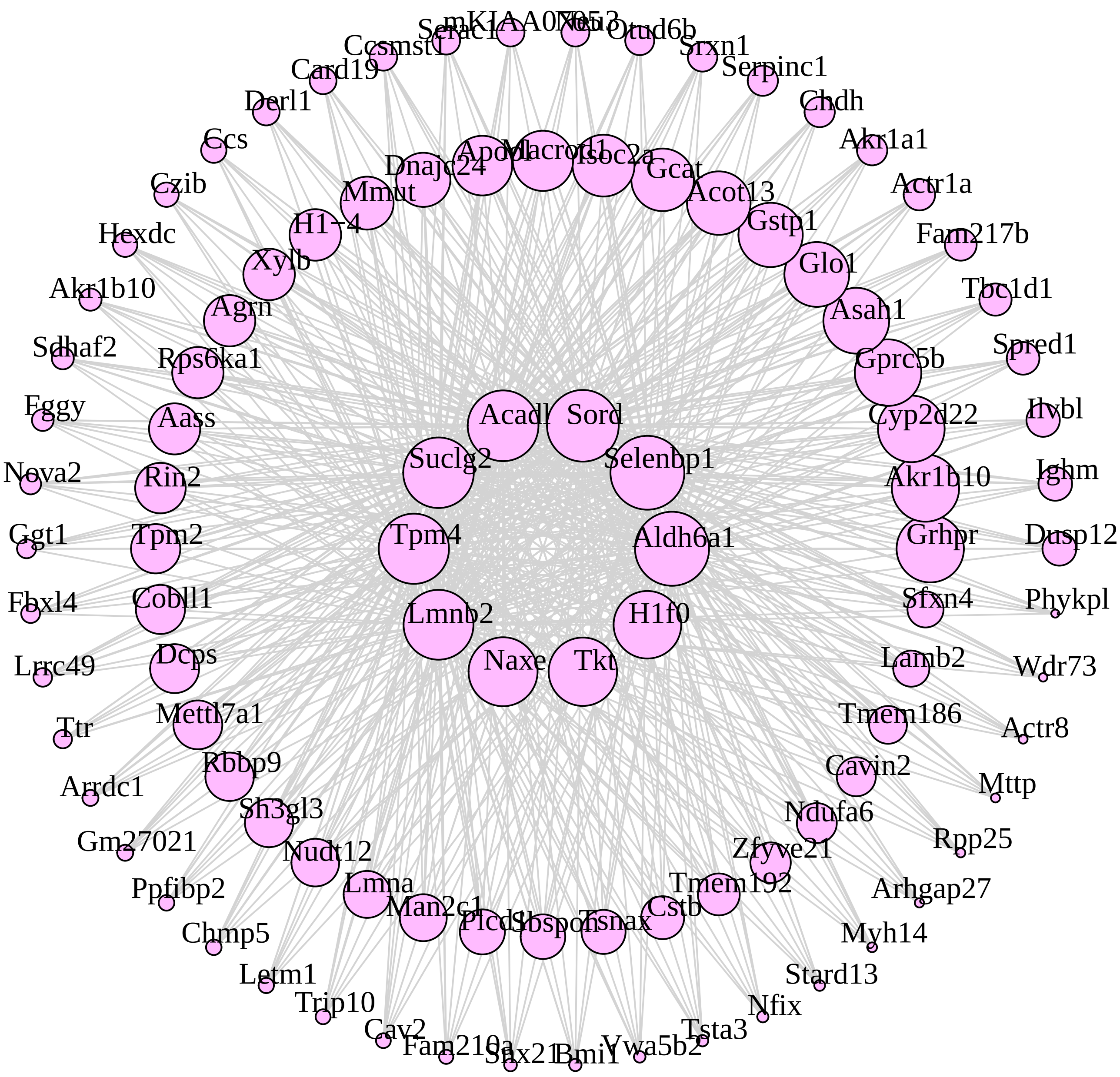

**M44 floralwhite module**

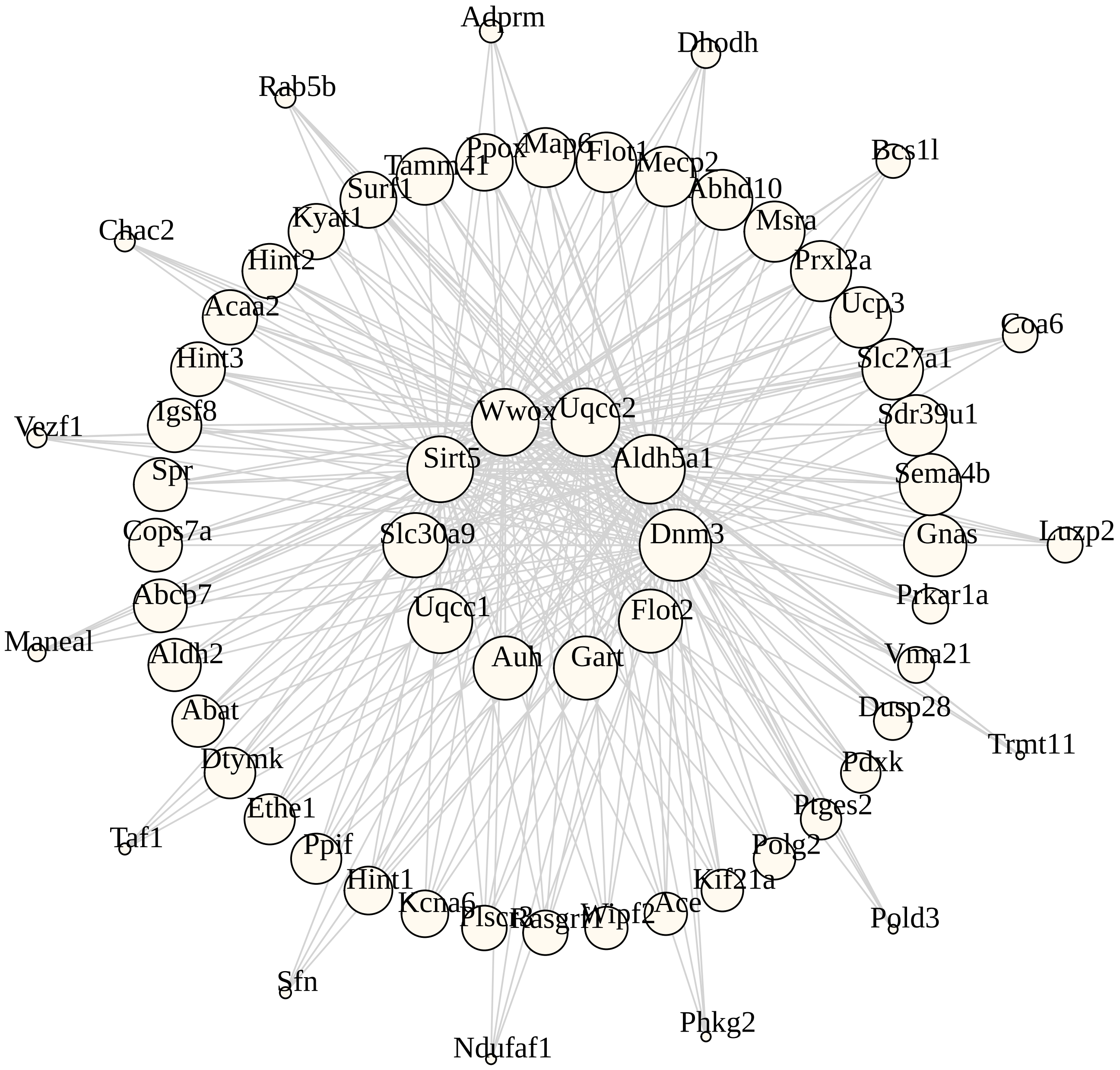

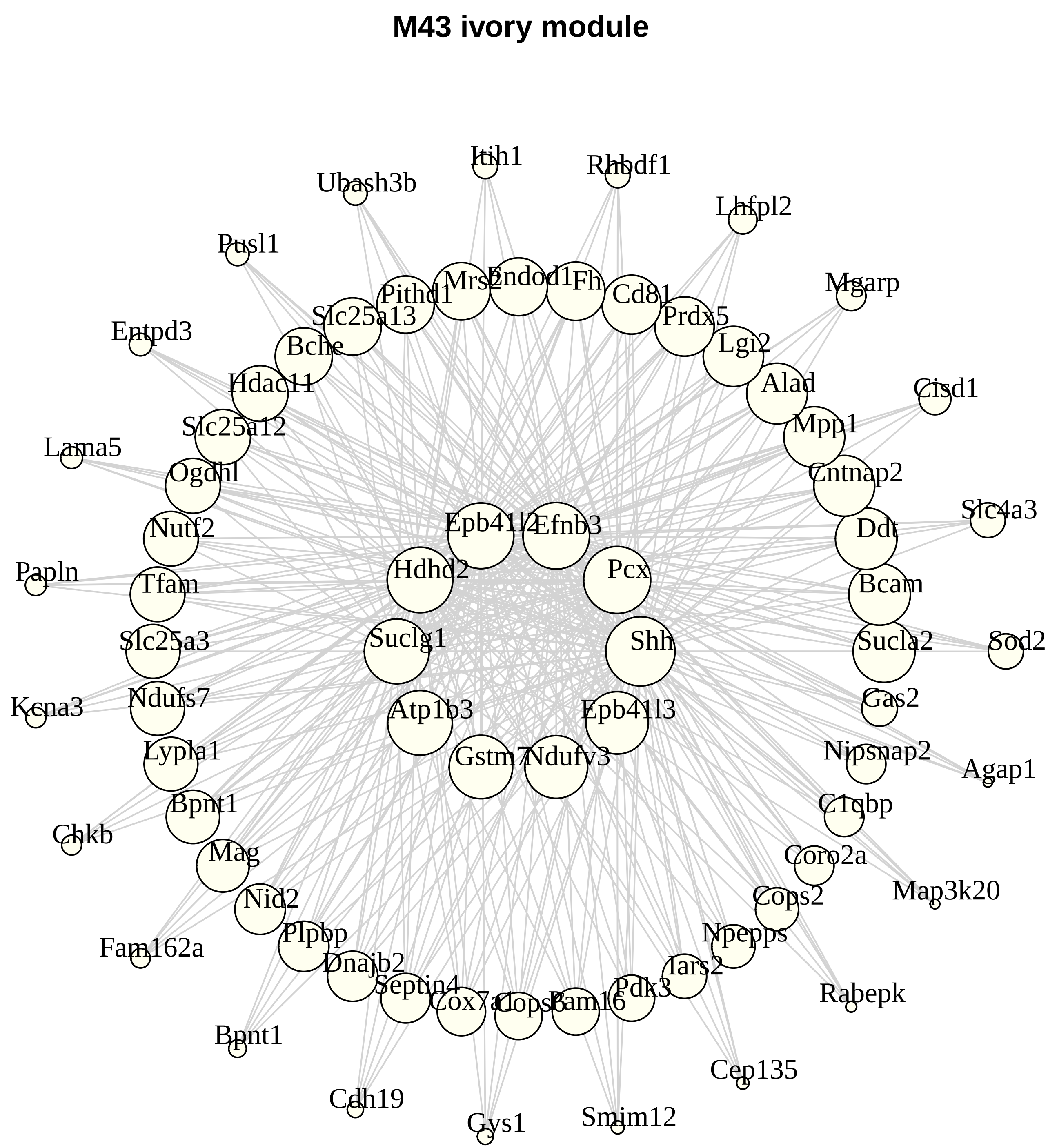
M43 ivory module
Itih1
Rhbdf1
Ubash3b
Lhfpl2
Pusl1
Mgarp
Endod1
Mrs2
Fh
Pithd1
Cd81
Slc25a13
Prdx5
Entpd3
Bche
Lgi2
Alad
Hdac11
Cisd1
Mpp1
Slc25a12
Lama5
Cntnap2
Ogdhl
Slc4a3
Epb41l2
Efnb3
Nutf2
Ddt
Papln
Hdhd2
Pcx
Tfam
Bcam
Shh
Suclg1
Sod2
Slc25a3
Sucla2
Ndufs7
Gas2
Kcna3
Atp1b3
Epb41l3
Lypla1
Nipsnap2
Gstm7
Ndufv3
Agap1
Bpnt1
C1qbp
Chkb
Mag
Coro2a
Map3k20
Nid2
Cops2
Plpbp
Npepps
Fam162a
Dnajb2
Iars2
Septin4
Pdk3
Rabepk
Cox7a1
Cops6
Pam16
Bpnt1
Cep135
Cdh19
Gys1
Smim12

**M42 lightcyan1 module**

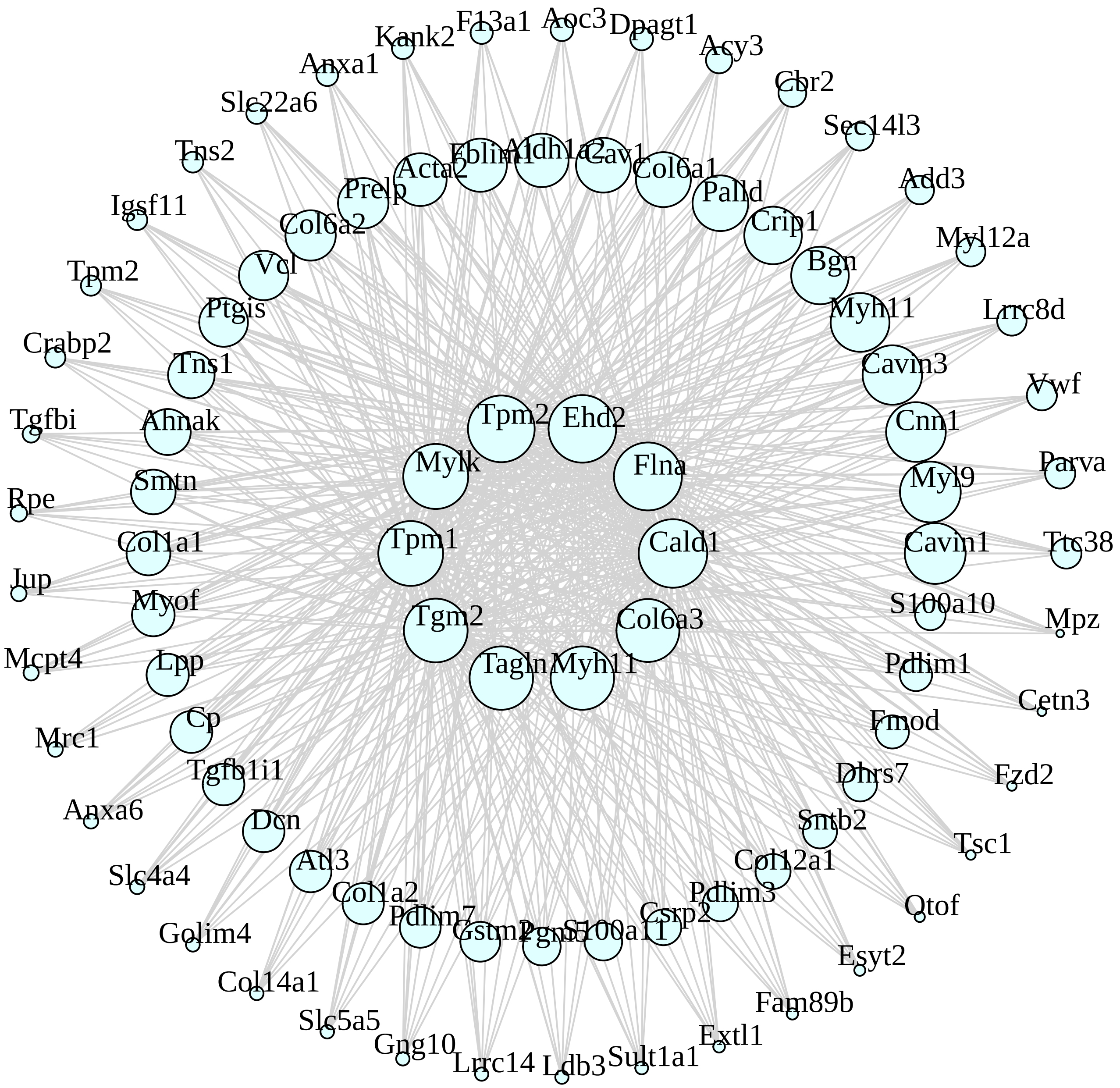

**M40 mediumpurple3 module**

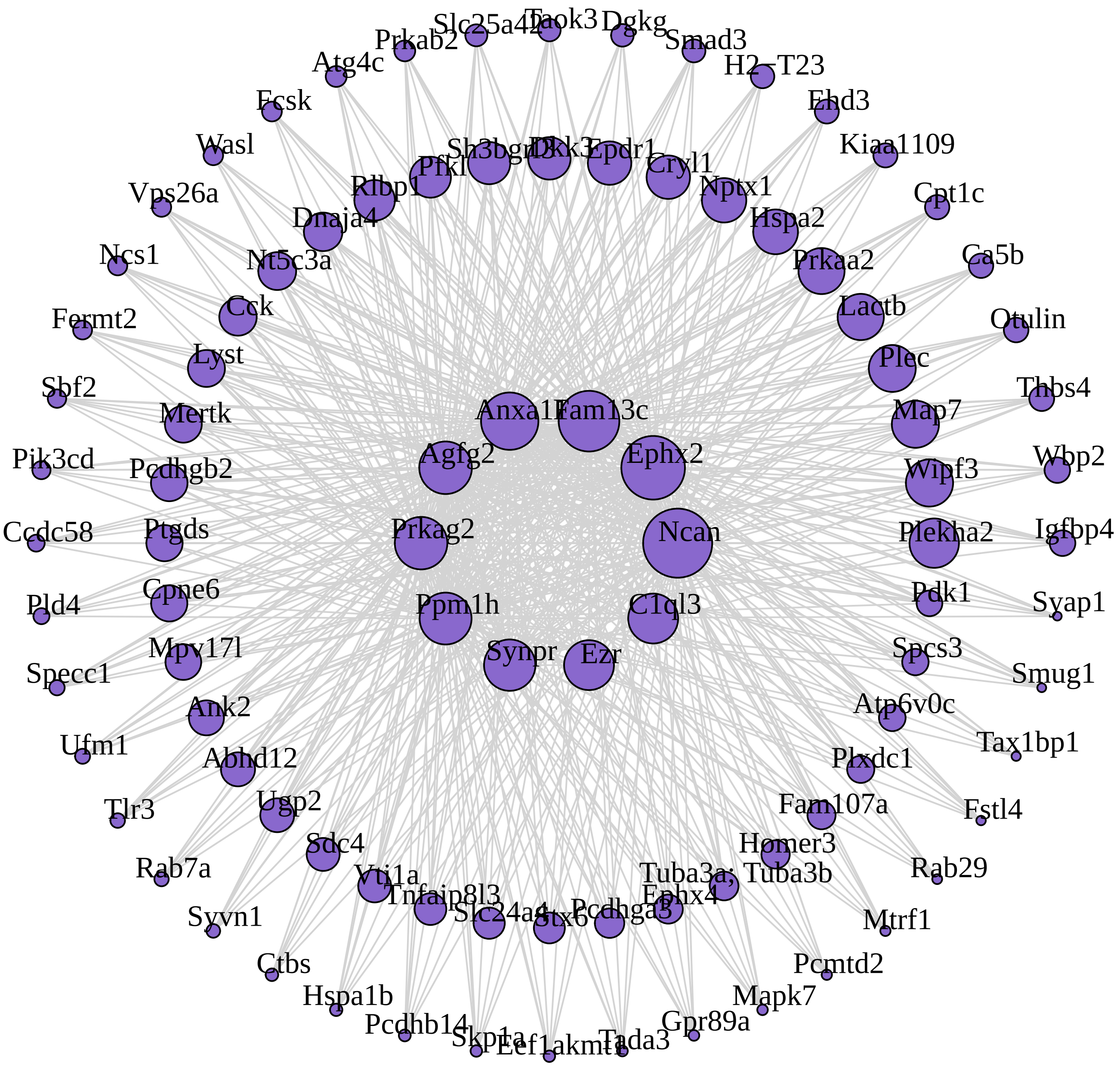

**M12 tan module**

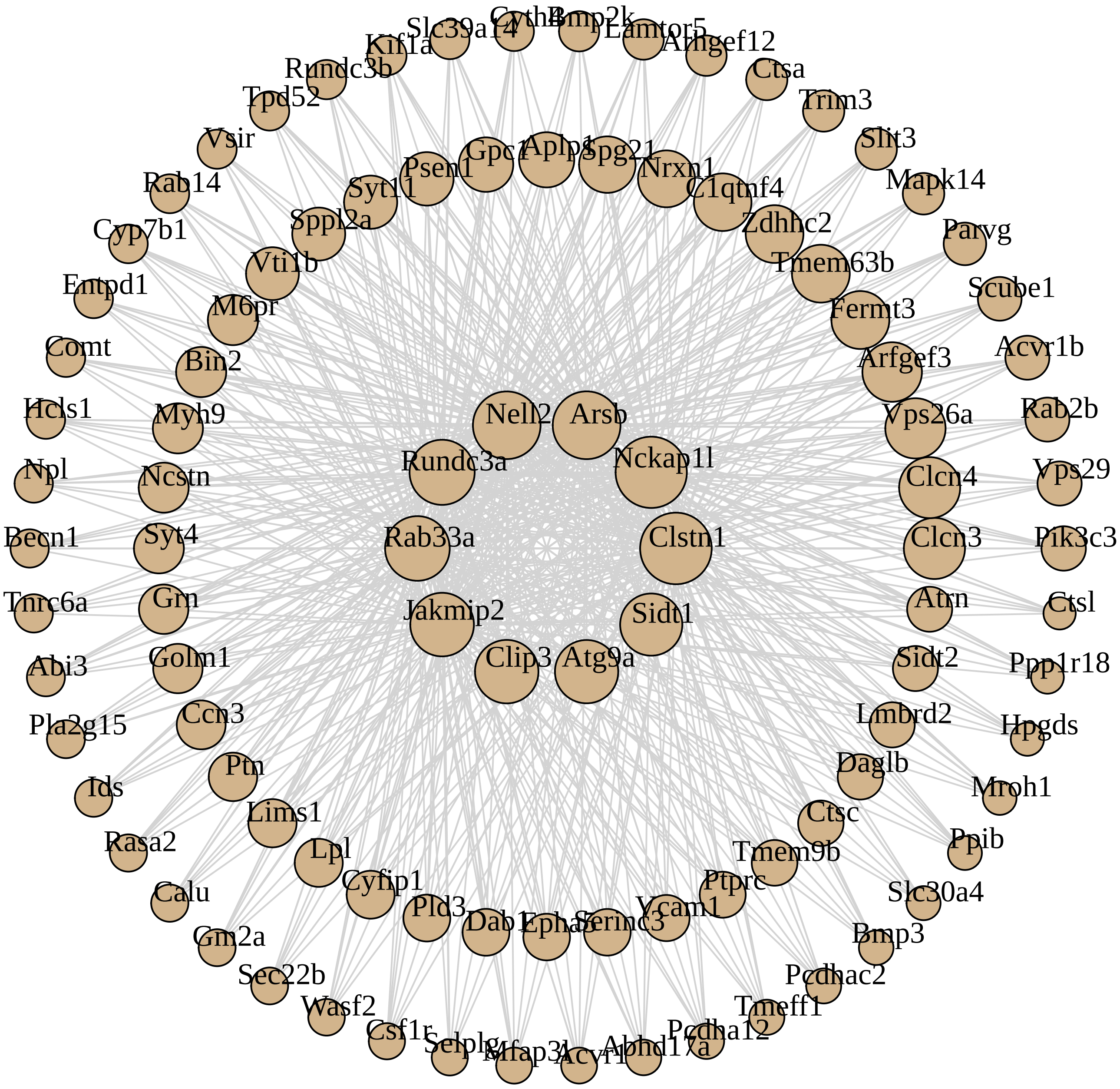

**M18 lightgreen module**

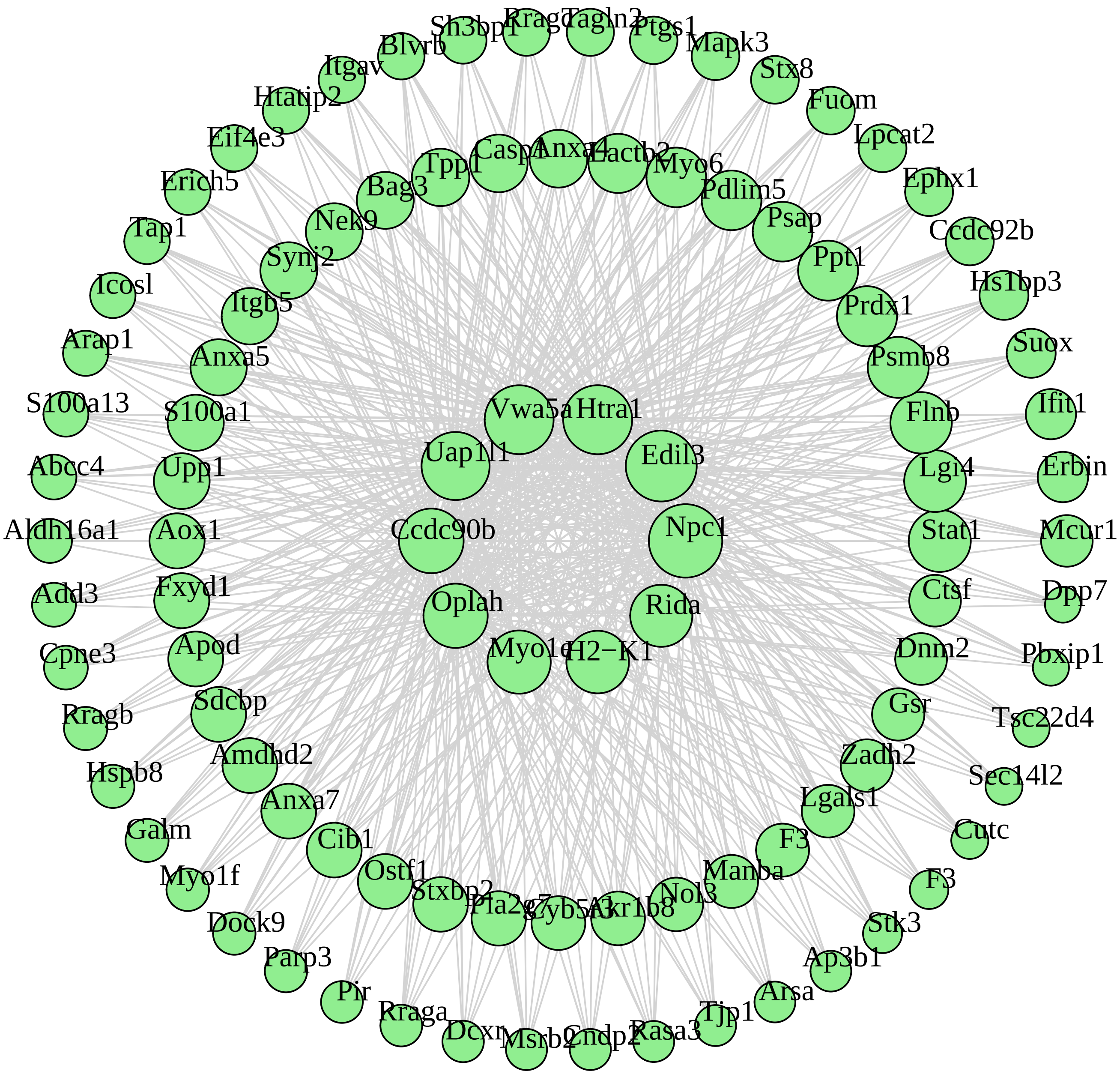

#### M50 thistle2 module

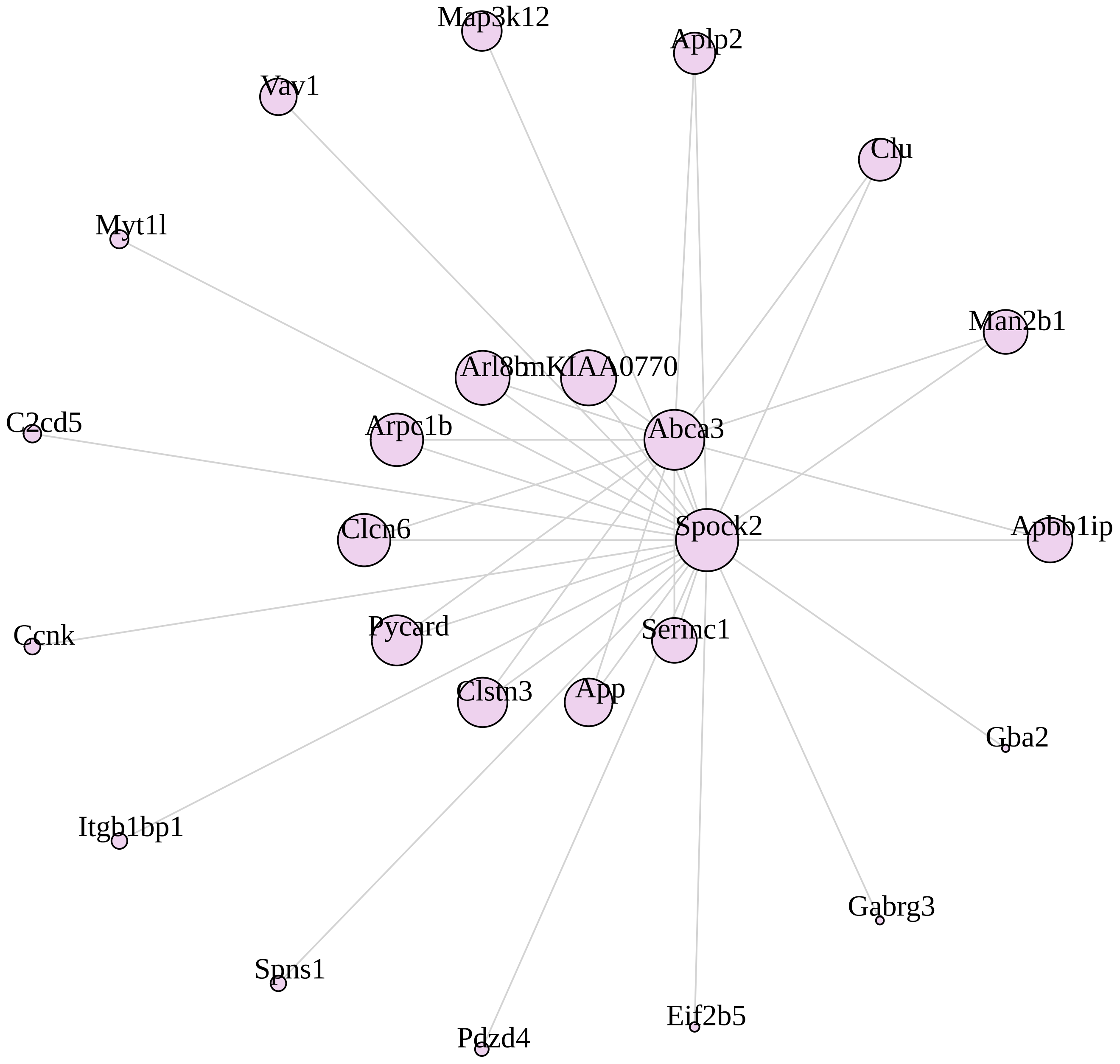

M20 royalblue module

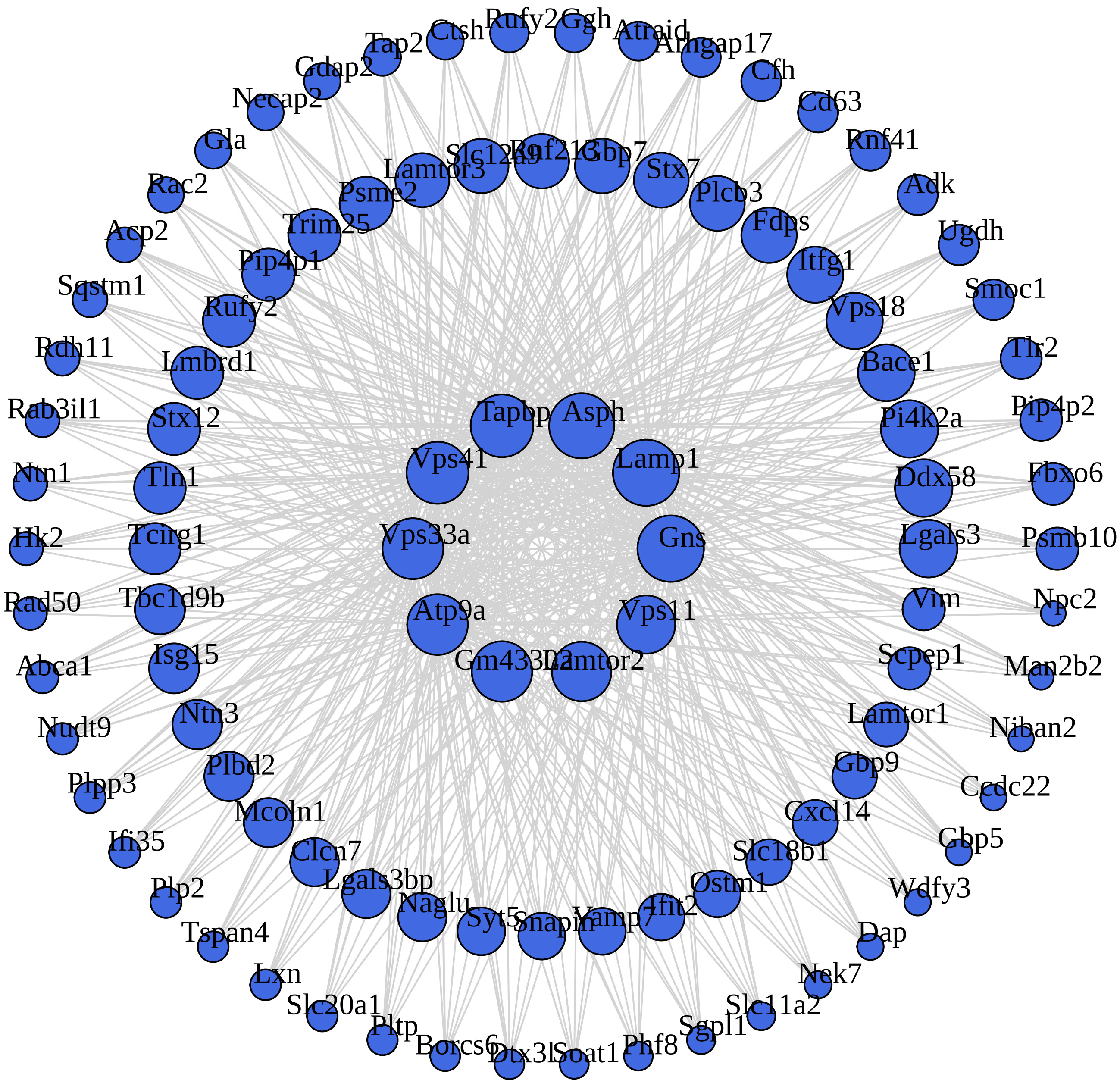

### M46 brown4 module

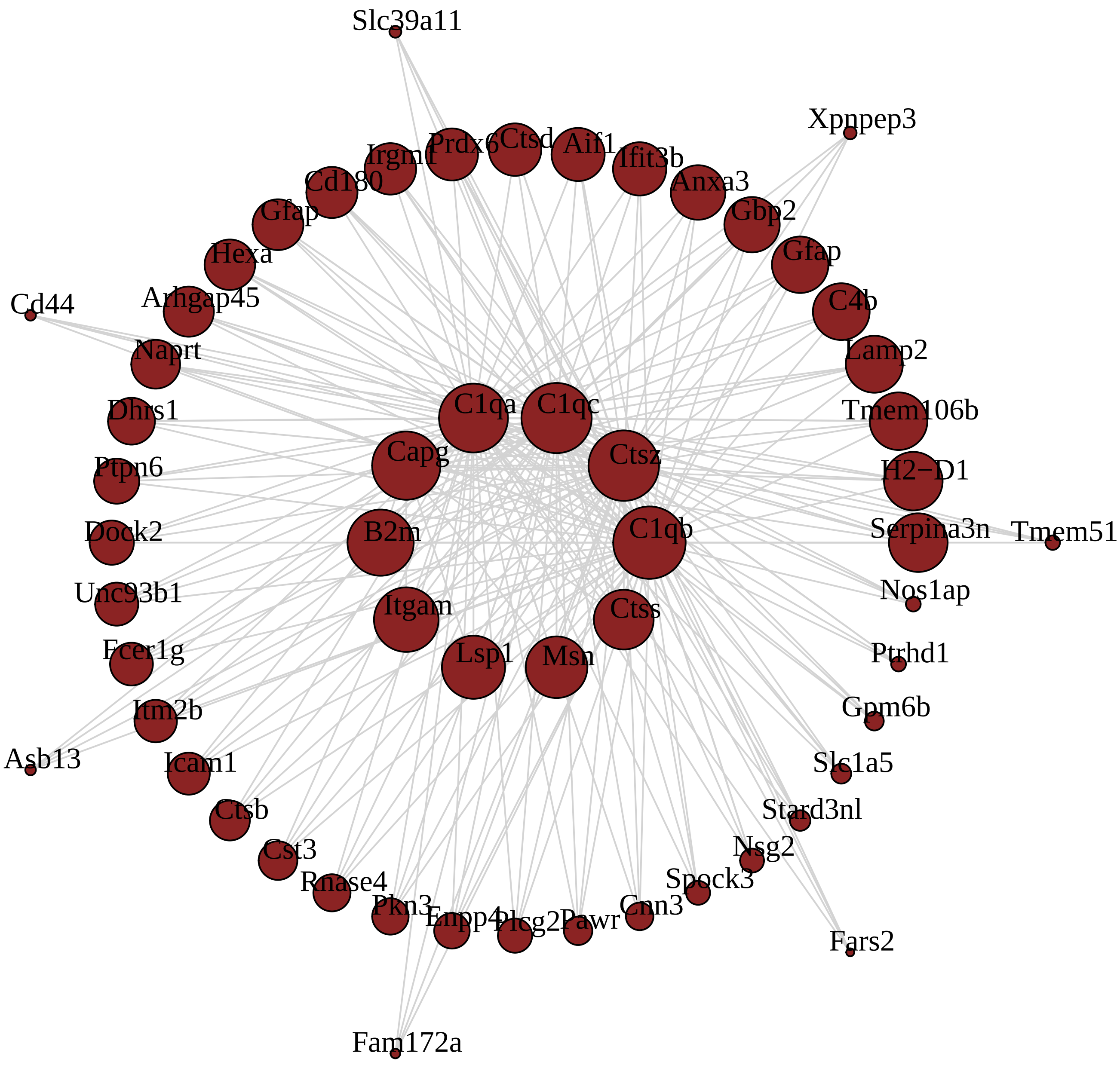

#### M51 thistle1 module

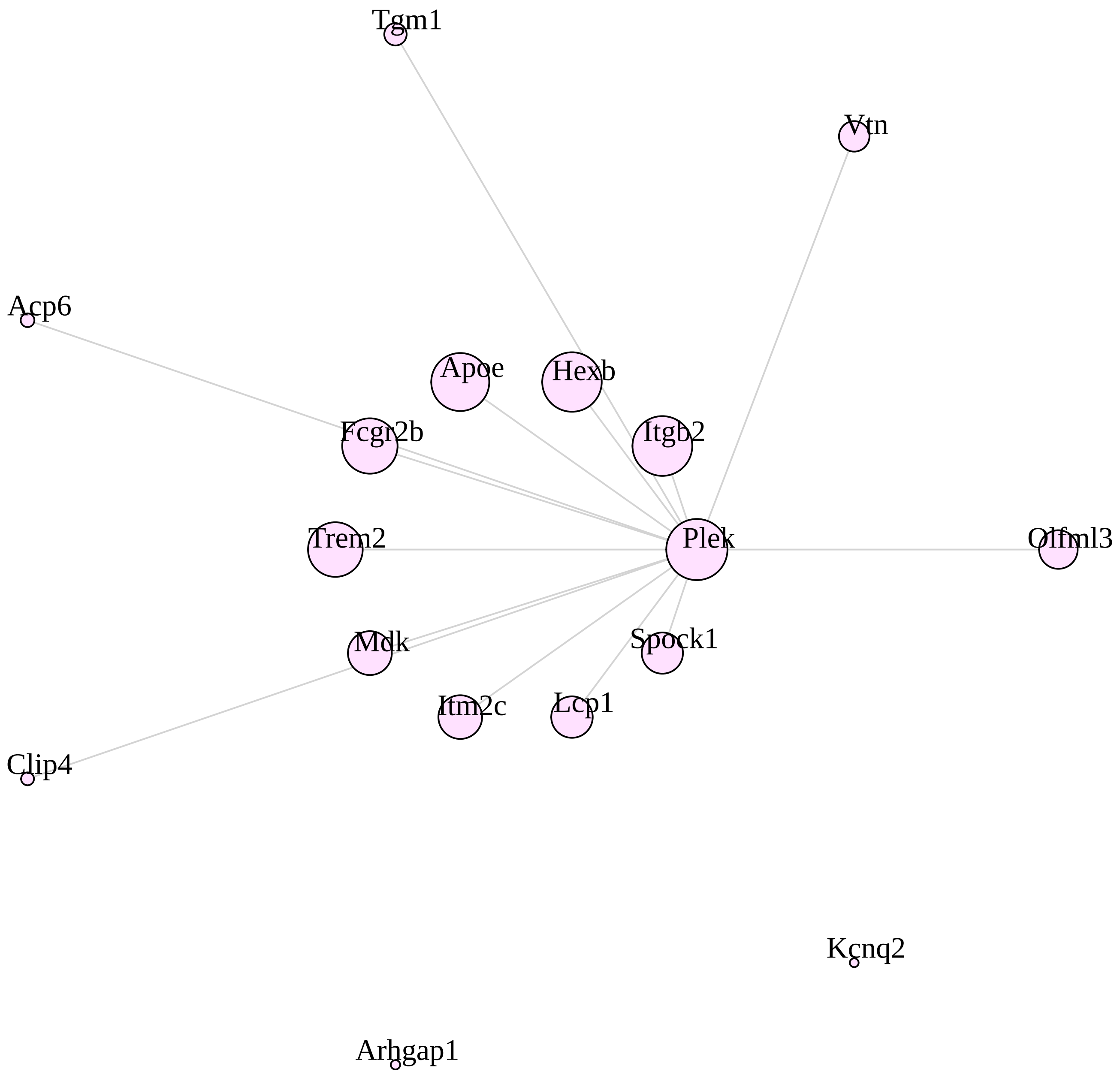

### M17 grey60 module

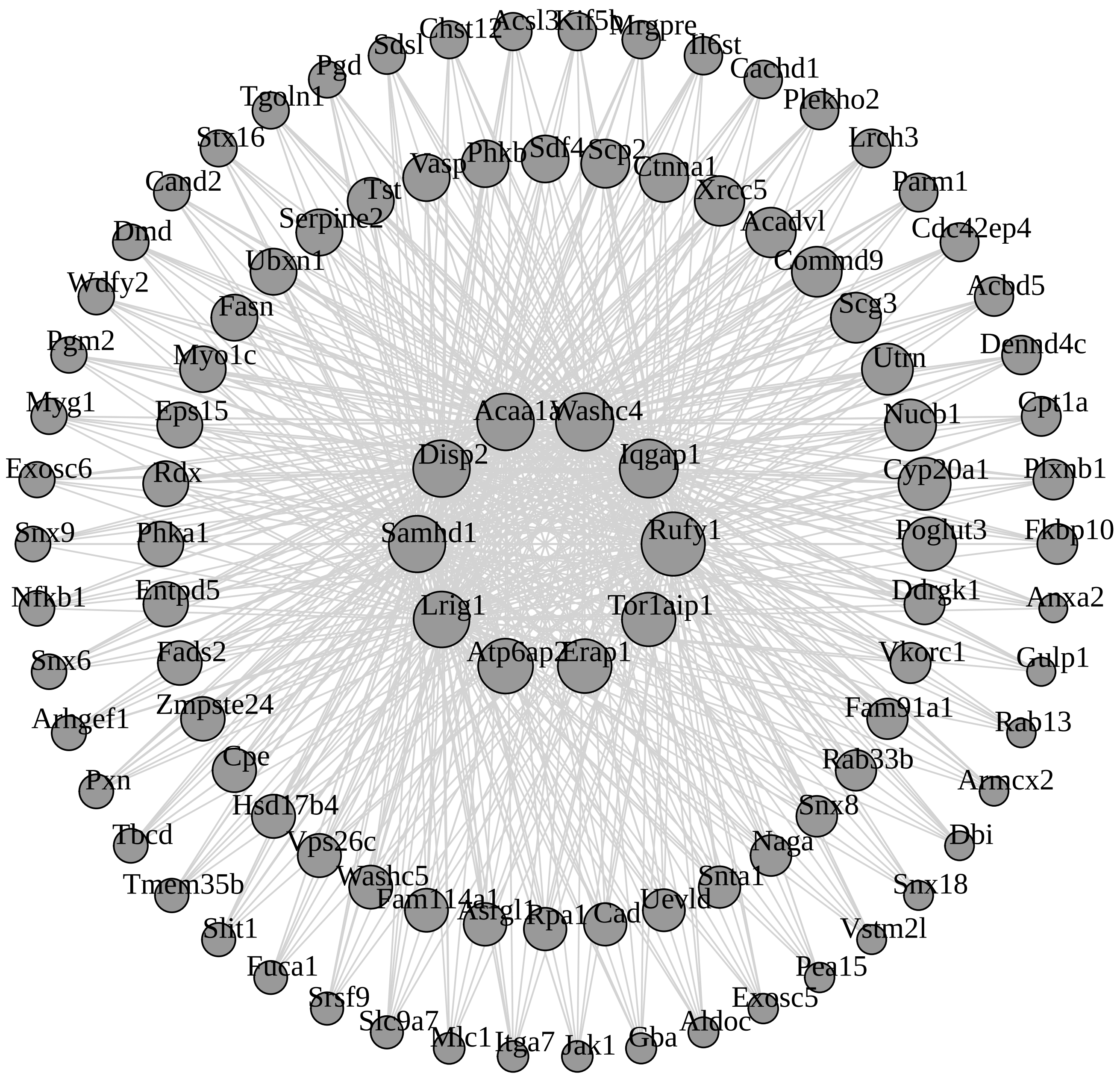

#### M36 yellowgreen module

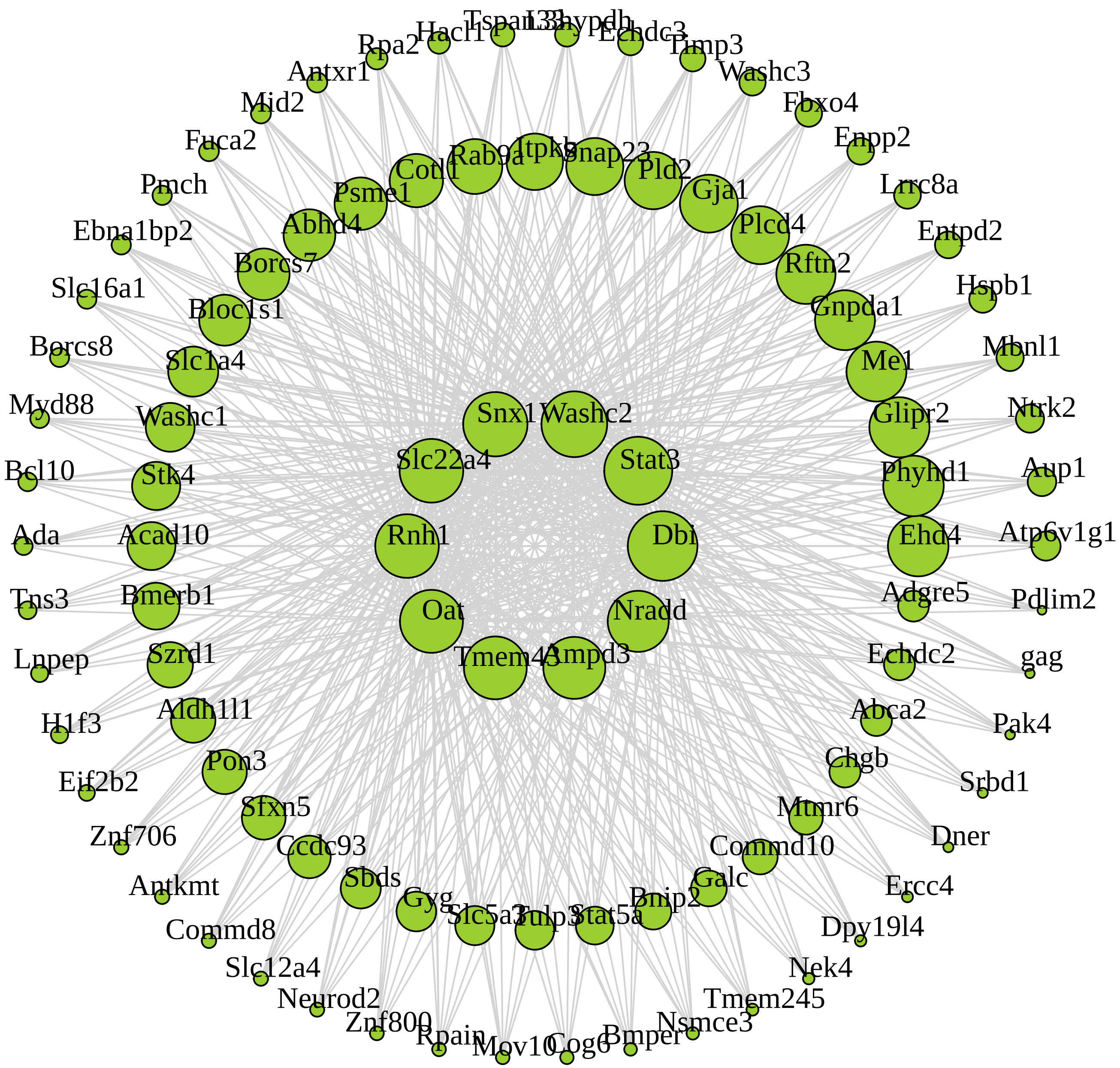

**M26 darkorange module**

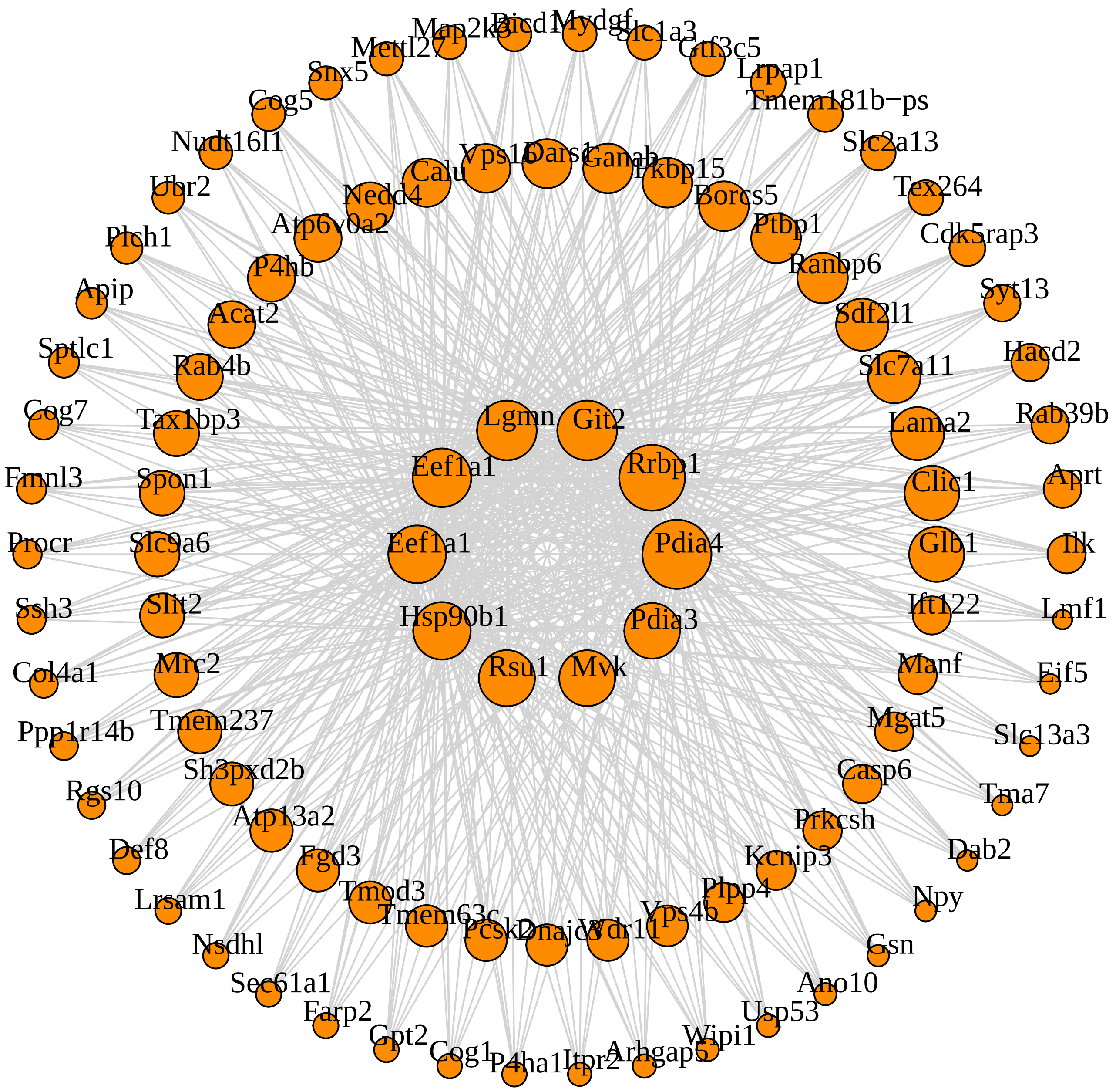

**M39 orangered4 module**

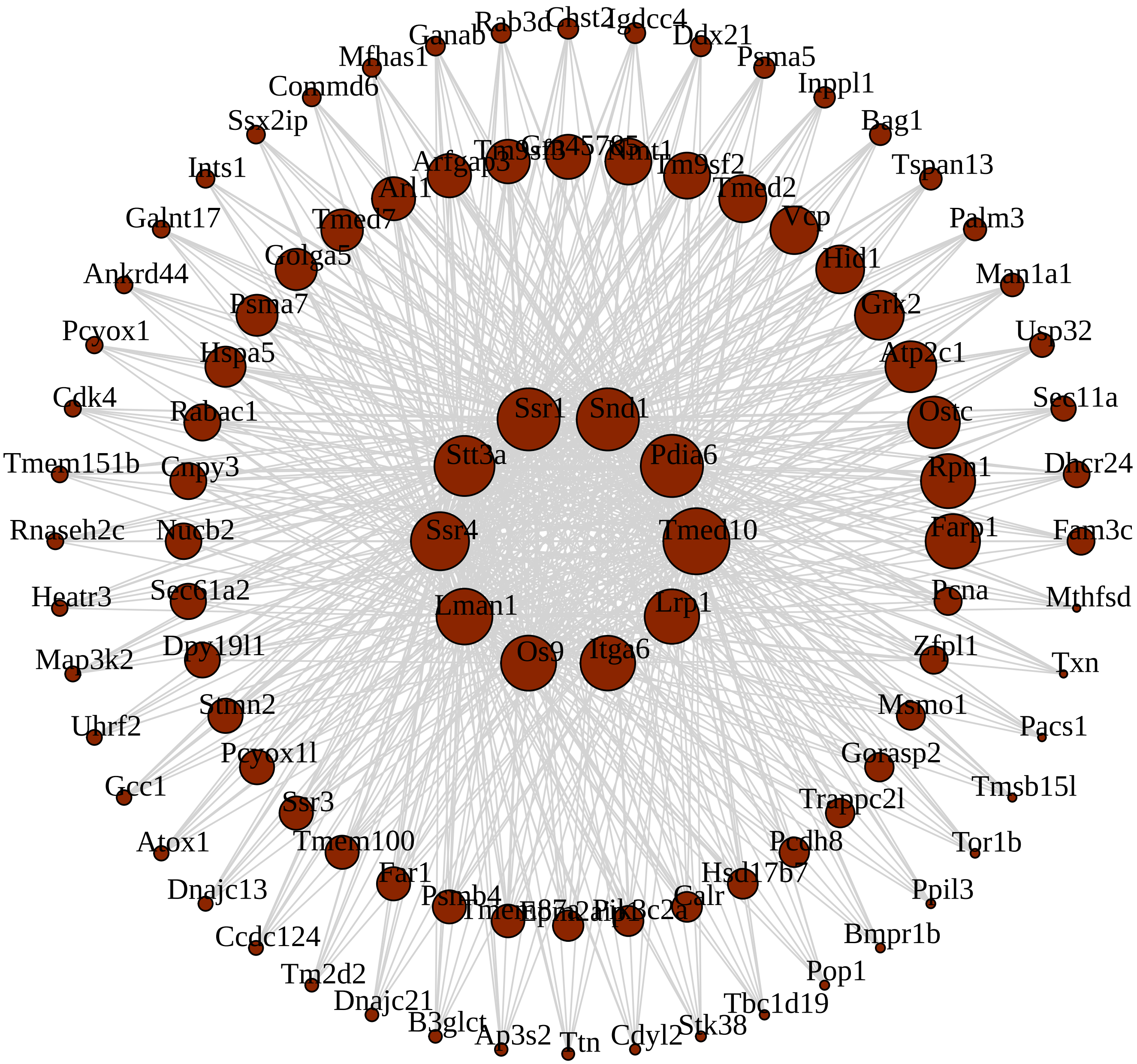
