## Supplemental Data 2 for "Network analysis of mouse brain proteomes resolves molecular and cellular endophenotypes of aging and Aβ pathology"

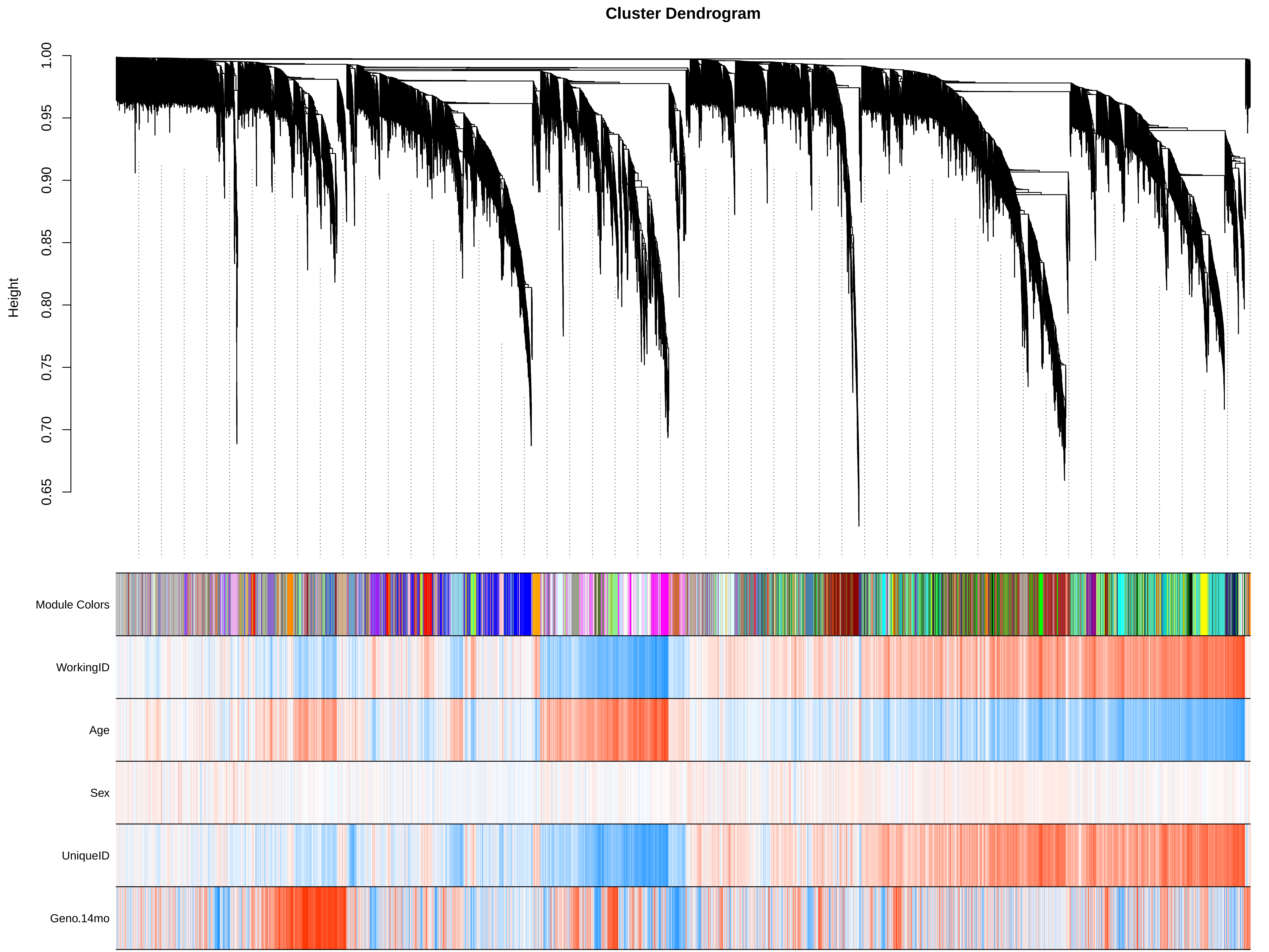
Cluster Dendrogram
Height
1.00
0.95
0.90
0.85
0.80
0.75
0.70
0.65
Module Colors
WorkingID
Age
Sex
UniqueID
Geno.14mo

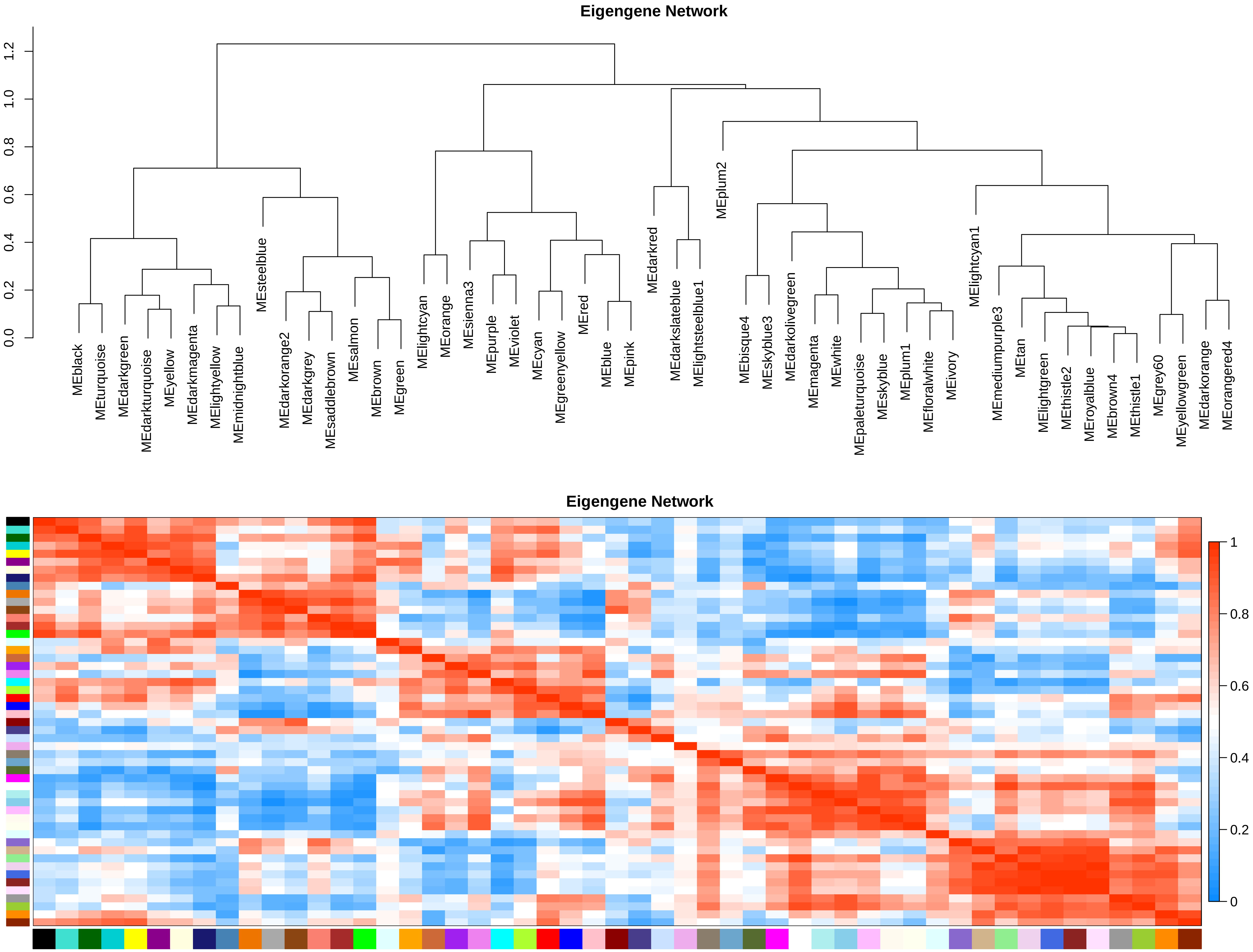

Eigengene Network
0.0
0.2
0.4
0.6
0.8
1.0
1.2
MEblack
MEturquoise
MEdarkgreen
MEdarkturquoise
MEyellow
MEdarkmagenta
MElightyellow
MEmidnightblue
MEsteelblue
MEdarkorange2
MEdarkgrey
MEsaddlebrown
MEsalmon
MEbrown
MEgreen
MElightcyan
MEorange
MEsienna3
MEpurple
MEviolet
MEcyan
MEgreenyellow
MEred
MEblue
MEpink
MEdarkred
MEdarkslateblue
MElightsteelblue1
MEplum2
MEbisque4
MEskyblue3
MEdarkolivegreen
MEmagenta
MEwhite
MEpaleturquoise
MEskyblue
MEplum1
MEfloralwhite
MEivory
MElightcyan1
MEmediumpurple3
MEtan
MElightgreen
MEthistle2
MEroyalblue
MEbrown4
MEthistle1
MEgrey60
MEyellowgreen
MEdarkorange
MEorangered4
Eigengene Network
1
0.8
0.6
0.4
0.2
0

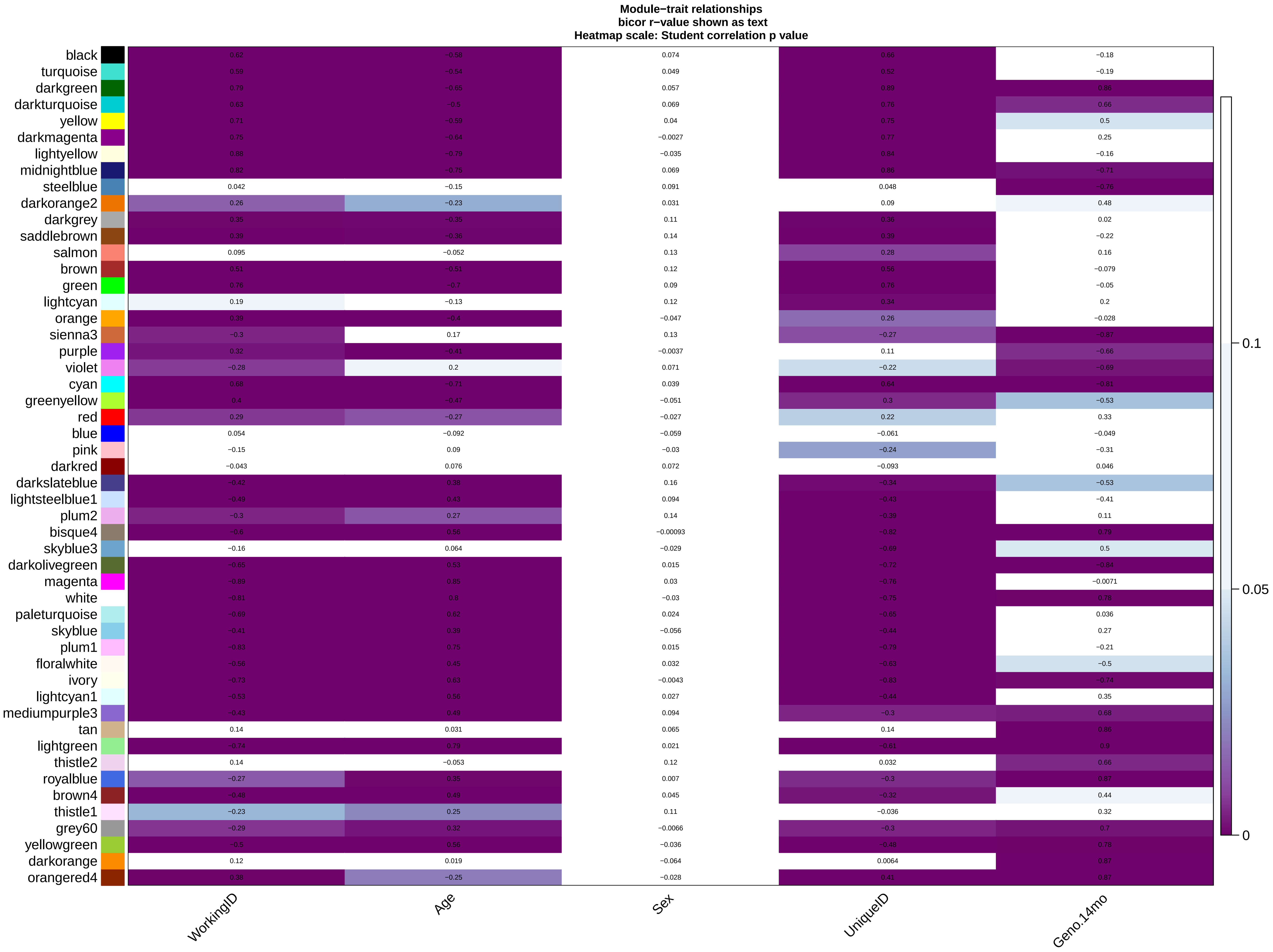
Module−trait relationships
bicor r−value shown as text
Heatmap scale: Student correlation p value
black
turquoise
darkgreen
darkturquoise
yellow
darkmagenta
lightyellow
midnightblue
steelblue
darkorange2
darkgrey
saddlebrown
salmon
brown
green
lightcyan
orange
sienna3
purple
violet
cyan
greenyellow
red
blue
pink
darkred
darkslateblue
lightsteelblue1
plum2
bisque4
skyblue3
darkolivegreen
magenta
white
paleturquoise
skyblue
plum1
floralwhite
ivory
lightcyan1
mediumpurple3
tan
lightgreen
thistle2
royalblue
brown4
thistle1
grey60
yellowgreen
darkorange
orangered4
WorkingID
Age
Sex
UniqueID
Geno.14mo
0.1
0.05
0

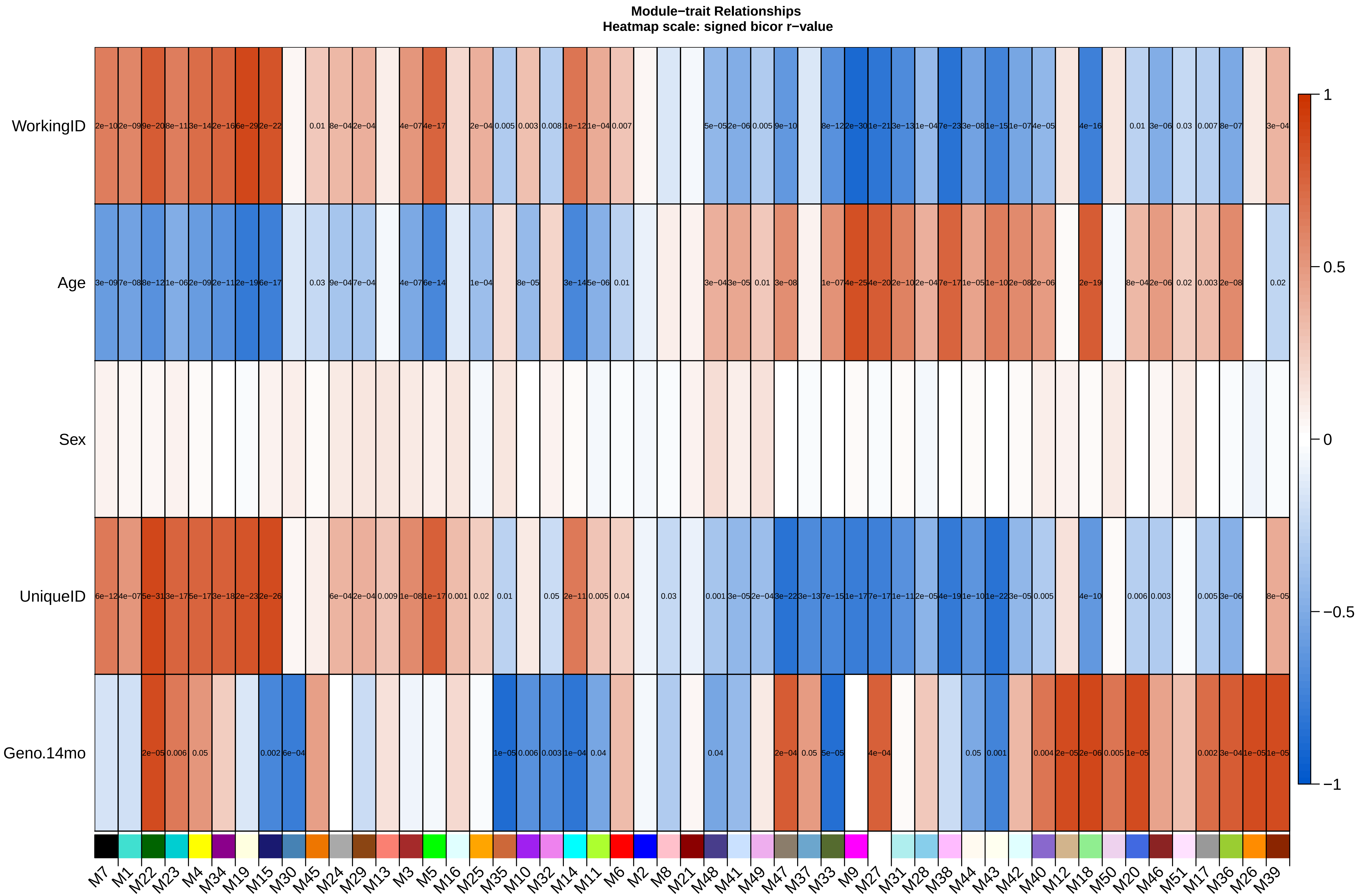
Module−trait Relationships
Heatmap scale: signed bicor r−value
WorkingID
Age
Sex
UniqueID
Geno.14mo
1
0.5
0
−0.5
−1
M7 M1 M22 M23 M4 M34 M19 M15 M30 M45 M24 M29 M13 M3 M5 M16 M25 M35 M10 M32 M14 M11 M6 M2 M8 M21 M48 M41 M49 M47 M37 M33 M9 M27 M31 M28 M38 M44 M43 M42 M40 M12 M18 M50 M20 M46 M51 M17 M36 M26 M39

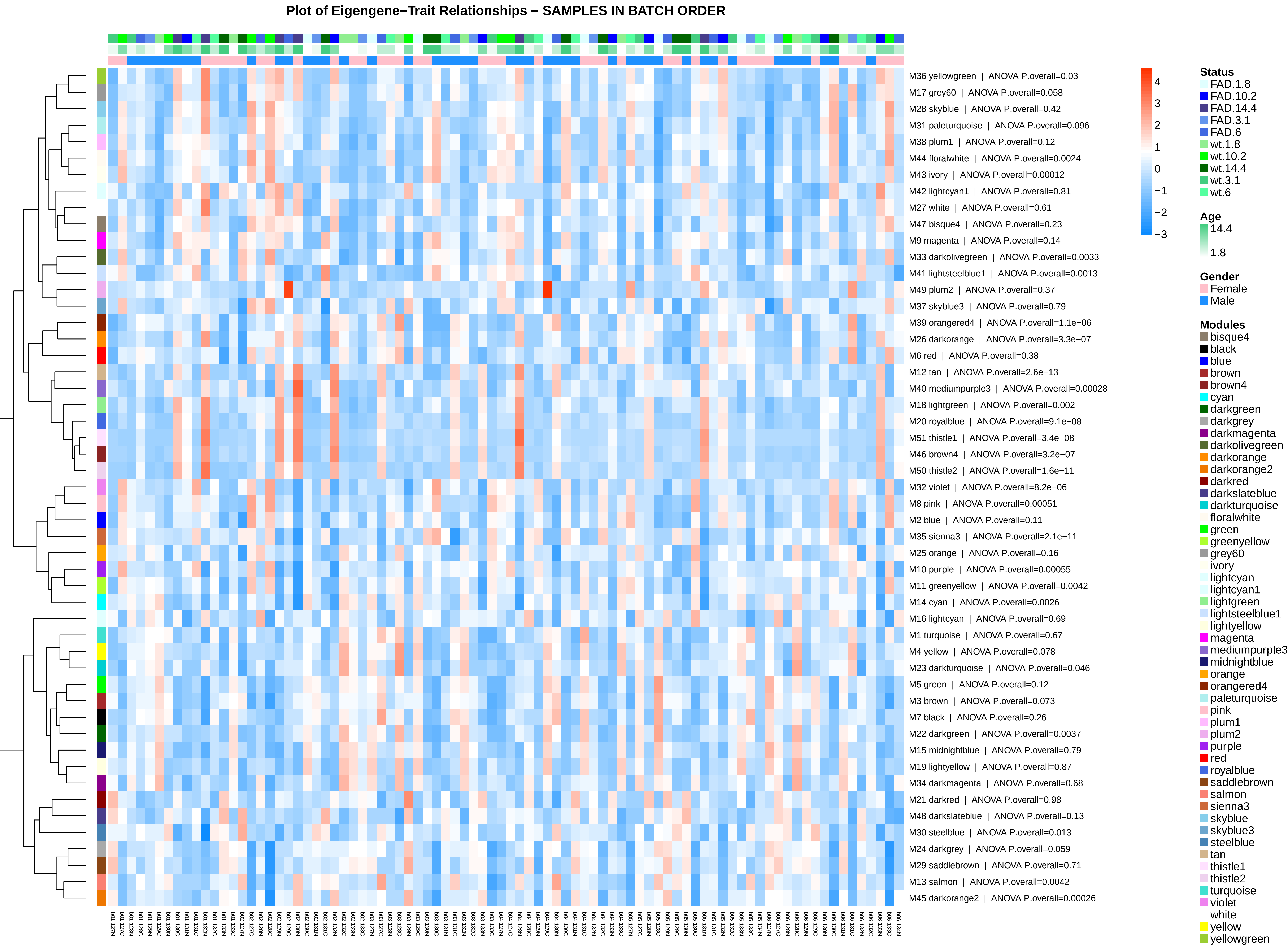
Plot of Eigengene−Trait Relationships − SAMPLES IN BATCH ORDER
M36 yellowgreen | ANOVA P.overall=0.03
M17 grey60 | ANOVA P.overall=0.058
M28 skyblue | ANOVA P.overall=0.42
M31 paleturquoise | ANOVA P.overall=0.096
M38 plum1 | ANOVA P.overall=0.12
M44 floralwhite | ANOVA P.overall=0.0024
M43 ivory | ANOVA P.overall=0.00012
M42 lightcyan1 | ANOVA P.overall=0.81
M27 white | ANOVA P.overall=0.61
M47 bisque4 | ANOVA P.overall=0.23
M9 magenta | ANOVA P.overall=0.14
M33 darkolivegreen | ANOVA P.overall=0.0033
M41 lightsteelblue1 | ANOVA P.overall=0.0013
M49 plum2 | ANOVA P.overall=0.37
M37 skyblue3 | ANOVA P.overall=0.79
M39 orangered4 | ANOVA P.overall=1.1e−06
M26 darkorange | ANOVA P.overall=3.3e−07
M6 red | ANOVA P.overall=0.38
M12 tan | ANOVA P.overall=2.6e−13
M40 mediumpurple3 | ANOVA P.overall=0.00028
M18 lightgreen | ANOVA P.overall=0.002
M20 royalblue | ANOVA P.overall=9.1e−08
M51 thistle1 | ANOVA P.overall=3.4e−08
M46 brown4 | ANOVA P.overall=3.2e−07
M50 thistle2 | ANOVA P.overall=1.6e−11
M32 violet | ANOVA P.overall=8.2e−06
M8 pink | ANOVA P.overall=0.00051
M2 blue | ANOVA P.overall=0.11
M35 sienna3 | ANOVA P.overall=2.1e−11
M25 orange | ANOVA P.overall=0.16
M10 purple | ANOVA P.overall=0.00055
M11 greenyellow | ANOVA P.overall=0.0042
M14 cyan | ANOVA P.overall=0.0026
M16 lightcyan | ANOVA P.overall=0.69
M1 turquoise | ANOVA P.overall=0.67
M4 yellow | ANOVA P.overall=0.078
M23 darkturquoise | ANOVA P.overall=0.046
M5 green | ANOVA P.overall=0.12
M3 brown | ANOVA P.overall=0.073
M7 black | ANOVA P.overall=0.26
M22 darkgreen | ANOVA P.overall=0.0037
M15 midnightblue | ANOVA P.overall=0.79
M19 lightyellow | ANOVA P.overall=0.87
M34 darkmagenta | ANOVA P.overall=0.68
M21 darkred | ANOVA P.overall=0.98
M48 darkslateblue | ANOVA P.overall=0.13
M30 steelblue | ANOVA P.overall=0.013
M24 darkgrey | ANOVA P.overall=0.059
M29 saddlebrown | ANOVA P.overall=0.71
M13 salmon | ANOVA P.overall=0.0042
M45 darkorange2 | ANOVA P.overall=0.00026
4
3
2
1
0
−1
−2
−3
Status
FAD.1.8
FAD.10.2
FAD.14.4
FAD.3.1
FAD.6
wt.1.8
wt.10.2
wt.14.4
wt.3.1
wt.6
Age
14.4
1.8
Gender
Female
Male
Modules
bisque4
black
blue
brown
brown4
cyan
darkgreen
darkgrey
darkmagenta
darkolivegreen
darkorange
darkorange2
darkred
darkslateblue
darkturquoise
floralwhite
green
greenyellow
grey60
ivory
lightcyan
lightcyan1
lightgreen
lightsteelblue1
lightyellow
magenta
mediumpurple3
midnightblue
orange
orangered4
paleturquoise
pink
plum1
plum2
purple
red
royalblue
saddlebrown
salmon
sienna3
skyblue
skyblue3
steelblue
tan
thistle1
thistle2
turquoise
violet
white
yellow
yellowgreen

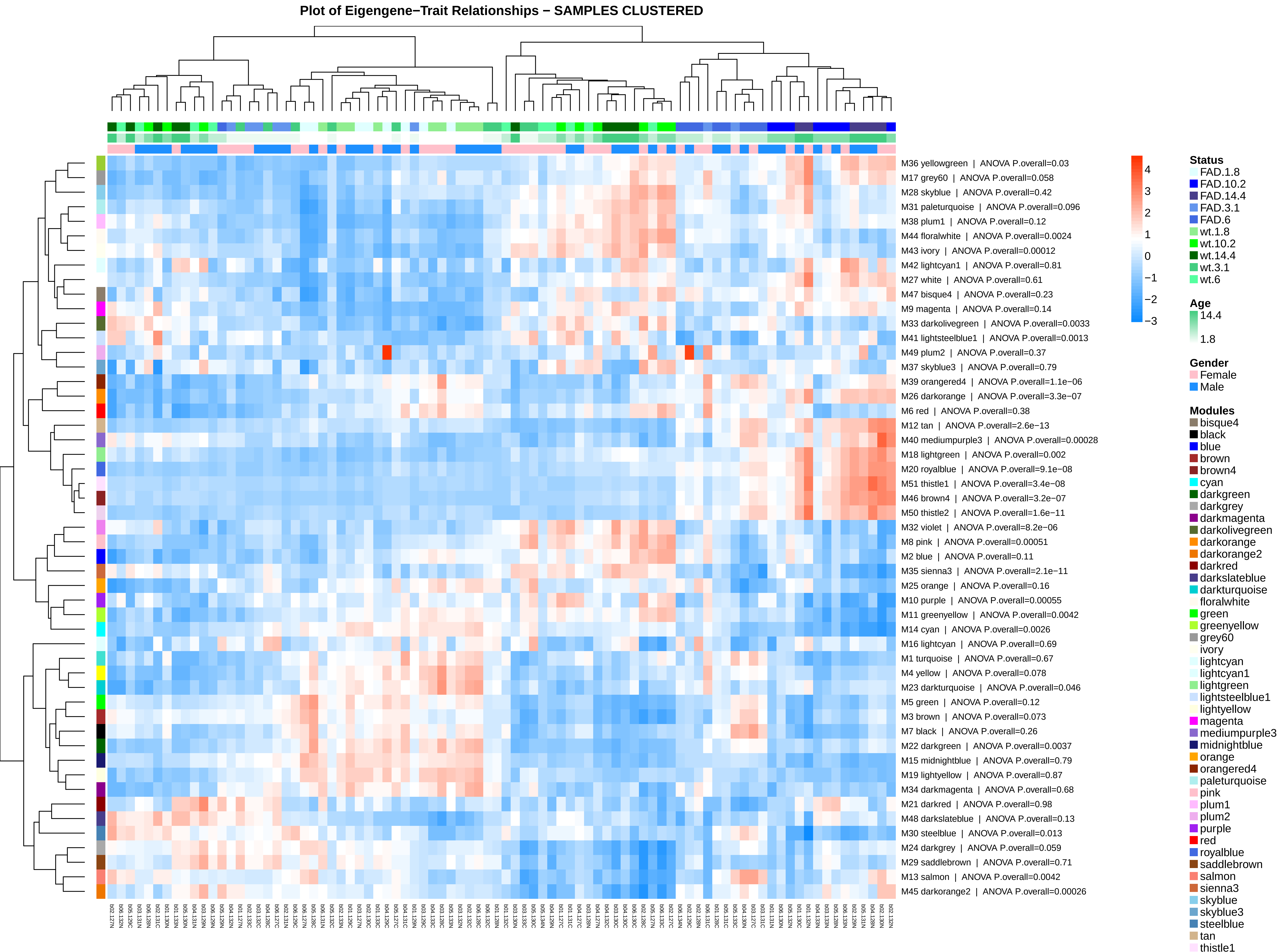
Plot of Eigengene−Trait Relationships − SAMPLES CLUSTERED
M36 yellowgreen | ANOVA P.overall=0.03
M17 grey60 | ANOVA P.overall=0.058
M28 skyblue | ANOVA P.overall=0.42
M31 paleturquoise | ANOVA P.overall=0.096
M38 plum1 | ANOVA P.overall=0.12
M44 floralwhite | ANOVA P.overall=0.0024
M43 ivory | ANOVA P.overall=0.00012
M42 lightcyan1 | ANOVA P.overall=0.81
M27 white | ANOVA P.overall=0.61
M47 bisque4 | ANOVA P.overall=0.23
M9 magenta | ANOVA P.overall=0.14
M33 darkolivegreen | ANOVA P.overall=0.0033
M41 lightsteelblue1 | ANOVA P.overall=0.0013
M49 plum2 | ANOVA P.overall=0.37
M37 skyblue3 | ANOVA P.overall=0.79
M39 orangered4 | ANOVA P.overall=1.1e−06
M26 darkorange | ANOVA P.overall=3.3e−07
M6 red | ANOVA P.overall=0.38
M12 tan | ANOVA P.overall=2.6e−13
M40 mediumpurple3 | ANOVA P.overall=0.00028
M18 lightgreen | ANOVA P.overall=0.002
M20 royalblue | ANOVA P.overall=9.1e−08
M51 thistle1 | ANOVA P.overall=3.4e−08
M46 brown4 | ANOVA P.overall=3.2e−07
M50 thistle2 | ANOVA P.overall=1.6e−11
M32 violet | ANOVA P.overall=8.2e−06
M8 pink | ANOVA P.overall=0.00051
M2 blue | ANOVA P.overall=0.11
M35 sienna3 | ANOVA P.overall=2.1e−11
M25 orange | ANOVA P.overall=0.16
M10 purple | ANOVA P.overall=0.00055
M11 greenyellow | ANOVA P.overall=0.0042
M14 cyan | ANOVA P.overall=0.0026
M16 lightcyan | ANOVA P.overall=0.69
M1 turquoise | ANOVA P.overall=0.67
M4 yellow | ANOVA P.overall=0.078
M23 darkturquoise | ANOVA P.overall=0.046
M5 green | ANOVA P.overall=0.12
M3 brown | ANOVA P.overall=0.073
M7 black | ANOVA P.overall=0.26
M22 darkgreen | ANOVA P.overall=0.0037
M15 midnightblue | ANOVA P.overall=0.79
M19 lightyellow | ANOVA P.overall=0.87
M34 darkmagenta | ANOVA P.overall=0.68
M21 darkred | ANOVA P.overall=0.98
M48 darkslateblue | ANOVA P.overall=0.13
M30 steelblue | ANOVA P.overall=0.013
M24 darkgrey | ANOVA P.overall=0.059
M29 saddlebrown | ANOVA P.overall=0.71
M13 salmon | ANOVA P.overall=0.0042
M45 darkorange2 | ANOVA P.overall=0.00026
4
3
2
1
0
−1
−2
−3
Status
FAD.1.8
FAD.10.2
FAD.14.4
FAD.3.1
FAD.6
wt.1.8
wt.10.2
wt.14.4
wt.3.1
wt.6
Age
14.4
1.8
Gender
Female
Male
Modules
bisque4
black
blue
brown
brown4
cyan
darkgreen
darkgrey
darkmagenta
darkolivegreen
darkorange
darkorange2
darkred
darkslateblue
darkturquoise
floralwhite
green
greenyellow
grey60
ivory
lightcyan
lightcyan1
lightgreen
lightsteelblue1
lightyellow
magenta
mediumpurple3
midnightblue
orange
orangered4
paleturquoise
pink
plum1
plum2
purple
red
royalblue
saddlebrown
salmon
sienna3
skyblue
skyblue3
steelblue
tan
thistle1

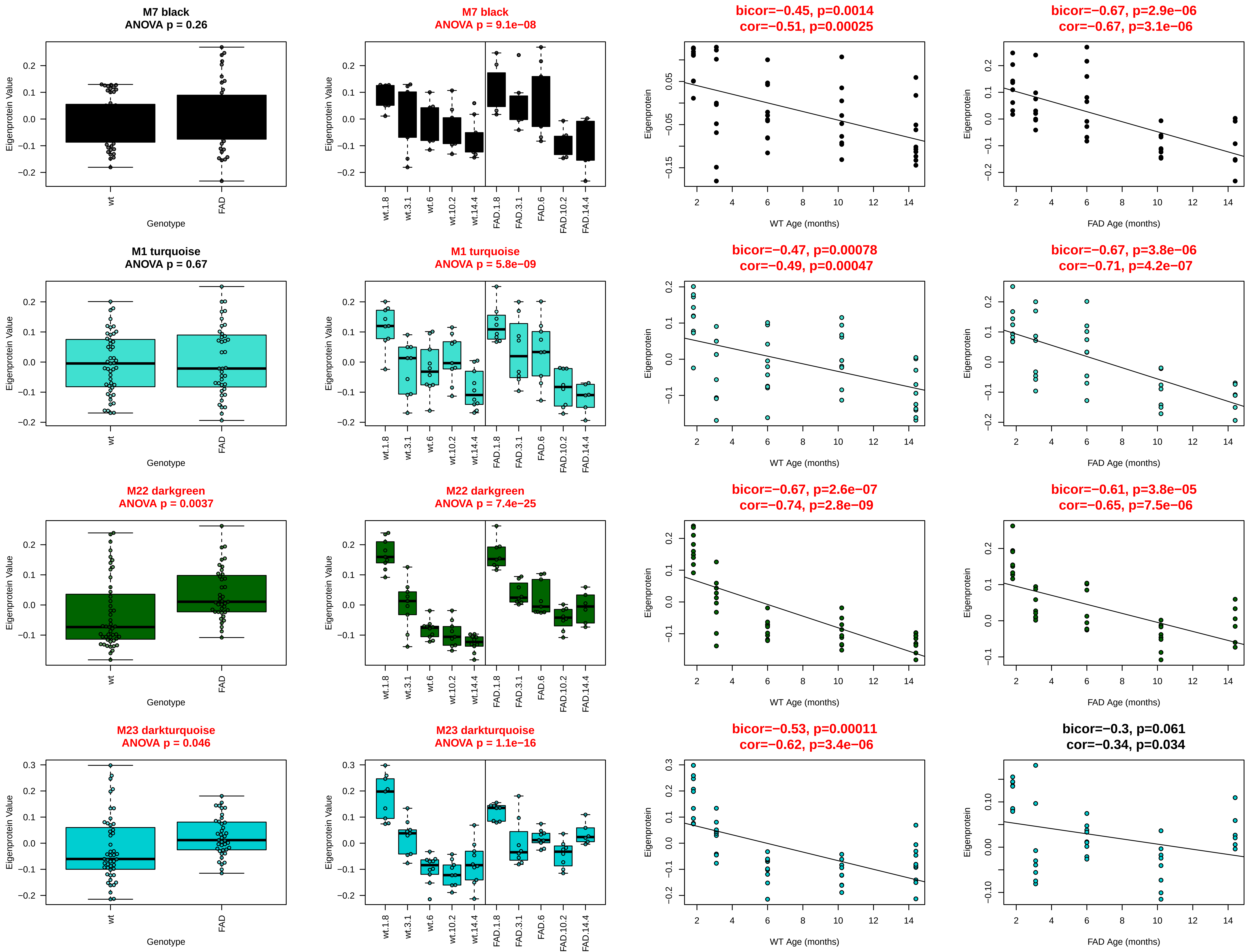

M7 black
ANOVA p = 0.26
M7 black
ANOVA p = 9.1e−08
bicor=−0.45, p=0.0014
cor=−0.51, p=0.00025
bicor=−0.67, p=2.9e−06
cor=−0.67, p=3.1e−06
M1 turquoise
ANOVA p = 0.67
M1 turquoise
ANOVA p = 5.8e−09
bicor=−0.47, p=0.00078
cor=−0.49, p=0.00047
bicor=−0.67, p=3.8e−06
cor=−0.71, p=4.2e−07
M22 darkgreen
ANOVA p = 0.0037
M22 darkgreen
ANOVA p = 7.4e−25
bicor=−0.67, p=2.6e−07
cor=−0.74, p=2.8e−09
bicor=−0.61, p=3.8e−05
cor=−0.65, p=7.5e−06
M23 darkturquoise
ANOVA p = 0.046
M23 darkturquoise
ANOVA p = 1.1e−16
bicor=−0.53, p=0.00011
cor=−0.62, p=3.4e−06
bicor=−0.3, p=0.061
cor=−0.34, p=0.034
Eigenprotein Value
Eigenprotein
Genotype
wt
FAD
wt.1.8
wt.3.1
wt.6
wt.10.2
wt.14.4
FAD.1.8
FAD.3.1
FAD.6
FAD.10.2
FAD.14.4
WT Age (months)
FAD Age (months)

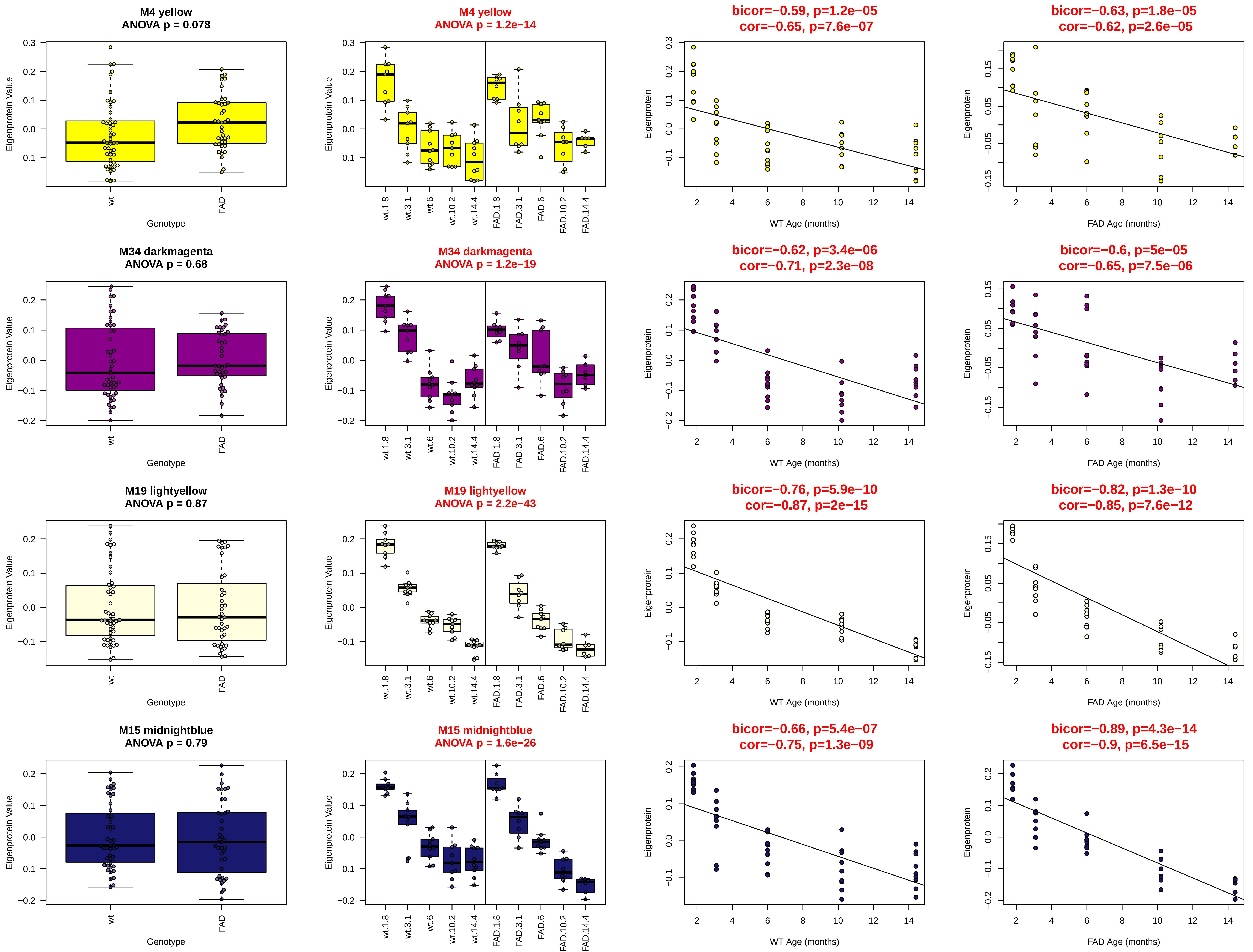

M4 yellow
ANOVA p = 0.078
M4 yellow
ANOVA p = 1.2e−14
bicor=−0.59, p=1.2e−05
cor=−0.65, p=7.6e−07
bicor=−0.63, p=1.8e−05
cor=−0.62, p=2.6e−05
M34 darkmagenta
ANOVA p = 0.68
M34 darkmagenta
ANOVA p = 1.2e−19
bicor=−0.62, p=3.4e−06
cor=−0.71, p=2.3e−08
bicor=−0.6, p=5e−05
cor=−0.65, p=7.5e−06
M19 lightyellow
ANOVA p = 0.87
M19 lightyellow
ANOVA p = 2.2e−43
bicor=−0.76, p=5.9e−10
cor=−0.87, p=2e−15
bicor=−0.82, p=1.3e−10
cor=−0.85, p=7.6e−12
M15 midnightblue
ANOVA p = 0.79
M15 midnightblue
ANOVA p = 1.6e−26
bicor=−0.66, p=5.4e−07
cor=−0.75, p=1.3e−09
bicor=−0.89, p=4.3e−14
cor=−0.9, p=6.5e−15
Eigenprotein Value
Eigenprotein
Genotype
wt
FAD
wt.1.8
wt.3.1
wt.6
wt.10.2
wt.14.4
FAD.1.8
FAD.3.1
FAD.6
FAD.10.2
FAD.14.4
WT Age (months)
FAD Age (months)

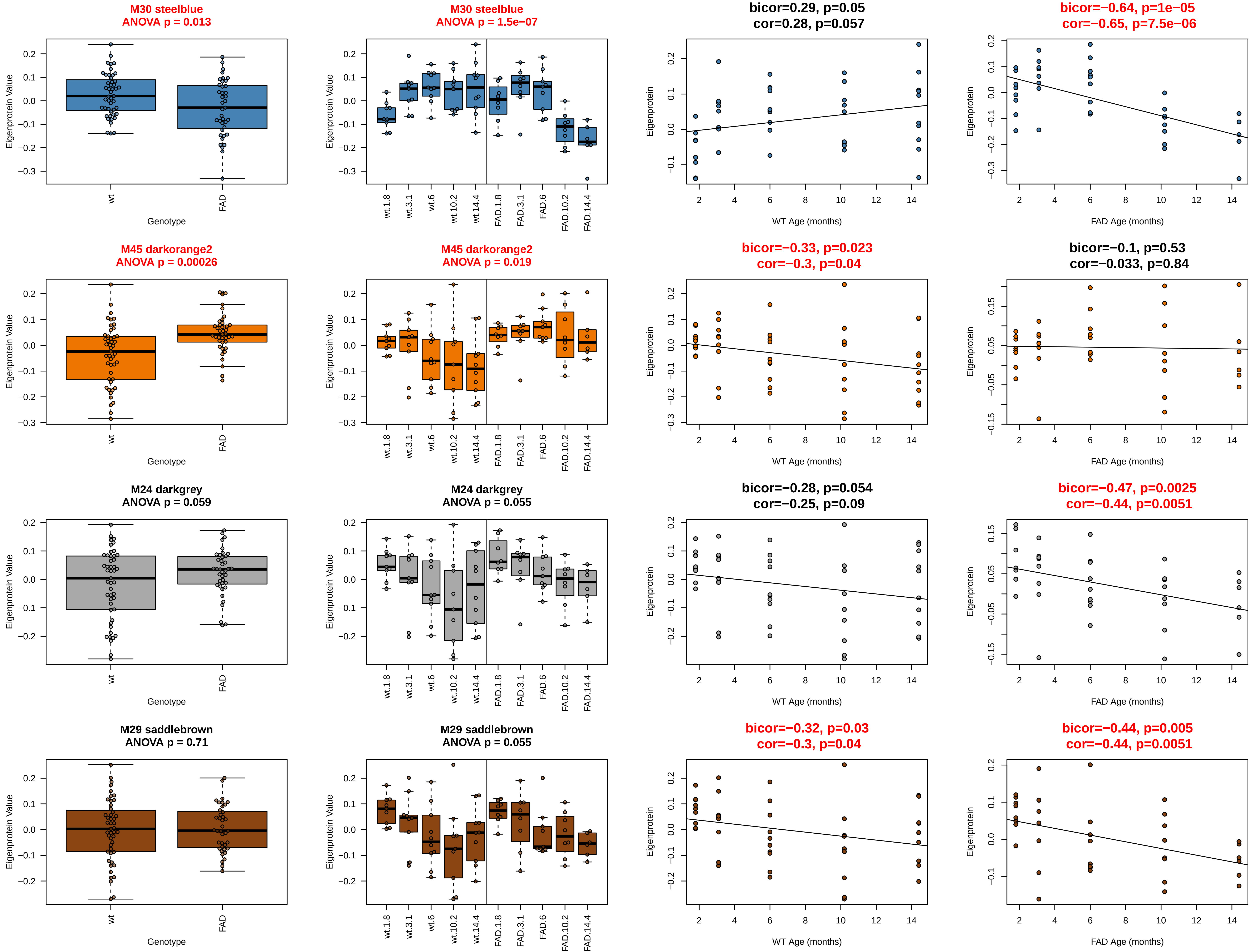
M30 steelblue
ANOVA p = 0.013
Eigenprotein Value
Genotype
wt
FAD
M30 steelblue
ANOVA p = 1.5e−07
wt.1.8
wt.3.1
wt.6
wt.10.2
wt.14.4
FAD.1.8
FAD.3.1
FAD.6
FAD.10.2
FAD.14.4
bicor=0.29, p=0.05
cor=0.28, p=0.057
Eigenprotein
WT Age (months)
bicor=−0.64, p=1e−05
cor=−0.65, p=7.5e−06
FAD Age (months)
M45 darkorange2
ANOVA p = 0.00026
M45 darkorange2
ANOVA p = 0.019
bicor=−0.33, p=0.023
cor=−0.3, p=0.04
bicor=−0.1, p=0.53
cor=−0.033, p=0.84
M24 darkgrey
ANOVA p = 0.059
M24 darkgrey
ANOVA p = 0.055
bicor=−0.28, p=0.054
cor=−0.25, p=0.09
bicor=−0.47, p=0.0025
cor=−0.44, p=0.0051
M29 saddlebrown
ANOVA p = 0.71
M29 saddlebrown
ANOVA p = 0.055
bicor=−0.32, p=0.03
cor=−0.3, p=0.04
bicor=−0.44, p=0.005
cor=−0.44, p=0.0051

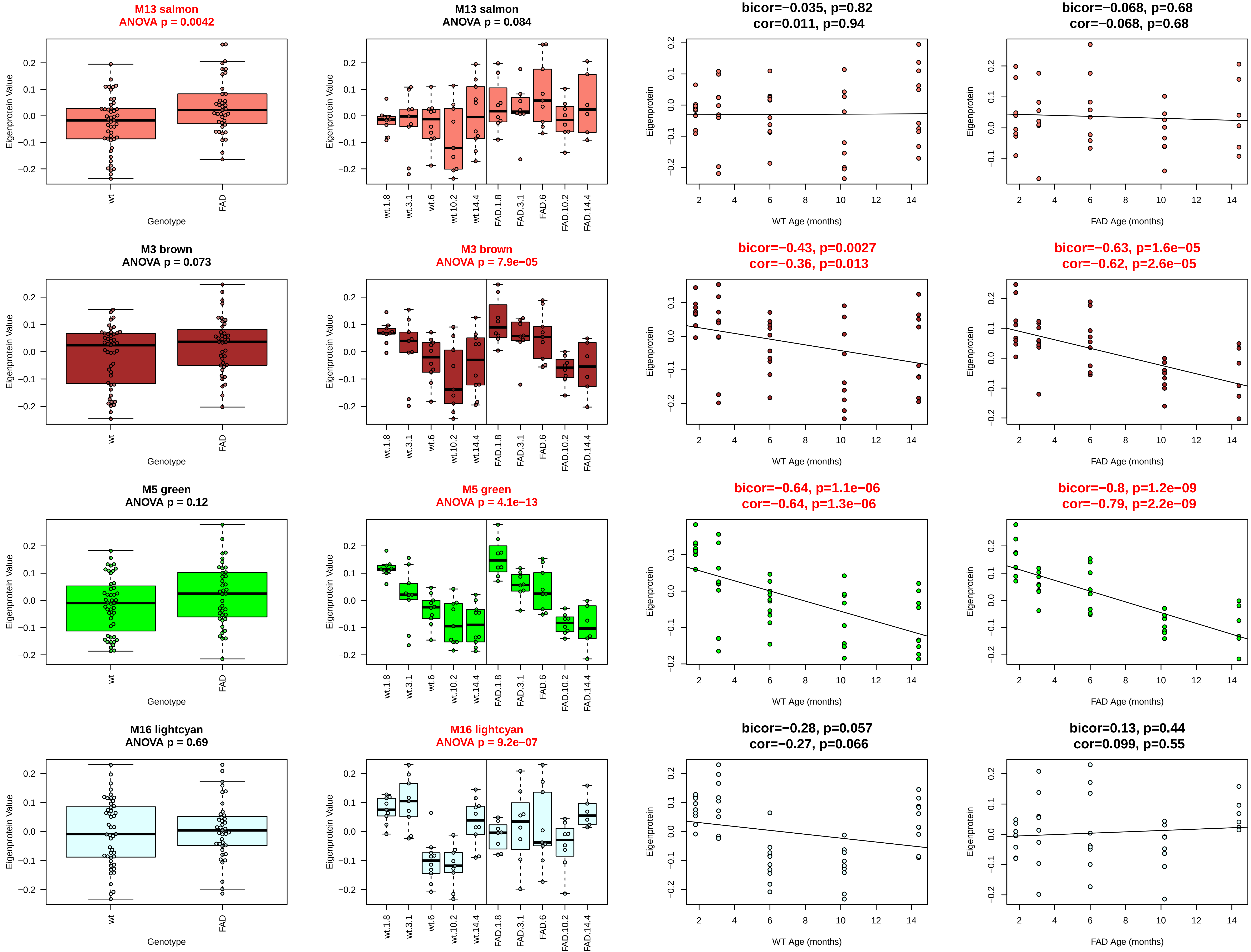
M13 salmon
ANOVA p = 0.0042
M13 salmon
ANOVA p = 0.084
bicor=−0.035, p=0.82
cor=0.011, p=0.94
bicor=−0.068, p=0.68
cor=−0.068, p=0.68
M3 brown
ANOVA p = 0.073
M3 brown
ANOVA p = 7.9e−05
bicor=−0.43, p=0.0027
cor=−0.36, p=0.013
bicor=−0.63, p=1.6e−05
cor=−0.62, p=2.6e−05
M5 green
ANOVA p = 0.12
M5 green
ANOVA p = 4.1e−13
bicor=−0.64, p=1.1e−06
cor=−0.64, p=1.3e−06
bicor=−0.8, p=1.2e−09
cor=−0.79, p=2.2e−09
M16 lightcyan
ANOVA p = 0.69
M16 lightcyan
ANOVA p = 9.2e−07
bicor=−0.28, p=0.057
cor=−0.27, p=0.066
bicor=0.13, p=0.44
cor=0.099, p=0.55
Eigenprotein Value
Eigenprotein
wt
FAD
Genotype
wt.1.8
wt.3.1
wt.6
wt.10.2
wt.14.4
FAD.1.8
FAD.3.1
FAD.6
FAD.10.2
FAD.14.4
WT Age (months)
FAD Age (months)

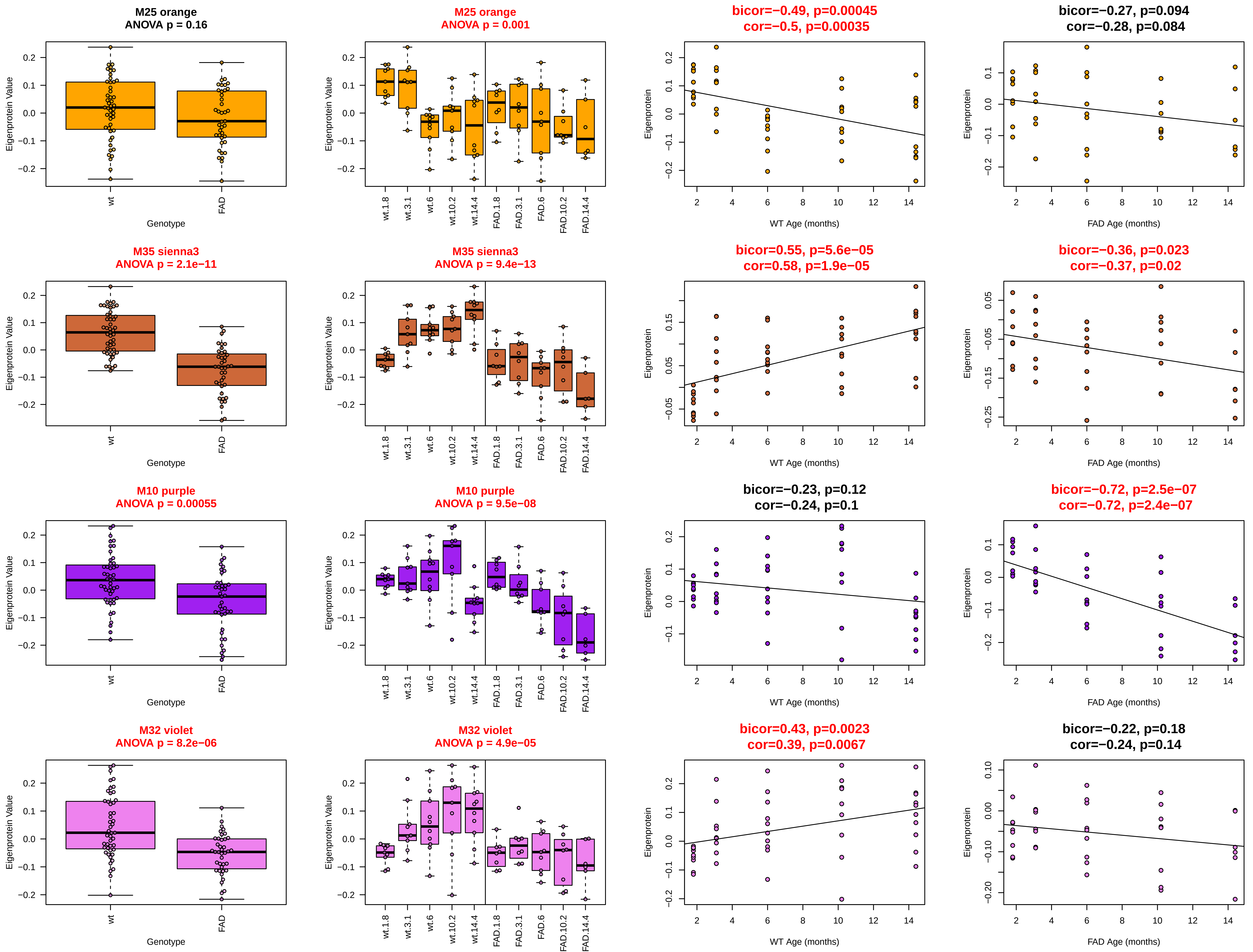

M25 orange
ANOVA p = 0.16
M25 orange
ANOVA p = 0.001
bicor=−0.49, p=0.00045
cor=−0.5, p=0.00035
bicor=−0.27, p=0.094
cor=−0.28, p=0.084
M35 sienna3
ANOVA p = 2.1e−11
M35 sienna3
ANOVA p = 9.4e−13
bicor=0.55, p=5.6e−05
cor=0.58, p=1.9e−05
bicor=−0.36, p=0.023
cor=−0.37, p=0.02
M10 purple
ANOVA p = 0.00055
M10 purple
ANOVA p = 9.5e−08
bicor=−0.23, p=0.12
cor=−0.24, p=0.1
bicor=−0.72, p=2.5e−07
cor=−0.72, p=2.4e−07
M32 violet
ANOVA p = 8.2e−06
M32 violet
ANOVA p = 4.9e−05
bicor=0.43, p=0.0023
cor=0.39, p=0.0067
bicor=−0.22, p=0.18
cor=−0.24, p=0.14
Eigenprotein Value
Eigenprotein
Genotype
wt
FAD
wt.1.8
wt.3.1
wt.6
wt.10.2
wt.14.4
FAD.1.8
FAD.3.1
FAD.6
FAD.10.2
FAD.14.4
WT Age (months)
FAD Age (months)

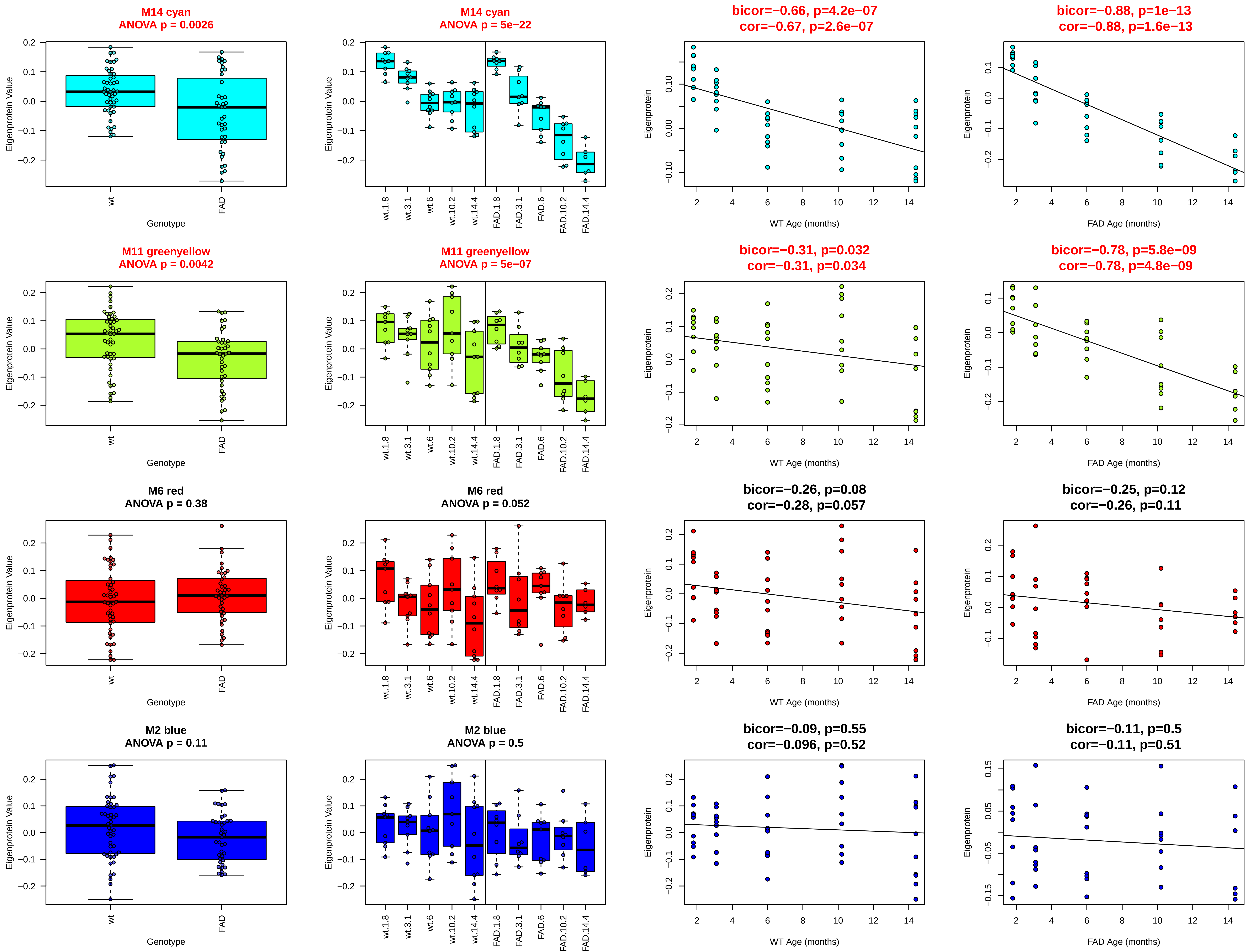
M14 cyan
ANOVA p = 0.0026
M14 cyan
ANOVA p = 5e−22
bicor=−0.66, p=4.2e−07
cor=−0.67, p=2.6e−07
bicor=−0.88, p=1e−13
cor=−0.88, p=1.6e−13
M11 greenyellow
ANOVA p = 0.0042
M11 greenyellow
ANOVA p = 5e−07
bicor=−0.31, p=0.032
cor=−0.31, p=0.034
bicor=−0.78, p=5.8e−09
cor=−0.78, p=4.8e−09
M6 red
ANOVA p = 0.38
M6 red
ANOVA p = 0.052
bicor=−0.26, p=0.08
cor=−0.28, p=0.057
bicor=−0.25, p=0.12
cor=−0.26, p=0.11
M2 blue
ANOVA p = 0.11
M2 blue
ANOVA p = 0.5
bicor=−0.09, p=0.55
cor=−0.096, p=0.52
bicor=−0.11, p=0.5
cor=−0.11, p=0.51
Eigenprotein Value
Eigenprotein
Genotype
wt
FAD
wt.1.8
wt.3.1
wt.6
wt.10.2
wt.14.4
FAD.1.8
FAD.3.1
FAD.6
FAD.10.2
FAD.14.4
WT Age (months)
FAD Age (months)

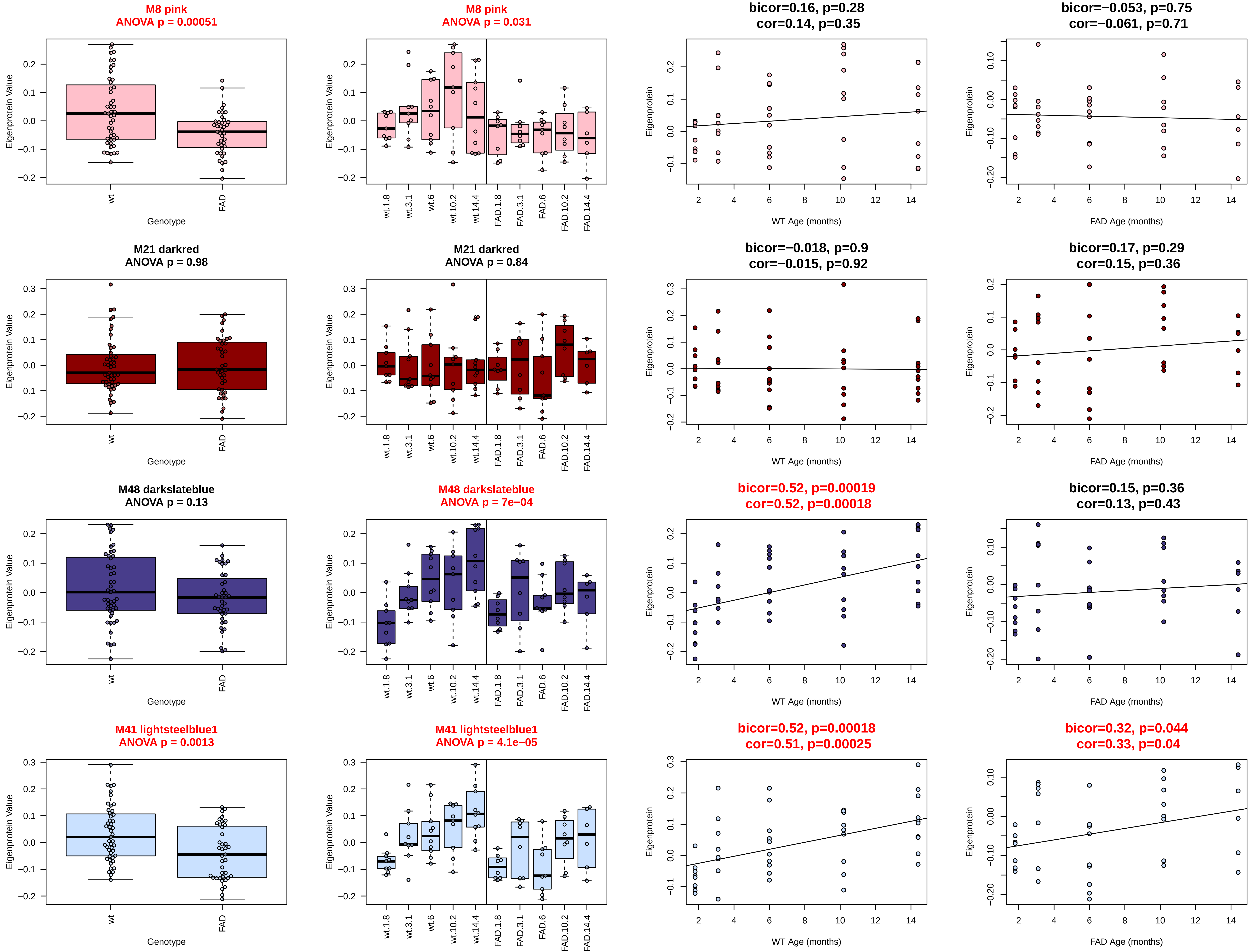

M8 pink
ANOVA p = 0.00051
M8 pink
ANOVA p = 0.031
bicor=0.16, p=0.28
cor=0.14, p=0.35
bicor=−0.053, p=0.75
cor=−0.061, p=0.71
M21 darkred
ANOVA p = 0.98
M21 darkred
ANOVA p = 0.84
bicor=−0.018, p=0.9
cor=−0.015, p=0.92
bicor=0.17, p=0.29
cor=0.15, p=0.36
M48 darkslateblue
ANOVA p = 0.13
M48 darkslateblue
ANOVA p = 7e−04
bicor=0.52, p=0.00019
cor=0.52, p=0.00018
bicor=0.15, p=0.36
cor=0.13, p=0.43
M41 lightsteelblue1
ANOVA p = 0.0013
M41 lightsteelblue1
ANOVA p = 4.1e−05
bicor=0.52, p=0.00018
cor=0.51, p=0.00025
bicor=0.32, p=0.044
cor=0.33, p=0.04
Eigenprotein Value
Eigenprotein
Genotype
wt
FAD
wt.1.8
wt.3.1
wt.6
wt.10.2
wt.14.4
FAD.1.8
FAD.3.1
FAD.6
FAD.10.2
FAD.14.4
WT Age (months)
FAD Age (months)

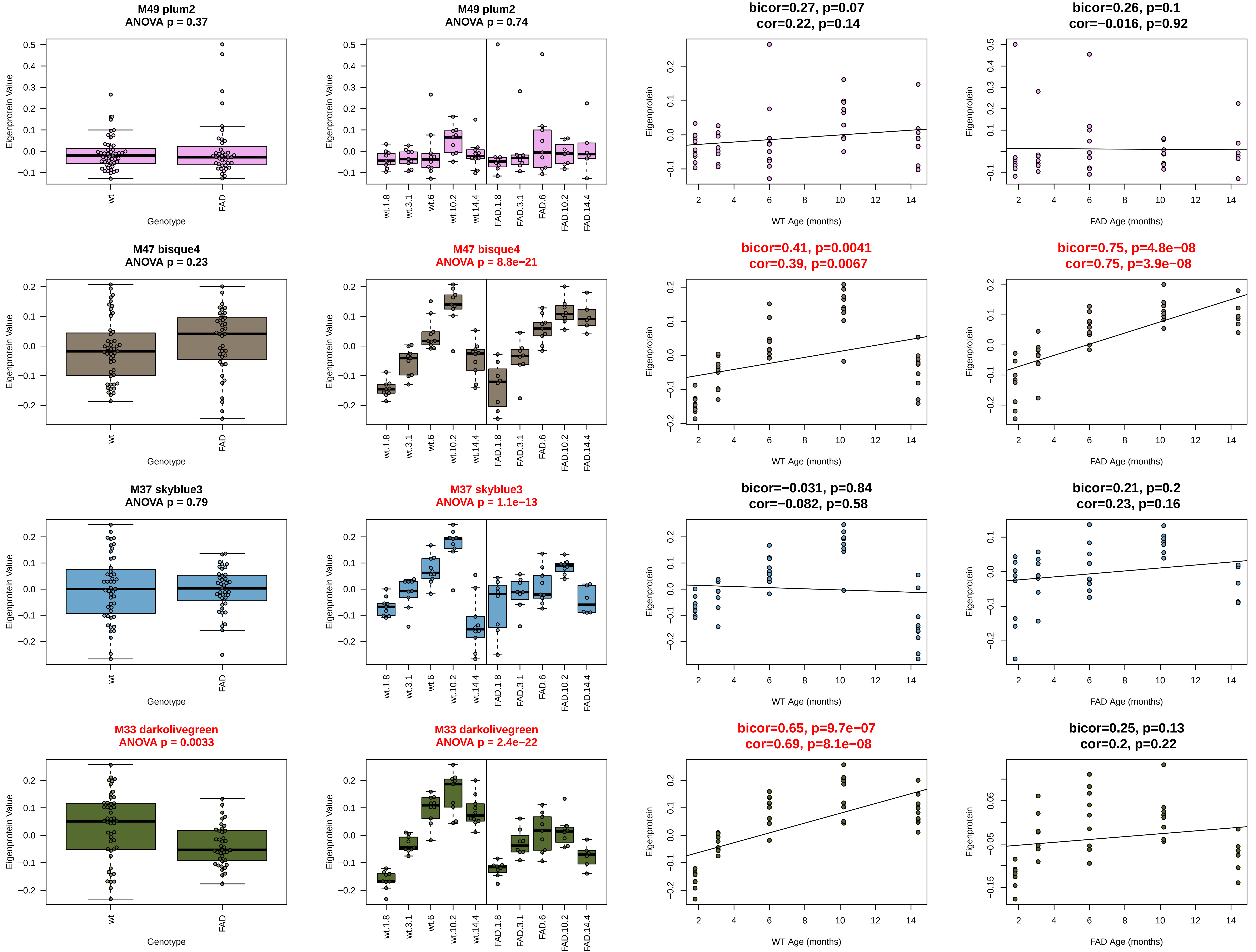
M49 plum2
ANOVA p = 0.37
M49 plum2
ANOVA p = 0.74
bicor=0.27, p=0.07
cor=0.22, p=0.14
bicor=0.26, p=0.1
cor=−0.016, p=0.92
M47 bisque4
ANOVA p = 0.23
M47 bisque4
ANOVA p = 8.8e−21
bicor=0.41, p=0.0041
cor=0.39, p=0.0067
bicor=0.75, p=4.8e−08
cor=0.75, p=3.9e−08
M37 skyblue3
ANOVA p = 0.79
M37 skyblue3
ANOVA p = 1.1e−13
bicor=−0.031, p=0.84
cor=−0.082, p=0.58
bicor=0.21, p=0.2
cor=0.23, p=0.16
M33 darkolivegreen
ANOVA p = 0.0033
M33 darkolivegreen
ANOVA p = 2.4e−22
bicor=0.65, p=9.7e−07
cor=0.69, p=8.1e−08
bicor=0.25, p=0.13
cor=0.2, p=0.22
Eigenprotein Value
Eigenprotein
Genotype
wt
FAD
wt.1.8
wt.3.1
wt.6
wt.10.2
wt.14.4
FAD.1.8
FAD.3.1
FAD.6
FAD.10.2
FAD.14.4
WT Age (months)
FAD Age (months)

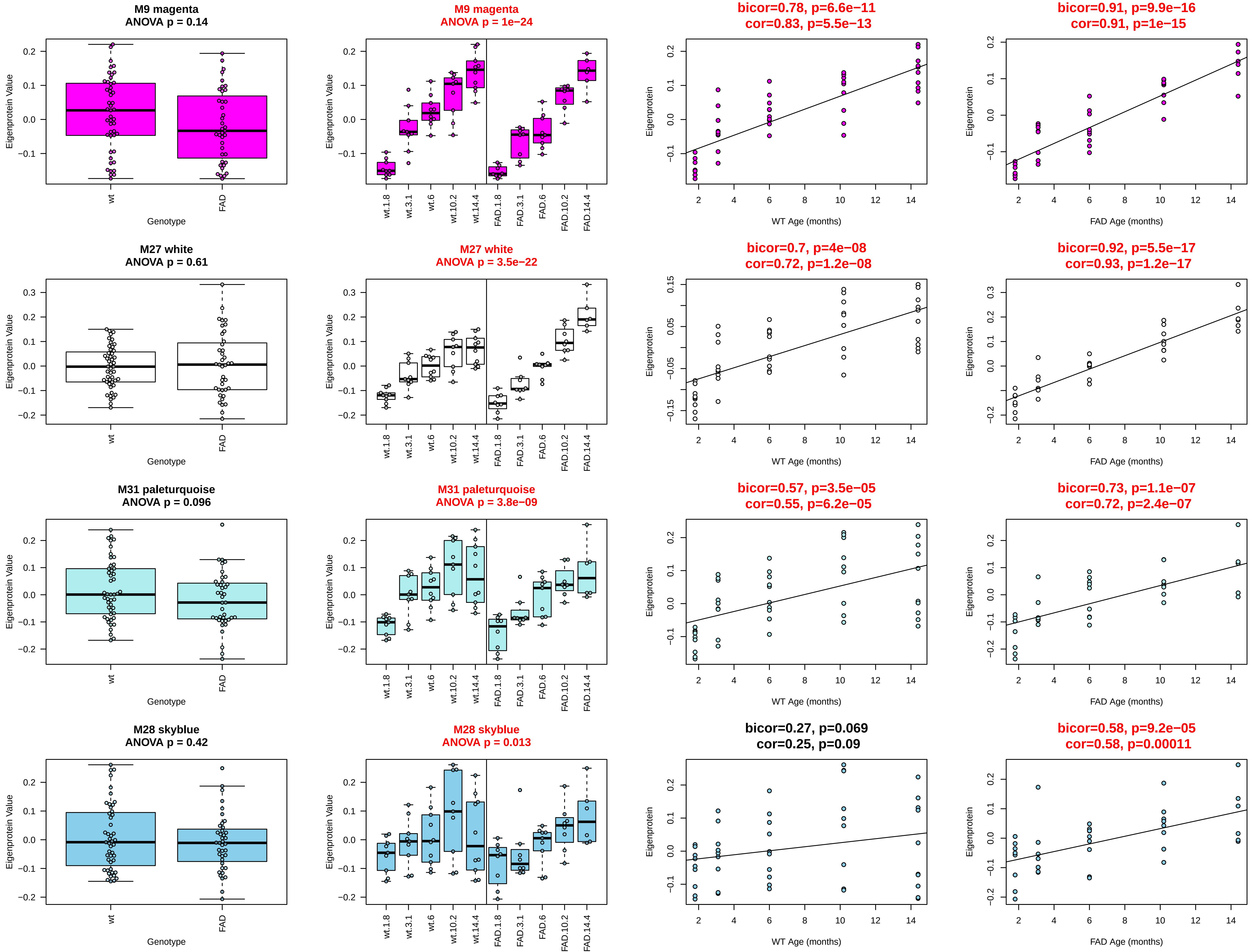

M9 magenta
ANOVA p = 0.14
M9 magenta
ANOVA p = 1e−24
bicor=0.78, p=6.6e−11
cor=0.83, p=5.5e−13
bicor=0.91, p=9.9e−16
cor=0.91, p=1e−15
M27 white
ANOVA p = 0.61
M27 white
ANOVA p = 3.5e−22
bicor=0.7, p=4e−08
cor=0.72, p=1.2e−08
bicor=0.92, p=5.5e−17
cor=0.93, p=1.2e−17
M31 paleturquoise
ANOVA p = 0.096
M31 paleturquoise
ANOVA p = 3.8e−09
bicor=0.57, p=3.5e−05
cor=0.55, p=6.2e−05
bicor=0.73, p=1.1e−07
cor=0.72, p=2.4e−07
M28 skyblue
ANOVA p = 0.42
M28 skyblue
ANOVA p = 0.013
bicor=0.27, p=0.069
cor=0.25, p=0.09
bicor=0.58, p=9.2e−05
cor=0.58, p=0.00011
Eigenprotein Value
Eigenprotein
Genotype
wt
FAD
wt.1.8
wt.3.1
wt.6
wt.10.2
wt.14.4
FAD.1.8
FAD.3.1
FAD.6
FAD.10.2
FAD.14.4
WT Age (months)
FAD Age (months)

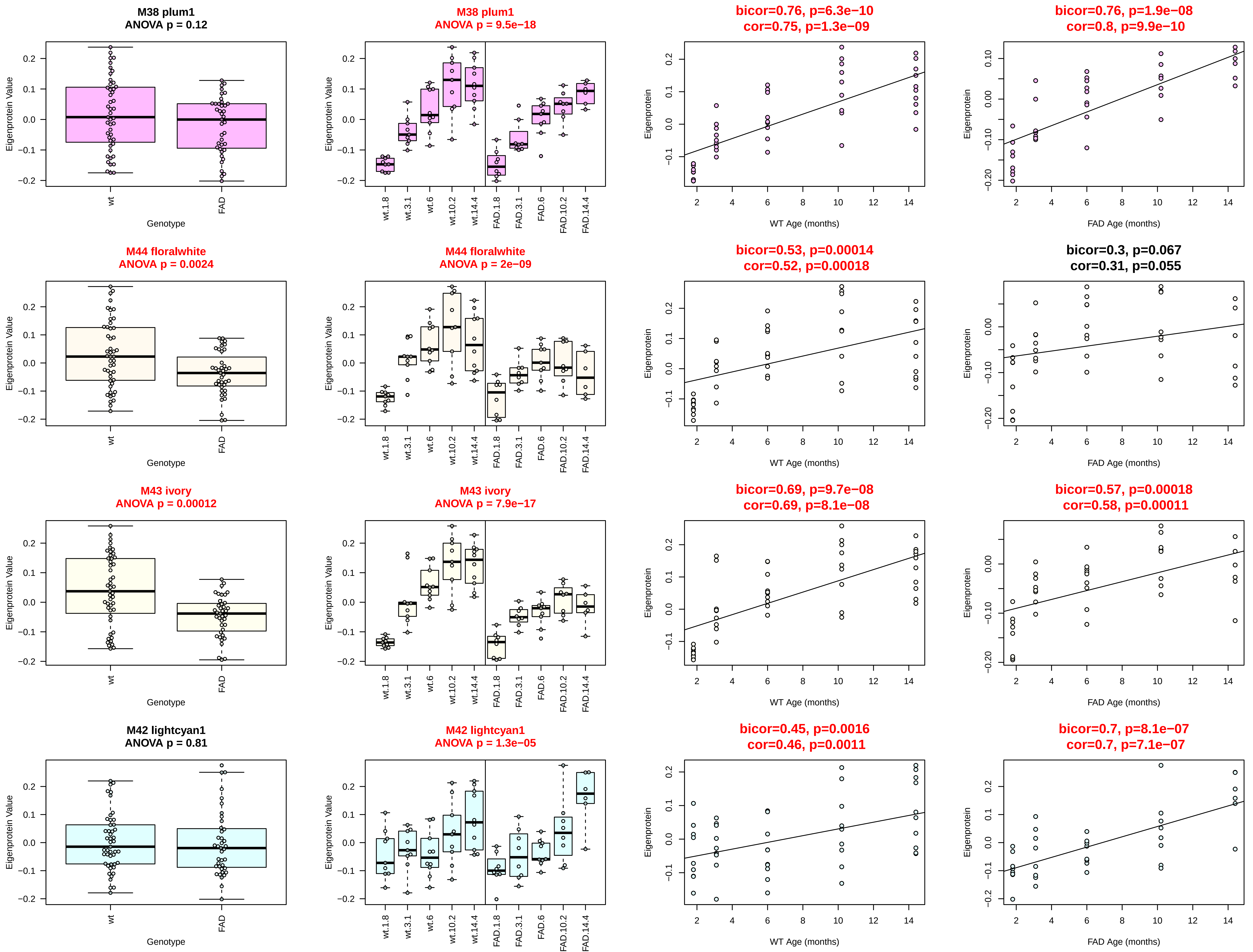

M38 plum1
ANOVA p = 0.12
M38 plum1
ANOVA p = 9.5e−18
bicor=0.76, p=6.3e−10
cor=0.75, p=1.3e−09
bicor=0.76, p=1.9e−08
cor=0.8, p=9.9e−10
M44 floralwhite
ANOVA p = 0.0024
M44 floralwhite
ANOVA p = 2e−09
bicor=0.53, p=0.00014
cor=0.52, p=0.00018
bicor=0.3, p=0.067
cor=0.31, p=0.055
M43 ivory
ANOVA p = 0.00012
M43 ivory
ANOVA p = 7.9e−17
bicor=0.69, p=9.7e−08
cor=0.69, p=8.1e−08
bicor=0.57, p=0.00018
cor=0.58, p=0.00011
M42 lightcyan1
ANOVA p = 0.81
M42 lightcyan1
ANOVA p = 1.3e−05
bicor=0.45, p=0.0016
cor=0.46, p=0.0011
bicor=0.7, p=8.1e−07
cor=0.7, p=7.1e−07
Eigenprotein Value
Eigenprotein
Genotype
wt
FAD
wt.1.8
wt.3.1
wt.6
wt.10.2
wt.14.4
FAD.1.8
FAD.3.1
FAD.6
FAD.10.2
FAD.14.4
WT Age (months)
FAD Age (months)

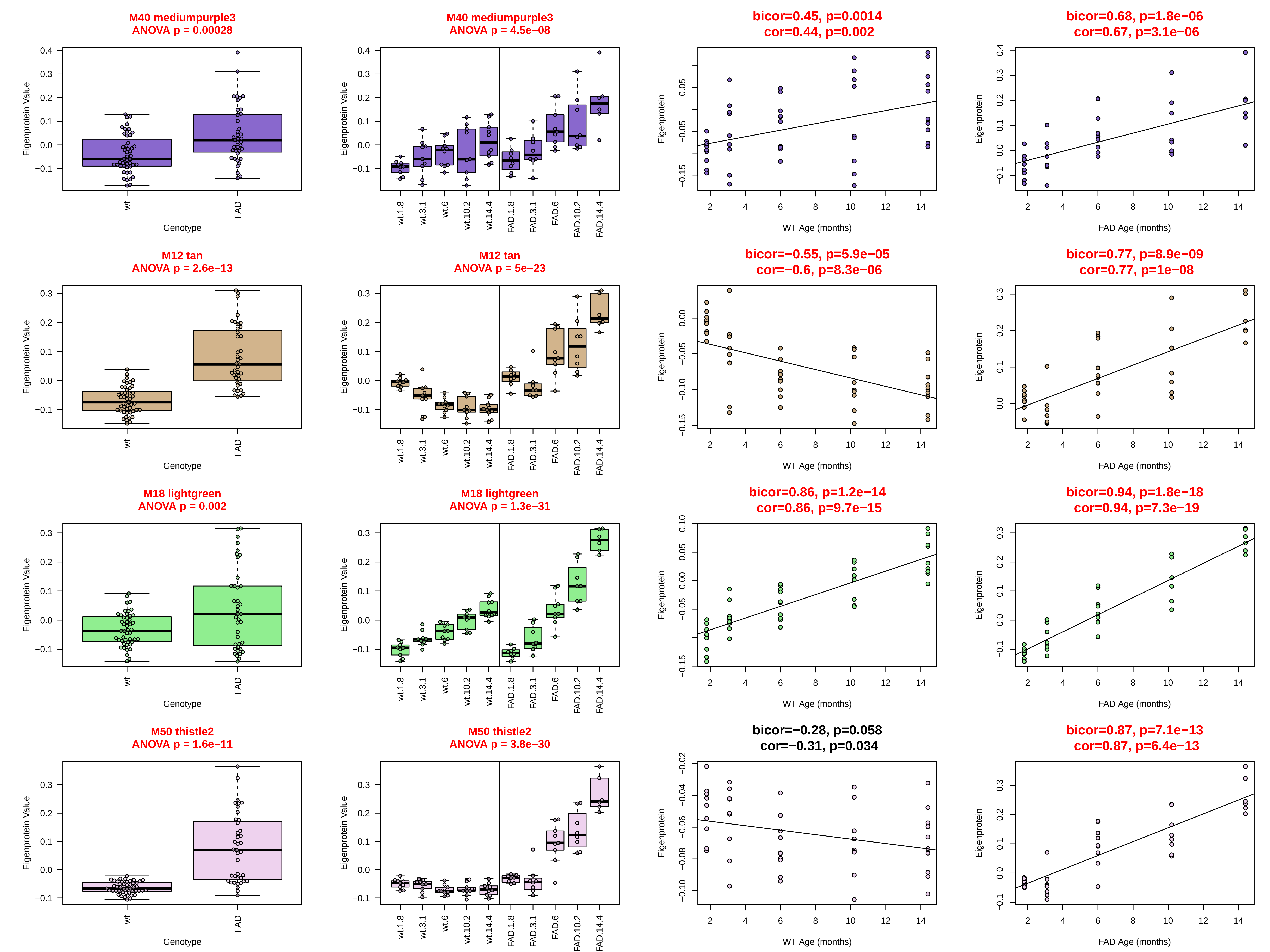

M40 mediumpurple3
ANOVA p = 0.00028
M40 mediumpurple3
ANOVA p = 4.5e−08
bicor=0.45, p=0.0014
cor=0.44, p=0.002
bicor=0.68, p=1.8e−06
cor=0.67, p=3.1e−06
M12 tan
ANOVA p = 2.6e−13
M12 tan
ANOVA p = 5e−23
bicor=−0.55, p=5.9e−05
cor=−0.6, p=8.3e−06
bicor=0.77, p=8.9e−09
cor=0.77, p=1e−08
M18 lightgreen
ANOVA p = 0.002
M18 lightgreen
ANOVA p = 1.3e−31
bicor=0.86, p=1.2e−14
cor=0.86, p=9.7e−15
bicor=0.94, p=1.8e−18
cor=0.94, p=7.3e−19
M50 thistle2
ANOVA p = 1.6e−11
M50 thistle2
ANOVA p = 3.8e−30
bicor=−0.28, p=0.058
cor=−0.31, p=0.034
bicor=0.87, p=7.1e−13
cor=0.87, p=6.4e−13
Eigenprotein Value
Eigenprotein
Genotype
wt
FAD
wt.1.8
wt.3.1
wt.6
wt.10.2
wt.14.4
FAD.1.8
FAD.3.1
FAD.6
FAD.10.2
FAD.14.4
WT Age (months)
FAD Age (months)

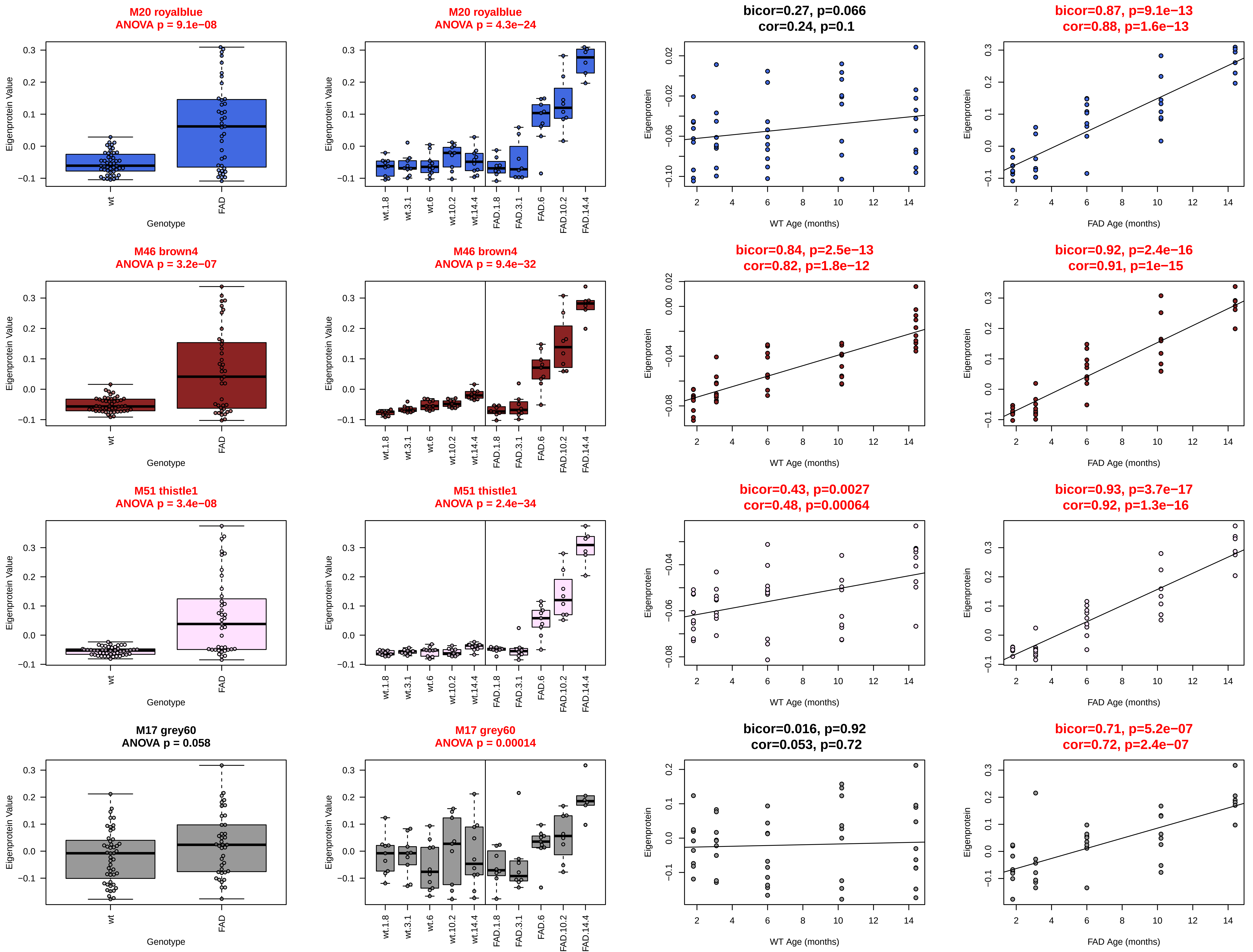
M20 royalblue
ANOVA p = 9.1e−08
M20 royalblue
ANOVA p = 4.3e−24
bicor=0.27, p=0.066
cor=0.24, p=0.1
bicor=0.87, p=9.1e−13
cor=0.88, p=1.6e−13
M46 brown4
ANOVA p = 3.2e−07
M46 brown4
ANOVA p = 9.4e−32
bicor=0.84, p=2.5e−13
cor=0.82, p=1.8e−12
bicor=0.92, p=2.4e−16
cor=0.91, p=1e−15
M51 thistle1
ANOVA p = 3.4e−08
M51 thistle1
ANOVA p = 2.4e−34
bicor=0.43, p=0.0027
cor=0.48, p=0.00064
bicor=0.93, p=3.7e−17
cor=0.92, p=1.3e−16
M17 grey60
ANOVA p = 0.058
M17 grey60
ANOVA p = 0.00014
bicor=0.016, p=0.92
cor=0.053, p=0.72
bicor=0.71, p=5.2e−07
cor=0.72, p=2.4e−07
Eigenprotein Value
Eigenprotein
Genotype
wt
FAD
wt.1.8
wt.3.1
wt.6
wt.10.2
wt.14.4
FAD.1.8
FAD.3.1
FAD.6
FAD.10.2
FAD.14.4
WT Age (months)
FAD Age (months)

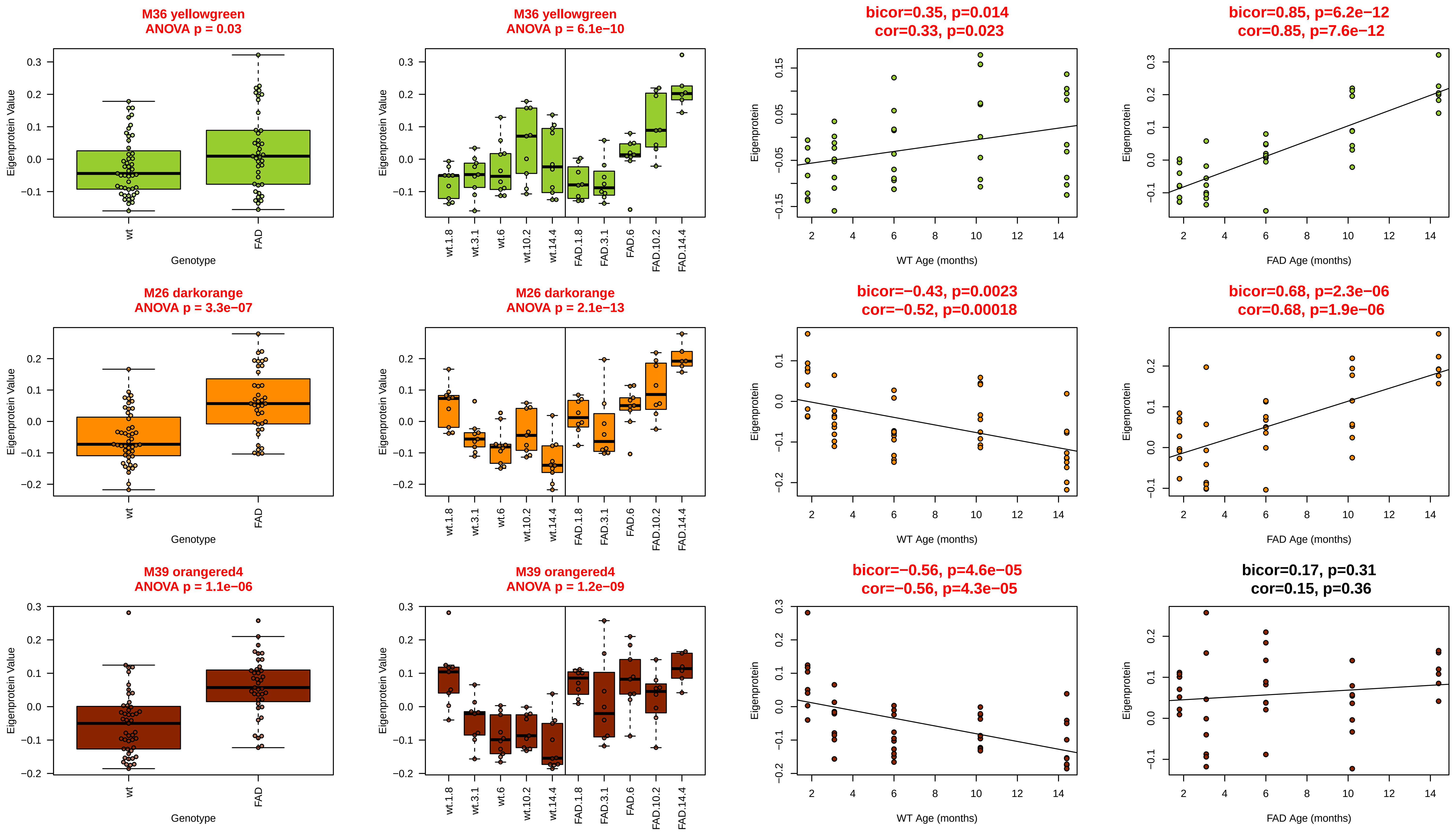
M36 yellowgreen
ANOVA p = 0.03
M36 yellowgreen
ANOVA p = 6.1e−10
bicor=0.35, p=0.014
cor=0.33, p=0.023
bicor=0.85, p=6.2e−12
cor=0.85, p=7.6e−12
M26 darkorange
ANOVA p = 3.3e−07
M26 darkorange
ANOVA p = 2.1e−13
bicor=−0.43, p=0.0023
cor=−0.52, p=0.00018
bicor=0.68, p=2.3e−06
cor=0.68, p=1.9e−06
M39 orangered4
ANOVA p = 1.1e−06
M39 orangered4
ANOVA p = 1.2e−09
bicor=−0.56, p=4.6e−05
cor=−0.56, p=4.3e−05
bicor=0.17, p=0.31
cor=0.15, p=0.36
Eigenprotein Value
Eigenprotein
Genotype
wt
FAD
wt.1.8
wt.3.1
wt.6
wt.10.2
wt.14.4
FAD.1.8
FAD.3.1
FAD.6
FAD.10.2
FAD.14.4
WT Age (months)
FAD Age (months)
