## Supplemental Data 3 for "Network analysis of mouse brain proteomes resolves molecular and cellular endophenotypes of aging and Aβ pathology"

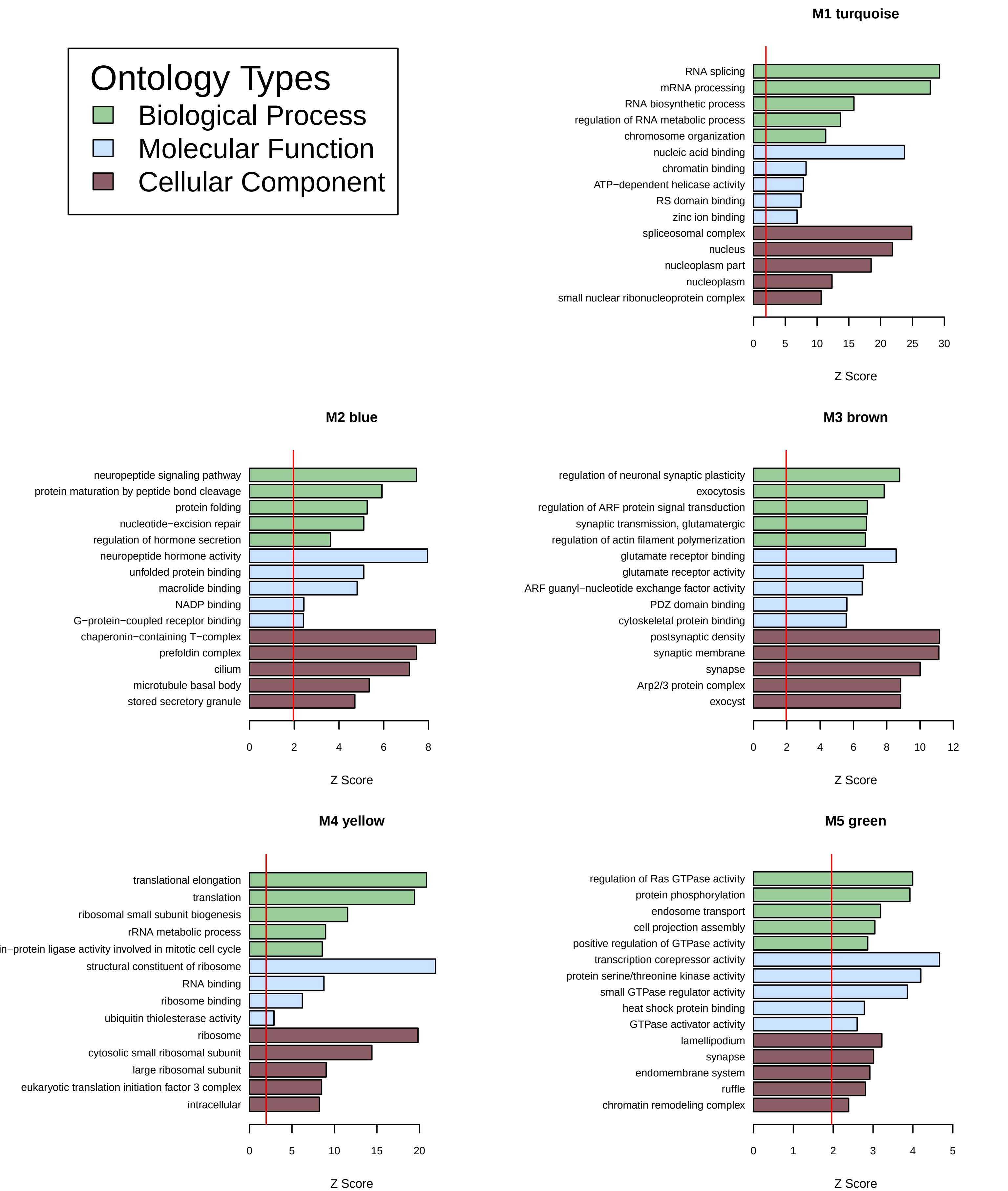
Ontology Types
Biological Process
Molecular Function
Cellular Component
M1 turquoise
RNA splicing
mRNA processing
RNA biosynthetic process
regulation of RNA metabolic process
chromosome organization
nucleic acid binding
chromatin binding
ATP−dependent helicase activity
RS domain binding
zinc ion binding
spliceosomal complex
nucleus
nucleoplasm part
nucleoplasm
small nuclear ribonucleoprotein complex
0 5 10 15 20 25 30
Z Score
M2 blue
neuropeptide signaling pathway
protein maturation by peptide bond cleavage
protein folding
nucleotide−excision repair
regulation of hormone secretion
neuropeptide hormone activity
unfolded protein binding
macrolide binding
NADP binding
G−protein−coupled receptor binding
chaperonin−containing T−complex
prefoldin complex
cilium
microtubule basal body
stored secretory granule
0 2 4 6 8
Z Score
M3 brown
regulation of neuronal synaptic plasticity
exocytosis
regulation of ARF protein signal transduction
synaptic transmission, glutamatergic
regulation of actin filament polymerization
glutamate receptor binding
glutamate receptor activity
ARF guanyl−nucleotide exchange factor activity
PDZ domain binding
cytoskeletal protein binding
postsynaptic density
synaptic membrane
synapse
Arp2/3 protein complex
exocyst
0 2 4 6 8 10 12
Z Score
M4 yellow
translational elongation
translation
ribosomal small subunit biogenesis
rRNA metabolic process
protein ligase activity involved in mitotic cell cycle
structural constituent of ribosome
RNA binding
ribosome binding
ubiquitin thiolesterase activity
ribosome
cytosolic small ribosomal subunit
large ribosomal subunit
eukaryotic translation initiation factor 3 complex
intracellular
0 5 10 15 20
Z Score
M5 green
regulation of Ras GTPase activity
protein phosphorylation
endosome transport
cell projection assembly
positive regulation of GTPase activity
transcription corepressor activity
protein serine/threonine kinase activity
small GTPase regulator activity
heat shock protein binding
GTPase activator activity
lamellipodium
synapse
endomembrane system
ruffle
chromatin remodeling complex
0 1 2 3 4 5
Z Score

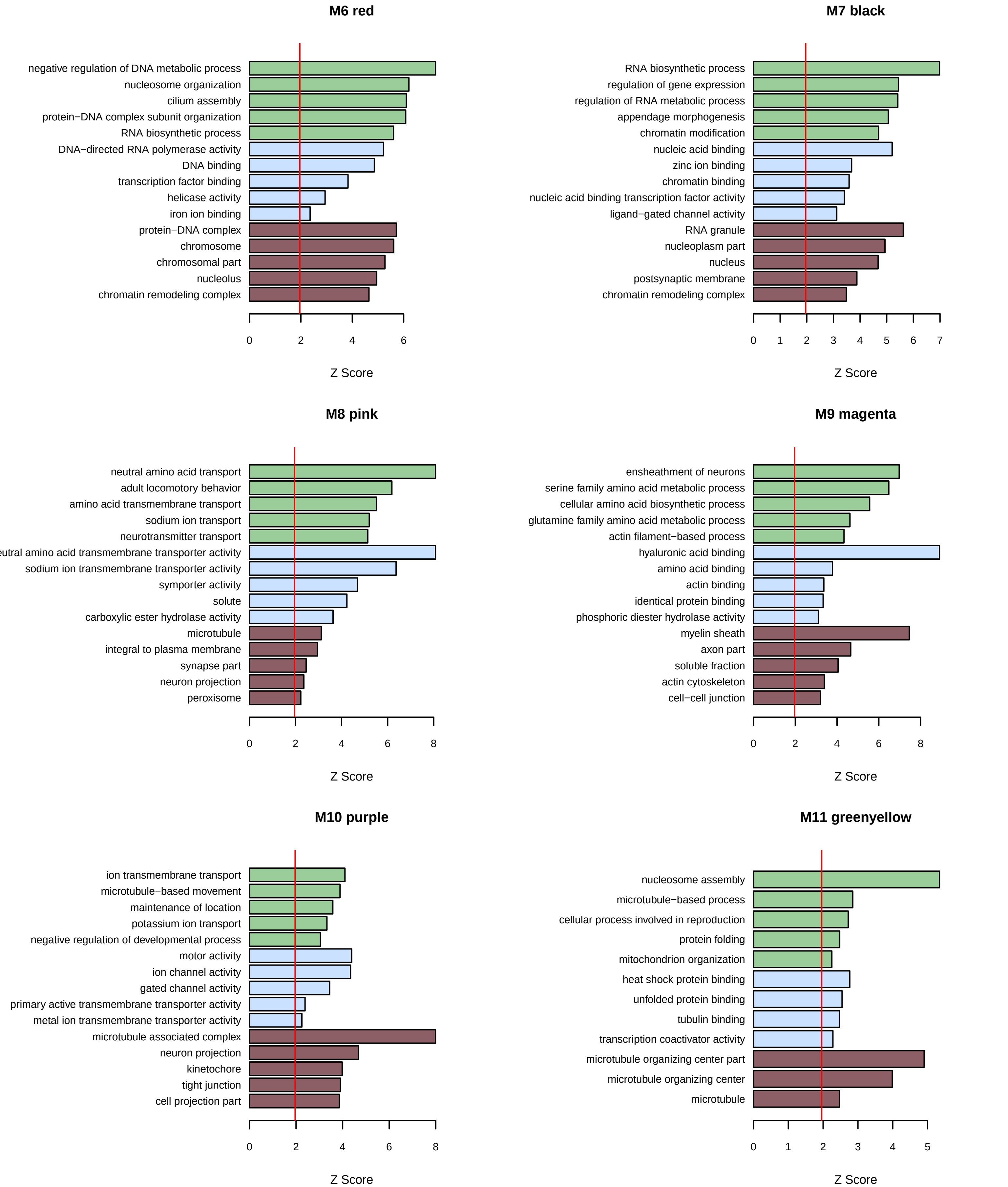
M6 red
negative regulation of DNA metabolic process
nucleosome organization
cilium assembly
protein−DNA complex subunit organization
RNA biosynthetic process
DNA−directed RNA polymerase activity
DNA binding
transcription factor binding
helicase activity
iron ion binding
protein−DNA complex
chromosome
chromosomal part
nucleolus
chromatin remodeling complex
0
2
4
6
Z Score
M7 black
RNA biosynthetic process
regulation of gene expression
regulation of RNA metabolic process
appendage morphogenesis
chromatin modification
nucleic acid binding
zinc ion binding
chromatin binding
nucleic acid binding transcription factor activity
ligand−gated channel activity
RNA granule
nucleoplasm part
nucleus
postsynaptic membrane
chromatin remodeling complex
0
1
2
3
4
5
6
7
Z Score
M8 pink
neutral amino acid transport
adult locomotory behavior
amino acid transmembrane transport
sodium ion transport
neurotransmitter transport
eutral amino acid transmembrane transporter activity
sodium ion transmembrane transporter activity
symporter activity
solute
carboxylic ester hydrolase activity
microtubule
integral to plasma membrane
synapse part
neuron projection
peroxisome
0
2
4
6
8
Z Score
M9 magenta
ensheathment of neurons
serine family amino acid metabolic process
cellular amino acid biosynthetic process
glutamine family amino acid metabolic process
actin filament−based process
hyaluronic acid binding
amino acid binding
actin binding
identical protein binding
phosphoric diester hydrolase activity
myelin sheath
axon part
soluble fraction
actin cytoskeleton
cell−cell junction
0
2
4
6
8
Z Score
M10 purple
ion transmembrane transport
microtubule−based movement
maintenance of location
potassium ion transport
negative regulation of developmental process
motor activity
ion channel activity
gated channel activity
primary active transmembrane transporter activity
metal ion transmembrane transporter activity
microtubule associated complex
neuron projection
kinetochore
tight junction
cell projection part
0
2
4
6
8
Z Score
M11 greenyellow
nucleosome assembly
microtubule−based process
cellular process involved in reproduction
protein folding
mitochondrion organization
heat shock protein binding
unfolded protein binding
tubulin binding
transcription coactivator activity
microtubule organizing center part
microtubule organizing center
microtubule
0
1
2
3
4
5
Z Score

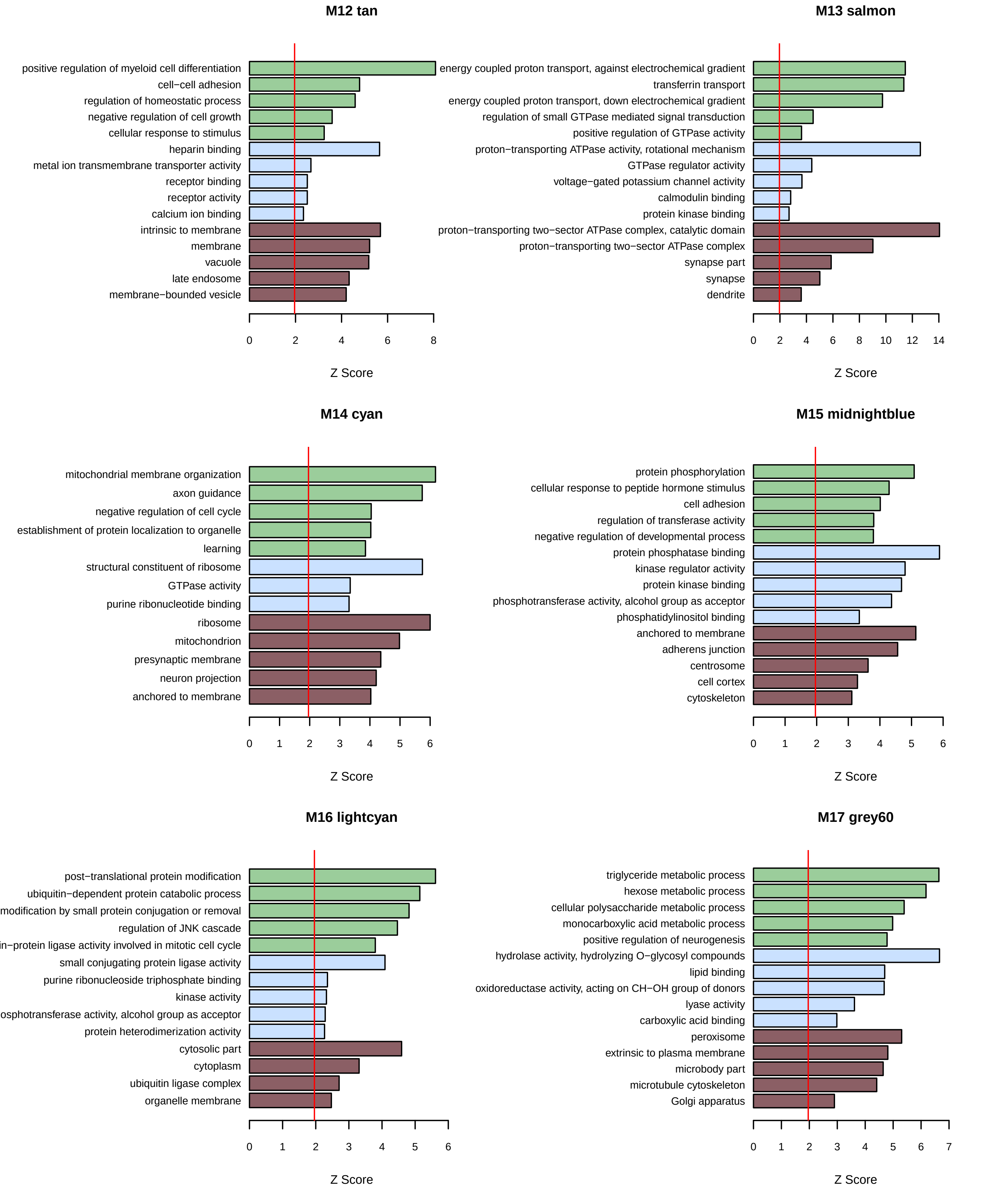
M12 tan
positive regulation of myeloid cell differentiation
cell−cell adhesion
regulation of homeostatic process
negative regulation of cell growth
cellular response to stimulus
heparin binding
metal ion transmembrane transporter activity
receptor binding
receptor activity
calcium ion binding
intrinsic to membrane
membrane
vacuole
late endosome
membrane−bounded vesicle
0 2 4 6 8
Z Score
M13 salmon
energy coupled proton transport, against electrochemical gradient
transferrin transport
energy coupled proton transport, down electrochemical gradient
regulation of small GTPase mediated signal transduction
positive regulation of GTPase activity
proton−transporting ATPase activity, rotational mechanism
GTPase regulator activity
voltage−gated potassium channel activity
calmodulin binding
protein kinase binding
proton−transporting two−sector ATPase complex, catalytic domain
proton−transporting two−sector ATPase complex
synapse part
synapse
dendrite
0 2 4 6 8 10 12 14
Z Score
M14 cyan
mitochondrial membrane organization
axon guidance
negative regulation of cell cycle
establishment of protein localization to organelle
learning
structural constituent of ribosome
GTPase activity
purine ribonucleotide binding
ribosome
mitochondrion
presynaptic membrane
neuron projection
anchored to membrane
0 1 2 3 4 5 6
Z Score
M15 midnightblue
protein phosphorylation
cellular response to peptide hormone stimulus
cell adhesion
regulation of transferase activity
negative regulation of developmental process
protein phosphatase binding
kinase regulator activity
protein kinase binding
phosphotransferase activity, alcohol group as acceptor
phosphatidylinositol binding
anchored to membrane
adherens junction
centrosome
cell cortex
cytoskeleton
0 1 2 3 4 5 6
Z Score
M16 lightcyan
post−translational protein modification
ubiquitin−dependent protein catabolic process
modification by small protein conjugation or removal
regulation of JNK cascade
in−protein ligase activity involved in mitotic cell cycle
small conjugating protein ligase activity
purine ribonucleoside triphosphate binding
kinase activity
osphotransferase activity, alcohol group as acceptor
protein heterodimerization activity
cytosolic part
cytoplasm
ubiquitin ligase complex
organelle membrane
0 1 2 3 4 5 6
Z Score
M17 grey60
triglyceride metabolic process
hexose metabolic process
cellular polysaccharide metabolic process
monocarboxylic acid metabolic process
positive regulation of neurogenesis
hydrolase activity, hydrolyzing O−glycosyl compounds
lipid binding
oxidoreductase activity, acting on CH−OH group of donors
lyase activity
carboxylic acid binding
peroxisome
extrinsic to plasma membrane
microbody part
microtubule cytoskeleton
Golgi apparatus
0 1 2 3 4 5 6 7
Z Score

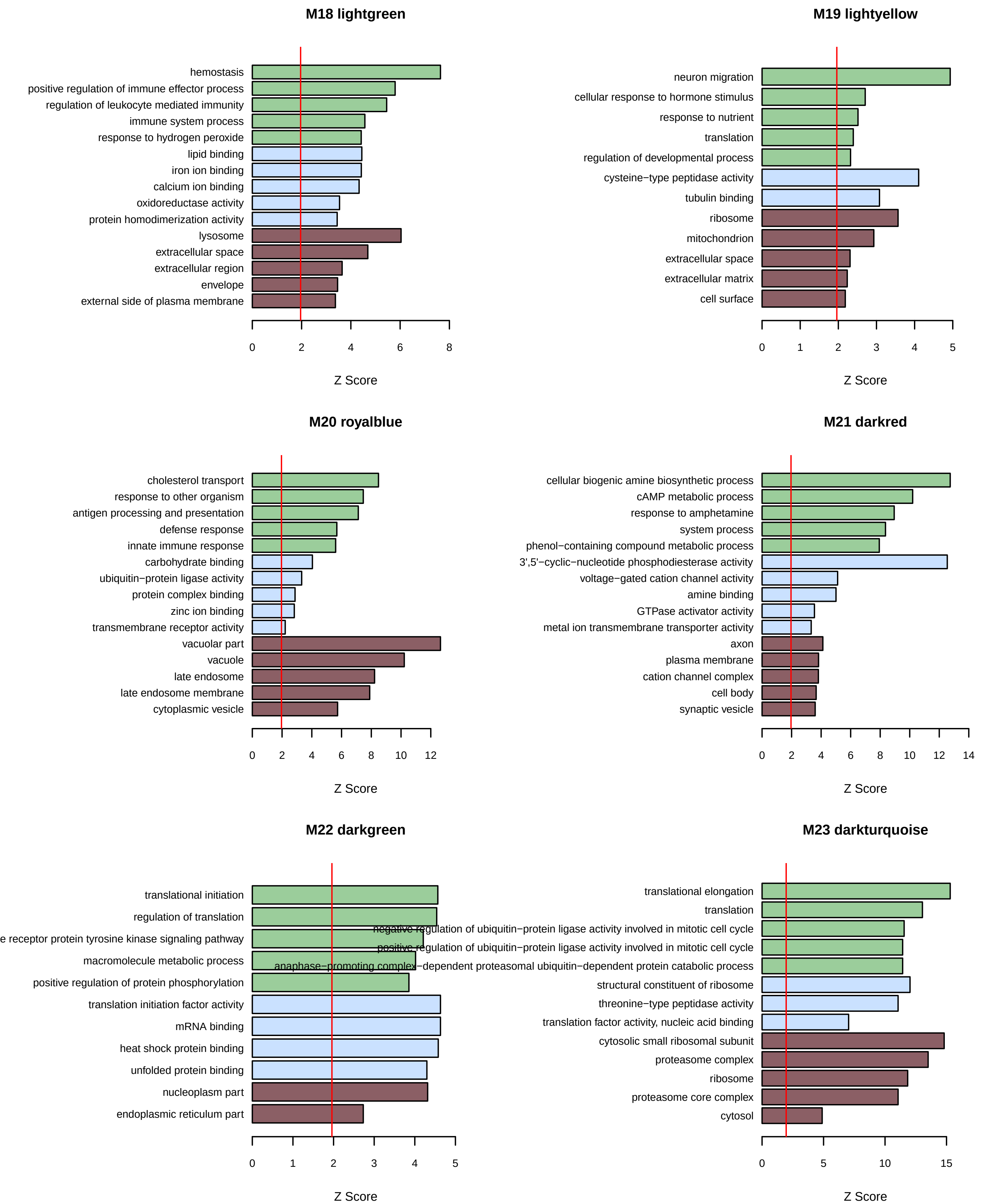
M18 lightgreen
hemostasis
positive regulation of immune effector process
regulation of leukocyte mediated immunity
immune system process
response to hydrogen peroxide
lipid binding
iron ion binding
calcium ion binding
oxidoreductase activity
protein homodimerization activity
lysosome
extracellular space
extracellular region
envelope
external side of plasma membrane
0 2 4 6 8
Z Score
M19 lightyellow
neuron migration
cellular response to hormone stimulus
response to nutrient
translation
regulation of developmental process
cysteine−type peptidase activity
tubulin binding
ribosome
mitochondrion
extracellular space
extracellular matrix
cell surface
0 1 2 3 4 5
Z Score
M20 royalblue
cholesterol transport
response to other organism
antigen processing and presentation
defense response
innate immune response
carbohydrate binding
ubiquitin−protein ligase activity
protein complex binding
zinc ion binding
transmembrane receptor activity
vacuolar part
vacuole
late endosome
late endosome membrane
cytoplasmic vesicle
0 2 4 6 8 10 12
Z Score
M21 darkred
cellular biogenic amine biosynthetic process
cAMP metabolic process
response to amphetamine
system process
phenol−containing compound metabolic process
3',5'−cyclic−nucleotide phosphodiesterase activity
voltage−gated cation channel activity
amine binding
GTPase activator activity
metal ion transmembrane transporter activity
axon
plasma membrane
cation channel complex
cell body
synaptic vesicle
0 2 4 6 8 10 12 14
Z Score
M22 darkgreen
translational initiation
regulation of translation
receptor protein tyrosine kinase signaling pathway
macromolecule metabolic process
positive regulation of protein phosphorylation
translation initiation factor activity
mRNA binding
heat shock protein binding
unfolded protein binding
nucleoplasm part
endoplasmic reticulum part
0 1 2 3 4 5
Z Score
M23 darkturquoise
translational elongation
translation
negative regulation of ubiquitin−protein ligase activity involved in mitotic cell cycle
positive regulation of ubiquitin−protein ligase activity involved in mitotic cell cycle
anaphase−promoting complex−dependent proteasomal ubiquitin−dependent protein catabolic process
structural constituent of ribosome
threonine−type peptidase activity
translation factor activity, nucleic acid binding
cytosolic small ribosomal subunit
proteasome complex
ribosome
proteasome core complex
cytosol
0 5 10 15
Z Score

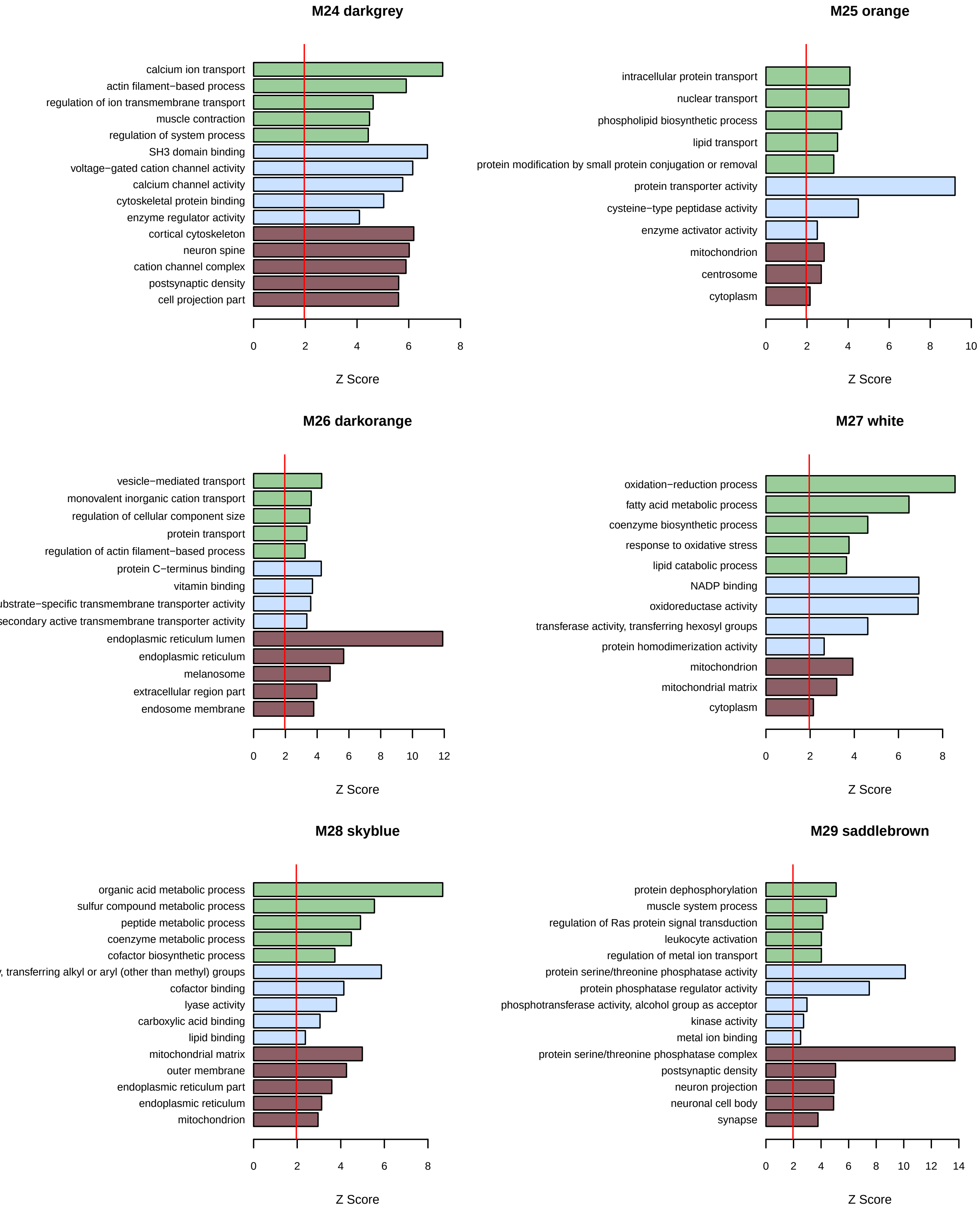

M24 darkgrey
calcium ion transport
actin filament−based process
regulation of ion transmembrane transport
muscle contraction
regulation of system process
SH3 domain binding
voltage−gated cation channel activity
calcium channel activity
cytoskeletal protein binding
enzyme regulator activity
cortical cytoskeleton
neuron spine
cation channel complex
postsynaptic density
cell projection part
0 2 4 6 8
Z Score
M25 orange
intracellular protein transport
nuclear transport
phospholipid biosynthetic process
lipid transport
protein modification by small protein conjugation or removal
protein transporter activity
cysteine−type peptidase activity
enzyme activator activity
mitochondrion
centrosome
cytoplasm
0 2 4 6 8 10
Z Score
M26 darkorange
vesicle−mediated transport
monovalent inorganic cation transport
regulation of cellular component size
protein transport
regulation of actin filament−based process
protein C−terminus binding
vitamin binding
ubstrate−specific transmembrane transporter activity
secondary active transmembrane transporter activity
endoplasmic reticulum lumen
endoplasmic reticulum
melanosome
extracellular region part
endosome membrane
0 2 4 6 8 10 12
Z Score
M27 white
oxidation−reduction process
fatty acid metabolic process
coenzyme biosynthetic process
response to oxidative stress
lipid catabolic process
NADP binding
oxidoreductase activity
transferase activity, transferring hexosyl groups
protein homodimerization activity
mitochondrion
mitochondrial matrix
cytoplasm
0 2 4 6 8
Z Score
M28 skyblue
organic acid metabolic process
sulfur compound metabolic process
peptide metabolic process
coenzyme metabolic process
cofactor biosynthetic process
, transferring alkyl or aryl (other than methyl) groups
cofactor binding
lyase activity
carboxylic acid binding
lipid binding
mitochondrial matrix
outer membrane
endoplasmic reticulum part
endoplasmic reticulum
mitochondrion
0 2 4 6 8
Z Score
M29 saddlebrown
protein dephosphorylation
muscle system process
regulation of Ras protein signal transduction
leukocyte activation
regulation of metal ion transport
protein serine/threonine phosphatase activity
protein phosphatase regulator activity
phosphotransferase activity, alcohol group as acceptor
kinase activity
metal ion binding
protein serine/threonine phosphatase complex
postsynaptic density
neuron projection
neuronal cell body
synapse
0 2 4 6 8 10 12 14
Z Score

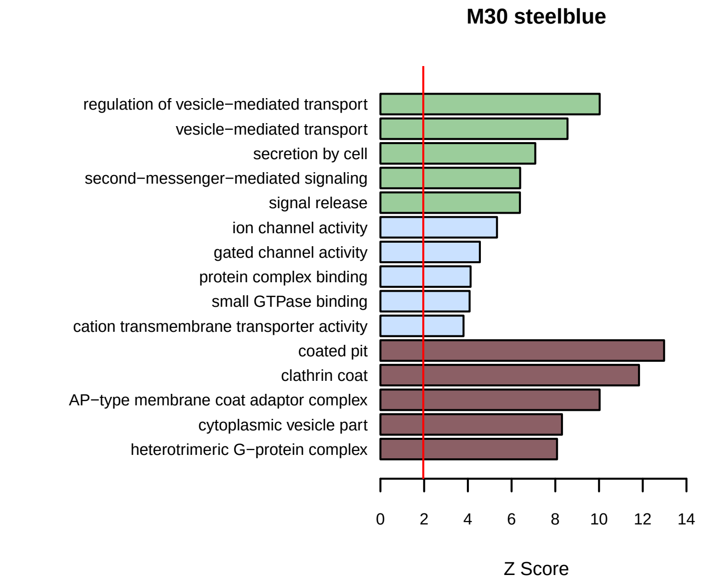
M30 steelblue
regulation of vesicle−mediated transport
vesicle−mediated transport
secretion by cell
second−messenger−mediated signaling
signal release
ion channel activity
gated channel activity
protein complex binding
small GTPase binding
cation transmembrane transporter activity
coated pit
clathrin coat
AP−type membrane coat adaptor complex
cytoplasmic vesicle part
heterotrimeric G−protein complex
0 2 4 6 8 10 12 14
Z Score

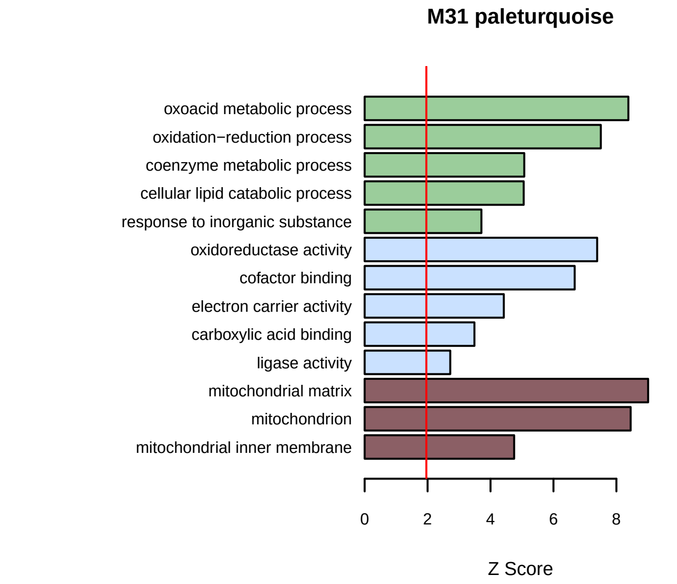
M31 paleturquoise
oxoacid metabolic process
oxidation−reduction process
coenzyme metabolic process
cellular lipid catabolic process
response to inorganic substance
oxidoreductase activity
cofactor binding
electron carrier activity
carboxylic acid binding
ligase activity
mitochondrial matrix
mitochondrion
mitochondrial inner membrane
0 2 4 6 8
Z Score

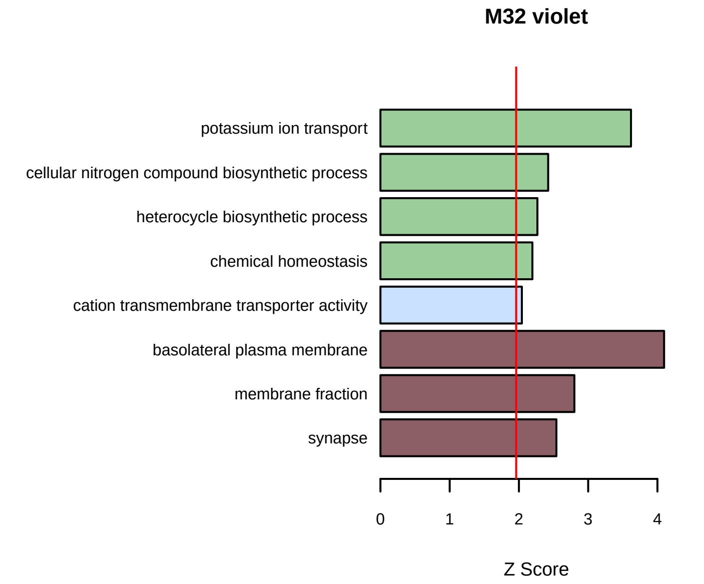
M32 violet
potassium ion transport
cellular nitrogen compound biosynthetic process
heterocycle biosynthetic process
chemical homeostasis
cation transmembrane transporter activity
basolateral plasma membrane
membrane fraction
synapse
0 1 2 3 4
Z Score

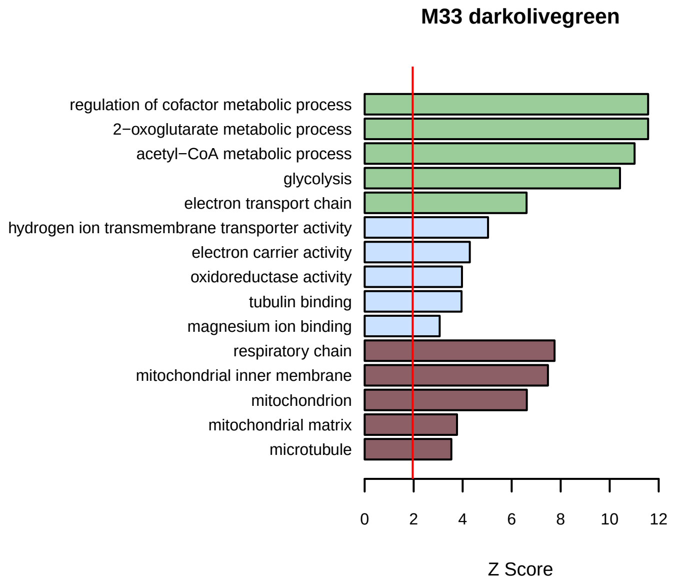
M33 darkolivegreen
regulation of cofactor metabolic process
2−oxoglutarate metabolic process
acetyl−CoA metabolic process
glycolysis
electron transport chain
hydrogen ion transmembrane transporter activity
electron carrier activity
oxidoreductase activity
tubulin binding
magnesium ion binding
respiratory chain
mitochondrial inner membrane
mitochondrion
mitochondrial matrix
microtubule
0 2 4 6 8 10 12
Z Score

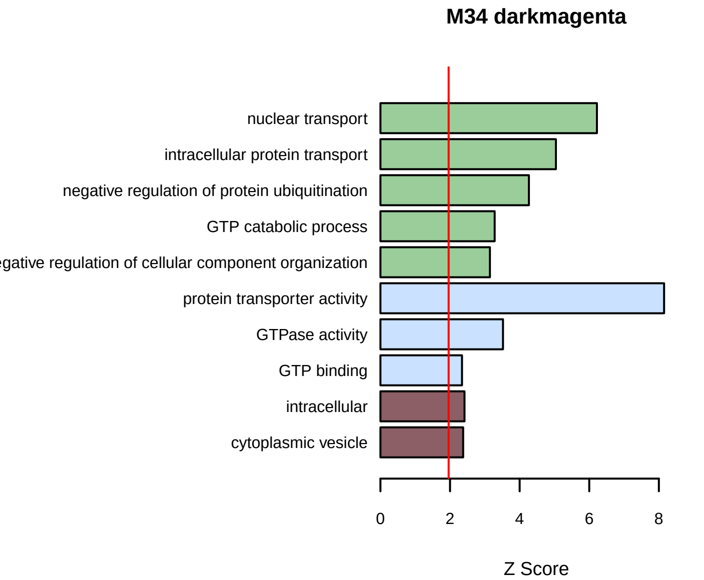
M34 darkmagenta
nuclear transport
intracellular protein transport
negative regulation of protein ubiquitination
GTP catabolic process
egative regulation of cellular component organization
protein transporter activity
GTPase activity
GTP binding
intracellular
cytoplasmic vesicle
0 2 4 6 8
Z Score

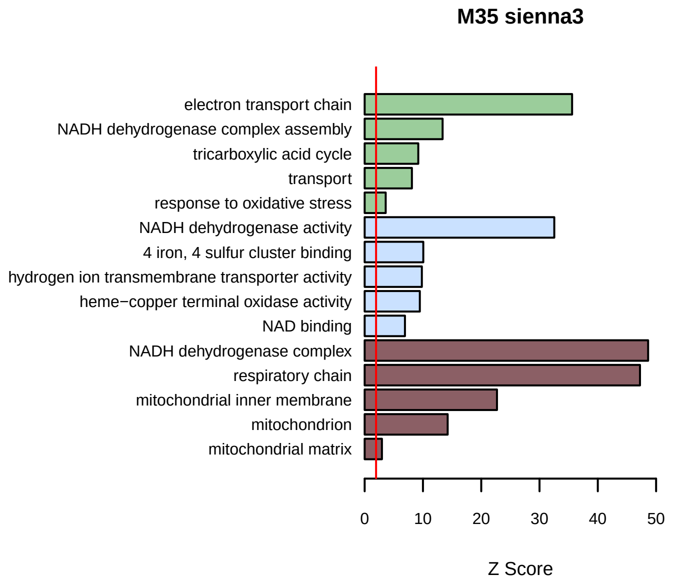
M35 sienna3
electron transport chain
NADH dehydrogenase complex assembly
tricarboxylic acid cycle
transport
response to oxidative stress
NADH dehydrogenase activity
4 iron, 4 sulfur cluster binding
hydrogen ion transmembrane transporter activity
heme−copper terminal oxidase activity
NAD binding
NADH dehydrogenase complex
respiratory chain
mitochondrial inner membrane
mitochondrion
mitochondrial matrix
0 10 20 30 40 50
Z Score

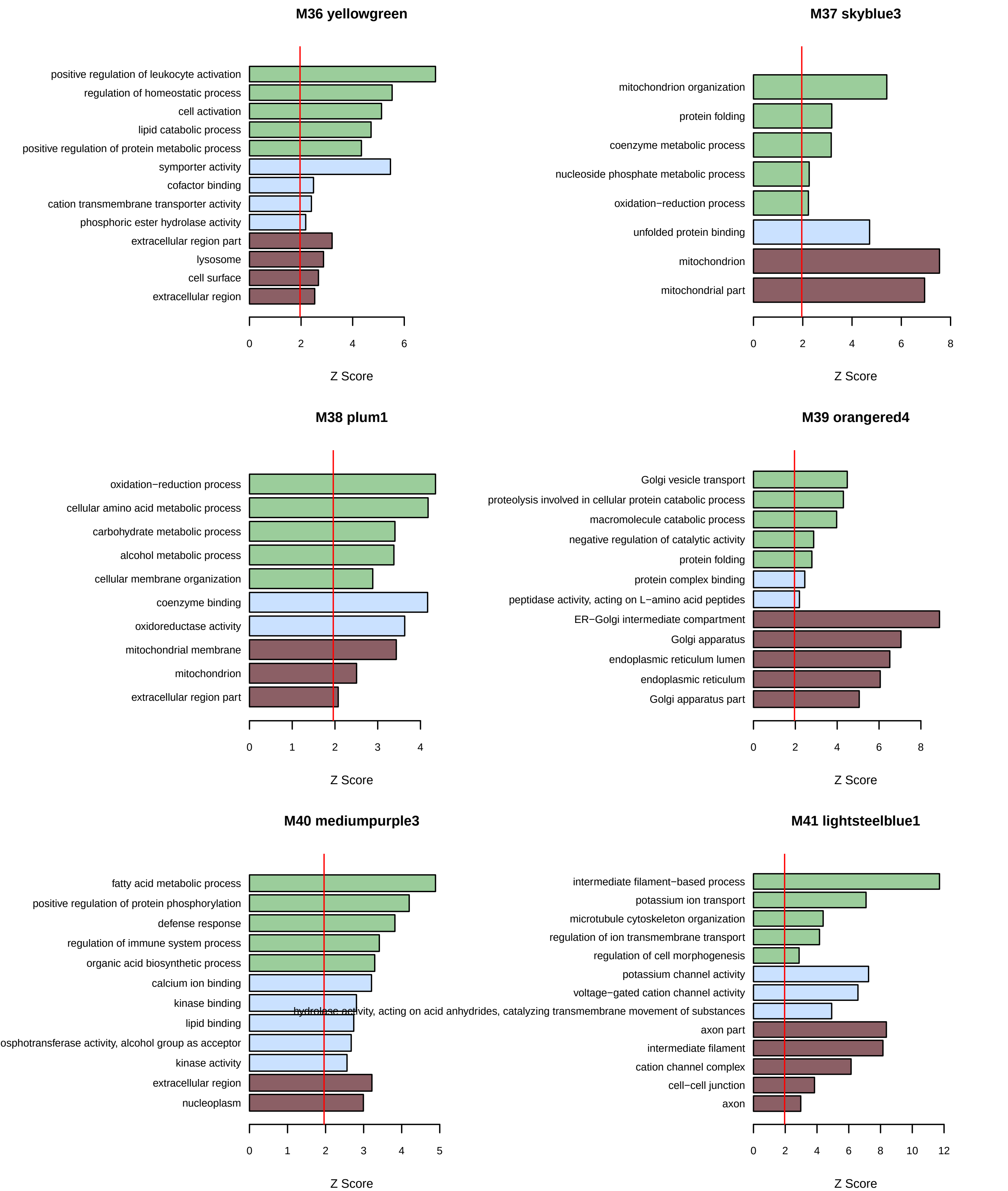
M36 yellowgreen
positive regulation of leukocyte activation
regulation of homeostatic process
cell activation
lipid catabolic process
positive regulation of protein metabolic process
symporter activity
cofactor binding
cation transmembrane transporter activity
phosphoric ester hydrolase activity
extracellular region part
lysosome
cell surface
extracellular region
0
2
4
6
Z Score
M37 skyblue3
mitochondrion organization
protein folding
coenzyme metabolic process
nucleoside phosphate metabolic process
oxidation−reduction process
unfolded protein binding
mitochondrion
mitochondrial part
0
2
4
6
8
Z Score
M38 plum1
oxidation−reduction process
cellular amino acid metabolic process
carbohydrate metabolic process
alcohol metabolic process
cellular membrane organization
coenzyme binding
oxidoreductase activity
mitochondrial membrane
mitochondrion
extracellular region part
0
1
2
3
4
Z Score
M39 orangered4
Golgi vesicle transport
proteolysis involved in cellular protein catabolic process
macromolecule catabolic process
negative regulation of catalytic activity
protein folding
protein complex binding
peptidase activity, acting on L−amino acid peptides
ER−Golgi intermediate compartment
Golgi apparatus
endoplasmic reticulum lumen
endoplasmic reticulum
Golgi apparatus part
0
2
4
6
8
Z Score
M40 mediumpurple3
fatty acid metabolic process
positive regulation of protein phosphorylation
defense response
regulation of immune system process
organic acid biosynthetic process
calcium ion binding
kinase binding
lipid binding
osphotransferase activity, alcohol group as acceptor
kinase activity
extracellular region
nucleoplasm
0
1
2
3
4
5
Z Score
M41 lightsteelblue1
intermediate filament−based process
potassium ion transport
microtubule cytoskeleton organization
regulation of ion transmembrane transport
regulation of cell morphogenesis
potassium channel activity
voltage−gated cation channel activity
hydrolase activity, acting on acid anhydrides, catalyzing transmembrane movement of substances
axon part
intermediate filament
cation channel complex
cell−cell junction
axon
0
2
4
6
8
10
12
Z Score

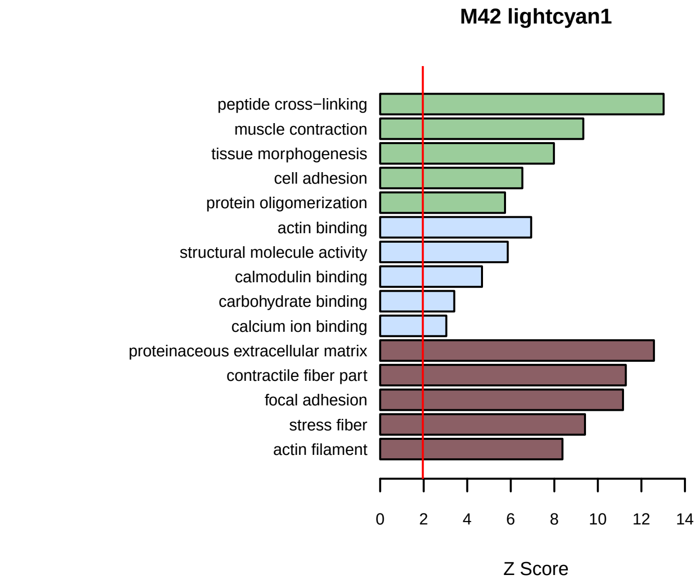
M42 lightcyan1
peptide cross−linking
muscle contraction
tissue morphogenesis
cell adhesion
protein oligomerization
actin binding
structural molecule activity
calmodulin binding
carbohydrate binding
calcium ion binding
proteinaceous extracellular matrix
contractile fiber part
focal adhesion
stress fiber
actin filament
0 2 4 6 8 10 12 14
Z Score

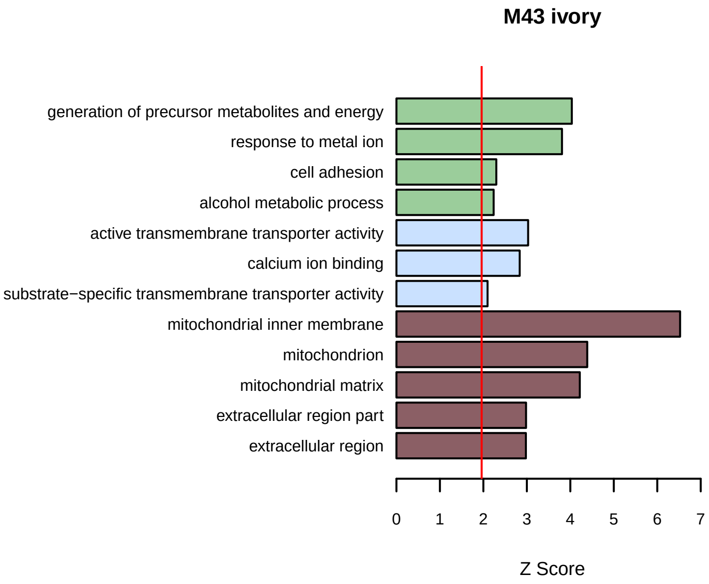
M43 ivory
generation of precursor metabolites and energy
response to metal ion
cell adhesion
alcohol metabolic process
active transmembrane transporter activity
calcium ion binding
substrate−specific transmembrane transporter activity
mitochondrial inner membrane
mitochondrion
mitochondrial matrix
extracellular region part
extracellular region
0 1 2 3 4 5 6 7
Z Score

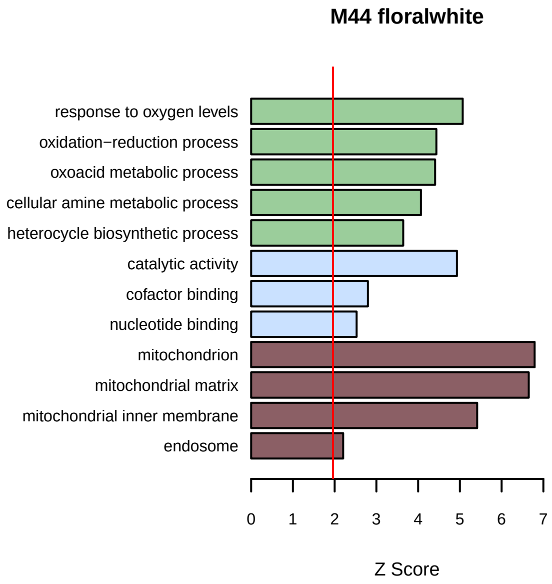
M44 floralwhite
response to oxygen levels
oxidation−reduction process
oxoacid metabolic process
cellular amine metabolic process
heterocycle biosynthetic process
catalytic activity
cofactor binding
nucleotide binding
mitochondrion
mitochondrial matrix
mitochondrial inner membrane
endosome
0 1 2 3 4 5 6 7
Z Score

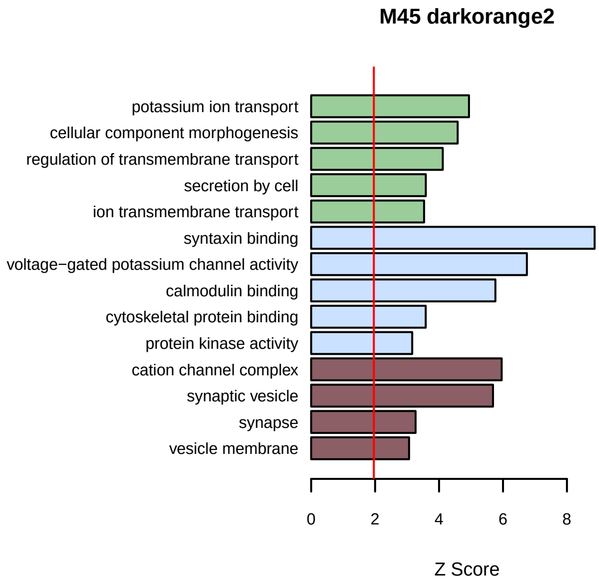
M45 darkorange2
potassium ion transport
cellular component morphogenesis
regulation of transmembrane transport
secretion by cell
ion transmembrane transport
syntaxin binding
voltage−gated potassium channel activity
calmodulin binding
cytoskeletal protein binding
protein kinase activity
cation channel complex
synaptic vesicle
synapse
vesicle membrane
0 2 4 6 8
Z Score

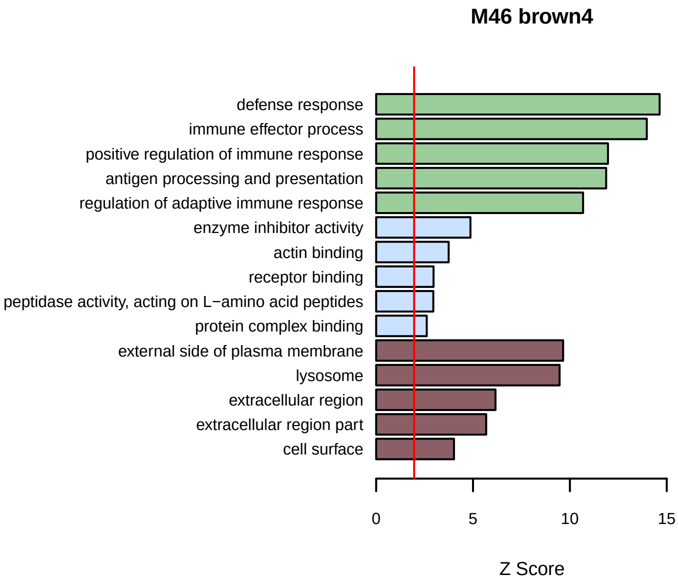
M46 brown4
defense response
immune effector process
positive regulation of immune response
antigen processing and presentation
regulation of adaptive immune response
enzyme inhibitor activity
actin binding
receptor binding
peptidase activity, acting on L−amino acid peptides
protein complex binding
external side of plasma membrane
lysosome
extracellular region
extracellular region part
cell surface
0 5 10 15
Z Score

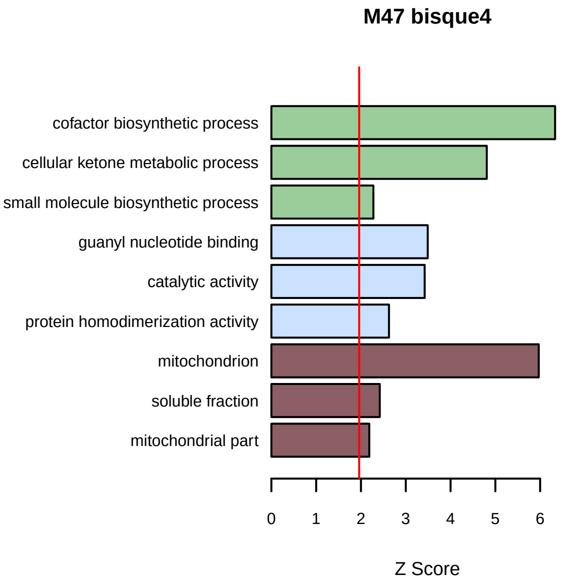
M47 bisque4
cofactor biosynthetic process
cellular ketone metabolic process
small molecule biosynthetic process
guanyl nucleotide binding
catalytic activity
protein homodimerization activity
mitochondrion
soluble fraction
mitochondrial part
0 1 2 3 4 5 6
Z Score

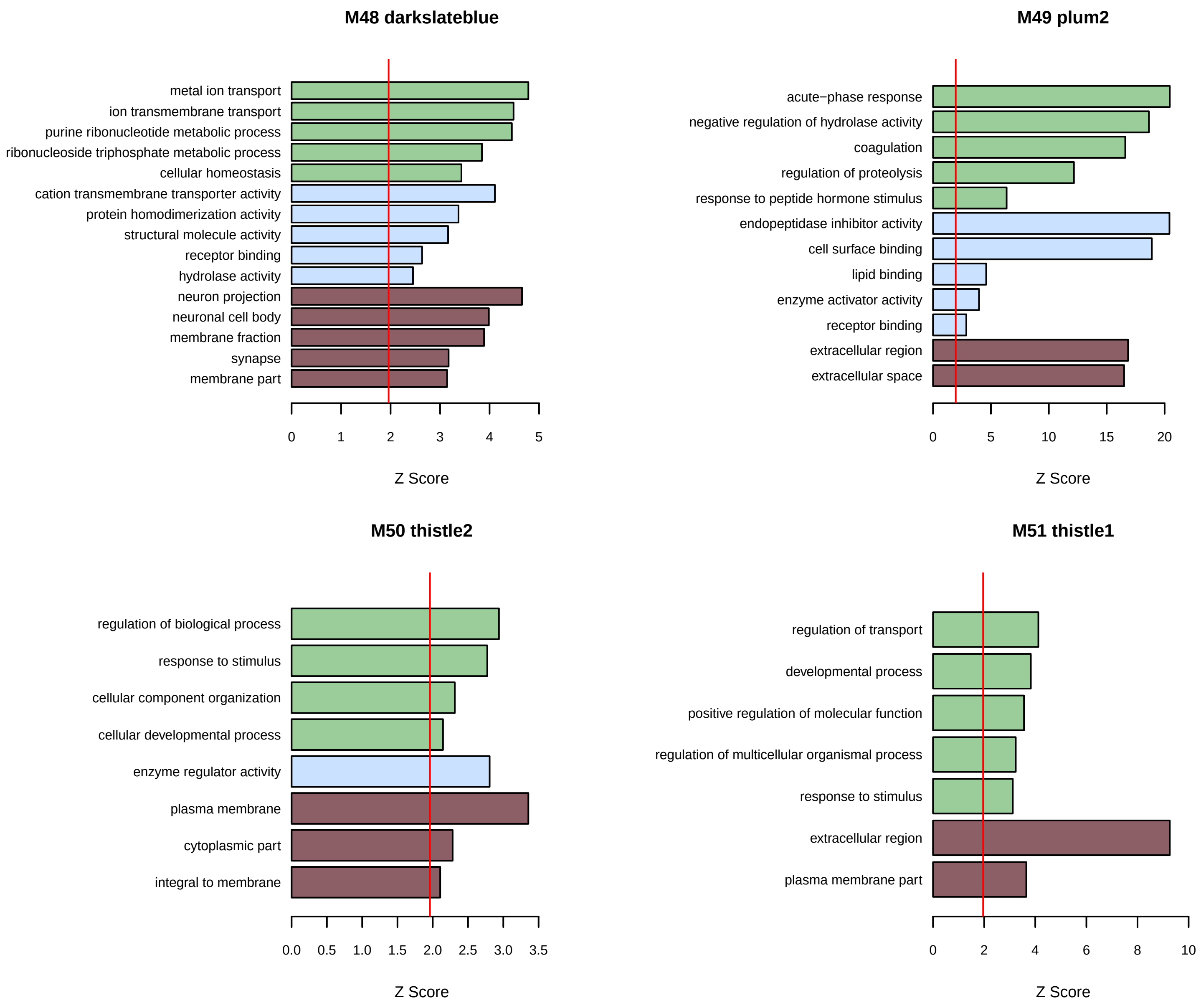
M48 darkslateblue
metal ion transport
ion transmembrane transport
purine ribonucleotide metabolic process
ribonucleoside triphosphate metabolic process
cellular homeostasis
cation transmembrane transporter activity
protein homodimerization activity
structural molecule activity
receptor binding
hydrolase activity
neuron projection
neuronal cell body
membrane fraction
synapse
membrane part
0 1 2 3 4 5
Z Score
M49 plum2
acute−phase response
negative regulation of hydrolase activity
coagulation
regulation of proteolysis
response to peptide hormone stimulus
endopeptidase inhibitor activity
cell surface binding
lipid binding
enzyme activator activity
receptor binding
extracellular region
extracellular space
0 5 10 15 20
Z Score
M50 thistle2
regulation of biological process
response to stimulus
cellular component organization
cellular developmental process
enzyme regulator activity
plasma membrane
cytoplasmic part
integral to membrane
0.0 0.5 1.0 1.5 2.0 2.5 3.0 3.5
Z Score
M51 thistle1
regulation of transport
developmental process
positive regulation of molecular function
regulation of multicellular organismal process
response to stimulus
extracellular region
plasma membrane part
0 2 4 6 8 10
Z Score
