## Supplementary Figures for "Network analysis of mouse brain proteomes resolves molecular and cellular endophenotypes of aging and Aβ pathology"

Supplementary Figures: 6

Supplementary Datasheets: 3

Supplementary Tables: 12

Keywords: aging, proteomics, amyloid-beta, mitochondria, metabolism, synapse, translation

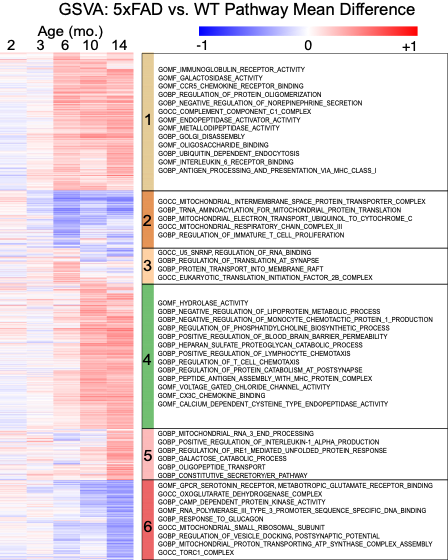
**Supplementary Figure 1. Gene-set variation analysis of the 5xFAD brain proteome across age.** Heatmap of the mean difference in GSVA enrichment score between 5xFAD and WT mice at 2, 3, 6, 10 and 14 months, computed over gene ontology gene sets. Red, higher in 5xFAD; blue, lower. Rows are hierarchically clustered into six groups, numbered and color-coded at left; representative gene sets for each group are listed at right. Immune, complement, antigen-presentation and lysosomal sets (groups 1 and 4) rise progressively from 6 months, while mitochondrial, respiratory-chain and synaptic sets (groups 2 and 6) fall at the oldest ages.

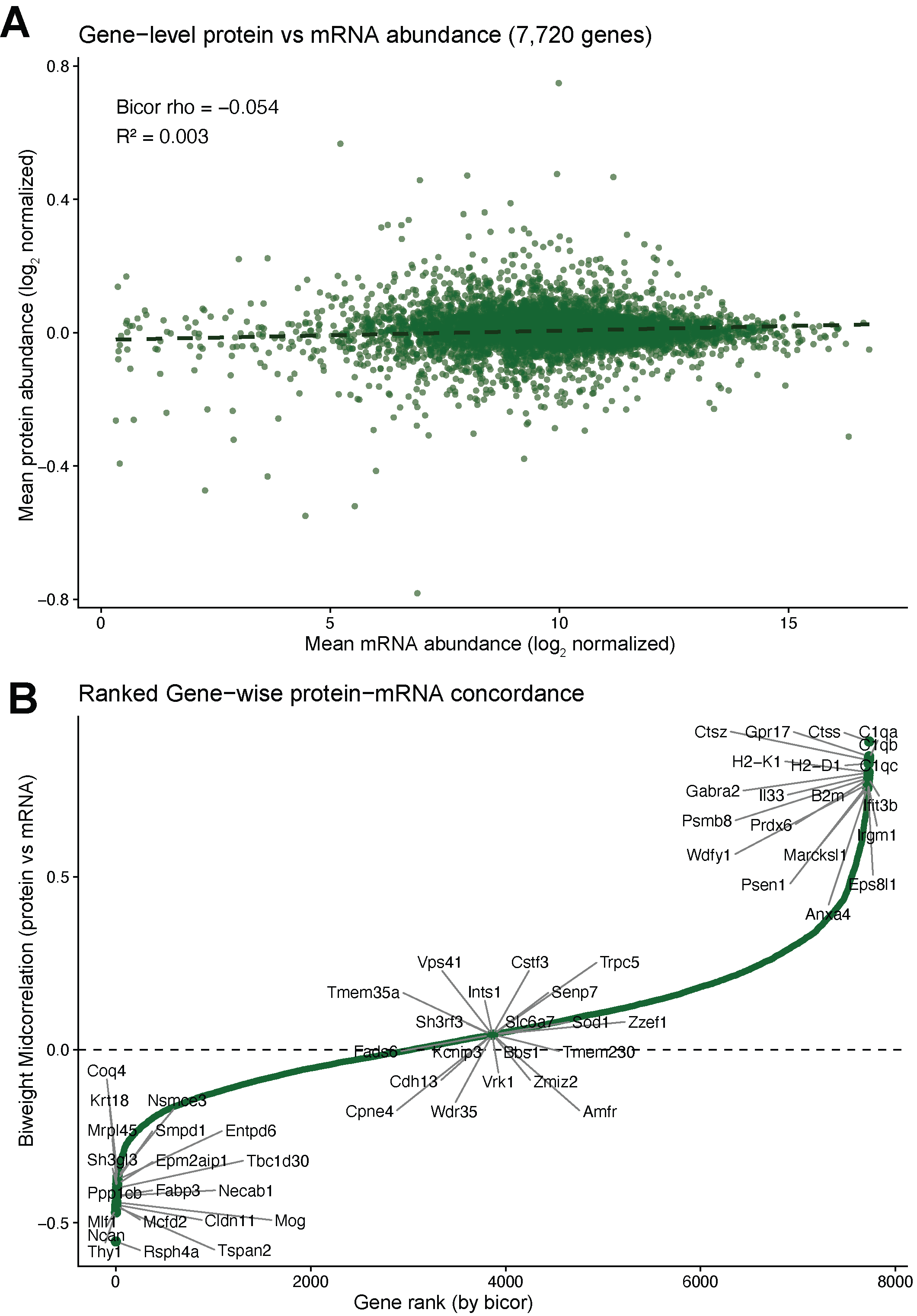

**Supplementary Figure 2. Protein and mRNA abundance are largely uncorrelated at the gene level.** (A) Mean protein abundance versus mean mRNA abundance across the 7,720 genes with paired proteomic and transcriptomic measurements. Biweight midcorrelation rho = -0.054, R2 = 0.003; dashed line, linear fit. (B) Genes ranked by gene-wise protein-mRNA biweight midcorrelation. Genes at the positive and negative extremes of the distribution are labelled. The most concordant genes are dominated by complement, MHC class I and microglial transcripts; the most discordant include myelin and neuronal surface proteins.

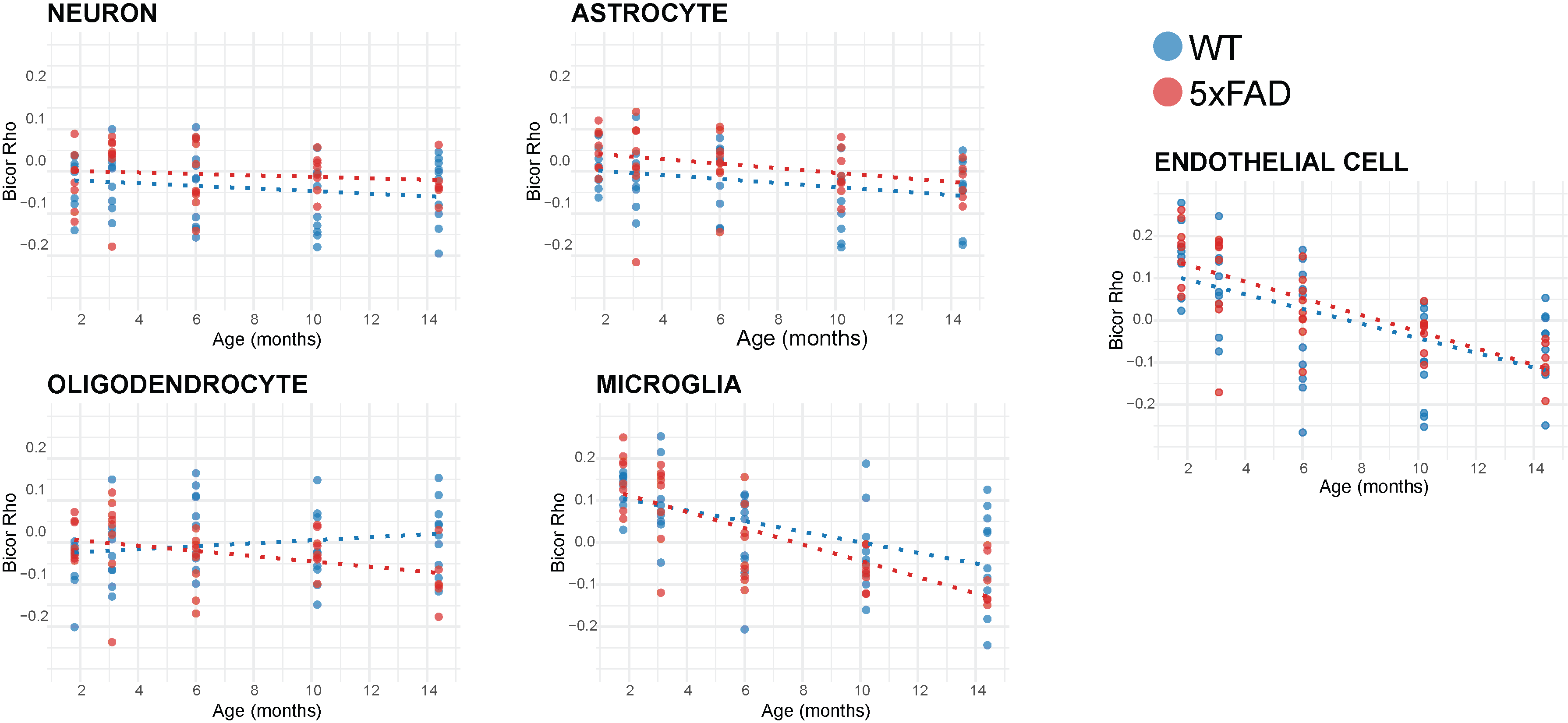

**Supplementary Figure 3. Protein-mRNA concordance declines with age in a cell type-specific manner.** Gene-wise protein-mRNA correlation (biweight midcorrelation rho) in each mouse, computed separately over neuronal, astrocytic, oligodendrocytic, microglial and endothelial marker genes, plotted against age. WT blue, 5xFAD red; dashed lines are per-genotype linear fits. Panels expand the cell-type trajectories summarized in Fig. 2F; endothelial and microglial concordance declines most steeply with age.

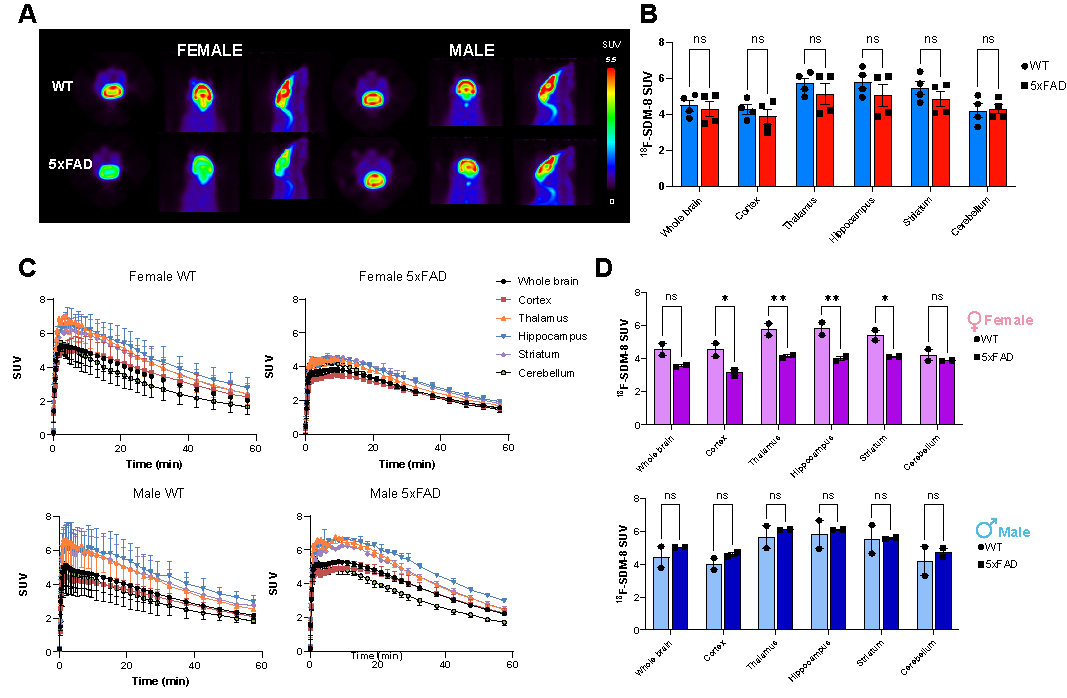
**Supplementary Figure 4. Synaptic density measured by SV2A PET is unchanged in 5xFAD mice when sexes are combined.** (A) Representative [18F]SDM-8 PET images in female and male WT and 5xFAD mice, shown in three orthogonal planes on a common SUV scale. (B) Regional [18F]SDM-8 standardized uptake value in whole brain, cortex, thalamus, hippocampus, striatum and cerebellum with sexes combined. No region differs by genotype. (C) Regional time-activity curves across the 60-min scan for female and male WT and 5xFAD mice. (D) Regional SUV analyzed separately in females (top) and males (bottom). Significant genotype differences are confined to female mice. ns, not significant; *P < 0.05; **P < 0.01. Data are mean +/- SEM with individual mice shown.

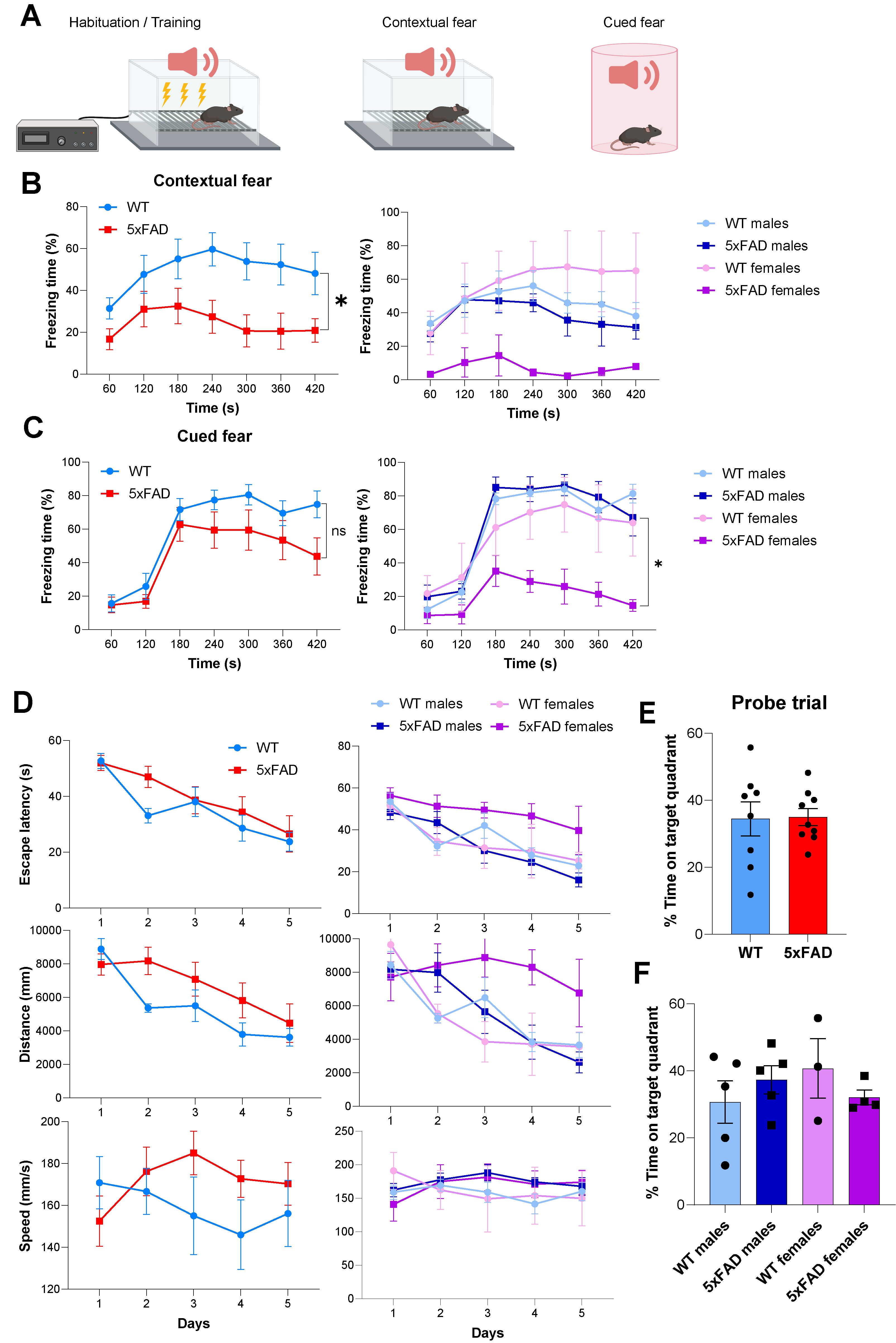

**Supplementary Figure 5. 5xFAD mice show impaired contextual fear memory with preserved spatial learning.** (A) Fear-conditioning paradigm: tone-shock training followed by contextual and cued fear testing. (B) Contextual and (C) cued fear conditioning. Freezing across the test session in WT and 5xFAD mice (left) and split by sex (right). (D) Morris water maze acquisition across five training days: escape latency, path length and swim speed, by genotype (left) and by sex (right). (E) Morris water maze probe trial: time spent in the target quadrant, by genotype. (F) Probe trial performance split by sex. Data are mean +/- SEM with individual mice shown. *P < 0.05.

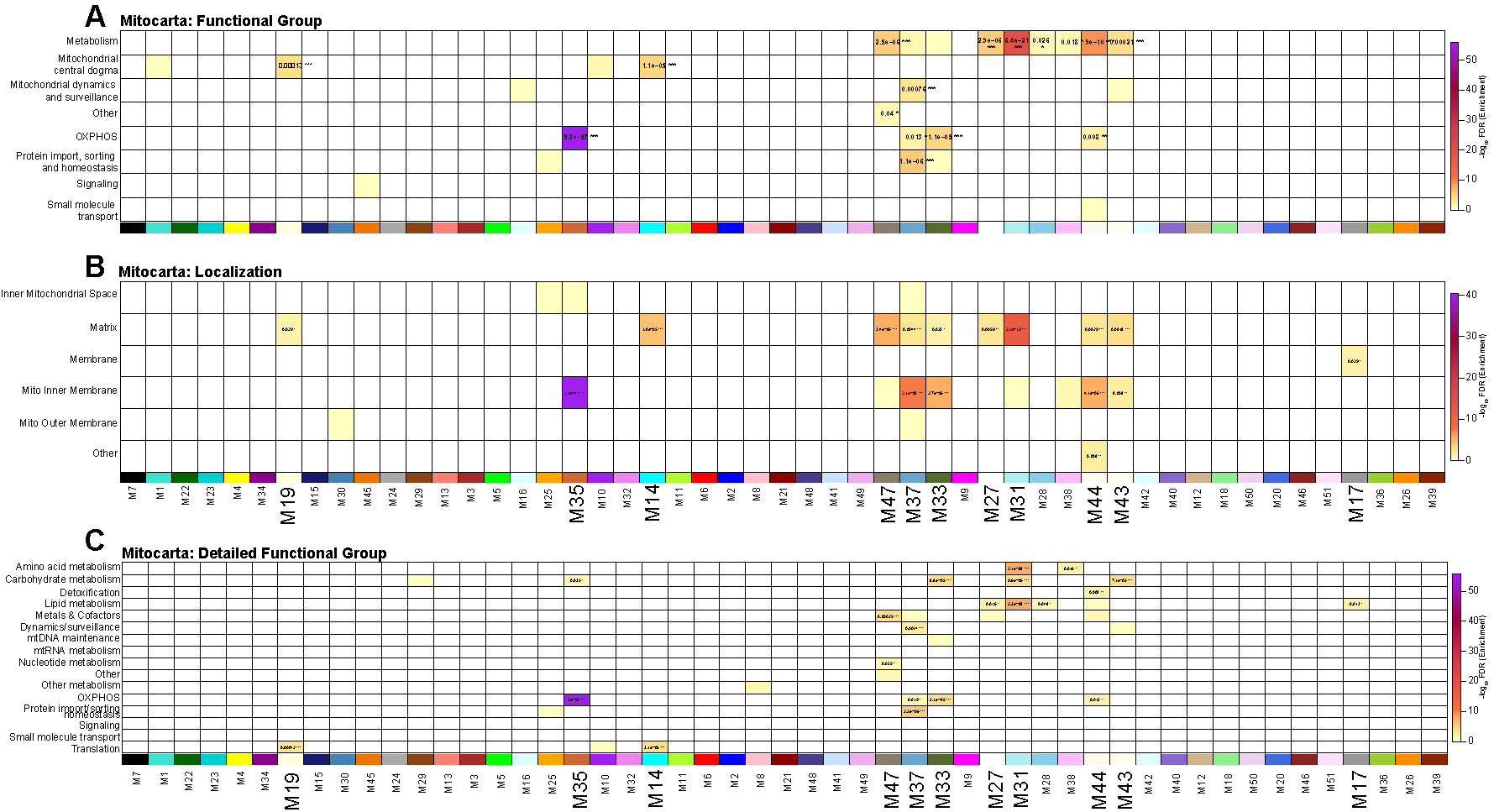

**Supplementary Figure 6. MitoCarta3.0 annotation enrichment across the 51 protein co-expression modules.** Enrichment of each module for MitoCarta3.0 (A) functional groups, (B) sub-mitochondrial localizations and (C) detailed functional categories. One-sided Fisher exact test; color denotes -log10 BH-adjusted P and the adjusted P value is printed in significant cells. Modules are ordered as in Fig. 1F with the module color bar above each heatmap. M35 is by far the most strongly enriched module (OXPHOS, adjusted P = 9.8e-57; inner mitochondrial membrane, adjusted P = 2.9e-41), supporting its assignment as the complex I / respiratory chain module in Fig. 4.
